# TP53 status aligns with morphology and metaplastic programs in colorectal cancer organoids, and MDM2 P2 promoter usage reflects p53 activity

**DOI:** 10.64898/2026.09.21.753357

**Authors:** Takahiro Tsukui, Kento Shin, Ryoji Yao, Koji Tsuda

## Abstract

**Background:** In colorectal cancer (CRC), TP53 mutation accompanies chromosomal instability. How TP53 status relates to tumor morphology and lineage programs, and whether a p53-activity readout could refine MDM2-inhibitor use, are unclear. MDM2 is transcribed from a constitutive (P1) and a p53-inducible (P2) promoter.

**Methods:** Sixty-three organoid samples from 21 CRC patients were classified by deep-learning morphology (VGG16), array-based MDM2 Splicing Index and TP53 targeted resequencing. In organoids, p53 activity was scored from 12 direct p53 targets and promoter usage as the array P2 index. TCGA-COAD/READ and cell lines used RNA-seq transcript ratios (P2_index; cell-line P2 index). Validation used same-patient arrays (GSE128213), TCGA-COAD/READ (n = 624), GSE39582 and an independent biobank (GSE294511).

**Results:** TP53 mutations were almost confined to P1-dominant samples (40 of 42 mutants). Morphology followed TP53 status (median Type1 fraction 0.854 (mutant) versus 0.422, specimen level; P = 0.013 adjusted for chromosome 20q and aneuploidy). With only five samples discordant, TP53 status and promoter usage could not be separated (P = 0.97). The morphology gene score followed TP53 status in 88 GSE294511 organoids (adjusted P = 6.1×10⁻⁴) and TCGA-COAD/READ, identifying mucinous carcinoma; gastric-metaplasia genes followed TP53 status, not P2, in both cohorts but not discovery organoids. The p53-target module of the P2 signature followed P2 beyond TP53 status in both organoid cohorts, whereas the independent 12-gene p53 activity score did so in GSE294511 (P = 0.012) but not discovery organoids (P = 0.094). P2-dominant samples, including two TP53-mutant ones, were p53-active, and nutlin induced P2 only in p53-proficient cells. In TCGA, P2_index was higher in wild-type tumors but did not follow p53 activity beyond total MDM2. In wild-type cell lines P2 followed MDM2 dependency beyond total MDM2, and in wild-type GSE294511 organoids idasanutlin sensitivity, though not beyond total MDM2.

**Conclusions:** In CRC organoids, morphology followed TP53 status, with no detectable contribution of MDM2 promoter usage, as did gastric-metaplastic programs in an independent biobank and tumors. In organoids, P2 usage reflected p53 activity, beyond genotype mainly in the biobank and two TP53-mutant P2-dominant specimens; whether it improves on MDM2 expression for MDM2-inhibitor selection requires testing.

## 1. Background

Molecular subtypes increasingly guide the treatment of colorectal cancer (CRC), but few biomarkers that can be read from a biopsy are linked to a specific drug class. TP53 is mutated in most chromosomally unstable (CIN) CRCs, which frequently gain and lose chromosome segments (Section 3.1.4). Yet TP53 genotype is an imperfect proxy for p53 function, and although TP53 mutations are known to be unevenly distributed across the histological and molecular types of CRC, how far TP53 status itself, rather than the features that accompany it, relates to the morphology and lineage programs of the tumor is not well defined. MDM2, the principal negative regulator of p53 and the target of MDM2 inhibitors (drugs that block MDM2 binding and so release wild-type p53), is transcribed from a constitutive promoter (P1) and from a promoter induced by wild-type p53 (P2).

This study asks how TP53 status, p53 activity and the balance between the two MDM2 promoters (the P1/P2 ratio) relate to two contrasting CRC phenotypes. The first is an autonomously proliferative CIN type whose differentiation core corresponds to consensus molecular subtype 2 (CMS2). The second is an environment-adaptive type with gastric-metaplasia features (acquisition of gastric mucous-cell-like traits by colorectal tumors; Section 3.3.1) that, at the signature level, resembles the microsatellite-instability (MSI)-like program, MSI being hypermutability from defective mismatch repair. We also ask whether P2 usage could serve as a readout (measurable indicator) of p53 activity linked to MDM2-inhibitor therapy.

With an estimated 2.0 million new cases worldwide in 2024, colorectal cancer is the third most commonly diagnosed cancer [1]. CRC is molecularly heterogeneous. The consensus molecular subtype (CMS) classification of Guinney et al. [2] divided CRC into four subtypes with distinct prognosis and therapeutic responsiveness. However, much remains unresolved about the molecular mechanisms that determine why particular subtypes show dramatic phenotypic transformation (metaplasia, a switch to another differentiated cell type) or distinctive phenotypes.

MDM2 has long been known as the principal negative regulator of p53 [3,4], and its transcription is controlled by two major promoters. Wild-type p53 induces MDM2 expression [5]. The P2 promoter carries p53 response elements and is directly induced by the activation of p53 [6,7], whereas P1 drives p53-independent basal transcription. In mice, point mutations in the two p53 response elements of the Mdm2 P2 promoter show that this feedback is dispensable for p53 stability, development and longevity but limits p53 activity after DNA damage [8]. Phelps et al. [9] showed that P2 can also be activated in a p53-independent manner through response elements for multiple transcription factors, revealing a complex mechanism of P2 regulation in cancer cells.

Organoid morphology offers a window onto this molecular basis from the side of the observable phenotype. Organoids are three-dimensional cultures grown from a patient’s tumor or normal tissue. Recent studies have quantified organoid morphology by computational image analysis and linked it to the underlying cell state and drug response (deep learning in breast cancer organoids [10]; high-content image-feature profiling of normal intestinal organoid regeneration [11]). These studies suggest that morphological differences may reflect the underlying molecular programs. From this perspective, the present study started from the question of what molecular axis underlies morphologically contrasting CRC organoids.

We studied two groups of CRC organoids that carry the same diagnosis but differ markedly in morphology and in their MDM2 P1/P2 usage ratio. Our integrative analysis combined deep-learning (VGG16) morphological classification of organoids, MDM2 Splicing Index analysis, gene-expression profiling, multilayer Ingenuity Pathway Analysis (IPA, a commercial knowledge-base tool that infers pathways and upstream regulators from expression changes; all 8 modules), EnrichR Gene Ontology (GO) analysis (EnrichR being a gene-set enrichment analysis tool), and TP53 targeted resequencing (Section 2). The patient-derived organoids analyzed here (21 patients, 63 samples, the HCT patient group) derive from the same patient-derived organoid biobank (the HCT series) as the cohort of Okamoto et al. (2022) [12], with largely overlapping patients. In that cohort, Okamoto et al. reported inter-patient heterogeneity using artificial intelligence (AI)-based 6-morphology typing. We noted that these diverse organoid morphologies appeared to separate broadly into two groups: one dominated by a compact glandular morphology and one dominated by round (cystic–mucinous) morphology. We undertook this study to identify the molecular axis that defines this separation. This study independently reanalyzes this biobank-matched cohort. It complements and extends the previous work by linking organoid morphology to TP53 status, to a transcript-level molecular axis (the MDM2 mRNA isoforms, P1/P2), and to prognosis and therapeutic stratification.

The working hypothesis that motivated this study was that MDM2 P1/P2 promoter choice is not only a phenotypic consequence of differences in genome-instability pathways. It may also contribute to the organ identity (lineage) of cancer cells, to the physical properties of the tumor tissue (inferred here from gene expression), and to the mode of interaction with the microenvironment. We tested this hypothesis against TP53 status and p53 activity. The study aims to provide a candidate readout of p53 activity that, after validation in biopsy tissue, could refine the molecular discrimination and therapeutic stratification of colorectal cancer. Gene-dependency maps of patient-derived tumor organoids are already being assembled on a large scale [13]. This study does not add the dependency map itself. Instead, it adds a candidate readout of p53 activity at a single locus (MDM2 P2 usage) that may refine TP53-based stratification of those dependencies. Full-length versions of the sections condensed in this article are provided in the Supplementary Report (Additional file 1).

The dual-promoter biology of MDM2 raises two further questions. First (Q1), mucinous carcinomas (a tumor type with abundant extracellular mucus) carry fewer p53-pathway alterations [14], and in the intrinsic consensus molecular subtype (iCMS) classification of CRC epithelial cells, TP53 mutations are associated with the chromosomally unstable iCMS2 class, whereas almost all MSI tumors fall into iCMS3 [15]. Whether these associations reflect TP53 status itself or the features that accompany it has not been separated, and it is unknown which genome-wide axis organoid morphology follows and whether this axis corresponds to chromosomal instability, TP53 status or mucinous histology in patient tumors. Second (Q2), TP53 genotype is an imperfect proxy for p53 function [16]. Wild-type tumors can lack p53 transcriptional activity, and whether expression of p53-target genes (genes directly activated by p53) predicts MDM2-inhibitor response beyond genotype is debated [17–19].

Transcriptome-based classifiers indicate that a proportion of TP53 wild-type tumors have lost p53 function [20,21], and a TP53-loss phenocopy signature (an expression state resembling that of TP53-mutant tumors in the absence of the mutation) is also present in many tumors without biallelic TP53 alterations [22]. Because P2 carries p53 response elements, its usage could reflect p53 activity directly at the locus of the drug target rather than TP53 genotype. The Results address these two questions in three parts. Section 3.1 asks which genome-wide axis morphology and lineage programs follow (Q1). Section 3.2 asks whether P2 usage reflects p53 activity rather than TP53 genotype and whether it relates to MDM2-inhibitor sensitivity (Q2). Section 3.3 characterizes the two expression programs that underlie these readouts.

## 2. Methods

This section summarizes each method; the full-length methodological descriptions are provided in the Supplementary Report (Additional file 1), Section R-2, and additional details of the methods of Sections 3.1 and 3.2 (Sections 3.1.4, 3.1.5 and 3.2) are given there and in the README of Supplementary Methods (Additional file 2), which also indexes all deposited analysis code. The methods are ordered to follow the Results: shared procedures first, then those specific to Section 3.1 (morphology and TP53 status), Section 3.2 (MDM2 P2 usage and p53 activity) and Section 3.3 (signature characterization); the transcript-level quantification of promoter usage in TCGA (P2_index; Section 3.2.2) and the colibactin analysis (Section 3.3.3) are described after the external cohorts, under their own headings.

Three measures of MDM2 promoter usage are used throughout, and the qualifying term (array, P2_index or cell-line) is always stated in the text and figure legends. The array P2 index (discovery organoids and the public HTA2.0 arrays of GSE128213) is the mean of the exon-2 probe sets minus the mean of the exon-1 probe sets, each taken relative to the MDM2 transcript cluster (a transcript cluster is the gene-level group of probe sets on the HTA2.0 array; RMA, a standard normalization of array intensities; probe sets defined under “Splicing Index analysis of MDM2 mRNA isoforms with TAC”, below), a signed difference on the log2 RMA scale. P2_index (TCGA-COAD/READ tumors, from RNA-seq transcript TPM, transcripts per million, an abundance unit normalized for transcript length and sequencing depth) is ΣTPM(P2)/ΣTPM(P1+P2), a ratio between 0 and 1. The cell-line P2 index (DepMap transcript TPM; DepMap is a public resource of cell-line expression, mutation and gene-dependency data) uses the same transcript-ratio formula as P2_index. The array measure is therefore on a different scale from the two transcript-level ratios, and MDM2 itself was excluded from every signature correlated with any of the three.

### Patient samples and generation of organoids

Colorectal cancer tissues (primary tumors and liver, lung, lymph node and ovarian metastases) were obtained from 21 patients, and organoids were generated from 63 samples [23]. Two arrays registered under the patient number HCT84 (HCT83-4LM, one of the 63 samples, and one array not used for discovery) derive from patient HCT83, as shown by the originating laboratory’s sample records and by expression-based identity checks (Section R-2); they are assigned to HCT83 in all patient-level analyses except one. In the genome-wide survey (Section 3.1.4), HCT83-4LM was counted as a separate patient (22 patients) when the direction of each cluster was checked in patient means; with 21 patients, the 124 morphology-only clusters were unchanged, the MDM2-axis-only clusters changed from 930 to 919, and the same chromosome arms were enriched (Genome-wide survey of morphology and TCGA histology, below). Specimen names follow the titles in the public database (GSE128213) except HCT27-1T (titled HCT27-1T-2, another scan of the same specimen); their correspondence with array file names and GSM accessions is given in the Availability of data and materials section. The cohort derives from the same biobank (the HCT series) as that of Okamoto et al. (2022) [12], with largely overlapping patients; the present study is an independent re-analysis, and the biobank, its patient-matched primary–metastasis design, transcriptome profiling and mutational profile are described by Okamoto et al. (2021) [24]. Eight of the 63 specimens were obtained after chemotherapy, and the passage number at array profiling ranged from 1 to 22 (median 3; recorded for 43 specimens; Section R-2); MSI/MMR status was not determined in this study (Section 4.6). Sample names give the patient number and the sampling site and occasion (e.g., HCT25-1T; LM, liver; LuM, lung; LN, lymph node; Ov, ovarian metastasis). The 15q11-q13 imprinted locus of the same cohort is analyzed in a companion paper (T. Tsukui, R. Yao, and K. Tsuda, unpublished observations) and is not interpreted here (this paper reports only the expression differences of its transcripts and the cluster score as one of the programs of Section 3.1.4); a second companion paper (T. Tsukui, R. Yao, and K. Tsuda, unpublished observations) examines p53 activity in TP53 wild-type tumors and refers to results reported here.

### Whole-transcriptome analysis by microarray

The 63 samples were profiled on the Affymetrix Human Transcriptome Array 2.0 (HTA2.0); for each sample, 50 organoids were pooled for RNA extraction and hybridized to one array, following the biobank-originating laboratory’s protocol [24]. Because HTA2.0 carries exon-covering (PSR) and junction-spanning (JUC) probes, exon usage can be quantified as with exon arrays [25]. Two preprocessing streams were used: sst-RMA (RMA with signal space transformation) values from Transcriptome Analysis Console (TAC 4.02, the array manufacturer’s analysis software) for the 33 two-group comparisons, the definition of differentially expressed genes (DEGs), the MDM2 Splicing Index and all IPA analyses; and RMA-normalized per-specimen values (R package oligo [26]; Code2, Additional file 2) for the EMT score, PCA/UMAP, the within-patient paired analysis and the clustering of the 39-gene panel. For replication (Sections 3.1.2 and 3.2.1), 19 arrays of the same biobank not used for discovery were normalized together with the 63 discovery arrays by RMA, and only the 14 identical to files public in GSE128213 (2 normal and 12 tumor) were analyzed, with each gene z-scored against the 63 discovery arrays (R71, R85 and R89; Section R-2); one unpublished tumor array (HCT55-1T) was added only in a sensitivity analysis.

### Image classification by deep learning (VGG16 transfer learning)

Organoid images were acquired by the procedure used for this biobank [12] (organoids embedded in Matrigel, a basement-membrane matrix gel, in 48-well plates, imaged with a stereomicroscope equipped with a monochrome charge-coupled device (CCD) camera from 4 days after plating) and were classified by transfer learning based on VGG16 [27]. Of 194 manually labeled images of three morphologies (Type0 84, Type1 53, Type5 57), 129 were used for training and validation and 65 as the test set (image-level random split with a fixed seed); test accuracy was 98.5% (64/65) (Section 3.1.1; Table 1). Type1 (compact glandular) and Type5 (dispersed mucinous) are the two opposite morphologies of interest, and Type0 (the differentiated, single-lumen type of the same biobank’s 6-morphology typing) is a reference category. The 792 images of the 63 samples were classified with training-matched input normalization (Code1/Code1b, Additional file 2), and per-specimen class fractions were calculated, the non-Type1 fraction being the sum of the Type0 and Type5 fractions; no majority-vote labels were created (Additional file 3: Supplementary Table S2; details in the Supplementary Report, Section R-3).

**Table 1.**
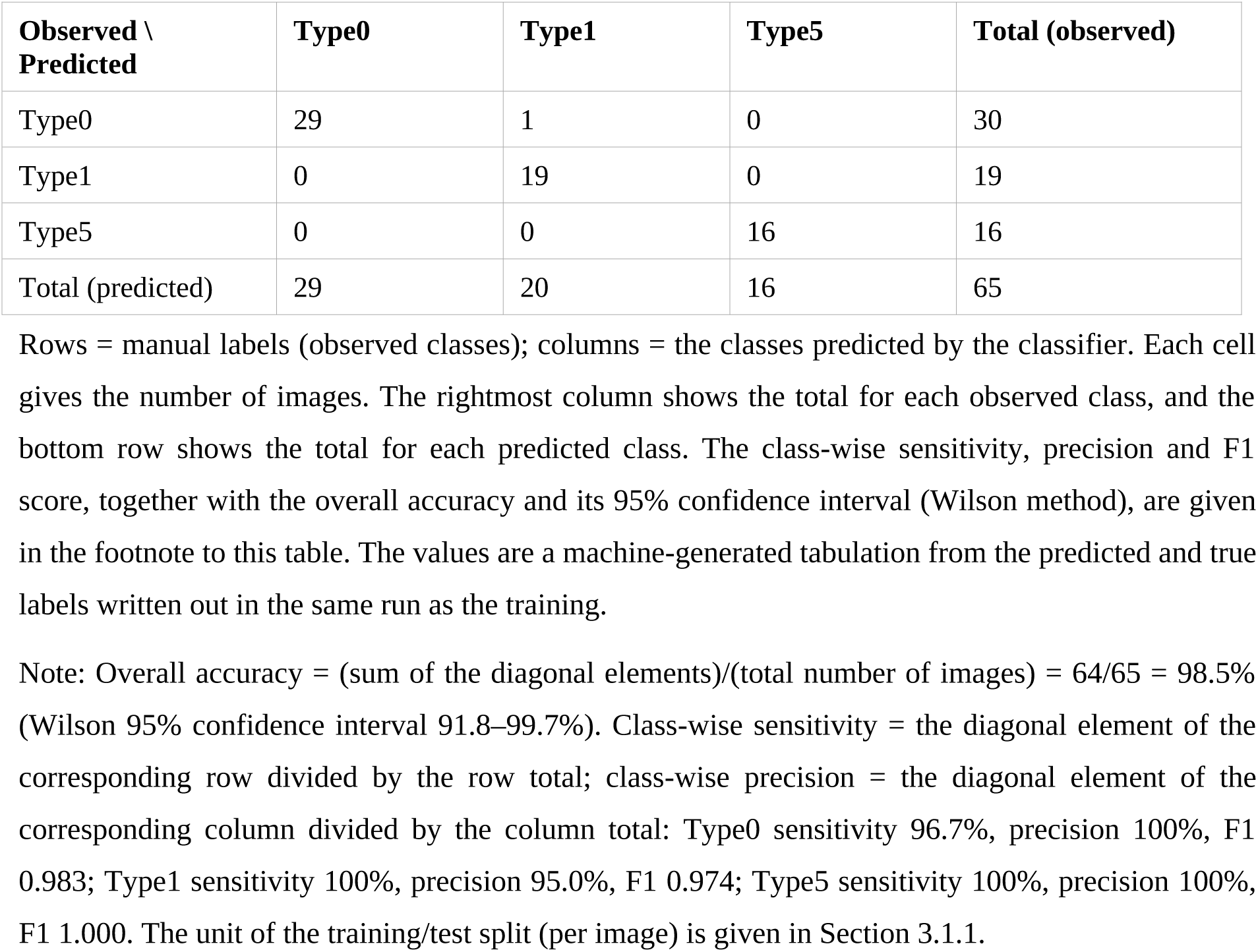
Confusion matrix of the deep-learning classifier (VGG16 transfer learning) on the test set.

| Observed \ Predicted | Type0 | Type1 | Type5 | Total (observed) |
| --- | --- | --- | --- | --- |
| Type0 | 29 | 1 | 0 | 30 |
| Type1 | 0 | 19 | 0 | 19 |
| Type5 | 0 | 0 | 16 | 16 |
| Total (predicted) | 29 | 20 | 16 | 65 |

Rows = manual labels (observed classes); columns = the classes predicted by the classifier. Each cell gives the number of images. The rightmost column shows the total for each observed class, and the bottom row shows the total for each predicted class. The class-wise sensitivity, precision and F1 score, together with the overall accuracy and its 95% confidence interval (Wilson method), are given in the footnote to this table. The values are a machine-generated tabulation from the predicted and true labels written out in the same run as the training.
Note: Overall accuracy = (sum of the diagonal elements)/(total number of images) = 64/65 = 98.5% (Wilson 95% confidence interval 91.8–99.7%). Class-wise sensitivity = the diagonal element of the corresponding row divided by the row total; class-wise precision = the diagonal element of the corresponding column divided by the column total: Type0 sensitivity 96.7%, precision 100%, F1 0.983; Type1 sensitivity 100%, precision 95.0%, F1 0.974; Type5 sensitivity 100%, precision 100%, F1 1.000. The unit of the training/test split (per image) is given in Section 3.1.1.

### Splicing Index analysis of MDM2 mRNA isoforms with TAC

The Splicing Index (SI) of TAC — (exon intensity / gene intensity in condition 1) / (exon intensity / gene intensity in condition 2) — compares MDM2 exon usage after removing gene-level expression; TAC reports SI as a signed linear value (a ratio below 1 is given as −1/ratio), so SI > 0 means relatively more RNA of the probe region on the tissue-1 side and SI < 0 on the tissue-2 side. MDM2 has 12 major exons [28]; P1-initiated transcripts start at exon 1 and skip exon 2, whereas P2-initiated transcripts use exon 2 instead of exon 1. Exon 1 is detected by three PSR probes (PSR12007995, PSR12007996, PSR12007998) and exon 2 by JUC12004243 and PSR12008003 (coordinates in the Supplementary Report, Section R-2). Each specimen’s dominant promoter was assigned from the MDM2 Splicing Index — P1-dominant when the exon-1 probes were relatively enriched on its side of the comparison and P2-dominant when the exon-2 probes were — and the call was checked by single-specimen comparisons against a common reference specimen of the opposite type (HCT64-1T for P1-dominant and HCT25-1T for P2-dominant specimens), taken from the 33 two-group comparisons (below) and from 30 additional single-specimen split comparisons (analyses 110–144; Additional file 4: Supplementary Table S3).

### TP53 mutation analysis

TP53 status was determined from targeted resequencing performed by the biobank-originating laboratory [24] (HaloPlex target capture, MiSeq sequencing, GATK HaplotypeCaller variant calling, ANNOVAR annotation), with three TP53 isoform transcripts as references and conversion to canonical numbering (e.g., R136H on NM_001126118 corresponds to R175H). Variants were classified by the ACMG/AMP criteria [29] (standard guidelines for rating variant pathogenicity) with reference to the public variant databases ClinVar [30] and dbSNP [31]; intronic and untranslated-region variants and variants also found in non-tumor tissue were not counted as somatic mutations. Per-specimen variant tables were available for 35 of the 63 samples; for the others, status was taken from the per-organoid mutation maps published for the same biobank (Fig. 1A in ref. [24] and Fig. S5 in ref. [12]), and two samples of patient HCT25 that appear in neither were assigned the patient’s mutation, present in all six of its other samples. Seven samples whose status could not be settled were treated as undetermined and excluded from analyses using TP53 status, and sensitivity analyses assigned them to either group (Additional file 5: Supplementary Table S11, which gives the source of each call).

**Figure 1.**
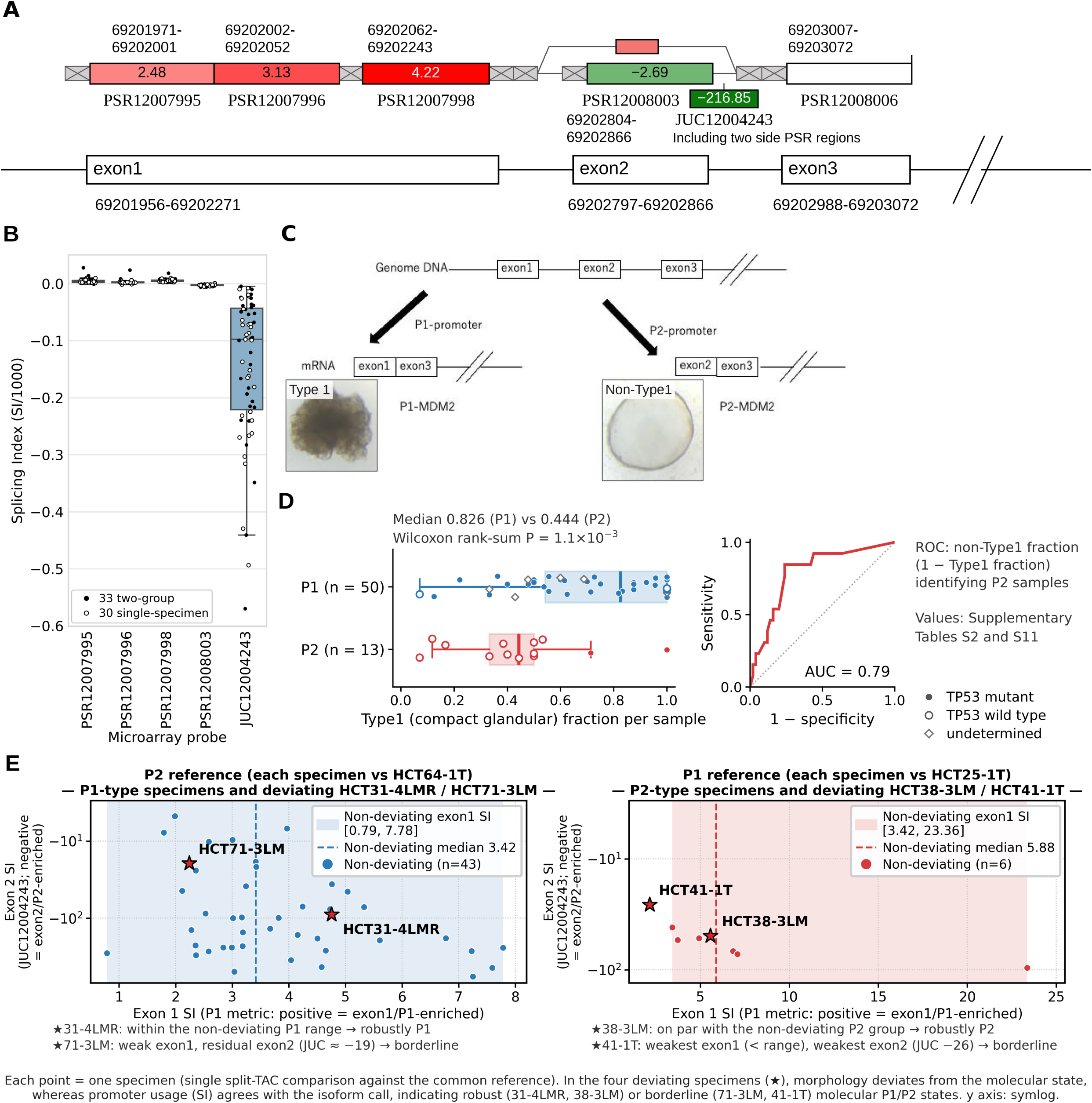
Deep-learning classification of organoid morphology and its correspondence with MDM2 mRNA isoforms. (A) The 5′ region of MDM2 (exons 1–3) and the HTA2.0 probes used for the Splicing Index (SI): exon 1, PSR12007995, PSR12007996 and PSR12007998; exon 2, JUC12004243 and PSR12008003. Probe boxes show SI values of a representative comparison (analysis 49 in Additional file 17: Supplementary Table S12); unnumbered boxes, probes not used for the SI. Coordinates, chr12, GRCh37/hg19 (GRCh38 in Section 2). (B) MDM2 SI across the 63 comparison sets of Supplementary Table S12: 33 two-group comparisons (analyses 49–109; filled dots) and 30 single-specimen split comparisons (analyses 110–144; open dots); box, IQR; whiskers, 1.5×IQR; y axis, SI/1,000. Exon 1 probes were positive on the tissue-1 side and exon 2 probes negative on the tissue-2 side, with four exceptions (PSR12007995 in analysis 108, PSR12008003 in analysis 115, PSR12007996 in analyses 122 and 129). (C) Schematic of P1- and P2-initiated MDM2 transcripts, with representative compact glandular (Type1) and cystic (non-Type1) organoids classified by the VGG16 model. (D) Morphology–isoform correspondence (Section 3.1.2): per-sample Type1 fraction of P1-dominant (n = 50) and P2-dominant (n = 13) samples (median 0.826 versus 0.444, Wilcoxon rank-sum P = 1.1×10⁻3; box, IQR; whiskers, 1.5×IQR), and ROC curve for identifying P2-dominant samples by the non-Type1 fraction (AUC 0.79). Marker fill, TP53 status (filled, mutant; open, wild type; diamonds, undetermined; Section 3.1.3). Values in Additional file 3: Supplementary Table S2 and Additional file 5: Supplementary Table S11. (E) Molecular type, by single-specimen split SI, of the 4 specimens deviating from the typical morphology of their group (Section 3.1.2); x axis, mean SI of the three exon 1 probes. Only specimens with a single-specimen comparison against the common reference (HCT64-1T or HCT25-1T) are plotted: 45 of 50 P1-dominant and 8 of 13 P2-dominant.

### Genome-wide survey of morphology and TCGA histology

For the 39,438 sufficiently expressed transcript clusters, rank partial correlations with the non-Type1 fraction and with the array P2 index (adjusted for scan period, the month of array scanning taken from the dates of the CEL files [the raw array intensity files], six categories, as a proxy for processing batch) were computed, and clusters with false discovery rate (FDR) < 0.05 and the same direction in patient means were retained (22 patients with HCT83-4LM counted separately; with 21 patients, the 124 morphology-only clusters [44 on 20q] were unchanged, the MDM2-axis-only clusters changed from 930 to 919, and the same arms were enriched). Exon usage was screened genome-wide in the same arrays (each PSR and JUC probe set expressed relative to its transcript cluster; P2-versus P1-dominant specimens compared on patient-by-group means, with the same direction required in a specimen-level model adjusted for scan period; R68), and in the survey the same relative values (MDM2 excluded) were related to both axes with the criteria above (R77; thresholds in Section R-2). Transcript clusters were assigned to chromosome arms by centromere position; arm-level enrichment was tested by one-sided Fisher’s exact tests (42 arms; Benjamini–Hochberg), arm-level mean expression served as a proxy for copy number, and the genes on the two morphological sides were tested for bias toward ten prespecified cytoskeleton and adhesion GO terms (one-sided Fisher’s exact tests). The morphology gene score in primary TCGA-COAD/READ tumors (Toil gene TPM, the uniformly re-quantified RNA-seq expression distributed by UCSC Xena; genes with TPM ≥ 1 in at least half of the primary tumors, 39 on the cystic–mucinous side and 64 on the compact glandular side) was the mean z-score of the genes of the 141 morphology-associated clusters higher on the cystic–mucinous side minus that of the genes higher on the compact glandular side; histology was taken from the histological_type field of the TCGA Pan-Cancer clinical table (TCGA-CDR) [32] — tumors recorded as colon or rectal mucinous adenocarcinoma (43; the pathological diagnosis recorded by TCGA, not re-assessed here) were counted as mucinous carcinoma, and three tumors recorded as discrepant were excluded (372 tumors) — and molecular subtypes from the TCGA molecular subtype table of UCSC Xena [33] (a public cancer-genomics data portal). Histological terms follow the WHO classification, sixth edition (2026) [34]. Whether morphology and expression programs follow TP53 status or the P2 measure was tested with rank linear models (partial F test for the added term, which asks whether that term improves the fit) in the discovery organoids (56 specimens with determined TP53 status), the GSE294511 organoids with published TP53 status (88) and TCGA-COAD/READ primary tumors with MC3 TP53 calls (MC3 being the TCGA pan-cancer consensus somatic-mutation call set; 358), with 20q and aneuploidy (arm-level expression proxies: within each organoid cohort, an arm score was the mean z-score of the expressed transcript clusters [median log2 RMA ≥ 5] on the arm minus the sample’s genome-wide mean, over the 39 arms with at least 40 clusters and with the clusters of the p53-score genes and of 30 known p53 targets excluded; 20q was the 20q arm score and the aneuploidy index the standard deviation of a sample’s 39 arm scores; in TCGA, 20q was the mean z-score of the expressed 20q genes) or 20q and MSI (the HM-indel subtype of the molecular subtype table; 335 tumors with a molecular subtype) as covariates; a sensitivity analysis in TCGA (TP53 calls of Additional file 6: Supplementary Table S47; 327 tumors) added BRAF mutation (any non-silent BRAF mutation in the MC3 calls, not restricted to V600E) and the CpG island methylator phenotype (CIMP; widespread hypermethylation of CpG islands; CIMP-high when, on the 450K DNA-methylation arrays, the median β value of the probes in the CpG island of at least three of the five markers of Weisenberger et al. [35] — CACNA1G, IGF2, NEUROG1, RUNX3 and SOCS1 — exceeded 0.30; called by us, because the subtype table gives no CIMP call for CRC; Section R-2), and non-MSI, BRAF wild-type, CIMP-low tumors were compared by Mann–Whitney tests and with adjustment for 20q. Program scores were the mean within-cohort z-scores of previously defined gene sets (the P1 and P2 signature components, among them the metaplasia genes of the P2 signature — nine gastric-type genes [CTSE, REN, TFF1, TFF3, MUC5B, MUC5AC, MUC6, MUC17 and ANXA10] and the Paneth-cell genes DEFA5 and DEFA6; the p53 activity score; a nine-gene gastric marker set fixed before the GSE294511 analysis [CTSE, TFF1, TFF2, MUC5AC, MUC6, ANXA10, REN, MUC17 and VSIG1]; and the SNORD116 cluster; gene lists in Section R-2; in TCGA, only genes with TPM ≥ 1 in at least half of the tumors were used, leaving seven of the 11 metaplasia genes and four of the nine gastric genes; gastric metaplasia in this study denotes the expression of these gastric-type genes and was not assessed histologically or by immunostaining), and the genome-wide comparison of TP53 status and promoter usage in the discovery organoids regressed each of 19,676 expressed genes on TP53 status, on promoter usage and on both, with cluster-robust standard errors by patient (which allow for correlation among specimens of the same patient) and Benjamini– Hochberg correction. These analyses were added after the per-specimen TP53 calls had been finalized (post hoc), and the program analysis, the CIMP sensitivity analysis and the genome-wide comparison were reproduced by independent re-implementations (Additional file 7: Supplementary Table S48, sheet H).

### Independent organoid biobank (GSE294511)

For replication in independent patients, we used the public HTA2.0 arrays of a biobank of organoids from colorectal liver metastases established at another institution (GSE294511; 119 organoids from 50 patients and 50 matched tumor tissues) [36], together with the per-organoid TP53 status, hematoxylin and eosin (H&E)-based morphology (cystic, mixed or solid) and drug sensitivity scores (DSS; higher = more sensitive; idasanutlin and the 23 other drugs common to both drug libraries of that study) published with it. CEL files were processed as for the discovery cohort (oligo RMA; organoids and tissues normalized separately), and the array P2 index (Section 3.1.2), the p53 activity score and the morphology gene score (the mean z-score of the 44 clusters higher on the non-Type1 side minus that of the 97 higher on the Type1 side among the 141 morphology-associated clusters; also recomputed from the 124 morphology-only clusters) were computed with definitions fixed in the discovery cohort; total MDM2 was the MDM2 transcript cluster (TC12000606.hg.1). Because the organoids come from fewer patients, the linear models of drug sensitivity were refitted with standard errors clustered by patient and the partial correlations repeated on patient means (sensitivity analyses added after the prespecified analyses). The analysis plan (primary test: one-sided Spearman correlation between the morphology gene score and the array P2 index, with patient-mean and patient random-intercept analyses and a within-patient comparison in patients whose H&E morphology differed between organoids) was fixed before the data were examined, and the key values were recomputed independently from the raw files (Additional file 7: Supplementary Table S48, sheet F).

### p53 activity score

Activity of wild-type p53 was scored as the mean z-score of 12 direct p53 targets (CDKN1A, BAX, BBC3, TP53I3, SESN1, RRM2B, GDF15, FDXR, TIGAR, PHLDA3, SPATA18 and EDA2R) [37] (p53 activity score); MDM2 itself and three p53 targets that ranked high in the genome-wide exon-usage screen (FAS, ZMAT3 and TNFRSF10C) were not included, to limit circularity with the P2 measures, and five of the 12 scored genes (CDKN1A, BAX, BBC3, GDF15 and SPATA18) are shared with the P2 signature (Section 3.2.1). Genes were z-scored across the 63 discovery specimens for organoids and within each cohort for TCGA and cell lines (10 of the 12 genes were available in the GDSC cell-line drug-sensitivity data); the per-specimen scores of the discovery organoids are listed in Additional file 7: Supplementary Table S48, sheet M, and their computation is deposited with a gate on the published group statistics and an independent re-implementation (R164; Additional file 2). TP53 wild-type samples without p53 activity were defined as those with a p53 activity score at or below the 90th percentile of mutant samples. Whether promoter usage carried information beyond the amount of MDM2 transcript was tested by adding the P2 measure to total MDM2 expression (the HTA2.0 MDM2 transcript cluster TC12000606.hg.1, or the sum of all MDM2 transcripts) in linear models of the ranked p53 activity score (partial F test), and by partial Spearman correlation in cell lines.

### RNA-seq of nutlin-treated cells

From the bigWig files (indexed genome-wide coverage tracks) of GSE86221 (a SubSeries of GSE86222; HCT116 TP53 wild type and null, MCF7 and SJSA; DMSO or nutlin 10 µM for 12 h) [38], the region around MDM2 (hg19) was extracted, and the mean coverage of the P1-only exon 1, the P2-only first exon, the shared ATG exon and the last exon was computed (exon coordinates in Section R-2; boundaries checked against the Ensembl GRCh37 annotation). Because the bigWig files are the sum of two replicates, the P2/P1 ratio was the primary measure, and induction of MDM2 and CDKN1A was confirmed with the gene-level count tables (R82; fold changes in counts per million written out by R169).

### Transcription initiation (ENCODE PRO-cap)

Transcription initiation at the two promoters was examined in ENCODE PRO-cap signal tracks (PRO-cap maps the 5′ ends of nascent capped RNA, that is, sites of transcription initiation; GRCh38) of six normal colon and stomach tissues and of Caco-2 cells, obtained from the ENCODE portal [39,40] (the Encyclopedia of DNA Elements, a public functional-genomics resource): the plus-strand signal was summed over windows around the P1 (chr12:68,808,100–68,808,300) and P2 (68,808,974–68,809,074) start sites, files of the same experiment were added (a reprocessed copy of the same replicate was counted once), and the P2 share was P2/(P1+P2) (Additional file 7: Supplementary Table S48, sheet K). The TP53 genotype of Caco-2 (p.Glu204Ter, homozygous) was taken from Cellosaurus (CVCL_0025) [41] and ClinVar [30].

### p53 activity, protein levels and junction reads in TCGA tumors

In TCGA-COAD/READ, the p53 activity score was computed from TCGAbiolinks expression [42] and, as a sensitivity analysis, from UCSC Xena Toil gene TPM [43,44]. TP53 status was taken from the GDC masked somatic mutation calls (a tumor with any TP53 variant other than silent, intronic, untranslated-region or RNA-gene classes was counted as mutant, and tumors absent from the calls were excluded rather than counted as wild type), and in the main analyses wild-type status was also confirmed in the MC3 mutation calls [45]; hypermutation was defined as ≥ 400 non-synonymous mutations. Protein levels of BAX, p21 and p53 were taken from the TCGA Pan-Cancer Atlas reverse-phase protein array (RPPA; antibody-based protein quantification) data (UCSC Xena [43]; R120). MDM2 junction reads were counted from recount3 [46] (uniformly reprocessed public RNA-seq, which provides exon–exon junction counts) for the TCGA-COAD/READ tumors: reads joining exon 1 (P1) or exon 2 (the P2 first exon) to exon 3, with the P2 share = P2 / (P1 + P2) (R117d; Additional file 7: Supplementary Table S48, sheet D). P2_index of adjacent normal colorectal mucosa and of patient-matched normal–tumor pairs was computed from the same Toil recompute (51 normal samples; 32 pairs; R160; Additional file 7: Supplementary Table S48, sheet I). The per-tumor scores of two published classifiers of p53 function — the TP53-loss phenocopy score of Fito-Lopez et al. [20] and the classifier of Li et al. [21] (obtained 14 October 2026) — were matched to the same tumors (R159 and R159b; Additional file 7: Supplementary Table S48, sheet J). Counts and P values in TCGA-COAD/READ depend on the expression source and on the TP53 and hypermutation call sets, so analyses of the same tumors with other choices (including those of the companion study on p53 activity in TP53 wild-type tumors; T. Tsukui, R. Yao, and K. Tsuda, unpublished observations) give counts and P values that differ from those reported here (for example, for the correlation in Section 3.2.3; the range over 12 combinations is given in Additional file 7: Supplementary Table S48, sheet N).

### Cell-line P2 index and drug response

The cell-line P2 index was computed from DepMap 24Q2 transcript TPM [47,48] with the same transcription-start-site assignment as for TCGA, restricted to lines with sufficient MDM2 expression (TPM ≥ 5; R74). TP53 status was taken from GDSC (COSMIC) [49] for analyses with GDSC (691 lines, 208 wild type) and from the DepMap mutation table for the CRISPR and PRISM analyses (364 wild-type lines; the two definitions agreed for 686 of 691 lines). Drug response used nutlin-3a ln IC50 from GDSC1 and GDSC2 [49,50] and the PRISM secondary (19Q4) and primary (24Q2) screens [51] (pooled drug screens of barcoded cell lines); MDM2 dependency used DepMap CRISPR gene effects (Chronos scores, more negative meaning stronger dependency; DepMap 24Q4) [52–54]; rank partial correlations adjusted for the p53 activity score were also computed (R70 and R76). Nutlin-3 sensitivity (area under the dose–response curve) in CTRPv2 [55,56] (taken from the data files distributed with oncoPredict [57]) was analyzed in the same way (R93). MDM2 junction reads were also counted directly in the SG-NEx cell lines, sequenced with short reads and with Nanopore long reads [58], as for TCGA, and the six lines with 20 or more junction reads and a cell-line P2 index (DepMap) were compared (R137 and R142).

### MDM2 inhibitor sensitivity and dependency, drug-sensitivity prediction, immune estimates and TP53 functional classes

Nutlin-3 is the racemate and nutlin-3a its active enantiomer, with the same mechanism of action; drug names follow each data source. MDM2 exon 1 (P1) and exon 2 (P2) are both 5′-untranslated and the first in-frame AUG lies in the shared exon 3 in the human [6] and murine [59] genes, so the nutlin-binding site of full-length MDM2 does not depend on promoter choice; initiation at downstream AUGs can yield N-terminally truncated MDM2 that cannot bind p53 [59], which our RNA-level data could not assess, and the Splicing Index does not detect the cancer-associated MDM2 splice variants lacking internal coding exons (MDM2-A, -B, -C) [28]. GDSC1/GDSC2 dose–response data [49,50] were matched against 13 MDM2–p53 pathway drug names (four matched: nutlin-3a, Serdemetan, PRIMA-1MET and MIRA-1), and LN_IC50 values were compared between TP53 wild-type and mutant lines (DepMap 24Q4 [52,53]) and between MSI-High and MSS/MSI-L lines by Mann– Whitney tests with Hodges–Lehmann effect sizes; MDM2 dependency used DepMap CRISPR gene effects (1,178 lines). The nutlin-3 data of the organoid biobank of reference [13] (primary data deposited on Figshare with that study; 65 colorectal lines with mutation data) were re-analyzed with exact Wilcoxon tests, Cliff’s delta, covariate-adjusted linear models and GSVA (gene set variation analysis, which scores each sample for a gene set; R38). Drug sensitivity of the 624 TCGA specimens was predicted with oncoPredict [57] (805 GDSC2 lines × 198 drugs; ComBat [60]; Code15), and on measured data P1/P2 scores from GDSC2 cell-line expression were related to LN_IC50 and to DepMap CRISPR gene effects by partial Spearman correlation adjusted for cancer type and TP53 status, and additionally for RAS/BRAF status (R47–R47c; drug classes compared by Wilcoxon tests on the per-drug partial correlations, R47d; the P2–P1 separation index is defined in Section R-16). The immune microenvironment was estimated with quanTIseq [61] and MCP-counter [62] via immunedeconv [63] on P2-score tertiles, with MSI-adjusted sensitivity analyses, and immune checkpoint inhibitor (ICI) response was predicted with TIDEpy 1.3.8 [64,65] (Code13). TP53 functional classes used the TP53 Database (release 21) [66,67] with two gain-of-function (GOF) definitions: missense at six hotspot residues [68] or all missense with a non-functional transactivation class [69] (Code14).

### Two-group comparative expression analysis with TAC

Using TAC 4.02 (sst-RMA), 33 comparisons were made between samples with MDM2 P1 dominance (“tissue 1”) and P2 dominance (“tissue 2”) (TAC analyses 49–109; one sample per patient in each comparison; compositions in Additional file 4: Supplementary Table S3; in analyses 49 and 52 the sides retain the morphological grouping of the exploratory phase). DEGs were defined by reproducibility of direction rather than by P value: the same sign of the gene-level mean linear fold change in at least 70% of the comparisons and |mean linear fold change| of at least twofold. This gave 1,890 genes (1,013 higher in tissue 1 and 877 in tissue 2; Table 2; Additional file 8: Supplementary Table S4). The definition was robust to the thresholds and to sign randomization (empirical FDR 0.60%; Additional files 9–11: Supplementary Table S5, Supplementary Figures S1 and S2; Code5; Supplementary Results 7 in the Supplementary Information, Additional file 12). Because the comparisons reuse specimens (the 13 P2-dominant specimens derive from six patients), they are not independent replicates, and these criteria are descriptive rather than patient-level inference. The EMT score was the mean z-score of 11 EMT/mesenchymal genes (CDH2, VIM, FN1, SPP1, VCAN, TGFB2, TGM2, LOXL4, CDH6, ALDH1A3 and ONECUT2) minus that of two epithelial markers (CDH1, EPCAM); within-patient paired comparisons (17 patients) used limma duplicateCorrelation [70], and PCA and UMAP [71] used the 3,000 most variable genes (Codes 3 and 6).

**Table 2.** Key genes by functional category.

| Functional category | Key genes in tissue 1 (in rank order) | Key genes in tissue 2 (in rank order) |
| --- | --- | --- |
| Most differentially expressed genes | OLFM4 (+1252; 73%), SLC26A3 (+782; 88%), NPSR1 (+516; 97%), CALB1 (+338; 69.7%), ALDH1A1 (+272; 94%), SLC9A3 (+199; 94%) | CTSE (−1901; 100%), REN (−1281; 97%), UCA1 (−547; 97%), MUC17 (−478; 85%), TFF1 (−408; 79%), DEFA5 (−381; 85%), HULC (−264; 94%), MMP7 (−111; 100%) |
| Transport and absorption | SLC9A3, SLC26A3, SLC5A8, SLC7A5, SLC6A4, SLC6A20, ABCB1, ABCC4, CFTR, ANPEP, | SLC6A14, SLC2A3, SLC2A1, SLC28A3, SLC12A2, SLC4A4, SLC44A4, SLC40A1, SLC34A2 |
|  | DPP4 |  |
| Mucin and secretion | MUC4, MUC12 (colonic-type mucin), FCGBP, PIGR | MUC5B, MUC5AC, MUC17, MUC6 (gastric-type/ectopic mucin), MSLN, PLIN2 |
| Metabolism and detoxification | ALDH1A1, AKR1B10, GSTM4, GSTM1, UGT2A3, CYP4X1, CYP2W1, ALDH1L2, ALDH5A1, PYGB | CYP3A5, CYP2B6, HMGCS2 (ketone bodies), ALDH3A1, BAAT, UGT1A1 cluster, NQO1 |
| Cell cycle and proliferation | TOP2A, FOXM1, MCM4/5/6/8/10, MYBL2, AURKA/B, PLK1, CDK1/2, CDCA7/8, BUB1, TTK, STMN1 | CDKN1A, CCND2, CCNG1/2 (arrest type), ZMAT3 (p53 response), MDM2 (P2), RRM2B (p53 target) |
| Stem cells and epigenetics | LGR5, OLFM4, HELLS, DNMT1, SUZ12, BMI1, ATAD2/5 | SNORD116-1/2/3/5/7/8/9/14/15/16/17/18/19/20/24 (representative members; all members, including SNORD116-17/-19 with the largest effect sizes and SNORD116-12 with the highest consistency rate, are given in Additional file 36: Supplementary Table S16), SNORD115 cluster, HULC, UCA1, H19, UBD (FAT10) |
| Inflammation, immunity, and ECM | IFI30, IFI6, IRF8, BST2, TAP2, LYZ (limited). Note: the tissue-1 items in this row are comparison-specific findings based on comprehensive extraction and are not discussed in the main text of Section 3.3.1. The extent of immune cell infiltration itself was evaluated in Section 3.3.3 using TCGA specimens, and is higher in the P2-high group. | IL33, IL1B, CXCL14, CCL20, CXCL8, UBD (FAT10; the same molecule that is also listed in the epigenetics row of this table), TLR3, TLR4, MYD88, DUOX2, NOX1, MMP7, VCAN, MSLN |
| Apoptosis and stress | PRKDC, BRCA1, MSH2, NBN, FANCD2, ATM, TOPBP1 (repair systems predominant) | FAS, TNFRSF10B (DR5), TNFRSF10C, TNFSF10, BNIP3, BNIP3L, BAX, APAF1, CASP-related genes (executioners of p53-induced apoptosis) |

### Ingenuity Pathway Analysis (IPA) and EnrichR GO analysis

IPA (QIAGEN) [72] was run on each of the 33 comparisons (Canonical Pathway, Upstream, Disease and Bio Functions, Tox Function, Regulator Effects, Networks and Graphical Summary) and aggregated across comparisons. Upstream findings were adopted only with sign consistency of the activation z-score of at least 75% and |median z| ≥ 2; the other modules were used descriptively, and Regulator Effects integrated only the first 8 comparisons because of computational load. The top 400 DEGs on each side were tested for GO enrichment with EnrichR [73] (GO BP/CC/MF 2026; BH-adjusted P < 0.05). Settings and the sensitivity analysis of the adoption criteria (Additional file 13: Supplementary Table S10) are described in the Supplementary Note (Additional file 14), Section 1.

### Biomarker Detection

Using the Biomarker Detection function of IPA, 39 candidate biomarker genes discriminating the two tissue types were extracted, and hierarchical clustering of the 63 samples with these genes was carried out with an R script written by the authors (Manhattan distance, average linkage; the framework follows Eisen et al. [74], without row or column scaling).

### Signature validation and survival analysis in external independent cohorts

The organoid-derived P1 (21 genes) and P2 (29 genes) signatures were tested in two public cohorts: TCGA-COAD/READ primary tumors (624 specimens; STAR-Counts TPM via TCGAbiolinks [42]), classified with CMScaller [75] (563 classifiable), and GSE39582 [76] (GEOquery [77]), analyzed for the 519 tumors with MMR annotation (75 mismatch repair-deficient [dMMR], 444 mismatch repair-proficient [pMMR]) after removal of the 19 non-tumoral samples (the three P1/P2 genes absent from the platform did not change the conclusions in a sensitivity analysis recovering BBC3; R45 and R49). The P1 signature (CIN type) comprises intestinal epithelial identity and stemness (e.g., OLFM4, LGR5), the master transcription factor HNF4A, and the MYC/E2F/FOXM1-driven cell-cycle program; the P2 signature (resembling MSI-like/serrated tumors at the signature level only) comprises gastric and Paneth metaplasia genes (e.g., CTSE, REN, TFF1, DEFA5), wild-type p53 transcriptional targets (e.g., CDKN1A, DDB2, BAX, BBC3) and MDM2 itself, six immune checkpoint molecules (IDO1, PDCD1, LAG3, CTLA4, TIGIT, HAVCR2), and cancer-associated lncRNAs (full gene lists and selection rationale in the Supplementary Report, Section R-2; gene sets in Code8). Because the six immune checkpoint genes are expressed mainly by infiltrating lymphocytes, the CMS and MMR associations in bulk tumors are reported primarily for the immune-excluded P2 score (23 genes), with the full 29-gene score alongside. Scores were computed with GSVA [78] (single-sample gene set enrichment analysis [ssGSEA] as an alternative); CMS associations were tested by Kruskal–Wallis and dMMR/pMMR differences by Wilcoxon rank-sum tests, and the overlap between the DEGs and CMS class-specific marker genes by one-sided Fisher’s exact tests against two independent reference sets (Additional file 15: Supplementary Table S28). Survival analysis used the 591 TCGA patients with overall survival (OS) data (median-split Kaplan–Meier with log-rank tests; univariate and multivariate Cox [79] adjusted for age, stage and MSI; Code8). Tumor purity in the signature, survival and P2_index analyses was derived from the ESTIMATE score [80] (ESTIMATE infers stromal and immune content from bulk expression) with the published conversion (purity = cos(0.6049872018 + 0.0001467884 × ESTIMATE score); Code17), and the purity-adjusted analyses of Section 3.2.3 used ABSOLUTE purity (inferred from copy-number data). MSI status in TCGA-COAD/READ was taken from the HM-indel (MSI) subtype of the TCGA molecular subtype table in Section 3.1.4, from the subtype table returned by TCGAbiolinks in the survival analysis, from the GDC clinical field paper_MSI_status in the immune analyses and from the covariate table of Code9 (generated by Code16) in Section 3.2.2, and was available only for subsets of tumors (Sections R-2 and R-14); in GSE39582 the published MMR annotation and in cell lines the GDSC annotation (MSI-H and MSI versus MSS and MSI-L) were used.

### Quantification of MDM2 promoter usage in TCGA tumors (P2_index)

Promoter usage was quantified from transcript-level TCGA-COAD/READ expression (UCSC Xena, Toil RNA-seq recompute [43,44], GENCODE v23): MDM2 transcripts were assigned to P1 (8 transcripts) or P2 (23 transcripts) by transcription start site (coordinates in Section R-2), P2 lying within intron 1 [6,59] and containing SNP309 [7], and P2_index = ΣTPM(P2)/ΣTPM(P1+P2) was calculated per tumor (Code9; input matrix by Code16); a multivariable linear model with TP53 status, CMS, tumor purity and MDM2 copy number (GDC gene-level ASCAT3 [81]; Code19; total copy number ≥ 5 = high-level amplification) estimated independent contributions.

### Colibactin mutational signatures (exploratory)

An exploratory association of the P1/P2 axis with the colibactin mutational signatures SBS88 and ID18 (mutation patterns attributed to the genotoxin colibactin of pks-positive Escherichia coli) was examined in the Nunes cohort [82], with replication in TCGA-COAD/READ using sigminer [83] (R42–R44; Section R-2).

### Statistical analysis

All tests were two-sided unless stated otherwise (the Fisher’s exact tests of DEG–CMS marker overlap, chromosome-arm enrichment and GO-term bias, the patient-level permutation test and the primary test in the independent organoid biobank were one-sided), and multiple testing was controlled by the Benjamini–Hochberg procedure within each family of tests. Group comparisons used the Wilcoxon rank-sum (Mann–Whitney) test or, for more than two groups, the Kruskal–Wallis test; correlations used Spearman’s ρ, with rank-based partial correlation where covariates were adjusted for; odds ratios of 2 × 2 tables were tested by Fisher’s exact test (the exact-versus-normal-approximation rules for rank tests are given in the Supplementary Report, Section R-2). The specimen was the primary unit of analysis, because specimens from the same patient could differ in morphology and MDM2 isoform (HCT38, HCT41 and HCT67) and even in TP53 status (HCT67); because several specimens derive from the same patient, the morphology–isoform comparison was also evaluated at the patient level as a sensitivity analysis with linear and binomial mixed models with a patient random intercept (R package lme4 [84]) and with a permutation test of isoform labels between patients; in the independent organoid biobank, linear models with a patient random intercept were fitted by maximum likelihood in Python, and standard errors clustered by patient (CR1) were used where stated. Analyses used R 4.6.0 and Python 3.6–3.13 (the image classifier: Python 3.6.2 with Keras 2.1.2 and TensorFlow 1.4.1); package versions are recorded in the sessionInfo() files and in the README of Supplementary Methods (Additional file 2), whose script identifiers map each analysis to the deposited code.

### Use of large language models

The manuscript was developed from an initial draft written by T.T. A large language model (Anthropic Claude) was used during the preparation of this study for the following purposes: (i) drafting and editing the manuscript text, including the Supplementary Report, the Supplementary Information and the Supplementary Note, and translating it into English, on the basis of that initial draft, through an iterative exchange of ideas with the author, whose own revisions are incorporated throughout the text; (ii) writing, reviewing and debugging analysis scripts in R and Python — excluding the image-classification, array-normalization (RMA), enrichment-analysis (EnrichR) and clustering code and the two deposited Jupyter notebooks (application of the image classifier and annotation of the array data), which the authors wrote themselves — and (iii) cross-checking numerical consistency between the manuscript, the tables and the primary analysis outputs. The model was not used to generate, impute or modify any experimental or derived data, and all reported values were computed by the deposited analysis code and verified by the authors against the primary outputs. The authors reviewed all model-assisted output and take full responsibility for the content of this manuscript.

## 3. Results

### 3.1 Morphology and lineage programs track TP53 status

The Background asks which genome-wide axis the morphology and lineage programs of organoids (three-dimensional cultures of patients’ tumor cells) follow (Q1). Section 3.1 shows, in the discovery organoids and then in an independent biobank and patient tumors, that morphology and the metaplastic programs follow TP53 status rather than MDM2 promoter usage itself.

#### 3.1.1 Deep-learning classification of organoid morphology

We first define the terms used from here on. “Type1 (compact glandular type),” “Type5 (dispersed mucinous type)” and the reference class “Type0” are the morphological classes that the classifier assigns to single images; the classes other than Type1 together form the cystic–mucinous “round morphology,” and the per-specimen Type fractions (including the non-Type1 fraction) are the proportions of a specimen’s images assigned to each class, image-based quantities distinct from the expression signatures. “tissue 1” and “tissue 2” denote the group-level contrast between MDM2 P1-dominant and P2-dominant specimens (MDM2 mRNA derived relatively more from the constitutive or the p53-inducible promoter; Section 2), which largely coincide with TP53-mutant and wild-type specimens (Section 3.1.3); the labels “autonomous-proliferation type” and “environment-adaptive type” summarize their expression programs without implying that promoter usage determines them, and “P1/P2 signature (score)” names the expression signatures. The Type numbers are inherited from the 6-morphology typing of Okamoto et al. (2022) [12] (hence no Type2–4) and carry no meaning of magnitude.

The classifier was a VGG16 network (an image-recognition network; ImageNet-pretrained, all layers trainable) with a three-class softmax head and 224×224 inputs, trained by stochastic gradient descent with EarlyStopping and no augmentation; across the five training runs with available logs, test accuracy was 64/65 or higher (architecture and training conditions in the Supplementary Report, Additional file 1, Section R-3).

Table 1 gives the overall test accuracy, the sensitivity, precision and F1 score of each class, and the 3-class confusion matrix (true versus predicted classes). Only 1 of 65 images was misclassified (an observed Type0 predicted as Type1), and misclassification between the two tumor-side classes (Type1 and Type5) was 0.

The trained classifier was applied to the 792 organoid images of the 63 specimens, with input normalization matched to training, to give per-specimen Type0/Type1/Type5 fractions (Additional file 3: Supplementary Table S2). Of the 194 training/evaluation images, 49 (6.2%) were also among the 792; performance claims rest on the held-out test set, which this overlap does not affect.

This classifier has seven limitations, reported following the CLAIM [85] and TRIPOD+AI [86] guidelines (full statement in Section R-3; Section 4.6): a single institution and imaging condition without external validation; manual labels by the authors without inter-observer agreement; a small test set (Wilson 95% confidence interval of the 98.5% accuracy, 91.8–99.7%); an image-level (not patient-level) split that may optimistically bias accuracy; possible propagation of misclassification to group assignments; a preprocessing mismatch at the initial application, corrected by re-classifying all 792 images with training-matched normalization (which agreed better with the manual labels and preserved the P1/P2 separation; both conditions in Additional file 16: Supplementary Table S46); and a median of only 10 images per specimen (range 2–48). Time-lapse observation in the same biobank found stable morphology in 19 of 21 organoids over 3 days [12], making short-term fluctuation an unlikely confounder.

#### 3.1.2 MDM2 mRNA isoform usage corresponds to morphology at the specimen level

In the 63 specimens, we analyzed MDM2 with the Splicing Index (SI), which compares the relative use of each exon between two groups after removal of gene-level expression, across the 33 two-group comparisons and 30 additional single-specimen split comparisons (63 comparison sets). Exon 1 SI values were positive on the tissue-1 (P1-dominant) side and exon 2 values negative on the tissue-2 (P2-dominant) side, and the signs of the representative probes (PSR12007998 for exon 1 and JUC12004243 for exon 2) reversed between the two regions in every comparison set; a single non-representative probe had the opposite sign in each of only four sets (analyses 108, 115, 122 and 129) (Figure 1A–C; all SI values in Additional file 17: Supplementary Table S12; Section R-4): this is direct evidence that P1 and P2 promoter usage is shifted in opposite directions between tissue 1 and tissue 2. The isoform axis was found through exploration (TAC comparisons were first made between the two morphological groups, the Splicing Index emerged there, and later comparisons were made between isoform groups). Single-specimen split SI, which compares one specimen with a common reference of the opposite type, confirmed the dominant promoter for 53 specimens (45 of 50 P1-dominant and 8 of 13 P2-dominant), all with the expected signs of the representative probes (Additional files 3 and 4: Supplementary Tables S2 and S3).

Morphology and isoform corresponded clearly at the specimen level. The 50 P1-dominant specimens had higher Type1 fractions than the 13 P2-dominant specimens (median 0.826 versus 0.444; Wilcoxon P = 1.1×10⁻3), and the P2-dominant specimens had correspondingly higher non-Type1 fractions (area under the receiver operating characteristic curve [AUC] 0.79 for identifying P2 by the non-Type1 fraction; Type0 fraction P = 0.013; Figure 1D; Additional files 3 and 16: Supplementary Tables S2 and S46). The correspondence held in the three patients with only P2-dominant specimens (patient-level permutation P = 0.035, one-sided). In a sensitivity analysis for the non-independence of specimens from the same patient (the 13 P2-dominant specimens derive from six patients, three exclusively P2), the difference at the patient level was in the same direction (linear mixed model with a patient random intercept, P = 0.069; patient × isoform means, exact Wilcoxon P = 0.20; Section R-4). At this stage we could not determine whether morphology followed promoter usage itself or TP53 status, which largely coincided with it (Section 3.1.3); Sections 3.1.4 and 3.1.5 indicate that it follows TP53 status rather than promoter usage.

The four specimens that deviated most from their group — the two P1-dominant specimens with the lowest Type1 fractions (HCT71-3LM, HCT31-4LMR) and the two P2-dominant specimens with the highest (HCT41-1T, HCT38-3LM) — showed dissociation between morphology and isoform. Their patients, together with HCT67 (intra-patient P1→P2 switch; Section 4.2), are all among those in whom the independent typing of the same biobank [12] found organoids of both type A and type B within one patient, which supports real intra-patient morphological admixture rather than classifier error. HCT38-3LM had activated a cancer-cell-intrinsic epithelial–mesenchymal transition (EMT) program, a shift toward a motile, mesenchymal-like cell state (Additional files 18 and 19: Supplementary Tables S13 and S14). By single-specimen split SI, the molecular type of all four agreed with the isoform assignment, robustly for HCT31-4LMR and HCT38-3LM and borderline for HCT71-3LM and HCT41-1T (Figure 1E; Section R-4). These are exploratory findings based on a small number of cases.

The EMT score was higher in metastatic than in primary specimens (median +0.130 versus −0.590 in all 63 specimens; Wilcoxon P = 0.014; excluding the four morphology-deviating specimens, the medians are +0.05 and −0.56) but did not differ significantly between P2- and P1-dominant specimens (P = 0.079) or by TP53 status; we therefore treat it as an axis of progression along sampling site rather than a layer of the isoform contrast (Additional file 7: Supplementary Table S48, sheet L; sites in Additional file 20: Supplementary Table S1; Section 4.2.1). In a within-patient paired analysis of primary–metastasis pairs (17 patients; limma with duplicateCorrelation), no gene passed FDR < 0.05 and |log2FC| > 1, although all 17 EMT/mesenchymal genes examined in that paired analysis (a larger list that partly overlaps the 11 genes of the EMT score) were higher in metastases, none significantly so individually (Section R-4). The miR-200/ZEB1/2 axis showed a partial-EMT pattern: ZEB1 and ZEB2 leaned toward tissue 2 in 29 and 28 of the 33 comparisons, but only by a median of 1.1- and 1.3-fold, while miR-200c was also higher in tissue 2 (median 1.6-fold, 25 of 33; Additional file 21: Supplementary Table S15). In principal component analysis (PCA) and UMAP, P1- and P2-dominant specimens separated, and the patients with specimens of both types (HCT38, HCT41, HCT67) followed markedly different primary-to-metastasis trajectories (Additional files 22 and 23: Supplementary Figures S3 and S4).

To test the P1/P2 separation outside the discovery set, we used 10 organoid arrays from seven patients of the same biobank not used for discovery, normalized together with the 63 discovery arrays. A per-specimen array P2 index separated the discovery specimens without overlap and correlated with the P2 signature score (Spearman ρ = 0.61); in the 10 replication arrays it again correlated with the P2 signature (ρ = 0.82, P = 0.0038), and the three P2-dominant specimens had higher P2 scores than the seven P1-dominant ones (Mann–Whitney P = 0.017; Additional file 24: Supplementary Table S24; Section R-4). Because all these specimens come from patients already in the discovery cohort, this is replication in independent arrays and specimens, not in independent patients; replication in independent patients is reported in Section 3.1.5.

#### 3.1.3 TP53 status, MDM2 isoform, and morphology: a three-way correspondence

Targeted resequencing determined TP53 status for 56 of the 63 samples (Section 2). It identified seven pathogenic and two likely pathogenic mutations, consistent with TP53 mutations encountered clinically in colorectal cancer (CRC) and reported in the IARC TP53 database [66] (Additional file 5: Supplementary Table S11; Section R-8). The published mutation maps showed further mutations, truncating (protein-shortening) in two patients and non-truncating in five, without the amino-acid change; the benign codon 72 polymorphism P72R [31] was not treated as pathogenic. In patient HCT67, R175H was the only somatic coding mutation; it was detected in the primary tumor and the liver metastasis but not in the recurrent liver metastasis HCT67-4LMR.

The per-specimen three-way correspondence is given in Additional file 25: Supplementary Table S23 (a patient was called mutant when a pathogenic or likely pathogenic mutation was detected in at least one specimen and was assigned to the P1 or P2 group by the isoform of most of its specimens). TP53 mutations largely coincided with the P1 group (patient level, Fisher P = 0.0088; among specimens with determined status, 40 of 43 P1-dominant and 2 of 13 P2-dominant, P = 1.7×10⁻⁷). The mutations with a known amino-acid change all impair the ability of wild-type p53 to activate its targets (loss-of-function or dominant-negative [87]; some hotspot variants may also have gain-of-function activities [68]), so in most mutant cases the p53 that activates the MDM2 P2 promoter through its response elements [88] would be absent, consistent with MDM2 remaining at constitutive P1-dominant expression (the two TP53-mutant, P2-dominant specimens, which were p53-active, are examined in Section 3.2.1); the two wild-type P1-dominant patients suggest that P1 dominance may also arise by other routes (e.g., failure of P2 induction by EGR1). The P2-dominant patients (HCT27, HCT33, HCT64) had no pathogenic mutation (effectively wild-type TP53), consistent with tumorigenesis that does not pass through TP53 mutation, such as the serrated pathway, and with TP53 mutations not being enriched in CMS1 [2]. TP53 status was also associated with morphology: mutant specimens had a higher Type1 fraction than wild-type specimens (median 0.854 versus 0.422, P = 6.1×10⁻⁴; Figure 2E), to a degree similar to promoter usage, and three of the four morphology-deviating specimens followed TP53 status rather than promoter usage. These descriptive comparisons alone therefore cannot separate the morphology–isoform correspondence from TP53 status; Sections 3.1.4 and 3.1.5 do so by adjusted analyses and in an independent biobank.

**Figure 2.**
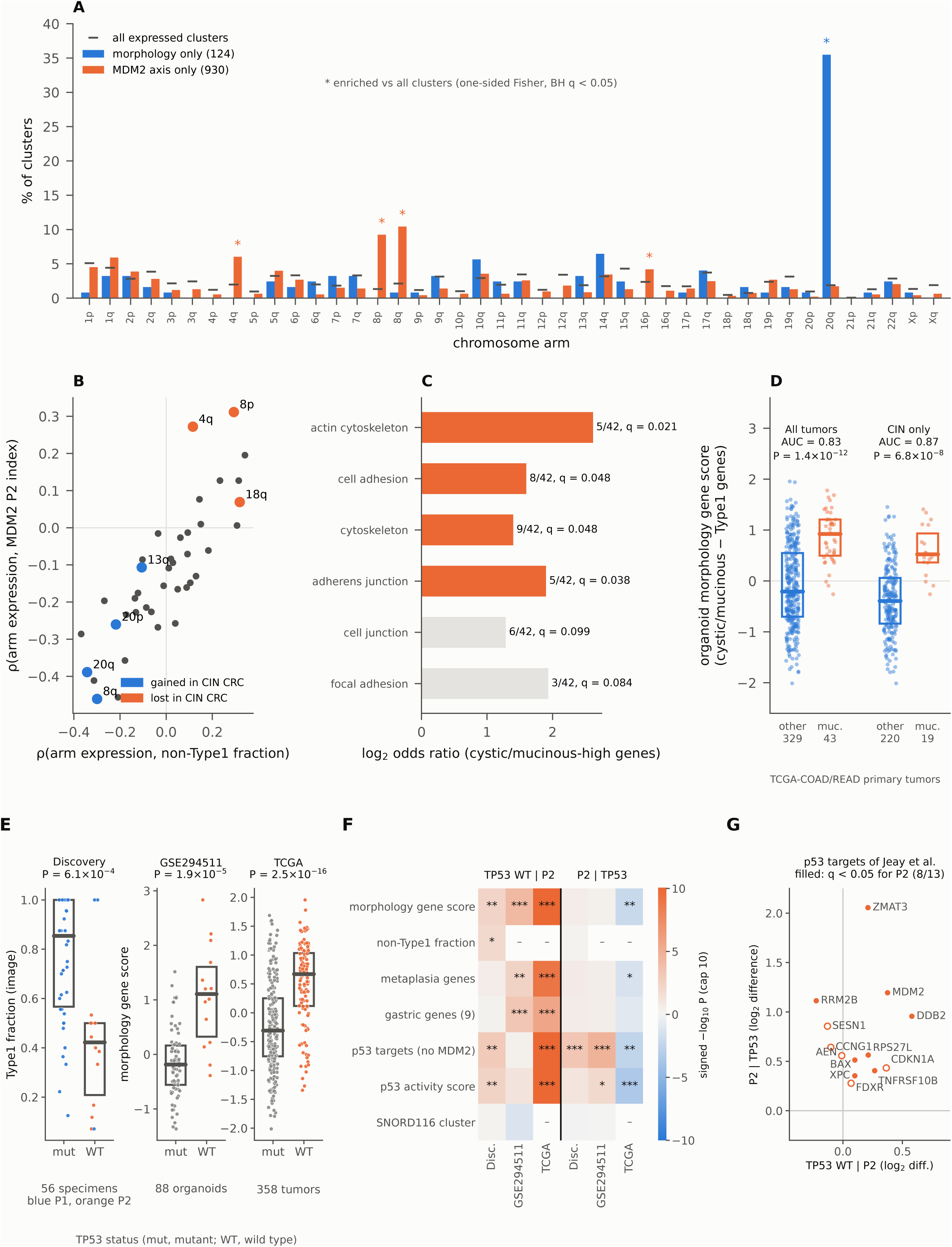
Morphology follows TP53 status, and the MDM2 axis follows p53 activity. (A) Proportion of transcript clusters on each chromosome arm associated with morphology only (124) or the MDM2 axis only (930). Ticks, all expressed clusters; *, BH q < 0.05, one-sided Fisher test. 44 of the 124 morphology-only clusters lie on 20q. (B) Correlation of arm-level mean expression with the two axes (non-Type1 fraction and array P2 index); dots, arms. (C) Cytoskeleton and adhesion GO terms enriched among genes higher on the cystic–mucinous side. (D) Morphology gene score and mucinous histology in primary TCGA-COAD/READ tumors: all, and chromosomally unstable (CIN) only. (E) Morphology by TP53 status in three cohorts: Type1 fraction of the 56 discovery specimens with determined status (colors, blue P1- and orange P2-dominant); morphology gene score of the 88 GSE294511 organoids with published TP53 status and of 358 primary TCGA-COAD/READ tumors with MC3 calls. Bars, medians; boxes, IQR; other cohorts: gray, mutant; orange, wild type; higher scores, cystic–mucinous side; Mann–Whitney P. (F) Programs following TP53 status or the P2 measure (array P2 index in organoids, P2_index in TCGA) in rank linear models without other covariates. Signed −log10 P: TP53 WT | P2 (wild-type status adjusted for P2; positive, higher in wild type) and P2 | TP53 (P2 adjusted for TP53 status; positive, higher with P2). *, P < 0.05; **, P < 0.01; ***, P < 0.001; colors capped at 10; –, not available. (G) The 13 p53-target genes of Jeay et al. [17] in the discovery organoids: coefficients for promoter usage adjusted for TP53 status (y axis) and for TP53 wild-type status adjusted for promoter usage (x axis) (log2 difference; patient-clustered standard errors). Filled, q < 0.05 for promoter usage (8 of 13). Values for E–G: Additional file 7, Supplementary Table S48, sheet H.

We mapped the fold changes of each comparison onto the comparison design. MDM2, CTSE, REN, and MMP7 were consistently higher on the tissue-2 side in all three P2 patients (HCT27, HCT33, HCT64; median fold change within each patient, MDM2 6.0 to 7.9, CTSE 21 to 2,353, REN 2.9 to 1,439 and MMP7 8.5 to 19.6), whereas EGR1 varied among them (1.5- and 1.6-fold higher on the tissue-2 side in HCT27 and HCT64 but 3.5-fold higher on the tissue-1 side in HCT33; Additional file 26: Supplementary Table S26; Additional file 27: Supplementary Figure S8; Additional file 28: Supplementary Table S18).

#### 3.1.4 A genome-wide test of which axis morphology follows

Organoid morphology corresponded to promoter usage (Section 3.1.2), and promoter usage largely coincided with TP53 status (Section 3.1.3), whereas morphology and the MDM2 axis themselves were only weakly correlated (non-Type1 fraction and array P2 index, ρ = 0.33). To ask which axis morphology follows, we related each of 39,438 sufficiently expressed transcript clusters (array units that roughly correspond to genes) to both axes by rank partial correlation (Section 2). Of the 124 clusters associated with morphology only, 44 lay on the long arm of chromosome 20 (20q) (background 1.9%, P = 6.5×10⁻⁴⁴), all higher on the compact glandular side; the 930 clusters associated with the MDM2 axis only were centered on p53 targets (FAS, DDB2, RPS27L, ZMAT3 and others, whose median fold changes between the P2- and P1-dominant sides of the 33 comparisons were 3.0, 10.5, 2.7 and 13.0, all higher on the P2 side; Additional file 8: Supplementary Table S4) and enriched on 8p, 8q and 4q, and weakly on 16p (Figure 2A). At the chromosome-arm level, the chromosomal-instability (CIN) expression pattern of colorectal cancer (inferred gains of 20q and 8q; losses of 8p, 18q and 4q; TP53 is mutated in most of these tumors) [89] matched the compact glandular, P1-dominant side, and the arm-level correlations with the two axes were of nearly the same direction and magnitude (Figure 2B; Section R-26). Thus, at the transcript-cluster level morphology corresponded preferentially to 20q and the MDM2 axis to p53 targets, whereas at the arm level both axes matched the same CIN pattern, which explains their overlap. Genes higher on the cystic– mucinous side were enriched for four of ten prespecified cytoskeleton and adhesion Gene Ontology (GO) terms (Figure 2C), genes on the opposite side were not, and no exon usage corresponded to morphology.

Here and below, the organoid morphology genes are the 141 transcript clusters whose expression followed morphology (the 124 associated with morphology only and the 17 associated with morphology and the MDM2 axis in the same direction), and the morphology gene score is the difference in expression between their two sides (Section 2; each cluster, with its correlations, chromosome arm and use in TCGA, is listed in Additional file 7: Supplementary Table S48, sheet O, and the gene symbols of each side are listed together at the end of that sheet, O3). They are a data-driven summary of the morphological axis rather than a predefined pathway: the 44 clusters higher on the cystic–mucinous side are almost all protein coding, none on 20q, and include genes of the actin cytoskeleton, cell adhesion and membrane trafficking (VCL, AHNAK, TRIOBP, FYN, CDC42BPB, INF2, ARHGAP5, PLEKHG3 and MICALL1; partial ρ with the non-Type1 fraction, for example, 0.58 for FYN and 0.52 for VCL) and the tetraspanins TSPAN14 and TSPAN15, whereas the 97 higher on the compact glandular side include 49 on 20q whose expression rises with 20q gain (among them RALY, PRPF6, EIF2S2, EIF6, PSMA7, CSE1L and RAE1; partial ρ −0.57 for RALY and PSMA7) and 21 without a gene annotation. The score thus contrasts a cytoskeleton–adhesion program with the expression consequence of 20q gain, and because no exon usage corresponded to morphology, it marks the expression state that accompanies a morphology rather than genes shown to cause it.

The organoid morphology genes also corresponded to histology in patient tumors. In 372 primary TCGA-COAD/READ tumors, the morphology gene score distinguished mucinous carcinomas, a tumor type with abundant extracellular mucus (43 tumors; AUC 0.83, P = 1.4×10⁻¹Z; Figure 2D), and this was not explained by microsatellite instability (MSI; Section R-26). The score ran opposite to the amount of 20q (ρ = −0.52) and was unrelated to the p53 activity score within TP53 wild-type tumors, but it was higher in wild-type than in mutant tumors (medians 0.67 versus −0.31, P = 2.5×10⁻¹⁶; Figure 2E), within the CIN and genomically stable (GS) subtypes but not within the MSI subtype, and after adjustment for 20q and MSI (P = 1.5×10⁻⁴; 335 tumors); mucinous carcinomas were more frequent among wild-type tumors (P = 7.8×10⁻⁵), in line with the fewer p53-pathway alterations reported in mucinous colorectal carcinomas [14]. In contrast, P2_index did not differ between mucinous and other carcinomas or among molecular subtypes, and after adjustment for TP53 status, 20q and MSI it ran opposite to the morphology gene score (P = 1.9×10⁻¹¹), possibly reflecting tissue composition (Section 3.2.3). Likewise, in the discovery organoids, the Type1 fraction differed by TP53 status after adjustment for 20q and aneuploidy (an arm-level, expression-based proxy for chromosome-arm gains and losses; P = 0.013), P2 dominance did not contribute after accounting for TP53 status and 20q (P = 0.97; Additional file 7: Supplementary Table S48), a test with little power because the two axes disagreed in only five of the 56 specimens, and in the patient-level sensitivity analysis (patient means of the 19 patients with uniform TP53 status) P = 0.13. The gastric-metaplasia programs in tumors are examined in Section 3.1.5.

To ask which expression programs follow TP53 status and which follow P2 usage, we scored previously defined gene sets in the discovery organoids, the GSE294511 organoids (Section 3.1.5) and TCGA, and compared the effect of TP53 status adjusted for the P2 measure (the array P2 index in organoids, P2_index in TCGA) with that of the P2 measure adjusted for TP53 status (rank regression; Section 2; Figure 2F; Additional file 7: Supplementary Table S48, sheet H). The morphology gene score followed TP53 status in all three cohorts, as did the image-based non-Type1 fraction in the discovery cohort, and so did the gastric-metaplasia genes of the P2 signature and the nine-gene gastric set in GSE294511 and TCGA, but not in the discovery organoids (56 specimens, 14 wild type). None of the morphology, metaplasia, gastric, P1 or SNORD116 scores was significantly positively associated with the P2 measure after adjustment for TP53 status or within wild-type samples. By contrast, the p53-target module of the P2 signature (MDM2 itself excluded) followed the P2 measure even after adjustment for TP53 status in both organoid cohorts (P = 5.8×10⁻⁴ and 2.7×10⁻⁵). Because this module was selected from the P1-versus-P2 comparison, we also tested the pre-specified 12-gene p53 activity score, which shares no selection step with it: it followed the P2 measure beyond TP53 status in GSE294511 (P = 0.012) but not significantly in the discovery organoids (P = 0.094; sheet H). In bulk tumors the p53-target module ran opposite to P2_index after adjustment for TP53 status; within TP53-mutant TCGA tumors the P2_index was inversely correlated with the 12-gene p53 activity score (ρ = −0.347, P = 3.7×10⁻⁸; sheet H), a relationship absent in mutant organoids (ρ = 0.05 in GSE294511; sheet F; 0.13 with the score rescored within the cohort, sheet H) and possibly reflecting tissue composition (Section 3.2.3). These results held at a false discovery rate below 0.05 across the programs of each cohort (all P values in sheet H; Section R-26); the P1 signature did not follow TP53 status consistently, and the SNORD116 cluster was associated with neither axis consistently.

We also asked whether the genome-wide expression differences between tissue 1 and tissue 2 (the 1,890 differentially expressed genes, DEGs) follow TP53 status or promoter usage. Because TP53 status and promoter usage disagreed in only five of the 56 specimens with determined status, this comparison has little power and is exploratory: the two sets of differences were closely correlated (ρ = 0.885), and genome-wide 55 genes followed TP53 status after adjustment for promoter usage and 145 followed promoter usage after adjustment for TP53 status (q < 0.05). Of 13 p53-target genes whose expression predicted sensitivity to the p53–MDM2 inhibitors NVP-CFC218 and NVP-CGM097 (drugs that block MDM2 binding to p53) better than TP53 status alone [17], 8 were among the 145 and none among the 55 (one-sided Fisher’s exact P = 8.9×10⁻¹⁵; Figure 2G). Because promoter usage was called from MDM2 exons and P2 is itself p53-inducible, this enrichment is not independent of the grouping; it nevertheless agrees with the program-level result above, whereas the bulk of the DEG program could not be assigned to either axis in this cohort (Section R-26).

#### 3.1.5 Morphology and metaplastic programs in an independent organoid biobank and in tumors

To test the main correspondences in independent patients, we analyzed the public HTA2.0 arrays of 119 organoids from 50 patients with colorectal liver metastases, established at another institution (Kryeziu et al. [36]; GEO GSE294511), under an analysis plan fixed before the data were examined (Section 2). Organoids whose morphology gene score lay further on the non-Type1 side had a higher array P2 index (Spearman ρ = 0.22, one-sided P = 0.0085); patient means were also correlated (ρ = 0.30, P = 0.037), but the slope in a patient random-intercept model was not significant, and within patients the two measures moved together in about half. The correspondence therefore held across samples, was weaker between patients and was not detected within patients (Section R-27).

Morphology followed TP53 status here as well: among the 88 organoids with published TP53 status, the morphology gene score lay further on the non-Type1 side in wild-type organoids (medians 1.11 versus −0.19, P = 1.9×10⁻⁵; patient means, P = 0.0029, 49 patients with uniform status; Figure 2E), also after adjustment for 20q and aneuploidy (P = 6.1×10⁻⁴), whereas the array P2 index did not contribute after accounting for TP53 status, 20q and aneuploidy (P = 0.51); given the weak correlations within each genotype group, the across-sample correspondence between morphology and P2 is largely explained by TP53 status (Additional file 7: Supplementary Table S48).

What replicated in this biobank was the expression program rather than the histological appearance, and in fact the two run in opposite directions: the non-Type1 pole of the morphology gene score, which marks the cystic–mucinous side in our image classifier, mapped here to the “solid” category of hematoxylin and eosin (H&E) histology (the gene score tended to be higher in solid organoids, P = 0.064), the published H&E categories did not differ in array P2 index, and the CDX2–KRT20 difference ran opposite to the array P2 index (ρ = −0.40, P = 8.9×10⁻⁶). The two morphological axes are thus only partly aligned; what reproduced across cohorts is the morphology gene program and its association with TP53 status, not the histological morphology itself (Section 3.1.4; Section 4.2.1). The gastric-metaplasia programs behaved likewise in tumors. In TCGA, the gastric-metaplasia genes of the P2 signature remained associated with TP53 status after further adjustment for 20q and MSI (P = 3.2×10⁻⁵), whereas the nine-gene gastric set did not (P = 0.18; Additional file 7: Supplementary Table S48, sheet H2, covariate-adjusted columns). With independent TP53 calls (Additional file 6: Supplementary Table S47), further adjustment for BRAF mutation and CIMP left the TP53 coefficients essentially unchanged (morphology gene score P = 5.8×10⁻⁵; metaplasia genes P = 1.8×10⁻⁴), whereas the nine-gene gastric set followed CIMP-high status rather than TP53 status; among tumors that were not of the MSI subtype, BRAF wild-type and CIMP-low, both scores remained higher in wild-type tumors (Section R-26). The organoid morphology program is thus reproduced in patient tumors as the histotype of mucinous carcinoma, and it follows TP53 status even after accounting for chromosomal instability, as do the metaplasia genes in tumors.

Morphology and the metaplastic programs therefore follow TP53 status rather than MDM2 promoter usage itself. We next ask whether MDM2 P2 promoter usage reflects p53 activity rather than genotype (Section 3.2).

### 3.2 MDM2 P2 promoter usage as a readout of p53 activity

Section 3.1 showed that morphology and the metaplastic programs follow TP53 status. We now ask whether MDM2 P2 promoter usage reports p53 genotype or p53 activity itself. Because the P2 signature contains p53 targets (Section 3.3.2), we measured p53 activity directly from 12 direct p53 targets (the p53 activity score; Section 2) rather than inferring it from genotype, and we quantified promoter usage with three cohort-specific measures, always naming which is meant: the array P2 index (exon-2 versus exon-1 signal on HTA2.0) in the discovery organoids and the independent biobank, the transcript-ratio P2_index (ΣTPM(P2)/ΣTPM(P1+P2)) in TCGA-COAD/READ tumors, and the cell-line P2 index (DepMap transcript quantification) in cell line panels.

#### 3.2.1 P2-dominant organoid specimens are p53-active, and P2 is induced in a p53-dependent manner

Among the 56 discovery specimens with determined TP53 status, the median p53 activity score was −0.30 in 42 TP53-mutant specimens, −0.35 in three wild-type P1-dominant specimens and +1.45 in 11 wild-type P2-dominant specimens (Figure 3A). Wild-type P2-dominant specimens were p53-active compared with mutants (P = 6.8×10⁻⁷, AUC 0.99; at the patient level, P = 5.2×10⁻⁴). None had a score at or below the 90th percentile of mutants, whereas two of the three wild-type P1-dominant specimens, both from HCT59 (variant of uncertain significance F113V), did (wild-type P2-dominant versus wild-type P1-dominant, P = 0.005). TP53 status and promoter usage largely coincided (P1-dominant, 40 mutant and 3 wild-type; P2-dominant, 2 mutant and 11 wild-type; Section 3.1.3), so genotype and activity cannot be fully separated here. Two observations nevertheless favor activity. First, the only mutant specimens with clearly positive scores were the two P2-dominant ones (HCT41-1T, +1.09, with R158H in its variant table; HCT38-3LM, +0.60, called from the published mutation maps of the same biobank [12,24] in the absence of a variant table; both above the 90th percentile of mutant specimens). Second, across the 56 specimens the array P2 index correlated with the p53 activity score (ρ = 0.53, P = 2.6×10⁻⁵). Within the 14 wild-type specimens, the array P2 index and the p53 activity score were correlated (ρ = 0.69, P = 0.0065; Figure 3B), but total MDM2 expression was correlated to a similar degree (ρ = 0.71), and neither P2 dominance nor the continuous array P2 index added significantly to total MDM2 (RZ 0.50 to 0.59, P = 0.15; 0.50 to 0.52, P = 0.50; Additional file 7: Supplementary Table S48). In each of the three patients with both P1 and P2 specimens (HCT38, HCT41 and HCT67), the P2 specimen had the highest p53 activity score (Figure 3C). The p53 suppressors MDM4, PPM1D and USP7 did not differ between groups (wild-type P2-dominant versus mutant specimens, difference in median log2 expression −0.19 to +0.03, P = 0.13 to 0.71; Kruskal– Wallis across the three groups, P ≥ 0.14; Additional file 2, R169). Assigning the seven specimens with undetermined TP53 status (all P1-dominant) to the mutant group left these results unchanged; assigning them instead to the wild-type group gave 10 wild-type P1-dominant specimens, eight at the mutant level (wild-type P2-dominant versus P1-dominant, P = 1.2×10⁻⁴; P2 dominance beyond total MDM2, P = 0.021; TP53 calls in Additional file 5: Supplementary Table S11). The P2 signature shares genes such as CDKN1A, BAX and BBC3 with the p53 activity score, and the P2-dominant group was defined in part through differences in p53 target expression, so the separation may partly reflect circularity.

**Figure 3.**
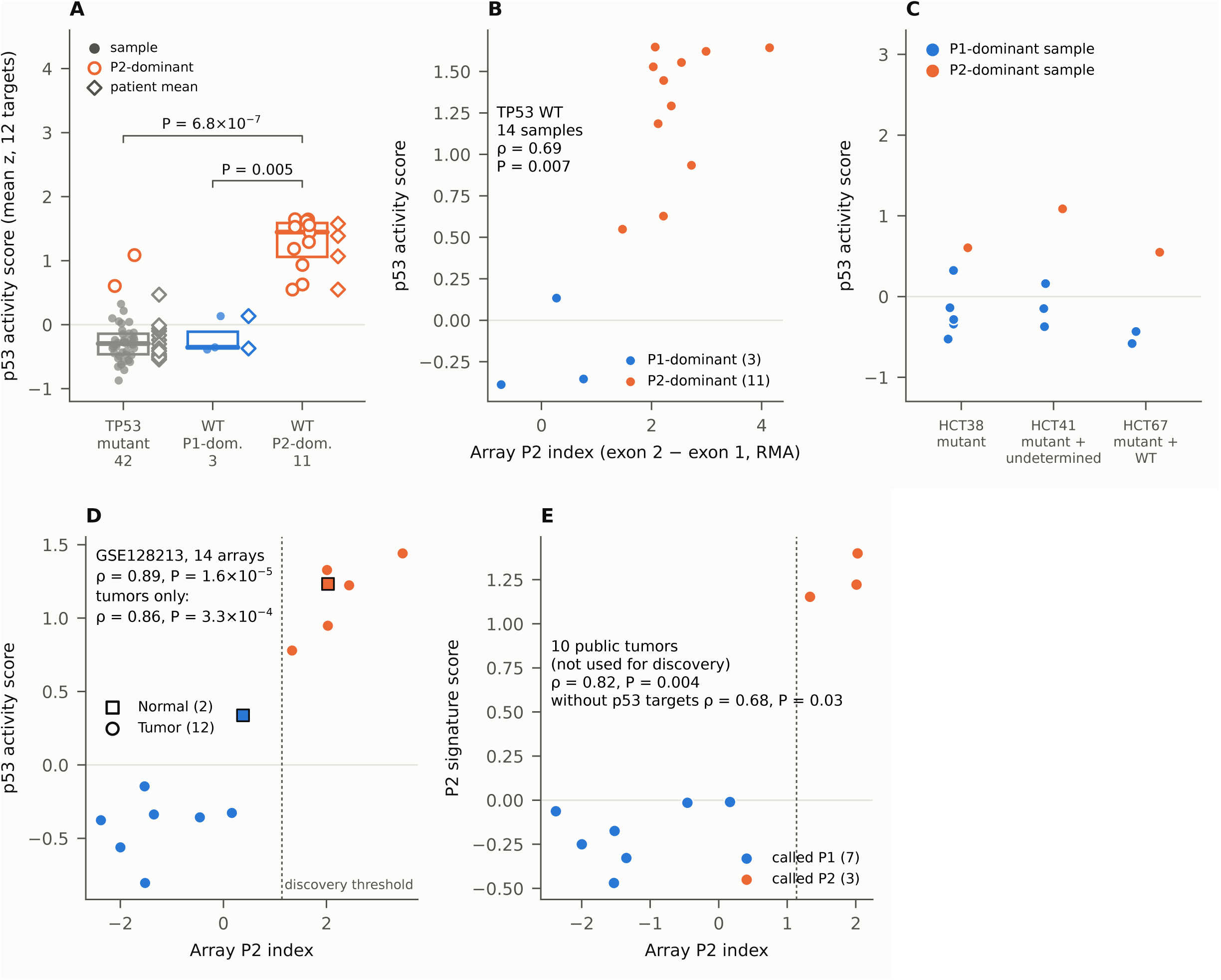
p53 activity and MDM2 P2 in organoids. (A) p53 activity score (mean z-score of 12 direct p53 targets) in TP53-mutant, wild-type P1-dominant and wild-type P2-dominant specimens (56 with determined TP53 status). Dots, specimens; open orange circles, P2-dominant specimens (the two in the mutant column carry TP53 mutations); diamonds, patient means; P values, wild-type P2-dominant versus TP53-mutant specimens (6.8×10⁻⁷) and versus wild-type P1-dominant specimens (0.005) (Mann–Whitney). (B) Array P2 index and p53 activity score in the 14 TP53 wild-type specimens. This index (Section 2) is the per-specimen mean RMA value of exon-2 minus that of exon-1 probe sets, each relative to the MDM2 transcript cluster. (C) The three patients with both P1 and P2 specimens (HCT38, HCT41 and HCT67). (D) Array P2 index and p53 activity score in 14 public HTA2.0 arrays not used for discovery (2 normal and 12 tumor; identical to GSE128213 files); dotted line, discovery threshold. (E) Array P2 index and P2 signature score in the 10 tumor arrays (P2/P1 calls by the discovery threshold).

In the 14 public HTA2.0 arrays not used for discovery (2 normal and 12 tumor; GSE128213), the array P2 index and the p53 activity score were again strongly correlated (ρ = 0.89, P = 1.6×10⁻⁵; Figure 3D). Two of these arrays carry the titles of discovery specimens of patient HCT33 (excluded from the replication of Section 3.1.2); both had a high array P2 index and a high p53 activity score, as in discovery. The correspondence with the P2 signature in the same 10 tumor arrays is shown in Figure 3E.

To test whether p53 activity drives P2 usage, we counted promoter-specific reads at the MDM2 locus in public RNA-seq of cells treated with nutlin (10 µM, 12 h) [38] (Figure 4A; Section 2). In TP53 wild-type HCT116 cells, nutlin increased reads on the P2-only first exon 29.5-fold but reads on the P1-only exon 1 only 2.8-fold, so that the P2/P1 ratio rose 10.5-fold (Figure 4B), whereas in TP53-null HCT116 cells neither the exons nor the ratio changed; MCF7 and SJSA showed the same direction, under DMSO P1 reads predominated in every line, and gene-level induction of MDM2 and CDKN1A was seen only in TP53 wild-type cells (nutlin versus DMSO, mean of two replicates: CDKN1A 25.3-fold and MDM2 16.7-fold in TP53 wild-type HCT116 but 1.7- and 1.2-fold in TP53-null HCT116; CDKN1A 28.6- and 45.9-fold and MDM2 18.9- and 42.9-fold in SJSA and MCF7; Figure 4C; Section R-25). When p53 is active, MDM2 transcription thus shifts strongly toward P2, so that P2 usage becomes a measurable indicator of p53 transcriptional activity; on the measures used here, this agrees with the known status of P2 as a direct target carrying p53 response elements [5,6]. At the level of transcription initiation, ENCODE PRO-cap of six normal colon and stomach tissues gave a median P2 share of 0.506 and TP53-mutant Caco-2 cells 0.565, whereas the P2 share of MDM2 mRNA for the same line in DepMap was 0.173 (Section 2; Additional file 7: Supplementary Table S48, sheet K); initiation at P2 was thus not reduced in this TP53-mutant line (interpretation in Section 4.2).

**Figure 4.**
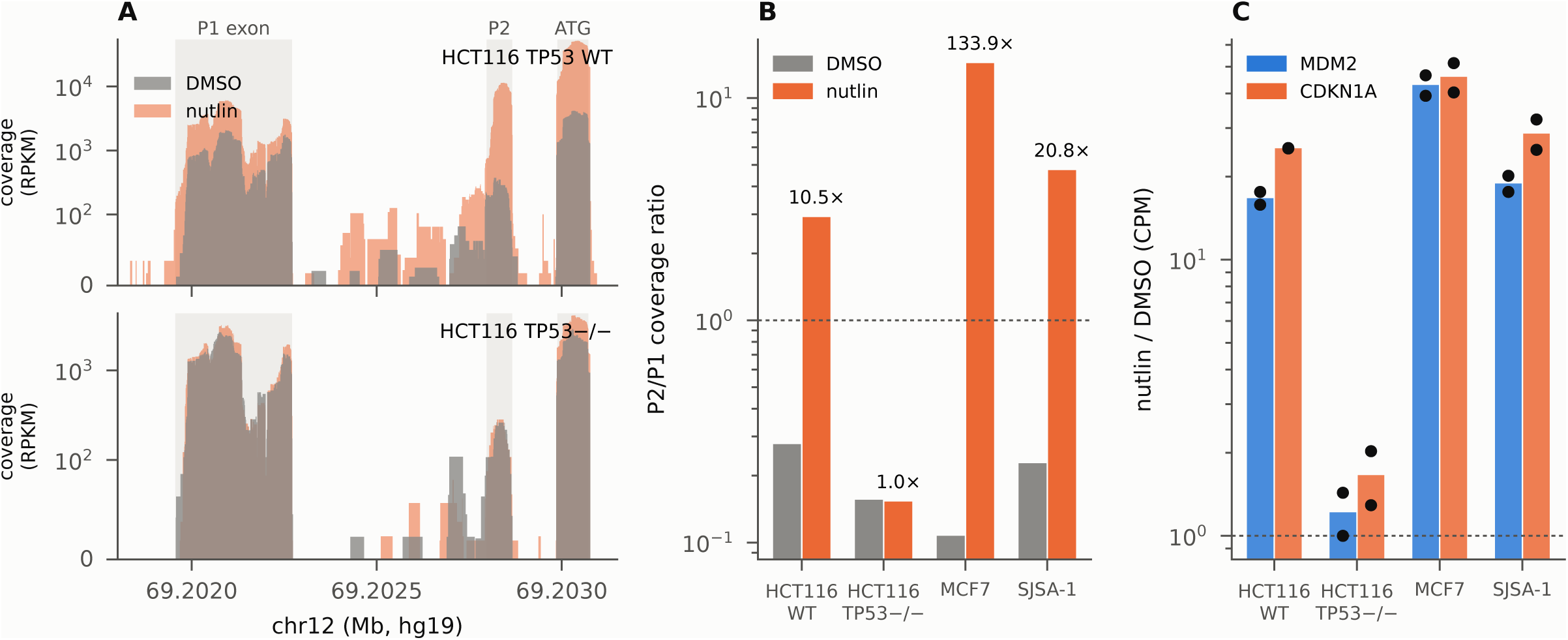
P2 is selectively induced in a p53-dependent manner (public RNA-seq, GSE86221). (A) Coverage at the MDM2 locus (P1-only exon 1, P2-only first exon and shared ATG exon). (B) P2/P1 ratio in four cell lines (HCT116 TP53 wild type and null, MCF7 and SJSA; DMSO or nutlin 10 µM for 12 h). (C) Gene-level induction of MDM2 and CDKN1A (two replicates).

#### 3.2.2 Direct quantification of P2 usage and the TP53 functional-class gradient in tumors

To examine promoter usage itself without relying on downstream signatures, we quantified it at the transcript level in TCGA-COAD/READ (P2_index). The P2_index was higher in TP53 wild-type than in mutant tumors (medians 0.481 in 132 versus 0.404 in 242; Wilcoxon P = 1.0×10⁻⁵; 374 analyzed), which fits functional wild-type p53 driving the p53-responsive P2 promoter; adjacent normal colorectal mucosa used mainly P1 (median P2_index 0.289, n = 51), and both wild-type and mutant tumors were above it (in 32 patient-matched pairs the tumor was higher in 23, Wilcoxon P = 6.1×10⁻⁴; Additional file 7: Supplementary Table S48, sheet I). The P2_index correlated positively with the p53 target module excluding MDM2 (Spearman ρ = +0.248, P = 1.1×10⁻⁶, n = 380) and, unadjusted, with the gastric/Paneth metaplasia module (Section 3.1.4), but not with the P1 signature; it differed among CMS subtypes (Kruskal–Wallis P = 1.48×10⁻⁴, n = 342; Figure 5), CMS4 being lowest, but did not differ significantly by MSI status in the small annotated subset of 32 specimens or among the CIN, GS and MSI subtypes, so MSI is not a surrogate for P2 usage, and it was not associated with overall survival (368 tumors; log-rank P = 0.82). P2 shares counted directly from recount3 junction reads were likewise higher in wild-type tumors (medians 0.316 versus 0.000, P = 2.8×10⁻¹⁶) and agreed with the P2_index in the tumors with 10 or more reads (ρ = 0.64; junction reads sparse, median 7 per tumor; sheet D). In a multivariable model, wild-type status remained associated with the P2_index (standardized coefficient +0.580, P = 5.3×10⁻⁷), the partial correlation with the p53 target module adjusted for purity and CMS was maintained (ρ = +0.253), and confounding by MDM2 amplification — which in astrocytic tumors preferentially drives P1 [90] — was excluded by adjustment and stratification. Overall, MDM2 promoter usage itself is linked, with modest effect sizes, to TP53 status and to the p53 target and P2 programs in an independent cohort (full analysis in Section R-12).

**Figure 5.**
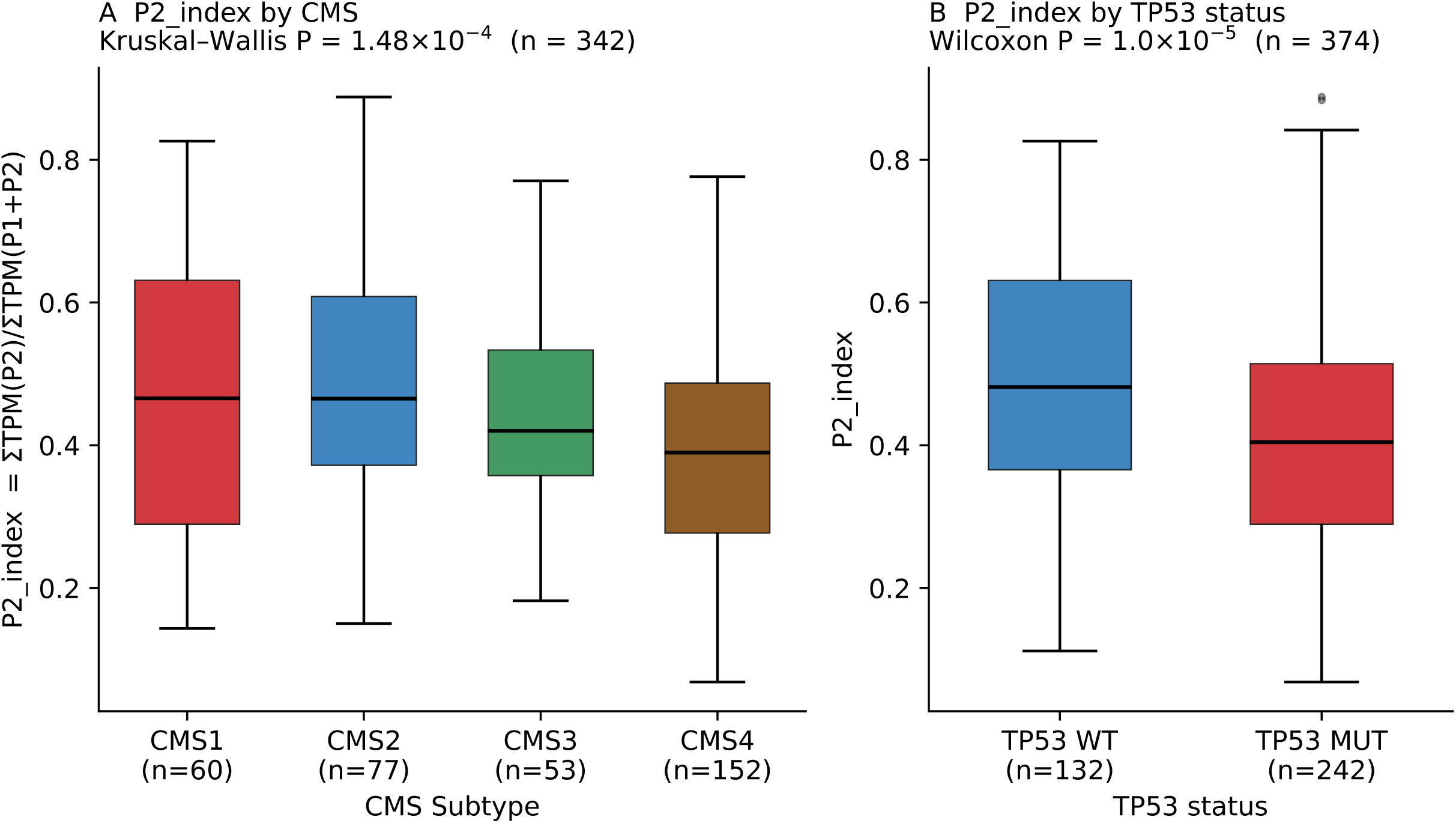
MDM2 P1/P2 usage (P2_index) by CMS subtype and TP53 status in TCGA-COAD/READ. (A) P2_index by CMS subtype (Kruskal–Wallis P = 1.48×10⁻⁴, n = 342). The difference arises because CMS4 is lowest; CMS1, CMS2 and CMS3 do not differ significantly. (B) P2_index by TP53 status: higher in wild-type tumors (Wilcoxon rank-sum P = 1.0×10⁻⁵, n = 374; 132 wild-type and 242 mutant). The 6 specimens without somatic mutation data were excluded, not assigned to wild type. Numerical details are in Section 3.2.2.

We annotated the 374 specimens with the TP53 Database (release 21) [66,67] using prespecified rules and two parallel gain-of-function definitions [68,69]. The P2_index differed among wild-type, GOF and loss-of-function (LOF) classes (Kruskal–Wallis P = 1.3×10⁻⁵ and 3.9×10⁻⁶ under the two definitions), with a gradient of truncating (median 0.364) < missense (0.423/0.422) < wild-type (0.481) preserved under both (LOF versus GOF, P = 0.015 and 0.013), whereas the p53 target module output differed only between wild-type and mutant, with no step within the mutant group. The P2_index therefore carries information not explained by p53 target output alone; the finding is, however, a truncating < missense < wild-type gradient, not a GOF-specific effect (hotspot versus non-hotspot missense not different; Additional file 29: Supplementary Table S37; full analysis in the Supplementary Report, Section R-15).

#### 3.2.3 P2 usage and p53 activity in bulk tumor tissue

In TCGA-COAD/READ, the P2_index showed a gradient across TP53 functional classes (Section 3.2.2; Figure 6A). Across all 132 TP53 wild-type tumors, its correlation with the p53 activity score was weak and depended on how expression was quantified (ρ = 0.19 with TCGAbiolinks expression; ρ = 0.03 with the Toil recompute; Additional file 7: Supplementary Table S48). We excluded hypermutated tumors (≥ 400 non-synonymous mutations; Section 2) from the next comparison, because among the 132 wild-type tumors the 41 hypermutated ones had a lower p53 activity score than the 91 non-hypermutated ones (median 0.13 versus 0.40, Mann–Whitney P = 0.008; with the Toil recompute 0.18 versus 0.41, P = 0.012; Additional file 7: Supplementary Table S48, sheet N); this restriction is an analytic choice, so the values without it are given above, and all 12 combinations of TP53 call, hypermutation definition and expression source are given in sheet N. In the 82 tumors that were not hypermutated and were confirmed as wild type in the MC3 calls, the P2_index and the p53 activity score were correlated (ρ = 0.42, P = 1.0×10⁻⁴; score higher in the middle and upper P2_index tertiles than in the lower; Figure 6B); this ρ = 0.42 is the upper end of a definition-dependent range (sheet N). In the same 82 tumors, total MDM2 expression followed the p53 activity score more closely (ρ = 0.57), and adding the P2_index did not improve the fit; across 12 combinations of TP53 call, hypermutation definition and expression source, ρ ranged from 0.03 to 0.42, and the P2_index added no significant positive information beyond total MDM2 in any of them (sheet N). In other words, in bulk tumors, P2 usage did not carry information on p53 activity beyond the amount of MDM2 transcript; the same held at the protein level, where BAX protein (reverse-phase protein array data of 69 tumors, via UCSC Xena [43]) followed the P2_index (ρ = 0.33) only through total MDM2 (partial ρ = 0.09 after adjustment for total MDM2; sheet E; Section R-25). The 12-target p53 activity score agreed with a published transcriptome-based classifier of p53 function: in the 118 wild-type tumors with a TP53-loss phenocopy score [20], the two were inversely correlated (ρ = −0.49, P = 2.0×10⁻⁸; AUC 0.80 for phenocopy status), whereas the classifier of Li et al. [21] called nearly all wild-type tumors functionally deficient (sheet J); the P2_index also correlated with the phenocopy score, but not after adjustment for total MDM2. The P2_index was inversely correlated with an immune-cell marker (PTPRC; ρ = −0.47 in wild-type tumors) but only weakly with tumor purity, so bulk P2_index may be influenced by tissue composition; generalization beyond colorectal cancer, and why some wild-type tumors lack p53 activity, are beyond the scope of this paper (Section 4.6) and are the subject of a companion study on p53 activity in TP53 wild-type tumors (T. Tsukui, R. Yao, and K. Tsuda, unpublished observations).

**Figure 6.**
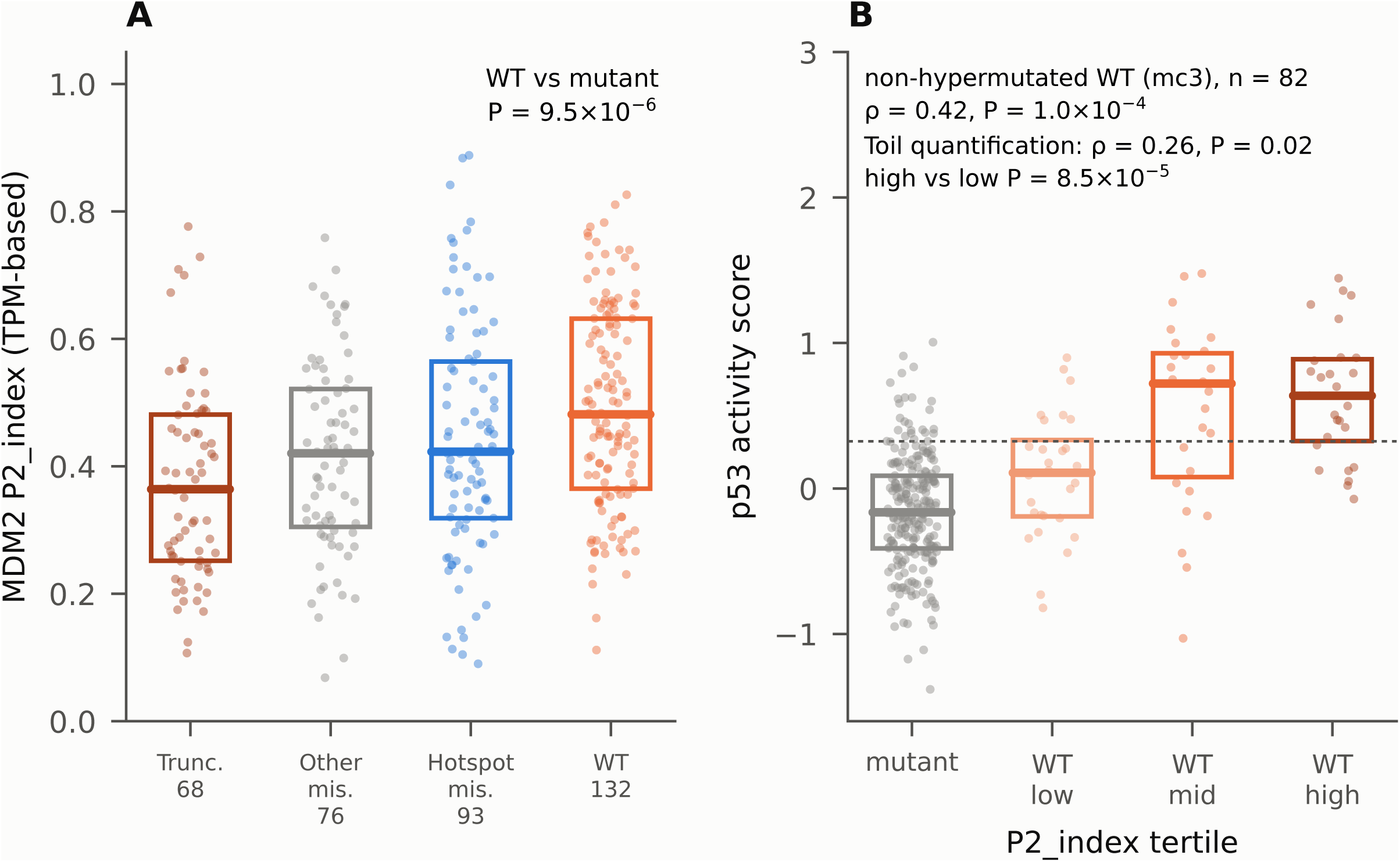
P2 usage and p53 activity in TCGA-COAD/READ. (A) P2_index by TP53 functional class (truncating, other missense, hotspot missense and wild type). Under definition A of Section 3.2.2, truncating is LOF and hotspot missense GOF; definition B counts as GOF all missense variants classed as non-functional in the TP53 Database. The 369 tumors with an assigned functional class are plotted. The in-figure wild-type-versus-mutant P (9.5×10⁻⁶) was computed on this plotting data set and differs slightly from the value in Section 3.2.2 (P = 1.0×10⁻⁵, n = 374). (B) p53 activity score across P2_index tertiles in the 82 tumors that were not hypermutated and were confirmed as wild type in MC3 (mutant status by MC3; dotted line, 90th percentile of mutant tumors; ρ with the Toil recompute shown alongside). The Spearman ρ of 0.42 shown is the upper end of a definition-dependent range (0.03–0.42 across 12 combinations of TP53 call, hypermutation definition and expression source; Additional file 7: Supplementary Table S48, sheet N).

#### 3.2.4 P2 usage, MDM2 dependency and MDM2-inhibitor sensitivity in cell lines

From DepMap transcript quantification [47,48] we computed the cell-line P2 index of 1,517 lines. In the 691 lines with known TP53 status it was higher in wild-type lines (median 0.231 versus 0.104, P = 1.8×10⁻33) and correlated with the p53 activity score among the 208 wild-type lines (ρ = 0.57, P = 1.4×10⁻¹⁹; Figure 7A); in six lines also sequenced with short and Nanopore long reads [58], the P2 share counted directly from MDM2 junction reads ranked the lines almost identically (ρ = 0.94 and 0.83; Additional file 7: Supplementary Table S48, sheet G). In TP53 wild-type lines, a higher cell-line P2 index was associated with greater sensitivity to nutlin-3a (GDSC1, 198 lines, ρ = −0.26, P = 1.7×10⁻⁴) [49,50], also after adjustment for the p53 activity score (partial correlation −0.19, P = 0.009; GDSC2, −0.15; Figure 7B), and with greater MDM2 dependency in CRISPR knockout screens [52,54] (364 lines, ρ = −0.33, P = 8.1×10⁻¹¹; tertile medians −0.77, −1.19 and −1.24; partial correlation adjusted for p53 activity −0.12, P = 0.022; Figure 7C). Total MDM2 expression, however, was associated with nutlin-3a sensitivity about as strongly, and after adjustment for it the P2 association weakened (GDSC1 partial correlation −0.15, P = 0.035; GDSC2 and nutlin-3 in CTRPv2 [55,56], not significant), whereas for MDM2 dependency the cell-line P2 index remained associated after adjustment for total MDM2 (partial correlation −0.27, P = 1.1×10⁻⁷; Section R-25).

**Figure 7.**
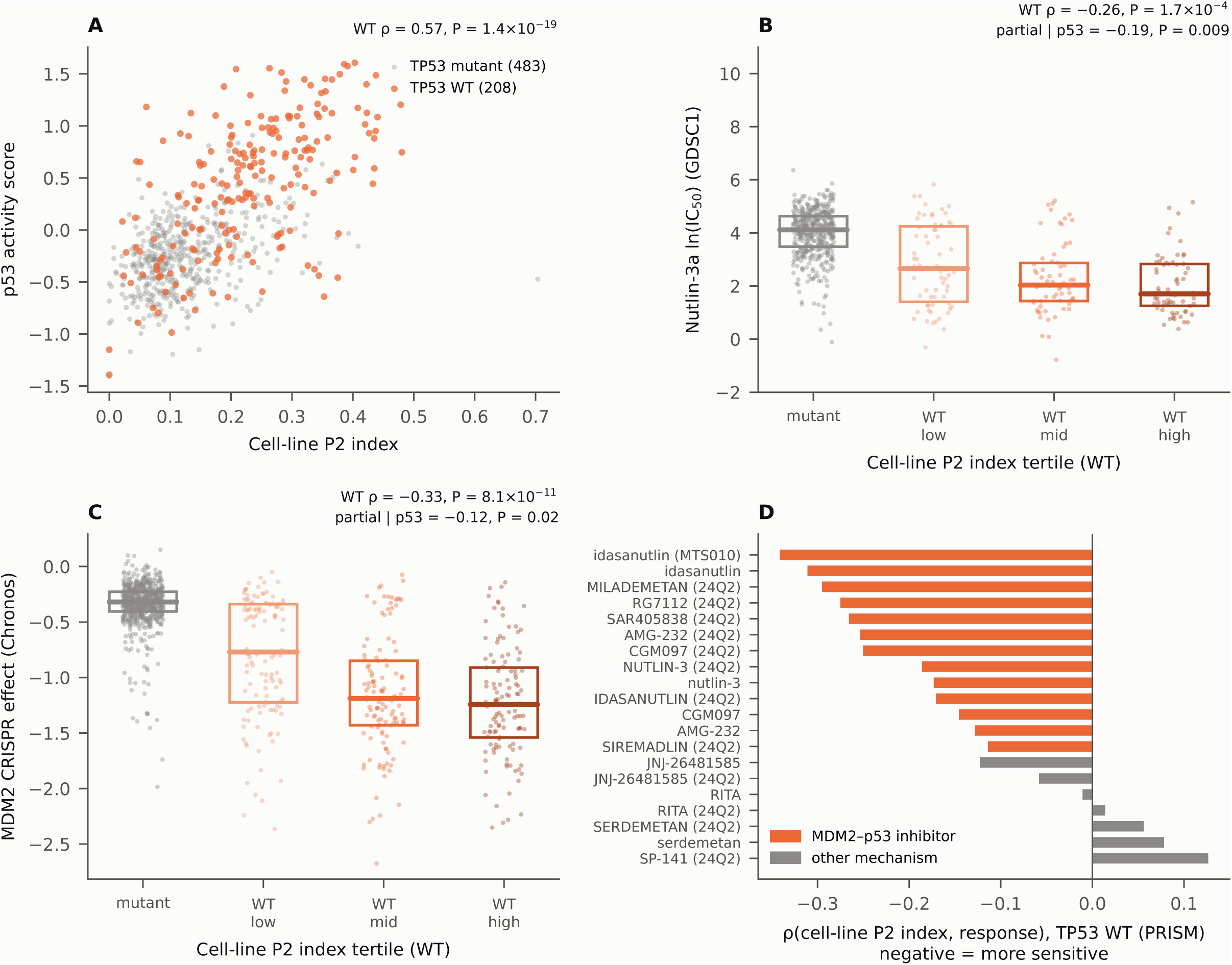
In cell lines, P2 follows MDM2 dependency and, less specifically, MDM2 inhibitor sensitivity. (A) Cell-line P2 index and p53 activity score (TP53 wild type, n = 208, orange; TP53 mutant, n = 483, gray). The index is ΣTPM of P2 transcripts/ΣTPM of P1 + P2 transcripts, computed as P2_index in TCGA (Section 2). (B) nutlin-3a ln IC50 by cell-line P2 index tertile (GDSC1). (C) MDM2 dependency in CRISPR screens (DepMap, Chronos). (D) Correlation by drug in PRISM (MDM2 inhibitors that release p53, and control drugs with other mechanisms).

This relationship was specific to the mechanism of action. In the PRISM drug screens [51], all eight agents that block the MDM2–p53 interaction and release p53 (idasanutlin, AMG-232, CGM097, SAR405838, RG7112, milademetan, nutlin-3 and siremadlin) were in the negative direction, seven at P < 0.05 in the primary screen and strongest for idasanutlin in the secondary screen, whereas drugs with other mechanisms (RITA, serdemetan, SP-141 and others) showed no association (Figure 7D). In PRISM, however, the association largely disappeared after adjustment for the p53 activity score, so most of the information in P2 overlaps with p53 activity, and the part beyond it was seen only in GDSC and CRISPR. TP53 wild-type lines without p53 activity (about 30%) were less sensitive to nutlin-3a (GDSC1: median ln IC50 3.12 versus 1.74, P = 3.8×10⁻⁷), also in lines confirmed as wild type by DepMap mutation data alone, so this is not explained by the misclassification of mutant lines raised as a concern [18]; for nutlin-3 in CTRPv2 the within-wild-type association did not reach significance (ρ = −0.14, P = 0.07). Section 3.2.5 examines organoids from independent patients. As context, we confirmed the well-established TP53-dependence of MDM2-inhibitor sensitivity, using wild-type TP53 as an imperfect proxy for p53 activity (not all wild-type tumors are p53-active, Section 3.2.3; Additional file 30: Supplementary Figure S10): wild-type lines were more sensitive to nutlin-3a (pan-cancer median LN_IC50 2.672 versus 5.167, Mann–Whitney P = 1.5×10⁻⁶¹; same direction in colorectal lines), and in the DepMap CRISPR screen (1,178 lines) MDM2 dependency was higher in wild-type lines (Additional file 31: Supplementary Table S38; Section R-13). In GDSC2, colorectal MSI-High lines were also more sensitive to nutlin-3a than MSS/MSI-L lines (median LN_IC50 2.489 in 15 versus 5.862 in 32 lines, P = 5.2×10⁻3; Supplementary Figure S10), a background observation consistent with their more frequent wild-type TP53; MSI-High status is not a proxy for P2 usage (Section 3.2.2). In a biobank of 256 patient-derived tumor organoids (Herranz-Ors et al. [13]), MDM2 dependency was likewise higher in wild-type colorectal organoids (effect size 0.463, adjusted P = 0.001), and in our independent re-analysis of its primary data (65 colorectal lines) the median nutlin-3 log(IC50) was 1.386 in the 17 wild-type versus 4.283 in the 48 mutant lines (exact Wilcoxon P = 7.4×10⁻⁸, Cliff’s delta = −0.804), preserved under adjustment for KRAS, MSI and MDM2 gain-of-function. Our P2 signature score was associated with nutlin-3 sensitivity (p53 target module excluding MDM2: ρ = −0.539, FDR = 5.1×10⁻⁵), but the partial correlation adjusted for TP53 status did not survive multiple-testing correction (ρ = −0.331, raw P = 0.0071, FDR = 0.056): the P2 axis was associated with MDM2-inhibitor efficacy in these 65 organoids but could not be shown to be independent of TP53 mutation status (per-line values in Additional files 32 and 33: Supplementary Tables S40 and S41; full analysis in Section R-13).

To evaluate drug sensitivity without preselecting drugs, oncoPredict [57] projected GDSC2 sensitivity (198 drugs) onto the 624 TCGA-COAD/READ specimens; the drug most strongly correlated with the P2 score was nutlin-3a (ρ = −0.659; negative = higher predicted sensitivity), ranking first of the 198 drugs and first on the P2–P1 separation index. An important limitation is that 171 of the 198 drugs reached FDR < 0.05 in the same direction, so the P2 score correlates with drug sensitivity in general (the rank, not the smallness of the nutlin-3a P value, is what should be interpreted), and because the P2 signature contains canonical p53 targets, a top rank for nutlin-3a is expected from its composition (Additional file 34: Supplementary Table S39; Section R-16). On measured data (GDSC2 cell-line expression versus LN_IC50, adjusting for cancer type and TP53 status), nutlin-3a again ranked first of 295 drug entries on the P2 score, and in DepMap the P2 score followed MDM2 dependency after accounting for lineage and TP53; on the P1 side, all seven inhibitors of the replication-stress checkpoint kinases ATR, CHK1 and WEE1 showed higher sensitivity with higher P1 scores (P = 4.3×10⁻⁷), whereas MEK/ERK/BRAF inhibitors unexpectedly correlated with the P2 score, and 5-FU, oxaliplatin and irinotecan showed no P1-specific association; these cell-line effects are small and exploratory (Supplementary Table S39, sheets S39b–S39d).

#### 3.2.5 p53 activity and idasanutlin sensitivity in an independent organoid biobank

In the GSE294511 biobank (Kryeziu et al. [36]), the array P2 index was higher in TP53 wild-type than in mutant organoids (P = 6.9×10⁻⁴) and, within the 14 wild-type organoids, was strongly correlated with the p53 activity score (ρ = 0.91, P = 5.5×10⁻⁶; discovery 0.69); total MDM2 also correlated with the score (ρ = 0.87), but the array P2 index remained associated after adjustment for it (partial ρ = 0.61, P = 0.028; patient means, 0.66, P = 0.053, 10 patients). The biobank authors reported idasanutlin to be more active in wild-type organoids [36]; among wild-type organoids, sensitivity followed the array P2 index (ρ = 0.70, P = 0.0049), the six with the lowest index being insensitive and the eight with an index above 0.3 sensitive. Across the 87 organoids with known TP53 status and idasanutlin sensitivity (49 patients), the array P2 index was associated with sensitivity after adjustment for TP53 status (P = 1.6×10⁻⁸; 0.0022 clustered by patient) and, more weakly, after further adjustment for the p53 activity score and the mean sensitivity to the 23 other drugs (clustered P = 0.069). Among wild-type organoids, however, total MDM2 followed idasanutlin sensitivity as closely (ρ = 0.69), and after adjustment for it the array P2 index was no longer associated (partial ρ = 0.26, P = 0.39); across the 87 organoids, total MDM2 remained (clustered P = 0.006) whereas the P2 term was weaker (clustered P = 0.16; Section R-27). Thus, in organoids from independent patients, P2 tended to remain associated with p53 activity after adjustment for total MDM2, whereas for MDM2-inhibitor sensitivity it could not be distinguished from total MDM2 among the few wild-type organoids, as in the cell-line results for nutlin-3a (Section 3.2.4).

The exon- and junction-level differences that accompanied P2 usage in the genome-wide exon-usage screen of the discovery cohort (39 probe sets in 11 genes, including FAS, ZMAT3, TNFRSF10C and ACTA2) all changed in the same direction with the array P2 index (39 of 39, P = 3.6×10⁻¹Z), as did 65% of the 1,890 DEGs and 31 of the 36 panel genes identifiable by symbol, and the gastric-metaplasia score was again higher toward P2 (ρ = 0.22; discovery 0.23). In the 50 matched tumor tissues, by contrast, the array P2 index was higher than in the organoids and its ranking agreed poorly with theirs, although within tissues the morphology gene score and the array P2 index were strongly correlated (ρ = 0.70); bulk tissue contains stroma and immune cells (Section 3.2.3; Section R-27). Taken together, P2 usage reflects p53 activity: it is induced in a p53-dependent manner, is graded with TP53 functional class, and follows MDM2 dependency even after accounting for total MDM2 transcript. In bulk tumors and for inhibitor sensitivity, however, its information rarely exceeds that of total MDM2, so its added clinical value over total MDM2 is limited. We next summarize the properties of the expression signatures that underlie both readouts (Section 3.3).

### 3.3 Characterization of the expression signatures

Sections 3.1 and 3.2 established two readouts, that is, measurable indicators: morphology and metaplastic programs, which follow TP53 status, and MDM2 P2 promoter usage, which reflects p53 activity. This section turns to the discovery-cohort signatures themselves. It characterizes the two expression programs that underlie those readouts.

#### 3.3.1 The two expression programs (tissue 1 and tissue 2)

The tissue-1 program centered on intestinal absorptive and metabolic identity. Markers that were highly reproducible across comparisons included OLFM4 (average fold change +1,252; median +4.0, the average being driven by the comparisons of one patient, HCT64; Additional file 26: Supplementary Table S26), SLC26A3 (+782), ALDH1A1 (+272), the brush-border enzymes MGAM2 (+87) and DPP4 (+25), and the crypt-base stem-cell marker LGR5 (+13) (average fold changes across the 33 comparisons; positive, higher in tissue 1; Additional file 8: Supplementary Table S4). They also included a set of proliferation- and mitosis-related genes biased toward tissue 1 (MYBL2 +9.66, TOP2A +6.54, FOXM1 +4.58, MKI67 +2.00). This set is the transcript-level readout of the MYC/E2F/FOXM1 axis (Table 2; WNT-pathway genes in Additional file 35: Supplementary Table S27). High OLFM4 is consistent with its role as an intestinal stem-cell marker [91,92] and with independent findings from the same biobank [12,24]. The full profile is given in the Supplementary Report (Additional file 1), Section R-5.

Pathway analysis matched the tissue-1 program to the CIN (CMS2) type, headed by “Processing of Capped Intron-Containing Pre-mRNA” (in 33/33 analyses). Upstream-regulator analysis (which infers regulators explaining the expression changes) adopted 121 tissue-1 activators, led by MYC (median z = +5.76, 32/33) and including E2F, FOXM1, MDM4, and AURKB (Table 3).

**Table 3.** Detailed comparison of the IPA Canonical Pathway analysis.

| Pathway category | Tissue 1 (major pathways) | Tissue 2 (major pathways) |
| --- | --- | --- |
| RNA processing, cell cycle, and DNA replication | Processing of Capped Intron-Containing Pre-mRNA (Rank1, 33/33); rRNA processing (Rank4, 29/33); rRNA modification (Rank10, 32/33); Cell Cycle Checkpoints (Rank2), Mitotic Metaphase/Anaphase/Prometaphase, Synthesis of DNA, DNA Replication Pre-Initiation, Regulation of Mitotic Cell Cycle, S Phase, G2/M phases | Cell Cycle G1/S Checkpoint Regulation, FOXO-mediated Transcription of Cell Cycle Genes, Oncogene Induced Senescence, Senescence Pathway |
| DNA repair | HDR (HRR/SSA), NHEJ, Mismatch Repair, Fanconi Anemia Pathway, Nucleotide Excision Repair, BER, DNA Double-Strand Break Repair, ATM Signaling | Ferroptosis Signaling Pathway (oxidative DNA damage), CGAS-STING Signaling (cytosolic DNA sensing) |
| Energy metabolism | Oxidative Phosphorylation, TCA Cycle, Pentose Phosphate Pathway, Citric acid cycle, Glycolysis, Mitochondrial translation, Complex I/III/IV assembly | Mitochondrial Dysfunction, HIF1α Signaling, Glycolysis I, NAFLD Signaling, Lipid metabolism |
| Inflammation and immunity | IL-12 Signaling, ISG15 antiviral mechanism, PKR-mediated signaling, TCR signaling, Regulation of Apoptosis. Note: the tissue-1 items in this row are comparison-specific findings based on comprehensive extraction and are not discussed in the main text or in Supplementary Note Section 2.1 (the same point as for the inflammation/immunity/ECM row of Table 2). | Pathogen Induced Cytokine Storm, IL-17 Signaling, NF-κB, Neuroinflammation, Immunogenic Cell Death, TNFR1/Death Receptor Signaling, Natural Killer Cell Signaling, Inflammasome, Neutrophil degranulation (Rank6 in the latter half, consistent in 30/33) |
| ECM, invasion, and EMT | (limited) | Extracellular Matrix Organization, Degradation of ECM, Regulation of EMT by Growth Factors, Integrin Signaling, Colorectal Cancer Metastasis Signaling |
| Growth factors and signaling | Hedgehog 'on/off', WNT/β-catenin, ESR-mediated signaling, EGFR, PPAR Signaling, Estrogen-mediated S-phase Entry | VEGF, ERBB, HER-2, NGF, IGF-1, JAK/STAT, MAPK/RAF, AKT/PIP3, IL-2/4/6/8/13/15/33, TGFB |
| Epigenetics | Nucleosome assembly, Chromatin organization, PRC2 methylates histones, DNA methylation, SUMOylation pathways, Histone Modification | (epigenetic alteration co-occurring with the inflammatory response) |
| Cancer-specific | Prostate/Chronic Myeloid Leukemia Signaling, SPINK1 Pancreatic Cancer, Senescence-Associated Secretory Phenotype | Colorectal Cancer Metastasis, SPINK1 General Cancer, FAT10 Cancer Signaling, Pancreatic Adenocarcinoma, Thyroid/Renal Cell Carcinoma, Tumor Microenvironment |

This table is a listing based on the rank order of frequency of appearance across the 33 analyses; no selection by effect size ( $|\text{median } z|$ ) was performed. For the median $z$ and sign consistency of each
pathway, see Additional file 41: Supplementary Table S6.

Genes consistently upregulated in tissue 2 included MDM2 (median linear fold change 6.0-fold, same direction in all 33 comparisons) and wild-type p53 target genes (CDKN1A/p21, DDB2, ZMAT3, FAS, GDF15, BAX, BBC3; median fold change from 13.0 for ZMAT3 and 10.5 for DDB2 to 3.4 for CDKN1A and 1.2 for BBC3; Table 4). They also included gastric metaplasia markers (CTSE, REN, TFF1, TFF3, ANXA10; for example, TFF1 with a median fold change of 82), Paneth-cell markers (DEFA5, DEFA6; median 117- and 138-fold), gel-forming mucins (MUC5AC, MUC5B, MUC6; MUC5B median 8.7-fold, MUC5AC and MUC6 less than 2-fold), and markers of inflammation, hypoxia, and invasion (Table 2; Table 4). The top tissue-2 differentially expressed genes (DEGs) were CTSE (−1,900.6), REN (−1,281.1), MUC17 (−477.6), TFF1 (−408.2), DEFA5 (−381.0), and the long noncoding RNAs (lncRNAs) UCA1 (−546.7) and HULC (−263.6) (average fold changes; negative, higher in tissue 2; the medians, for example −1,034 for CTSE and −35 for UCA1, are in Additional file 8: Supplementary Table S4). Together they describe a distinct program resembling the gastric mucosa. This program agrees in direction with the gastric pyloric-gland differentiation of serrated lesions (saw-toothed precursor lesions of colorectal cancer) and with gastric metaplasia as a point of origin of serrated tumorigenesis [93–99]. The 10-gene wild-type TP53 activation signature in Table 4 (Figure 8A) is a dedicated list chosen by explicit criteria: directional consistency ≥ 90%, direct p53 targets in the Fischer census [37], and use as indicators of wild-type p53 activity, P2 being a p53-inducible promoter [6,7]. The full gene-by-gene description is given in the Supplementary Report (Additional file 1), Section R-6.

**Figure 8.**
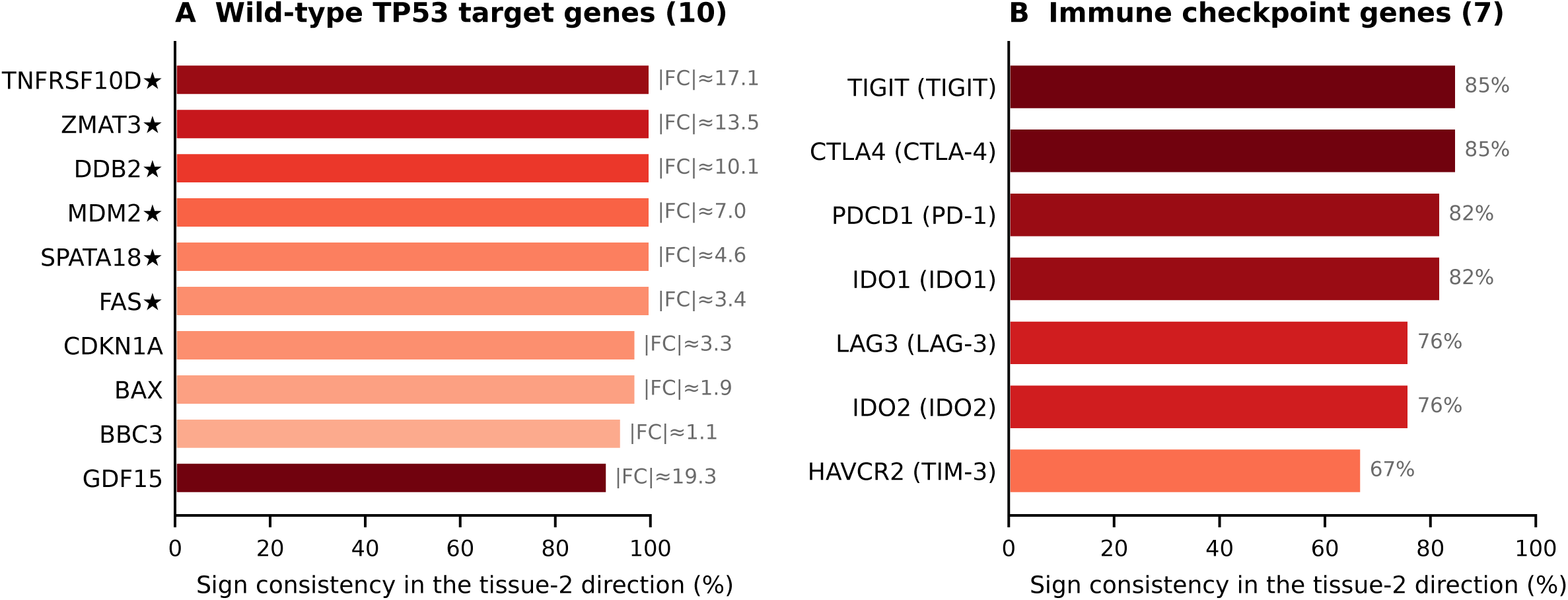
Wild-type TP53 pathway activation and expression of immune checkpoint molecules in P2 organoids. (A) The 10 wild-type TP53 target genes downstream of the MDM2 P2 pathway (TNFRSF10D, ZMAT3, DDB2, MDM2, SPATA18, FAS, CDKN1A, BAX, BBC3, GDF15) were all enriched on the tissue-2 side. Genes are in descending order of sign consistency in the tissue-2 direction, ties by descending |mean linear fold change|. Table 4 gives each gene’s mean and median fold change and sign consistency in the tissue-2 direction. Six genes (TNFRSF10D, ZMAT3, DDB2, MDM2, SPATA18 and FAS) were consistent in all 33 comparisons; CDKN1A and BAX were 97%, BBC3 94% and GDF15 91%. (B) Seven immune checkpoint molecules (TIGIT, CTLA4, PDCD1, IDO1, LAG3, IDO2, HAVCR2; same order rule: sign consistency in the tissue-2 direction, ties by descending |mean linear fold change|) also showed directionally consistent higher expression on the tissue-2 side. This is exploratory and below the DEG effect-size criterion. Consistency rates: TIGIT 84.8%, CTLA4 84.8%, PDCD1 81.8%, IDO1 81.8%, LAG3 75.8%, IDO2 75.8%, HAVCR2 66.7%. Values: panel A, Table 4; panel B, Additional file 46: Supplementary Table S30. Stars, genes consistent in all 33 comparisons; bar color, |mean linear fold change| in A (value at right) and sign consistency in B. |Mean linear fold change| is not printed in B because its values (−0.38 to −0.77) are below 1 in absolute value and would read as a decrease. This directional pattern provides a background for considering immunotherapies such as anti-PD-1, IDO1 inhibition, LAG-3 inhibition and TIGIT inhibition in cases with confirmed MSI-High/dMMR. However, this study does not claim that the P2-dominant group itself constitutes a population eligible for immune checkpoint inhibitors (Section 3.3.3).

**Table 4.**
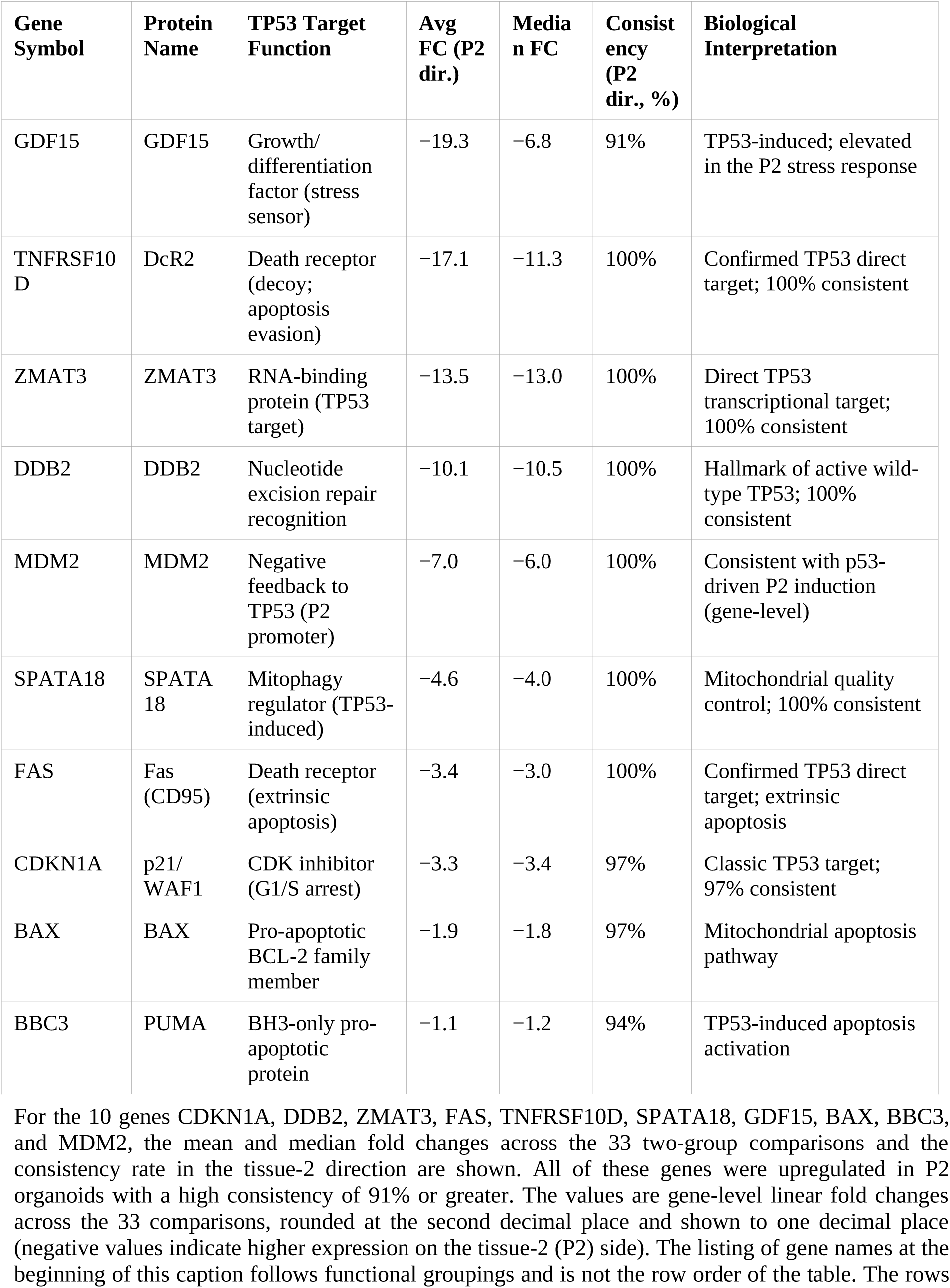
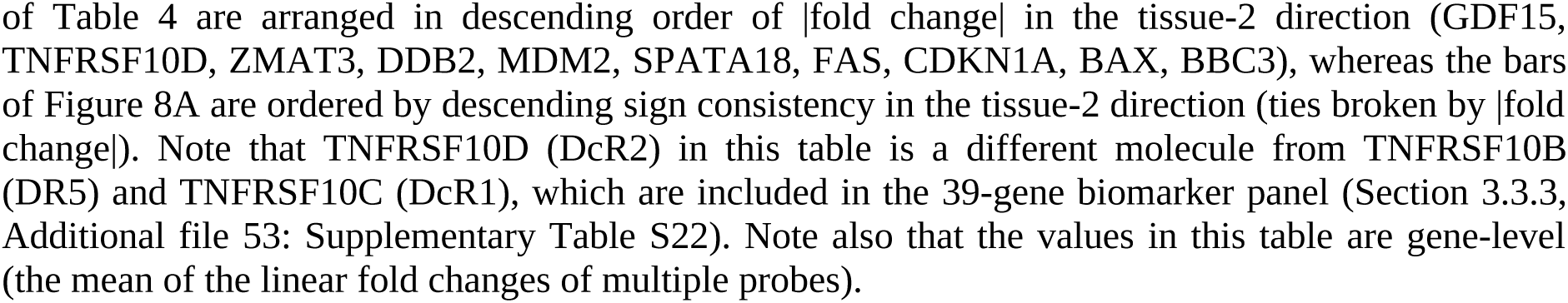
Wild-type TP53 pathway activation signature (10 p53 target genes in P2 organoids).

| Gene Symbol | Protein Name | TP53 Target Function | Avg FC (P2 dir.) | Median FC | Consistency (P2 dir., %) | Biological Interpretation |
| --- | --- | --- | --- | --- | --- | --- |
| GDF15 | GDF15 | Growth/ differentiation factor (stress sensor) | -19.3 | -6.8 | 91% | TP53-induced; elevated in the P2 stress response |
| TNFRSF10D | DcR2 | Death receptor (decoy; apoptosis evasion) | -17.1 | -11.3 | 100% | Confirmed TP53 direct target; 100% consistent |
| ZMAT3 | ZMAT3 | RNA-binding protein (TP53 target) | -13.5 | -13.0 | 100% | Direct TP53 transcriptional target; 100% consistent |
| DDB2 | DDB2 | Nucleotide excision repair recognition | -10.1 | -10.5 | 100% | Hallmark of active wild-type TP53; 100% consistent |
| MDM2 | MDM2 | Negative feedback to TP53 (P2 promoter) | -7.0 | -6.0 | 100% | Consistent with p53-driven P2 induction (gene-level) |
| SPATA18 | SPATA18 | Mitophagy regulator (TP53-induced) | -4.6 | -4.0 | 100% | Mitochondrial quality control; 100% consistent |
| FAS | Fas (CD95) | Death receptor (extrinsic apoptosis) | -3.4 | -3.0 | 100% | Confirmed TP53 direct target; extrinsic apoptosis |
| CDKN1A | p21/ WAF1 | CDK inhibitor (G1/S arrest) | -3.3 | -3.4 | 97% | Classic TP53 target; 97% consistent |
| BAX | BAX | Pro-apoptotic BCL-2 family member | -1.9 | -1.8 | 97% | Mitochondrial apoptosis pathway |
| BBC3 | PUMA | BH3-only pro-apoptotic protein | -1.1 | -1.2 | 94% | TP53-induced apoptosis activation |

For the 10 genes CDKN1A, DDB2, ZMAT3, FAS, TNFRSF10D, SPATA18, GDF15, BAX, BBC3, and MDM2, the mean and median fold changes across the 33 two-group comparisons and the consistency rate in the tissue-2 direction are shown. All of these genes were upregulated in P2 organoids with a high consistency of 91% or greater. The values are gene-level linear fold changes across the 33 comparisons, rounded at the second decimal place and shown to one decimal place (negative values indicate higher expression on the tissue-2 (P2) side). The listing of gene names at the beginning of this caption follows functional groupings and is not the row order of the table. The rows

Several of the top tissue-2 DEGs were C/D box small nucleolar RNAs (snoRNAs) of the imprinted SNORD116 cluster at 15q11-q13 (for example, SNORD116-17/-19, average fold change −662.4; probe-level values in Additional file 36: Supplementary Table S16). As a continuous score, the cluster followed neither P2 usage nor, consistently, TP53 status (Section 3.1.4). It is analyzed in the companion study on the 15q11-q13 locus (T. Tsukui, R. Yao, and K. Tsuda, unpublished observations) rather than interpreted here.

The tissue-2 pathway profile was one of inflammation, invasion, and adaptation to hypoxia, diametrically opposite to that of tissue 1. The top pathway was Pathogen Induced Cytokine Storm Signaling (sign consistency 100%, median z = −3.32). In the Upstream Analysis, the tissue-2 activators were led by NUPR1 (median z = −8.59, 94%) and wild-type TP53 (−7.14, 100%). Wild-type TP53 thus ranked in the top class of activated upstream regulators. EGR1, a transcription factor that can regulate MDM2 in a context-dependent manner [100], pointed the same way but fell below the threshold (−1.95). Together, these rankings are consistent with transcriptional induction of the MDM2 P2 promoter (Section 3.1.2; Figure 1). Full pathway results are in Additional file 14 (Supplementary Note Sections 2–4), and the source tables in Additional files 37–41 (Supplementary Tables S6–S9 and S31).

#### 3.3.2 Signature behavior in clinical cohorts: CMS, mismatch-repair status, and prognosis

We externally validated the organoid signatures in two clinical cohorts: TCGA-COAD/READ (n = 624 primary tumors) and GSE39582 (519 tumors with mismatch-repair annotation; 75 deficient [dMMR] and 444 proficient [pMMR]). We scored the P1/P2 signatures by gene set variation analysis (GSVA) [78], which gives each sample an enrichment score for a gene set. Among the 563 CMS-classified specimens, both scores differed significantly by CMS class (Kruskal–Wallis; P1 P = 7.2×10⁻¹⁵; P2 immune-excluded score P = 2.54×10⁻3⁰, highest in CMS3; full 29-gene P2 score P = 3.3×10⁻Z⁸, with CMS1 and CMS3 comparable). The class with the highest value depended on the functional module. The P1 differentiation core was highest in CMS2, whereas the whole P1 score leaned toward the highly proliferative CMS1. The P2 metaplasia-plus-p53 core was highest in CMS3, whereas the immune module was highest in CMS1 (module breakdown in Additional file 42: Supplementary Table S34; Additional file 43: Supplementary Figure S6). Single-sample gene set enrichment analysis (ssGSEA) [101] gave concordant rankings for all 11 scored modules. Thus, signatures derived from the organoids of 21 patients correspond to clinical CMS subtypes in a module-specific manner.

In GSE39582, the immune-excluded P2 score was significantly higher in dMMR tumors (Wilcoxon, P = 6.59×10⁻¹⁰; full 29-gene score P = 1.14×10⁻¹⁷), whereas the P1 score was higher in pMMR tumors (P = 0.017). The higher P2 score in dMMR tumors is consistent with an MSI-associated expression profile of the P2 signature. It does not extend to P2 promoter usage, which was not associated with MSI status (Sections 3.2.2 and 3.1.4).

In the TCGA-COAD/READ survival cohort (n = 591), Kaplan–Meier analysis showed a non-significant trend toward longer overall survival in the P1-high group (log-rank P = 0.114). The P2 score showed no difference (P = 0.104). By contrast, survival differed significantly among CMS subtypes (P = 0.0072), consistent with their reported prognostic differences [2] (Additional file 44: Supplementary Figure S7). After adjustment for age, stage, and MSI, the P1 score made no independent contribution (likelihood-ratio test P = 0.65; C-index = 0.785; n = 214). The association was attributable mainly to stage. The full analysis is given in the Supplementary Report (Additional file 1), Section R-10.

The association between the signatures and CMS was not an artifact of shared genes. They shared at most 3 genes per class and side with the CMScaller templates (reference gene sets that assign CMS classes), and every scored functional submodule was significantly associated with CMS on its own (Additional file 45: Supplementary Figure S9). The P2 score, and especially its p53-target module, was strongly associated with TP53 status in TCGA (P = 1.8×10⁻⁴¹ and 5.4×10⁻⁴⁷; module median +0.48 in wild type versus −0.41 in mutant). This was expected from its p53-target content and is a positive control rather than independent validation. The full analysis is given in the Supplementary Report (Additional file 1), Section R-11.

#### 3.3.3 Biomarker panel, immune microenvironment, and exploratory analyses

Seven immune checkpoint molecules (inhibitory regulators of immune responses) were consistently higher in direction on the tissue-2 side (TIGIT, CTLA4, PDCD1, IDO1, LAG3, IDO2, HAVCR2; in 67% to 85% of the comparisons). However, the shifts were small (median fold change 1.0 to 1.1), and all fell below the DEG effect-size criterion (Figure 8B; Additional file 46: Supplementary Table S30). Organoid cultures contain few if any immune cells. These small shifts therefore presumably reflect expression by the tumor cells, and the finding is exploratory. Six of the seven were adopted as the immune module of the P2 signature. The study does not propose the P2-dominant group as a population indicated for immune checkpoint inhibitors (see the microenvironment analysis below and Section 4.5).

We also ran the IPA Regulator Effects, Networks, and Graphical Summary analyses. The highest-scoring cascade (Consistency Score 25.93) ran from upstream regulators such as EGR1, BMP4, and HIF1A through TP53 and VEGFA. MDM2 coexisted in a protein–protein interaction module with FBXW7, CSNK2A1/2, and XPO1. These results contrasted the autonomously proliferative hub structure of tissue 1 with the antagonism in tissue 2 between p53-induced apoptosis and MDM2/AKT survival. Full pathway results are in Additional file 14 (Supplementary Note Sections 2–4). The source tables and the Graphical Summary are in Additional files 47 and 48 (Supplementary Tables S32 and S19), Additional file 49 (Supplementary Figure S14) and Additional files 50 and 51 (Supplementary Tables S20 and S21).

Using the 39 genes identified by IPA Biomarker Detection, hierarchical clustering (Manhattan distance, average linkage) divided the 63 samples into two clusters of 51 and 12 specimens (final merge height 57.93 versus 48.66; cophenetic correlation 0.833). The clusters corresponded to MDM2 promoter usage in 62 of 63 specimens (98.4%) (Additional file 52: Supplementary Figure S5). The 39 genes fall into seven post hoc functional categories. These include direct wild-type TP53 targets, invasion/extracellular-matrix (ECM) genes (MSLN, MMP7, SERPINB5, COL17A1, DCBLD2; MMP7 and DCBLD2, for example, were higher in tissue 2 in all 33 comparisons, by a median of 9.1- and 11.6-fold) [102] and gastric metaplasia markers (CTSE, REN) (gene annotations in Additional file 53: Supplementary Table S22; key items of the biomarker and GO analyses in Additional file 54: Supplementary Table S43). The panel was selected from the same P1-versus-P2 comparisons. This 62/63 agreement is therefore a within-sample (non-independent) result. It summarizes the p53-active/metaplastic contrast of these specimens rather than providing an independent classifier. Any assay derived from it would require independent and prospective validation, and the analysis is exploratory in a 21-patient cohort (Section 4.6). Full details are given in the Supplementary Report (Additional file 1), Section R-7.

In TCGA-COAD/READ, comparing the top and bottom P2-score tertiles, NK cells, the cytotoxicity score, B cells, and T cells (MCP-counter) and CD8-positive T cells (quanTIseq) were significantly higher in the P2-high group (P = 1.5×10⁻ZZ to 2.3×10⁻¹⁰; both methods estimate immune-cell abundance from bulk expression). These findings remained after adjustment for MSI (Additional files 55 and 56: Supplementary Tables S35 and S36). Immune infiltration in P2 is therefore not a restatement of MSI status. By contrast, TIDE-predicted response to immune checkpoint inhibitors (ICI) was not reproduced in the MSI-annotated specimens alone (P = 0.50) and is treated as exploratory. The full analysis is given in the Supplementary Report (Additional file 1), Section R-14. In one-sided Fisher’s exact tests against two independent reference sets, the tissue-1-side DEGs were enriched specifically in CMS2 markers in the TCGA-derived reference set B (OR = 3.92, BH-FDR = 8.7×10⁻⁷), but in the CMScaller reference set A they were enriched in CMS1, CMS2 and CMS3 markers to a similar degree (OR = 3.09–3.77). Tissue-2-side DEGs were enriched most strongly in CMS3 markers in reference set B (OR = 5.91, BH-FDR = 2.0×10⁻¹¹) while also carrying CMS1 markers (Additional file 15: Supplementary Table S28). Removing the P2 signature genes and a metaplasia/intestinal marker panel (47 genes) from the query hardly attenuated the CMS3 enrichment (OR = 5.91 → 5.30). The enrichment is therefore not an artifact of the signature definitions (Section 4.2.1). In the exploratory colibactin analysis, activities of the mutational signatures SBS88 and ID18 were only slightly denser on the P2-dominant side, with partial replication in TCGA-COAD/READ (values in Additional file 12; Section 4.6). This shift did not run through p53 activity: in TCGA-COAD/READ the activities were unrelated to the p53 activity score in TP53 wild-type tumors (132 tumors, Spearman ρ = −0.059 for SBS88 and −0.010 for ID18; 91 non-hypermutated wild-type tumors, −0.088 and −0.001), the positivity rate did not differ between wild-type tumors with and without evidence of p53 activity (3 of 57 versus 6 of 75 for SBS88, P = 0.73; 7 of 57 versus 6 of 75 for ID18, P = 0.56), and in the Nunes cohort the weak positive association with the p53-target score seen in MSS specimens did not survive adjustment for MSI status, stage, site and age (Additional file 7: Supplementary Table S48, sheet P).

## 4. Discussion

### 4.1 TP53 status, morphology and MDM2 isoforms: the central finding of this study

In the discovery organoids (21 patients, 63 specimens), morphology corresponded to MDM2 promoter usage specimen by specimen. However, TP53 status coincided with promoter usage (40 of the 42 mutant specimens were P1-dominant). When the two were compared, morphology followed TP53 status to a degree similar to promoter usage. Three of the four morphology-deviating specimens followed TP53 status rather than promoter usage (Figure 9C), and, within the narrow range in which the two axes disagreed (five specimens), P2 dominance added no detectable contribution to morphology after accounting for TP53 status (Sections 3.1.3 and 3.1.4). Like the morphology–isoform association, the TP53–morphology association in this cohort was established at the specimen level (patient-level sensitivity analysis in Section 3.1.4). In an independent biobank (GSE294511 [36]; 88 organoids with published TP53 status), where the association held on patient means, and in TCGA tumors, the morphology genes again followed TP53 status after accounting for 20q (the long arm of chromosome 20). In that biobank, the alignment of morphology with P2 was largely explained by TP53 status (Section 3.1.5). We therefore interpret the central finding as an alignment of morphology with TP53 status. The comparison of morphology against TP53 status and promoter usage was added after the per-specimen TP53 calls had been finalized and is therefore post hoc; the independent biobank (GSE294511) was analyzed under a plan fixed in advance, whose primary test concerned the P2 hypothesis (Section 3.1.5). TP53-mutant organoids tend to be compact and glandular, with expression features of chromosomal instability (frequent gains and losses of chromosome segments), and TP53 wild-type organoids tend to be round (cystic–mucinous). Gastric-metaplastic genes followed TP53 status in the independent biobank and in tumors. MDM2 P2 usage, which tracks p53 activity in organoids (the evidence that it does so beyond genotype resting mainly on the independent biobank and two TP53-mutant P2-dominant specimens; Sections 3.1.4, 3.2.1 and 3.2.5), reflects the p53 state rather than determining morphology. Within TCGA wild-type tumors, the morphology gene score was unrelated to the p53 activity score (ρ = −0.08, P = 0.37; 119 tumors; Additional file 7: Supplementary Table S48, sheet H2). Morphology therefore follows TP53 genotype, or the pathway it marks, rather than p53 function. At the proximal level, the genes higher on the cystic–mucinous side were enriched for cytoskeleton and adhesion terms (VCL, AHNAK, TRIOBP, FYN, CDC42BPB; partial ρ with the non-Type1 fraction 0.48 to 0.58; Section 3.1.4). This places the shape difference in a differentiation and cytoskeletal–adhesion program rather than an acute p53 response. At the distal level, two routes remain open. In the first, loss of wild-type p53 permits the chromosomal-instability route, with which the morphology genes align at the transcript-cluster level (20q). In the second, mutant p53 acquires a gain of function; the mouse intestinal-organoid comparison discussed in Section 4.4 favors this route. Comparing TP53-knockout with mutant-knock-in organoids of the same genetic background would separate the two. Whether TP53 loss itself drives the morphological change was not tested, and TP53 mutation is entangled with chromosomal instability. Nevertheless, adjustment for 20q, aneuploidy and MSI (and, in tumors, BRAF mutation and CIMP) did not remove the association. Among non-MSI, BRAF wild-type, CIMP-low tumors, however, the 20q-adjusted morphology difference was only marginal (Section R-26). Because the observation rests on molecular axes independent of the morphological typing of Okamoto et al. [12], it complements that work rather than reproducing it (Section 4.4). The full discussion is given in the Supplementary Report (Additional file 1), Section R-17.

**Figure 9.**
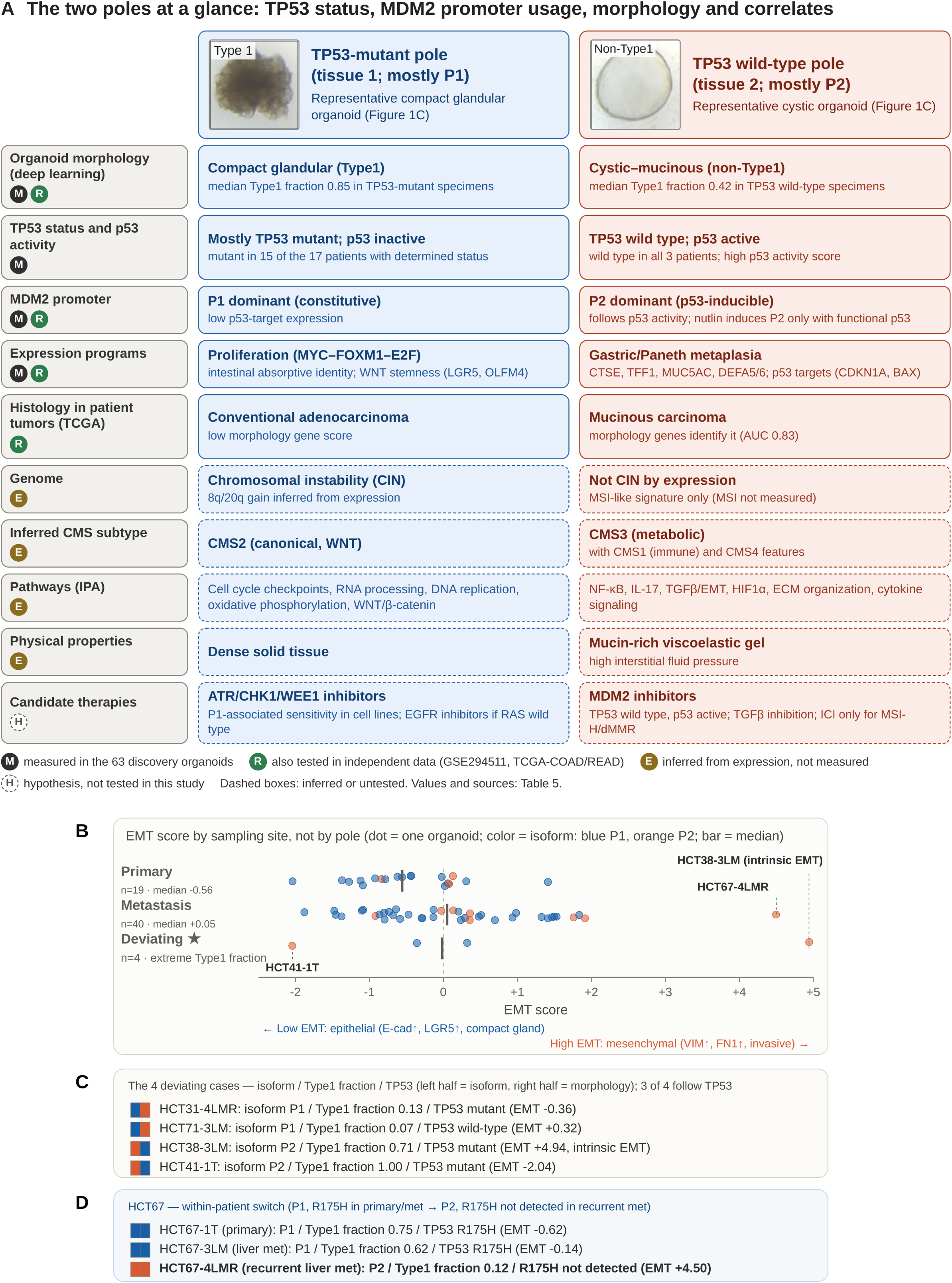
The two poles at a glance: TP53 status, MDM2 promoter usage, morphology and correlates. (A) Comparison of the TP53-mutant pole (tissue 1, mostly P1) and the TP53 wild-type pole (tissue 2, mostly P2), headed by the representative organoids of Figure 1C. Each row is tagged by the strength of its evidence: M, measured in the 63 discovery organoids; R, also tested in independent data (GSE294511 or TCGA-COAD/READ); E, inferred from expression and not measured; H, hypothesis not tested in this study (E and H rows in dashed boxes). The poles are tendencies. Morphology follows TP53 status (as do the metaplastic genes in the independent biobank and in tumors), the p53-target genes follow P2 usage in organoids (Section 3.1.4), and morphology co-varies with the other layers only partly (Section 4.2.1). Values and sources are in Table 5. (B) EMT score distribution of 59 specimens, with the 4 specimens deviating from the typical pattern of their group (Section 3.1.2) shown separately. Specimens are grouped by sampling site, the axis the score follows (primary n = 19, median −0.56; metastasis n = 40, median +0.05; P = 0.014 over all 63 specimens), and colored by MDM2 isoform (blue P1, orange P2). In all 63 specimens the P2–P1 difference was not significant (P = 0.079), and sampling site was balanced between isoform groups (Section 3.1.2). (C) The 4 deviating cases with their TP53 status: in morphology, three of the four follow TP53 status rather than promoter usage. (D) The within-patient P1→P2 switch in HCT67 (EMT score −0.62 → −0.14 → +4.50); R175H was detected in the primary tumor and the liver metastasis but not in the recurrent liver metastasis.

**Table 5.**
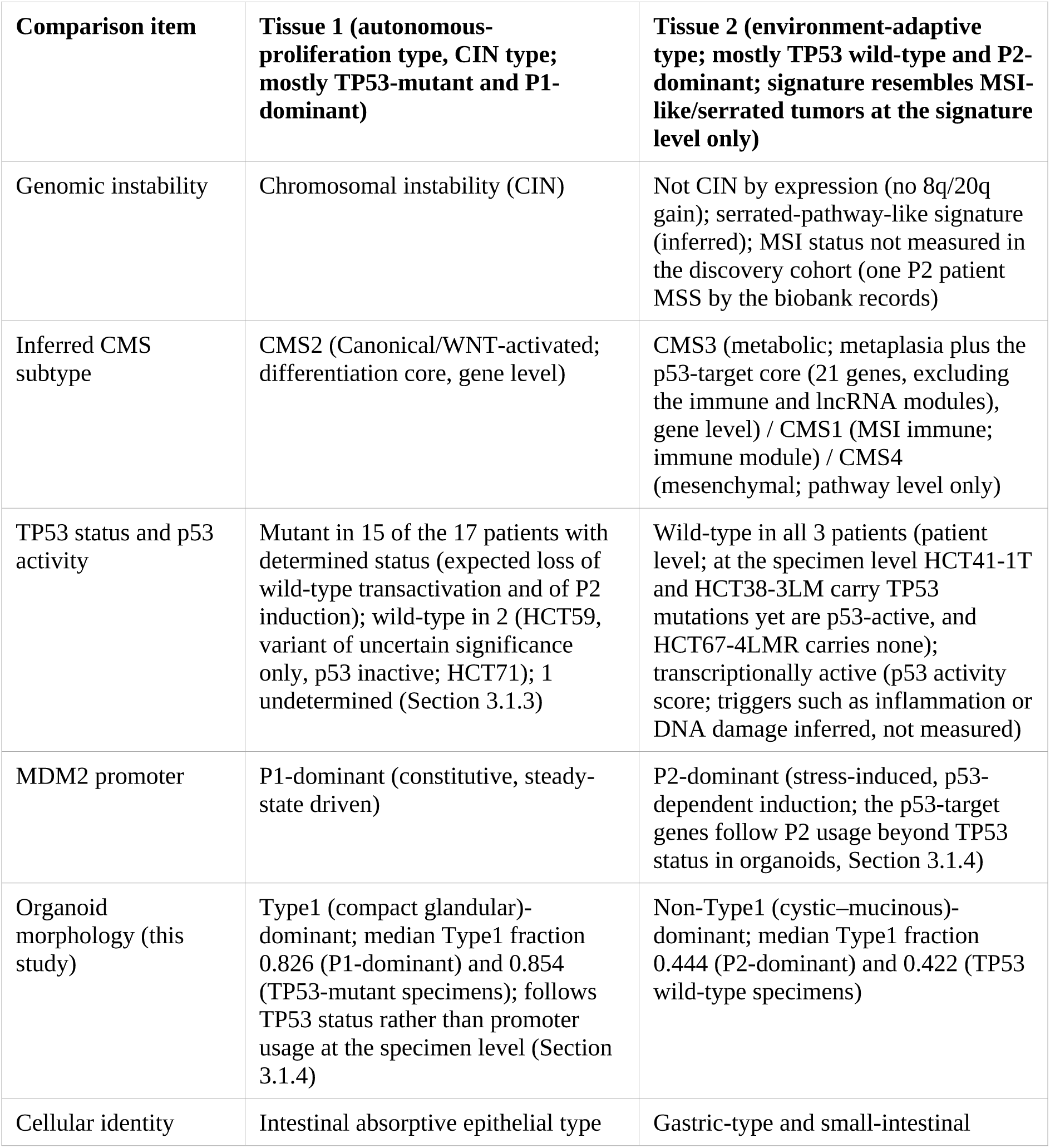
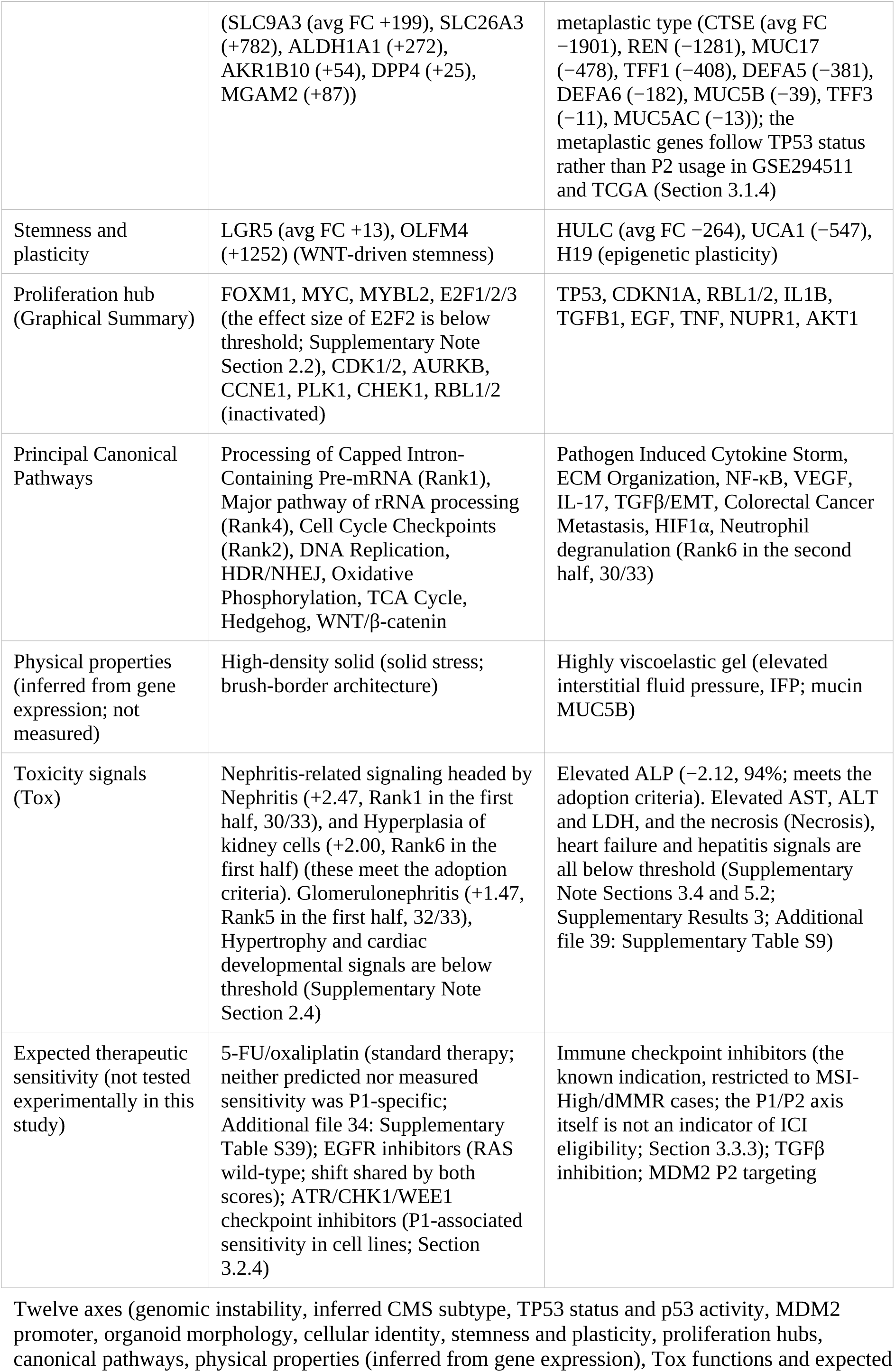
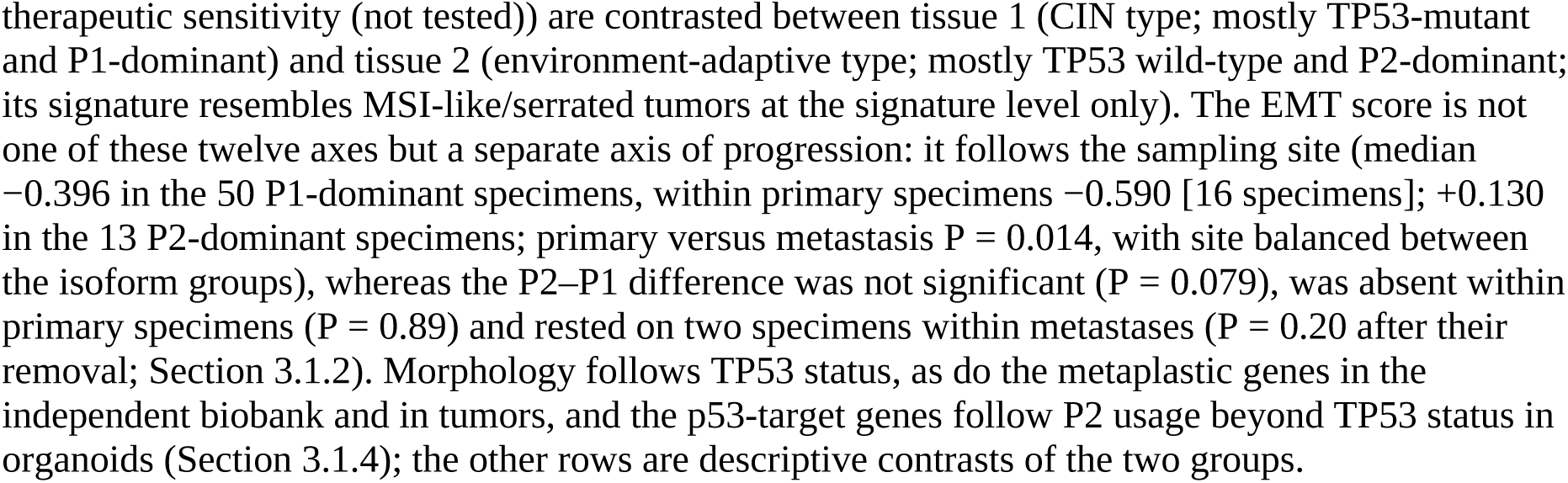
Multilayered comparison of the two colorectal cancer tissues.

| **Comparison item** | **Tissue 1 (autonomous-proliferation type, CIN type; mostly TP53-mutant and P1-dominant)** | **Tissue 2 (environment-adaptive type; mostly TP53 wild-type and P2-dominant; signature resembles MSI-like/serrated tumors at the signature level only)** |
| --- | --- | --- |
| Genomic instability | Chromosomal instability (CIN) | Not CIN by expression (no 8q/20q gain); serrated-pathway-like signature (inferred); MSI status not measured in the discovery cohort (one P2 patient MSS by the biobank records) |
| Inferred CMS subtype | CMS2 (Canonical/WNT-activated; differentiation core, gene level) | CMS3 (metabolic; metaplasia plus the p53-target core (21 genes, excluding the immune and lncRNA modules), gene level) / CMS1 (MSI immune; immune module) / CMS4 (mesenchymal; pathway level only) |
| TP53 status and p53 activity | Mutant in 15 of the 17 patients with determined status (expected loss of wild-type transactivation and of P2 induction); wild-type in 2 (HCT59, variant of uncertain significance only, p53 inactive; HCT71); 1 undetermined (Section 3.1.3) | Wild-type in all 3 patients (patient level; at the specimen level HCT41-1T and HCT38-3LM carry TP53 mutations yet are p53-active, and HCT67-4LMR carries none); transcriptionally active (p53 activity score; triggers such as inflammation or DNA damage inferred, not measured) |
| MDM2 promoter | P1-dominant (constitutive, steady-state driven) | P2-dominant (stress-induced, p53-dependent induction; the p53-target genes follow P2 usage beyond TP53 status in organoids, Section 3.1.4) |
| Organoid morphology (this study) | Type1 (compact glandular)-dominant; median Type1 fraction 0.826 (P1-dominant) and 0.854 (TP53-mutant specimens); follows TP53 status rather than promoter usage at the specimen level (Section 3.1.4) | Non-Type1 (cystic-mucinous)-dominant; median Type1 fraction 0.444 (P2-dominant) and 0.422 (TP53 wild-type specimens) |
| Cellular identity | Intestinal absorptive epithelial type | Gastric-type and small-intestinal |
|  | (SLC9A3 (avg FC +199), SLC26A3 (+782), ALDH1A1 (+272), AKR1B10 (+54), DPP4 (+25), MGAM2 (+87)) | metaplastic type (CTSE (avg FC -1901), REN (-1281), MUC17 (-478), TFF1 (-408), DEFA5 (-381), DEFA6 (-182), MUC5B (-39), TFF3 (-11), MUC5AC (-13)); the metaplastic genes follow TP53 status rather than P2 usage in GSE294511 and TCGA (Section 3.1.4) |
| Stemness and plasticity | LGR5 (avg FC +13), OLFM4 (+1252) (WNT-driven stemness) | HULC (avg FC -264), UCA1 (-547), H19 (epigenetic plasticity) |
| Proliferation hub (Graphical Summary) | FOXM1, MYC, MYBL2, E2F1/2/3 (the effect size of E2F2 is below threshold; Supplementary Note Section 2.2), CDK1/2, AURKB, CCNE1, PLK1, CHEK1, RBL1/2 (inactivated) | TP53, CDKN1A, RBL1/2, IL1B, TGFB1, EGF, TNF, NUPR1, AKT1 |
| Principal Canonical Pathways | Processing of Capped Intron-Containing Pre-mRNA (Rank1), Major pathway of rRNA processing (Rank4), Cell Cycle Checkpoints (Rank2), DNA Replication, HDR/NHEJ, Oxidative Phosphorylation, TCA Cycle, Hedgehog, WNT/β-catenin | Pathogen Induced Cytokine Storm, ECM Organization, NF-κB, VEGF, IL-17, TGFβ/EMT, Colorectal Cancer Metastasis, HIF1α, Neutrophil degranulation (Rank6 in the second half, 30/33) |
| Physical properties (inferred from gene expression; not measured) | High-density solid (solid stress; brush-border architecture) | Highly viscoelastic gel (elevated interstitial fluid pressure, IFP; mucin MUC5B) |
| Toxicity signals (Tox) | Nephritis-related signaling headed by Nephritis (+2.47, Rank1 in the first half, 30/33), and Hyperplasia of kidney cells (+2.00, Rank6 in the first half) (these meet the adoption criteria). Glomerulonephritis (+1.47, Rank5 in the first half, 32/33), Hypertrophy and cardiac developmental signals are below threshold (Supplementary Note Section 2.4) | Elevated ALP (-2.12, 94%; meets the adoption criteria). Elevated AST, ALT and LDH, and the necrosis (Necrosis), heart failure and hepatitis signals are all below threshold (Supplementary Note Sections 3.4 and 5.2; Supplementary Results 3; Additional file 39: Supplementary Table S9) |
| Expected therapeutic sensitivity (not tested experimentally in this study) | 5-FU/oxaliplatin (standard therapy; neither predicted nor measured sensitivity was P1-specific; Additional file 34: Supplementary Table S39); EGFR inhibitors (RAS wild-type; shift shared by both scores); ATR/CHK1/WEE1 checkpoint inhibitors (P1-associated sensitivity in cell lines; Section 3.2.4) | Immune checkpoint inhibitors (the known indication, restricted to MSI-High/dMMR cases; the P1/P2 axis itself is not an indicator of ICI eligibility; Section 3.3.3); TGFβ inhibition; MDM2 P2 targeting |

Twelve axes (genomic instability, inferred CMS subtype, TP53 status and p53 activity, MDM2 promoter, organoid morphology, cellular identity, stemness and plasticity, proliferation hubs, canonical pathways, physical properties (inferred from gene expression), Tox functions and expected
therapeutic sensitivity (not tested)) are contrasted between tissue 1 (CIN type; mostly TP53-mutant and P1-dominant) and tissue 2 (environment-adaptive type; mostly TP53 wild-type and P2-dominant; its signature resembles MSI-like/serrated tumors at the signature level only). The EMT score is not one of these twelve axes but a separate axis of progression: it follows the sampling site (median $-0.396$ in the 50 P1-dominant specimens, within primary specimens $-0.590$ [16 specimens]; $+0.130$ in the 13 P2-dominant specimens; primary versus metastasis $P = 0.014$ , with site balanced between the isoform groups), whereas the P2–P1 difference was not significant ( $P = 0.079$ ), was absent within primary specimens ( $P = 0.89$ ) and rested on two specimens within metastases ( $P = 0.20$ after their removal; Section 3.1.2). Morphology follows TP53 status, as do the metaplastic genes in the independent biobank and in tumors, and the p53-target genes follow P2 usage beyond TP53 status in organoids (Section 3.1.4); the other rows are descriptive contrasts of the two groups.

### 4.2 MDM2 P2 usage as a readout of p53 activity: a working model

Our data support the view that the change in MDM2 isoforms is a consequence and a readout (a measurable indicator) of p53 activity (Sections 3.2.1 and 3.1.4). We did not test whether promoter usage itself feeds back on cellular programs. In tissue 1, TP53 mutation or loss accompanies the chromosomal-instability (CIN) pathway [103,104]. This would abolish p53-dependent P2 activation and leave MDM2 transcribed mainly from P1. Here the MYC/FOXM1/E2F proliferative program and the glandular architecture accompany TP53 status and chromosomal instability rather than promoter usage. Public nascent-transcription data qualify this picture. In ENCODE PRO-cap, P2 accounted for about half of MDM2 initiation in normal colon and stomach, and for a larger share of initiation than of mRNA in p53-deficient Caco-2 cells (Section 3.2.1). Basal initiation at P2 therefore does not appear to require p53. Beyond inducing P2, p53 may also turn P2 initiation into productive, mRNA-yielding transcription. Because PRO-cap also captures paused and unstable transcripts, and only normal tissue and one p53-deficient line could be examined, both inferences remain hypotheses. In tissue 2, wild-type p53 is active and the P2 promoter is induced. Expression and IPA predictions suggest chronic inflammation, DNA damage and oxidative stress as possible triggers (not measured). This state co-occurs with suppression of p53-induced apoptosis and with a lineage conversion — a TGFβ/SMAD3-mediated partial EMT [105] and a gastric-type program that follows TP53 status — whose temporal order relative to P2 induction is not established (Section 4.3). TP53 status is consistent with the model. All three P2 patients (HCT27, HCT33, HCT64) were TP53 wild-type at the patient level (at the specimen level the P2 specimens HCT41-1T and HCT38-3LM carried TP53 mutations yet were p53-active, whereas HCT67-4LMR carried none), whereas TP53 mutations were found in 15 of the 17 P1-dominant patients with determined status (Section 3.1.3). Because genotype and promoter usage so largely coincide in this cohort, the organoids alone cannot show whether P1 dominance follows TP53 loss specifically or p53 inactivity more generally (Section 3.2.1). The coexistence of P1- and P2-dominant specimens within patients (HCT38, HCT41, HCT67; Figures 3C and 9D) is interpreted as a condition-dependent state. Because this study is cross-sectional, the direction and reversibility of any transition were not tested (Section 4.6). The Rank 2 Regulator Effects cascade further suggests a bidirectional relationship in which MDM2 is both an upstream regulator of p53 and a downstream target of mitotic-checkpoint factors (Supplementary Report, Section R-19; upstream and downstream factors contrasted in Additional file 57: Supplementary Table S44). The full discussion is given in the Supplementary Report (Additional file 1), Section R-18.

#### 4.2.1 A unified model of the four observational layers (Figure 9)

The four observational layers — TP53 mutation status, MDM2 promoter choice, the downstream programs, and organoid morphology — tend to vary together (Figure 9A; the two tissue types are contrasted in Table 5), and the data place TP53 status at their center. Per-specimen support is collected in Figure 10. For each of the 63 discovery specimens, the 119 organoid lines of the independent biobank and the 380 TCGA-COAD/READ tumors, it shows the morphology or sampling site, the MDM2 promoter measure, TP53 status and the p53 activity score as one annotated column per sample. The measure differs by cohort, so this is an overview rather than a cross-cohort replication; every drawn value is listed in Additional file 7: Supplementary Table S48, sheet M. At the TP53-mutant pole (tissue 1, P1), p53 inactivity (CIN; differentiation core CMS2) accompanies basal P1 transcription, maximal MYC–FOXM1–E2F signaling, retained epithelial character and a compact glandular morphology. At the TP53 wild-type pole (tissue 2, P2), active p53 accompanies activation of p53-inducible P2, gastric/Paneth metaplasia and a mucinous, cystic morphology; this pole is a composite subtype centered on CMS3 at the gene level, with CMS1-like immune features and CMS4-like features only at the pathway level. The EMT score is not one of these layers but a separate axis of progression. It follows the sampling site (higher in metastases, P = 0.014) rather than the two poles (P2–P1 difference P = 0.079; Section 3.1.2; Figure 9B). Within this pattern the layers separate into two readouts (Section 3.1.4). Morphology followed TP53 status beyond 20q, aneuploidy and MSI (and, in tumors, BRAF mutation and CIMP). The gastric-metaplastic genes followed TP53 status in the independent biobank and in tumors. Neither followed P2 usage, whereas the p53-target module followed P2 usage beyond TP53 status in organoids (although without circularity this was confirmed only in the independent biobank: pre-specified p53 activity score, P = 0.012; discovery P = 0.094; Sections 3.1.4 and 3.2.5). The results of Section 3.2 make the meaning of P2 concrete. P2-dominant organoid specimens were p53-active, including two that carried TP53 mutations. P2 was induced more strongly than P1 by nutlin, and not at all without p53. In non-hypermutated, MC3-confirmed wild-type tumors P2_index tracked p53 activity, though not beyond total MDM2 (Section 3.2.3). The P2 signature resembles the MSI-like/serrated program at the signature level, but P2 dominance is not equivalent to microsatellite instability: P2_index did not differ among the CIN, GS and MSI subtypes and was not associated with MSI status (Sections 3.2.2, 3.1.4 and 4.6). A detailed interpretation is provided in Supplementary Note Section 5.

**Figure 10.**
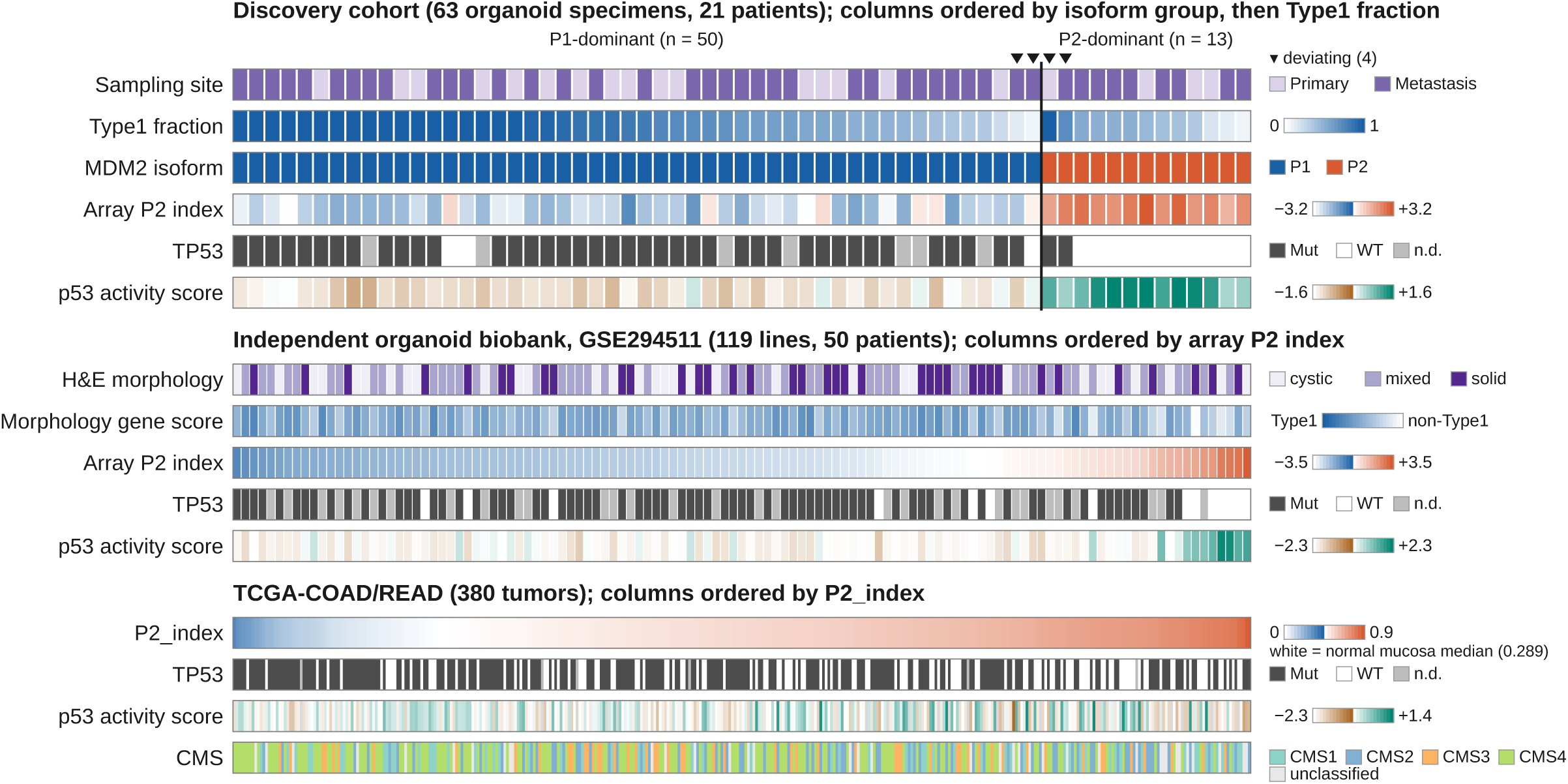
Per-specimen overview: morphology, MDM2 promoter usage, TP53 and p53 activity in the three cohorts. Each column is one sample and each row one annotation; color scales and category keys are at the right of each block. Top: the 63 discovery organoid specimens from 21 patients (Sections 3.1.2 and 3.2.1). They are grouped by MDM2 isoform call (P1-dominant n = 50, P2-dominant n = 13; vertical divider) and, within each group, ordered by descending Type1 fraction. The rows are sampling site, Type1 fraction, isoform call, the array MDM2 P2 index (exon 2 − exon 1, RMA; Section 2), TP53 status and the p53 activity score (Methods); ▾ marks the 4 specimens deviating from the typical pattern of their group (Section 3.1.2). Middle: the 119 organoid lines from 50 patients of the independent biobank GSE294511 (Sections 3.1.5 and 3.2.5), ordered by the array MDM2 P2 index. The rows are H&E morphology (solid, mixed, cystic), the morphology gene score (+ = non-Type1 side), the array P2 index, TP53 status and the p53 activity score (Supplementary Table S48, sheet F). Bottom: the 380 TCGA-COAD/READ tumors of Supplementary Table S47 (Additional file 6), ordered by P2_index. The rows are P2_index (color scale anchored with white at the normal-mucosa median of 0.289; Supplementary Table S48, sheet I), TP53 status, the p53 activity score (gray, the 18 tumors without a score) and the consensus molecular subtype (CMS). Mut, mutant; WT, wild type; n.d., not determined (in TCGA, tumors without somatic mutation data). Every drawn value is listed, with the source table of each block, in Additional file 7: Supplementary Table S48, sheet M. The drawing script, the drawn-value file and a cell-by-cell verification that recomputes every drawn color from the sources are deposited in Additional file 2.

### 4.3 Lineage plasticity in the TP53 wild-type, P2-dominant specimens

An exploratory observation of this study is that, in the TP53 wild-type, P2-dominant specimens, attenuated intestinal identity and acquired gastric metaplasia co-occur with a partial EMT program. Whether these are causally linked cannot be tested with this design. We offer two working hypotheses. First, a decrease of the nuclear chromatin repression machinery (B-type lamins, H3K9 writers, PRC2; Additional file 58: Supplementary Table S29) is consistent with reduced lamina tension permitting derepression of lineage loci [106–108]. Second, the TGFβ/SMAD3-driven partial EMT during the survival reprieve [109–112] is proposed to be held short of complete EMT by miR-200-mediated repression of ZEB1/2 [113,114]. More generally, lineage plasticity is a common pathway to therapeutic resistance [115]. In the injured mouse colon, wild-type p53 terminates the damage-induced regenerative, fetal-like state, and p53-deficient crypts remain locked in it [116]; whether the gastric-type program of the p53-active, P2-dominant specimens relates to such injury-associated states is not established. Each causal relationship requires functional validation (Section 4.6); the full discussion is given in the Supplementary Report (Additional file 1), Section R-20.

### 4.4 Comparison with previous studies

Several earlier studies bear on these findings. van de Wetering et al. [117] described diverse organoid morphologies without linking them to molecular subtypes. Fujii et al. [118] and Betge et al. [119] related CRC organoid phenotypes to niche-factor requirements and to drug responses. Zhao et al. [10] and Lukonin et al. [11] linked morphology to cell state in other systems. A CRC organoid study identified cystic and solid subtypes by image-based profiling and proposed the cystic, stem-like relapse phenotype as a candidate prognostic biomarker [120]. This study extends these frameworks to colorectal cancer. It is novel in linking morphology to TP53 status, to a transcript-level readout of p53 activity (MDM2 P1/P2) and to therapeutic stratification (detailed comparison in Additional file 59: Supplementary Table S33; full discussion in Section R-21). Morphological typing (Okamoto 2022 [12]), cellular composition (Okamoto 2021 [24]) and the alignment of morphology with TP53 status together with a promoter-level p53 readout (this study) are three independent, complementary analytical axes built on the same biobank. The original typing study reported no correlation of PDO morphology with the APC/TP53/KRAS profile. In our specimens, by contrast, TP53-mutant organoids had a higher Type1 fraction, beyond 20q and aneuploidy, and in an independent biobank the morphology gene score followed TP53 status (Sections 3.1.3, 3.1.4 and 3.1.5). The difference may reflect our two-class summary (Type1 versus non-Type1) and the continuous per-specimen comparison. In mouse Apc-mutant intestinal tumor organoids, which were round and cystic when Trp53 was wild-type or deleted, homozygous mutant p53 (R270H) induced complex glandular structures associated with invasiveness; and in nine human colorectal cancers, tumors with TP53 mutations around codon 273 had higher gland-branching scores than wild-type tumors [121]. Our human organoid data agree in direction, although they cannot distinguish the effects of mutant p53 from those of p53 loss. Our associations were observed at the specimen level, the unit at which morphology, isoform and TP53 status varied within patients (Sections 3.1.2 and 3.1.3), and the TP53– morphology association was replicated across patients in the larger independent biobank (Sections 3.1.5 and 4.1). The novelty is differentiated along three axes. First, although the dual-promoter biology of MDM2 is established [6,9,59], no report has used MDM2 promoter usage as a locus-level readout of p53 activity in CRC (supporting observations: P2-transcript induction with wild-type p53 stabilization in oral cancer [122]; independence of MDM2 amplification from SNP309 and TP53 status in CRC [123]). Second, recent CRC lineage-plasticity work centers on PROX1 [124] and HNF4A/ATRX [125], and the gastric-metaplasia origin framework of serrated tumorigenesis is established [98,99]; our data add that the gastric-metaplastic genes follow TP53 status rather than MDM2 promoter usage (in the independent biobank and in tumors), extending to organoids the lower frequency of p53-pathway alterations reported in mucinous colorectal carcinomas [14]. Third, whereas CMS [2], CRIS [126] and iCMS [15] are multigene-signature classifications, this study is methodologically orthogonal in unifying a CMS2-like type and a type centered on CMS3 with CMS1-like immune features through TP53 status and a single locus-level readout of p53 activity. In the independent biobank of Kryeziu et al. [36] (GSE294511), the H&E cystic, mixed and solid categories did not correspond to the array P2 index, and their cystic marker axis (CDX2 high, KRT20 low) was unrelated to our Type1/non-Type1 axis in the discovery cohort (ρ = 0.08, P = 0.55; Additional file 7: Supplementary Table S48, sheet F); the two classifications therefore overlap only partly (Section 3.1.5).

### 4.5 Clinical implications

Two exploratory therapeutic hypotheses follow from the data. First, because P2-dominant organoids retained (at the patient level) wild-type, transcriptionally active p53, MDM2 inhibitors are the most direct candidates for TP53 wild-type, P2-dominant tumors (Sections 3.2.4 and 3.2.5). In cell lines the P2 signature tracked measured nutlin-3a sensitivity and MDM2 dependency after adjustment for TP53 status, although the effect was small; the cell-line P2 index tracked MDM2 dependency beyond total MDM2, but nutlin-3a sensitivity only weakly beyond it (Section 3.2.4). In organoids from independent patients (GSE294511) idasanutlin sensitivity followed the array P2 index among wild-type organoids, though not separably from total MDM2 (Section 3.2.5). Whether P2_index predicts clinical response independently of TP53 status is untested (Section 4.6). Second, on the P1 side, ATR/CHK1/WEE1 checkpoint inhibitors showed the clearest P1-associated sensitivity in cell lines (Section 3.2.4). The 39-gene panel (Section 3.3.3; Supplementary Figure S5, Additional file 52) would need independent validation before use as an assay. Organoid morphology, which followed TP53 status in both organoid cohorts and its gene program in tumors, could at most pre-screen for TP53 status rather than for promoter usage; both require validation in biopsy tissue. The physical states inferred from gene expression (a dense solid in tissue 1, a viscoelastic mucin gel with elevated interstitial fluid pressure in tissue 2) may limit drug accessibility [127,128] and motivate anti-VEGF [129] or mucolytic combinations, but were not measured here (Additional file 60: Supplementary Table S45; Supplementary Note Section 5.3). Established CRC therapies — standard chemotherapy, anti-EGFR antibodies in RAS/BRAF wild-type left-sided tumors, anti-HER2 therapy in HER2-amplified cases [130], sotorasib plus panitumumab for KRAS G12C [131] and encorafenib plus cetuximab for BRAF V600E [132] — follow their own biomarkers. In cell lines, standard cytotoxics and EGFR inhibitors showed no P1-specific association, whereas MEK/ERK/BRAF inhibitors correlated with the P2 score (exploratory; Section 3.2.4). For MSI-High/dMMR tumors — a status that P2 dominance does not itself imply (Sections 3.2.2 and 4.2.1) — immune checkpoint inhibitors are indicated [133–137], and mucolytic pretreatment can restore organoid chemosensitivity [138]. For direct targeting of the P2 axis, MDM2 inhibitors [139] are the rational candidates for TP53 wild-type, P2-dominant tumors. Their clinical development is difficult as of October 2026: milademetan gave a 3.2% confirmed objective response rate in the phase II MANTRA-2 trial [140], the phase III MANTRA trial did not meet its primary endpoint [141], brigimadlin was discontinued [142,143], siremadlin development in acute myeloid leukemia was halted [144], idasanutlin failed in MIRROS [145,146], and navtemadlin remains in phase III (POIESIS) after its BOREAS trial did not meet its primary endpoint [144]. MDM2 degraders, which can overcome the feedback induction of MDM2 that limits MDM2 inhibitors, have shown more sustained p53-pathway activation and deep tumor regressions in TP53 wild-type Merkel cell carcinoma models [147], and would face the same question of patient selection. These trials select patients by TP53 wild-type status or MDM2 amplification alone. P2_index measures promoter usage itself and could, if it is shown to carry information beyond total MDM2 expression (Section 3.2.5), add a selection axis — as could the p53-target signature that predicted response to p53–MDM2 inhibitors better than TP53 status alone [17], eight of whose 13 genes followed P2 usage beyond TP53 status in our discovery organoids (exploratory; Section 3.1.4). MEK-plus-MDM2 [148] and MDM2-plus-anti-PD-1 [149] combinations are also rational in TP53 wild-type tumors. Further candidate agents and combinations are discussed in the Supplementary Report (Additional file 1), Section R-22.

### 4.6 Novelty and limitations of this study

The novelty of this study can be summarized in four points. (1) To our knowledge, this is the first observation in a human CRC organoid cohort that organoid morphology follows TP53 status (previously described in mouse intestinal tumor organoids and a small human series [121]) — compact and glandular in TP53-mutant and round, cystic–mucinous in wild-type organoids — beyond 20q and aneuploidy, in the discovery cohort (56 specimens from 20 patients; P = 0.013, specimen level) and in an independent biobank (GSE294511; P = 6.1×10⁻⁴; also on unadjusted patient means). The morphology gene program identified mucinous carcinoma and followed TP53 status beyond 20q, MSI, BRAF mutation and CIMP in patient tumors (Sections 3.1.3, 3.1.4 and 3.1.5; in tumors, mucinous histology has been linked to fewer p53-pathway alterations [14]). The correspondence of morphology with MDM2 promoter usage at the specimen level (median Type1 fraction 0.826 versus 0.444, P = 1.1×10⁻3) is consistent with this alignment with TP53 status. (2) We directly quantified promoter usage itself (P2_index) in TCGA, where it tracked TP53 status and showed a truncating < missense < wild-type gradient not seen in p53-target output (Section 3.2.2). (3) We provide evidence that, among TP53 wild-type samples, P2 dominance marks a state in which p53 is transcriptionally active. P2 was induced in a p53-dependent manner by nutlin, tracked p53 activity in non-hypermutated wild-type tumor tissue (though not beyond total MDM2) and in cell lines, and in cell lines tracked MDM2 dependency beyond both p53 activity and total MDM2, and tracked nutlin-3a sensitivity beyond p53 activity but only weakly beyond total MDM2 (Section 3.2). In an independent organoid biobank (GSE294511 [36]) P2 again tracked p53 activity, with a tendency beyond total MDM2, and followed idasanutlin sensitivity, though not separably from total MDM2 (Section 3.2.5). (4) We separate the P2 signature into a p53-target part that follows P2 usage beyond TP53 status in organoids (without circularity, confirmed in the independent biobank: pre-specified p53 activity score, P = 0.012; discovery P = 0.094; Sections 3.1.4 and 3.2.5) and a gastric-metaplastic part that, like morphology, follows TP53 status (in the independent biobank and in tumors) and not P2 (Section 3.1.4). Further contributions — the multilayered IPA/EnrichR framework, the interpretation of the toxicity indices and the link between expression-inferred physical properties and jamming/unjamming transitions of cell collectives [150] — are exploratory and are detailed in the Supplementary Report (Additional file 1), Section R-23.

The limitations are as follows (full statement in the Supplementary Report, Additional file 1, Section R-23).

i. The study is mainly bioinformatic, without cell-functional experiments that switch MDM2 isoforms.
ii. The cohort (21 patients, 63 samples) is small; the 13 P2-dominant specimens derive from six patients, so the 33 comparisons are not independent replicates, and the morphology–isoform difference was not significant after adjustment for patient (Section 3.1.2). Discriminative performance of the signatures in external cohorts (AUC, sensitivity, specificity) was not evaluated.
iii. Because the isoform axis emerged from comparisons that began with the morphological groups, the morphology–isoform correspondence is not fully independent; the single-specimen split SI and the four morphology-deviating specimens provide partial independent support.
iv. Because the P2-dominant group was defined partly through p53-target differences, its separation from mutant specimens by the p53 activity score (AUC 0.99) may partly reflect circularity (Section 3.2.1).
v. Variant tables were available for only 35 of the 63 discovery samples; the others were called from published mutation maps, and seven remained undetermined (Section 2).
vi. Because TP53 status and promoter usage largely coincided in the discovery cohort, the organoid analyses cannot fully separate p53 activity from TP53 genotype, and the conclusion that morphology follows TP53 status rather than promoter usage rests on post hoc, observational, adjusted analyses in 56 specimens (specimen level only), the independent biobank and bulk tumors (Sections 3.1.4 and 3.1.5) that cannot exclude that TP53 mutation marks the chromosomal-instability pathway more broadly than the 20q, aneuploidy, MSI, BRAF and CIMP covariates capture; within MSI tumors TP53 status was not associated with the morphology genes, and among non-MSI, BRAF wild-type, CIMP-low tumors the morphology difference was marginal after adjustment for 20q.
vii. Organoids lack stroma and immune cells and are grown in defined media, so culture conditions could themselves modulate p53 activity and morphology, which may differ from those of the tumor of origin.
viii. Chromosome-arm gains were inferred from arm-level mean expression, not measured by copy-number profiling, and chromosomal instability was not measured directly.
ix. The P1 score did not show prognostic independence after adjustment for stage, and P2_index showed no association with OS.
x. HTA2.0 is limited to exon-level resolution, and in RNA-seq P2_index and the cell-line P2 index are estimated from transcript quantification; junction-read P2 shares agreed with P2_index (ρ = 0.64 in tumors with 10 or more reads; Section 3.2.2), but transcription start sites were not mapped in tumors or organoids.
xi. MSI/MMR status was not determined in this study (documented for one patient only, HCT27, microsatellite-stable), so the resemblance of the P2 signature to MSI-like tumors is a signature-level resemblance that requires direct confirmation; however, P2_index did not differ among the CIN, GS and MSI subtypes in TCGA (P = 0.11; Section R-26), so there is no evidence that P2 usage itself tracks microsatellite instability.
xii. The predictive value of P2 beyond p53 activity is limited (on p53-target expression as a predictor of MDM2-inhibitor response, see [17–19]): the cell-line P2 index remained associated with nutlin-3a sensitivity and MDM2 dependency beyond the p53 activity score in GDSC and CRISPR but not in PRISM (Section 3.2.4), in the 65 organoids of Herranz-Ors et al. [13] the P2 signature was not associated with nutlin-3 sensitivity after adjustment for TP53 status (FDR = 0.056), and in 14 wild-type GSE294511 organoids P2 usage could not be separated from total MDM2 (Section 3.2.5); P2 usage was not examined in patients, and MDM2 dependency only in cell lines.
xiii. The colibactin analysis (Additional files 61–63: Supplementary Figures S11 and S12, Supplementary Table S42) is exploratory.
xiv. The discovery cohort combines primary tumors with metastases, and sampling site (including the higher EMT scores of metastases, Section 3.1.2), prior chemotherapy (eight specimens) and passage number were not adjusted for in the P1–P2 comparisons.
xv. The public arrays of Sections 3.1.2 and 3.2.1 (GSE128213) are other specimens from patients of the discovery cohort; the independent biobank (GSE294511) comprised liver metastases only and recorded morphology on H&E sections rather than with our classifier, and no public bright-field organoid images with matched expression data were found, so the classifier itself was not tested on external images, only the morphology gene score derived from it.
xvi. Across all TCGA wild-type tumors the correlation between P2 and p53 activity was weak and quantification-dependent (ρ = 0.03–0.19), the ρ = 0.42 of Section 3.2.3 being the upper end of a definition-dependent range; bulk P2_index was inversely related to immune-cell content (PTPRC ρ = −0.47), so it may be influenced by tissue composition, and generalization beyond colorectal cancer is outside the scope of this article.
xvii. The proportion of wild-type tumors judged p53-inactive depends on the classifier (55% here, of the 118 tumors with a phenocopy score; 9% with the phenocopy threshold of Fito-Lopez et al. [20] and 99% with the classifier of Li et al. [21]), although the continuous measures agree in direction (Section 3.2.3). (xviii) Adjacent normal mucosa uses mainly P1, and TP53-mutant tumors also exceeded it (Section 3.2.2), so P2 usage in tumors also has inputs other than p53 activity; within TP53-mutant tumors the P2_index even ran opposite to p53-target output (Section 3.2.3), a bulk pattern we interpret cautiously.
xviii. The GDSC validation uses TP53 status as a surrogate rather than identifying the P2-dominant group directly, and the P1/P2 axis has not been mapped onto the iCMS/IMF framework [15].

## 5. Conclusions

In CRC organoids, morphology followed TP53 status, with no detectable added contribution from MDM2 promoter usage. TP53-mutant organoids mostly formed compact glands, whereas wild-type ones more often had the cystic–mucinous (non-Type1) morphology (deep-learning VGG16 classification, test accuracy 98.5%, 64/65; P = 0.013 at the specimen level after accounting for 20q and aneuploidy). In an independent organoid biobank (GSE294511) and in TCGA tumors, TP53 status also matched gastric-metaplastic programs, and in TCGA the organoid morphology genes identified mucinous carcinoma. By contrast, in organoids MDM2 P2 promoter usage reflected p53 activity; the evidence that it does so beyond TP53 genotype came mainly from the independent biobank and from two TP53-mutant, P2-dominant specimens. The p53-target genes followed P2 usage in both organoid cohorts, nutlin induced P2 in a p53-dependent manner, and in cell lines P2 followed MDM2 dependency after accounting for total MDM2. Whether TP53 loss itself drives the morphological change remains to be tested experimentally, and whether P2 usage adds predictive value beyond TP53 status and total MDM2 expression for selecting patients for MDM2-inhibitor therapy remains to be tested prospectively.

## List of abbreviations

5-FU: 5-fluorouracil
ACMG: American College of Medical Genetics and Genomics
AI: artificial intelligence
ALP: alkaline phosphatase
ALT: alanine aminotransferase
AMP: Association for Molecular Pathology
AST: aspartate aminotransferase
AUC: area under the receiver operating characteristic curve
BER: base excision repair
BH: Benjamini–Hochberg
BP: biological process
C-index: concordance index
CC: cellular component
CCD: charge-coupled device
CCLE: Cancer Cell Line Encyclopedia
CDK: cyclin-dependent kinase
CIMP: CpG island methylator phenotype
CIN: chromosomal instability
CLAIM: Checklist for Artificial Intelligence in Medical Imaging
CMS: consensus molecular subtype
COAD: colon adenocarcinoma
COSMIC: Catalogue of Somatic Mutations in Cancer
CR1: cluster-robust variance estimator of type 1
CRC: colorectal cancer
CRIS: colorectal cancer intrinsic subtypes
CRISPR: clustered regularly interspaced short palindromic repeats
CTRPv2: Cancer Therapeutics Response Portal version 2
DcR1/DcR2: decoy receptor 1/2
DDBJ: DNA Data Bank of Japan
DEG: differentially expressed gene
DepMap: Cancer Dependency Map
dMMR: mismatch repair-deficient
DMSO: dimethyl sulfoxide
DR5: death receptor 5
DSS: drug sensitivity score
ECM: extracellular matrix
EMT: epithelial–mesenchymal transition
ENCODE: Encyclopedia of DNA Elements
ERK: extracellular signal-regulated kinase
FC: fold change
FDR: false discovery rate
GDC: Genomic Data Commons
GDSC: Genomics of Drug Sensitivity in Cancer
GEO: Gene Expression Omnibus
GO: Gene Ontology
GOF: gain of function
GS: genomically stable
GSM: Gene Expression Omnibus sample accession
GSVA: gene set variation analysis
H&E: hematoxylin and eosin
HDR: homology-directed repair
HRR: homologous recombination repair
HTA2.0: Human Transcriptome Array 2.0
IARC: International Agency for Research on Cancer
IC50: half-maximal inhibitory concentration
ICI: immune checkpoint inhibitor
iCMS: intrinsic consensus molecular subtype
IFP: interstitial fluid pressure
IMF: classification combining iCMS, microsatellite instability and fibrosis
IPA: Ingenuity Pathway Analysis
IQR: interquartile range
JGA: Japanese Genotype-phenotype Archive
JUC: junction probe
LDH: lactate dehydrogenase
LM: liver metastasis
LN: lymph node metastasis
lncRNA: long non-coding RNA
LOF: loss of function
LuM: lung metastasis
MAPK: mitogen-activated protein kinase
MC3: Multi-Center Mutation Calling in Multiple Cancers
MCP-counter: Microenvironment Cell Populations-counter
MD5: message-digest algorithm 5
MEK: mitogen-activated protein kinase kinase
MF: molecular function
MMR: mismatch repair
MSI: microsatellite instability
MSI-High: microsatellite instability-high
MSI-L: microsatellite instability-low
MSS: microsatellite stable
NAFLD: non-alcoholic fatty liver disease
NHEJ: non-homologous end joining
NK: natural killer
OR: odds ratio
OS: overall survival
Ov: ovarian metastasis
PCA: principal component analysis
PD-1: programmed cell death protein 1
PDO: patient-derived organoid
pMMR: mismatch repair-proficient
PRC2: Polycomb repressive complex 2
PRISM: Profiling Relative Inhibition Simultaneously in Mixtures
PRO-cap: precision nuclear run-on and sequencing of 5′-capped RNA
PSR: probe selection region
READ: rectum adenocarcinoma
RMA: robust multi-array average
RNA-seq: RNA sequencing
ROC: receiver operating characteristic
RPPA: reverse-phase protein array
SG-NEx: Singapore Nanopore Expression project
SI: Splicing Index
SNP: single-nucleotide polymorphism
SSA: single-strand annealing
snoRNA: small nucleolar RNA
ssGSEA: single-sample gene set enrichment analysis
sst-RMA: signal space transformation robust multi-array average
TAC: Transcriptome Analysis Console
TCGA: The Cancer Genome Atlas
TCGA-CDR: TCGA Pan-Cancer Clinical Data Resource
TCR: T-cell receptor
TIDE: Tumor Immune Dysfunction and Exclusion
TPM: transcripts per million
TRIPOD+AI: Transparent Reporting of a multivariable prediction model for Individual Prognosis Or Diagnosis, updated for artificial intelligence
U-CAN: Uppsala–Umeå Comprehensive Cancer Consortium
UCSC: University of California Santa Cruz
UMAP: uniform manifold approximation and projection
VGG16: Visual Geometry Group 16-layer network
WHO: World Health Organization
WT: wild type

## Declarations

### Ethics approval and consent to participate

The patient-derived organoids and associated data used in this study were obtained with written informed consent from all patients [24], under approval by the Ethics and Medical Research Committee of the Japanese Foundation for Cancer Research, Tokyo, Japan (approval number: 2013-1105). The study was conducted in accordance with the Declaration of Helsinki.

### Consent for publication

Not applicable. This manuscript does not contain identifying personal details, images or videos of any individual person; the organoid microscopy images shown in Figure 1 are de-identified and cannot be linked to individual patients.

### Clinical trial number

Not applicable.

### Availability of data and materials

The HTA2.0 microarrays of the discovery cohort (63 specimens from 21 patients) were obtained within the patient-derived organoid biobank (the HCT series) whose establishment was reported by Okamoto et al. (2021) [24]; the microarray data of this biobank have been deposited by the originating laboratory in the Gene Expression Omnibus (GEO) under accession number GSE128213 [24]. Comparing the CEL files of the 63 specimens analyzed here against all files deposited in GSE128213, the array data of 60 specimens were identical to the deposited files (matching MD5 checksums). Specimen names in this article follow the GSE128213 titles (except HCT27-1T, titled HCT27-1T-2; Section 2); for six specimens the array file names differ (the files HCT17-Lv3, HCT33-4LM, HCT38-5LMR, HCT47-3LM-2, HCT50-4LMRR and HCT84-4LM correspond to HCT17-3LM, HCT33-4LMR, HCT38-5LM, HCT47-3LM, HCT50-4LM and HCT83-4LM, respectively), and the correspondence between file names, titles and GSM accessions is given in Additional file 20: Supplementary Table S1, sheet 3. The CEL files of the remaining three specimens (HCT27-1T, HCT33-4LMR and HCT33-5LMR; deposited in GSE128213 under the titles HCT27-1T-2, HCT33-4LMR and HCT33-5LMR, respectively) are not identical to those deposited files (presumably separate scans of the same specimens); these scan data derive from primary patient material, will not be newly deposited in a public repository, and are kept private (not publicly available). For these three specimens, the array data deposited in GSE128213 under the titles listed above are available. In addition, the single-cell RNA-seq data of the same biobank have been deposited by the originating laboratory in the DDBJ Japanese Genotype-phenotype Archive (JGA) under accession code JGAS00000000139 [24] (controlled access; not analyzed in this study). The sources, versions and retrieval procedures of the public datasets used for external validation and comparison (TCGA-COAD/READ, GSE39582, GDSC, DepMap/CCLE, the primary data of the patient-derived organoid biobank [Figshare, https://doi.org/10.6084/m9.figshare.28339340], and others, including GSE86221 (a SubSeries of GSE86222), DepMap 24Q2, PRISM, CTRPv2, the SG-NEx short- and long-read alignments, the ENCODE PRO-cap signal tracks and the TCGA data from UCSC Xena) are described in the Methods. The 14 non-discovery HTA2.0 arrays used for replication (Sections 3.1.2 and 3.2.1) are likewise identical to GSE128213 files (matching MD5 checksums). The independent organoid biobank used for replication (Sections 3.1.5 and 3.2.5) is available in GEO under accession number GSE294511 [36]; its per-organoid TP53 status, morphology and drug sensitivity were taken from the supplementary tables of the original publication. All analysis code used to compute the reported values is provided in Supplementary Methods (Additional file 2); plotting scripts not included there are available from the corresponding authors on request. Per-tumor P2_index values for the TCGA-COAD/READ tumors, with TP53 status and CMS class, are provided in Additional file 6: Supplementary Table S47. Correlations between P2 usage and p53 activity in TCGA tumors and cell lines, and the values of the replication in the independent organoid biobank, are provided in Additional file 7: Supplementary Table S48. The microscopy images (including the training and classified images for the deep-learning classifier) are primary patient-derived data and are not publicly available, because consent of the patients themselves for their public release cannot be obtained. The trained classifier weights (.h5) are likewise not publicly available, because consent of the patients themselves for their public release cannot be obtained.

### Authors’ contributions

T.T. conceived and designed the study, performed the deep-learning image classification and all bioinformatic and statistical analyses, and wrote the manuscript. K.S. supported the image classification in the early phase of the study. R.Y. provided the data and gave advice on the study. K.T. provided the research environment and gave advice on the analysis methods. All authors read and approved the final manuscript.

## Acknowledgements

We thank the members of the Division of Cell Biology, Cancer Institute, Japanese Foundation for Cancer Research, for establishing the patient-derived organoid biobank and for performing the organoid culture and microarray experiments, and the patients who donated the specimens.

## Additional files

Additional files are numbered in the order of first citation in the main text; the numbers of the Supplementary Tables and Supplementary Figures (S numbers) are identifiers from the analysis records and do not follow the order of citation.

Additional file 1: Supplementary Report: full-length analyses and extended discussion. (File format: DOCX) Full-length versions of the main-text sections presented in condensed form in the article (Sections R-1 to R-27), with a reference list at the end. Contains no analyses beyond the full versions of those summarized in the article (the TCGA junction-read counts are summarized in Section 3.2.2).

Additional file 2: Supplementary Methods: deposited analysis code and its index. (File format: ZIP) All analysis code used to produce the reported values (271 files), together with an index (README) that maps each script to the reported results and states its inputs, outputs, dependencies and run order. The files comprise R and Python analysis scripts and the shell scripts used for data download, reusable R and Python functions that compute the p53 activity score and the MDM2 P2_index from an expression matrix (with their check against Supplementary Table S47), including those of the patient-assignment check, the replication in further arrays (with the public-data restriction), the MSI-coding check and the p53-activity and genome-wide analyses of Sections 3.1.4 and 3.2 (with the plotting scripts of Figures 2–4, 6 and 7), the scripts behind Supplementary Table S48 (including the splice-junction counts of sheet D, the protein-level comparison of sheet E and the direct counts in SG-NEx cell lines of sheet G and the ENCODE PRO-cap analysis of transcription initiation at MDM2 P1 and P2 of sheet K), the replication in an independent organoid biobank (GSE294511) and its sheet F, the recall of TP53 status and the comparison of TP53 status with promoter usage (Supplementary Tables S11, S23 and S48, sheet H, with the plotting scripts of Figures 2, 3 and 9), the plotting and table-assembly scripts of Figures 1B, 1D and 8 and Supplementary Figure S5, Supplementary Figures S10 and S12 and Supplementary Table S34, the post-processing of Supplementary Figures S3 and S4, Jupyter notebooks, execution-environment records, the prediction/label, fraction and robustness-record data files of the image classifier, the values drawn in Supplementary Figure S10, the stratification of the EMT score by sampling site, MDM2 isoform and TP53 status behind sheet L of Supplementary Table S48, the drawing and cell-by-cell verification of Figure 10 with the per-specimen p53-activity table behind its discovery block (sheet M of Supplementary Table S48), the definition-sensitivity analysis behind sheet N, the drawing of Figure 9A, the redrawing of Figures 3B and 10, the assembly of the list of morphology genes in sheet O, and the stratification of the colibactin mutational-signature analysis by TP53 status and p53 activity behind sheet P (R168).

Additional file 3: Supplementary Table S2. Organoid morphology subtype versus MDM2 mRNA isoform. (File format: XLSX) Six columns: sample, Type0/Type1/Type5 fractions (re-classification with training-matched normalization), number of images, MDM2 isoform. Discloses the full basis for the morphology-isoform correspondence (Section 3.1.2).

Additional file 4: Supplementary Table S3. Sample composition of the 63 TAC comparison sets. (File format: XLSX) Three columns: analysis ID (49-144; non-consecutive TAC-internal numbering), P1-side samples, P2-side samples. The 33 sets used for the differential-expression and IPA comparisons are analyses 49-109; the remaining 30 (analyses 110-144) are the single-specimen split comparisons for the Splicing Index. Shows that no patient is duplicated within a comparison.

Additional file 5: Supplementary Table S11. TP53 mutation status of the 63 samples. (File format: XLSX) 16 columns: number, sample, patient ID, sample type, Type1 fraction, MDM2 isoform, TP53 status used in the analyses, source of the call, mutation detected, pathogenic variant, variant (isoform notation), canonical variant (NM_000546), dbSNP rsID, mutation type, ClinVar classification and functional impact.

Additional file 6: Supplementary Table S47. Per-tumor MDM2 promoter usage (P2_index), TP53 status and CMS class in TCGA-COAD/READ. (File format: XLSX) One sheet: 380 primary tumors × 6 columns (specimen, P2_index, TP53 status, TP53 functional class under definitions A and B, CMS class), with the definition of P2_index (transcript assignment by transcription start site, GRCh38) and five notes, including the reproduction of the values reported in Section 3.2.2 and Figure 5.

Additional file 7: Supplementary Table S48. P2 usage and p53 activity in TCGA tumors, organoids and cell lines, and replication in an independent organoid biobank. (File format: XLSX) Eighteen sheets: (A) correlation between P2_index and the p53 activity score in TCGA-COAD/READ TP53 wild-type tumors by TP53 call, hypermutation and expression quantification (TCGAbiolinks and Toil), with purity-adjusted values; (B1, B2) CTRPv2 nutlin-3 sensitivity and the cell-line P2 index (summary and 568 cell lines); (C) whether P2 usage carries information beyond total MDM2 expression, for the p53 activity score in TCGA tumors and organoids and for MDM2 dependency and nutlin sensitivity in cell lines; (D) MDM2 P1/P2 usage counted from splice-junction reads (recount3) in TCGA-COAD/READ by TP53 status, with its agreement with P2_index; (E) protein levels (RPPA) of BAX, p21 and p53 versus the p53 activity score, P2_index and total MDM2 in TCGA-COAD/READ; (F) replication in an independent organoid biobank (GSE294511): summary and per-organoid values; (G) MDM2 P1/P2 junction reads counted directly in SG-NEx cell lines with short and Nanopore long reads, and the DepMap cell-line P2 index; (H) morphology and expression programs by TP53 status and P2 usage in the discovery organoids, GSE294511 and TCGA-COAD/READ, including adjustment for BRAF mutation and CIMP in TCGA and a genome-wide comparison in the discovery organoids; (I) P2_index in TCGA adjacent normal tissue, wild-type and mutant tumors and patient-matched pairs; (J) comparison with published transcriptome-based classifiers of p53 function; (K) transcription initiation at MDM2 P1 and P2 measured by ENCODE PRO-cap in normal colon and stomach tissue and in Caco-2 cells, with the cell-line P2 index of Caco-2 from DepMap; (L) EMT score by sampling site, MDM2 isoform and TP53 status in the 63 discovery organoids; (M) the per-sample values drawn in Figure 10 (discovery organoids, GSE294511 and TCGA-COAD/READ), with the source of each block; (N) the analysis of sheets A and C repeated across 12 combinations of TP53 call, hypermutation definition and expression source; (O) the organoid morphology genes — the 141 transcript clusters that make up the morphology gene score, with their correlations, chromosome arms and the genes used in TCGA, and the gene symbols of each side listed together (O3); (P) the colibactin mutational-signature activities (SBS88 and ID18) in TCGA-COAD/READ by TP53 status, hypermutation and p53 activity, and in the Nunes cohort (TP53 status not available) against the p53-target score in all and in MSS specimens, testing whether their shift toward the P2-dominant side runs through p53 activity; and a README.

Additional file 8: Supplementary Table S4. Complete gene-level list of differentially expressed genes (1,890 genes). (File format: XLSX) Twelve columns including gene symbol, description, mean/median linear fold change, log2FC, consistency, number of comparisons, direction and probe set IDs. 1,013 genes higher on the P1 side and 877 higher on the P2 side.

Additional file 9: Supplementary Table S5. Robustness of the differentially expressed gene definition. (File format: XLSX) Four sheets: threshold sensitivity analysis, permutation test, reproducibility of representative marker genes, and notes.

Additional file 10: Supplementary Figure S1. Robustness of the differentially expressed gene definition to threshold choice. (File format: PDF) Number of differentially expressed genes as a function of the consistency and fold-change thresholds, showing that the gene set is not an artifact of one arbitrary cut-off. Analysis code in Additional file 2.

Additional file 11: Supplementary Figure S2. Sign consistency versus log2 fold change for all genes. (File format: PDF) Scatter plot of per-gene sign consistency against log2 fold change, with the adopted criteria (consistency ≥ 70% and |log2FC| ≥ 1) marked. Per-gene values computed by Code5 (Additional file 2) from Additional file 36; the plotting script is available from the corresponding authors on request.

Additional file 12: Supplementary Information. (File format: DOCX) Supplementary Results 1-13 (supplementary methods, results and discussion), legends for the Supplementary Figures provided as Additional files 10, 11, 22, 23, 52, 43, 44, 27, 45, 30, 61, 62 and 49, and the index of the deposited analysis code, supporting the main text.

Additional file 13: Supplementary Table S10. Sensitivity analysis of the IPA selection criteria. (File format: XLSX) Stability of the extracted pathways, upstream regulators and functions when the sign-consistency, half-rank and |median z| thresholds are varied. The core hubs (FOXM1/MYC/E2F and TP53/CDKN1A) keep their assignment throughout.

Additional file 14: Supplementary Note: Molecular landscape of the two organoid groups. (File format: DOCX) The methods of the IPA and EnrichR analyses (Section 1) and their results for the two organoid groups (Sections 2–4), with an extended discussion of the molecular landscape that complements the main Discussion, covering the necrotic cell-death hypothesis, the colibactin findings and the mechanotransduction axis. References cited only in Additional files 12 and 14 are listed in those files.

Additional file 15: Supplementary Table S28. Overlap with CMS marker genes (Fisher exact test). (File format: XLSX) Eight columns: query gene set, CMS class, reference genes on the array, overlap (n and %), odds ratio, one-tailed P and BH-FDR. Eight tests (4 CMS classes × 2 directions) against two independent reference sets, plus a sensitivity analysis excluding this study’s own signature genes.

Additional file 16: Supplementary Table S46. Per-sample morphology-class fractions under both preprocessing conditions. (File format: XLSX) One sheet, 11 columns: sample, MDM2 isoform, number of images, Type0/Type1/Type5 fractions under the as-applied condition and under re-classification with training-matched normalization, and non-Type1 fractions. Documents the robustness of the two-group separation to image preprocessing (Section 3.1.2).

Additional file 17: Supplementary Table S12. MDM2 Splicing Index across the 63 TAC comparison sets. (File format: XLSX) Eight columns: analysis, P1-side samples, exon 1 SI (three probes), exon 2 SI (two probes including the junction probe), P2-side samples. Includes the 33 comparisons used for the differential-expression and IPA analyses (analyses 49-109) and the single-specimen split comparisons (analyses 110-144). Underlying data for Figure 1.

Additional file 18: Supplementary Table S13. HCT38-3LM-specific EMT and mesenchymal program. (File format: XLSX) Two sheets: (1) gene, fold change in analysis 76, median across the other 32 comparisons, category and description; (2) notes. Shows the program is tumor cell intrinsic rather than stromal.

Additional file 19: Supplementary Table S14. Detailed profiles of the 4 deviating samples (extreme Type1 fractions). (File format: XLSX) Two sheets: (1) deviating marker profile; (2) per-sample assessment. Shows that the deviation is biologically interpretable rather than measurement error.

Additional file 20: Supplementary Table S1. Sample summary and EMT scores for all 63 samples. (File format: XLSX) Three sheets. (1) Sample name, patient ID, sample type, MDM2 isoform, Type1 morphology fraction (re-classification with training-matched normalization), deviation flag, EMT score and expression of mesenchymal, epithelial and functional genes. (2) Legend and notes. (3) Specimen names, array file names, GSE128213 titles and GSM accessions of the 63 discovery arrays, with MD5 identity to the deposited files. This is the ledger that every downstream analysis refers to.

Additional file 21: Supplementary Table S15. miR-200/ZEB1/2 axis expression in P1 versus P2 samples. (File format: XLSX) Six columns: gene, class, mean/median linear fold change, sign consistency toward P2, interpretation. Supports the partial-EMT rather than full-EMT reading of P2 samples.

Additional file 22: Supplementary Figure S3. Principal component analysis trajectories of the 63 organoid samples. (File format: PDF) PCA of RMA-normalized expression for all 63 samples (PC1 14.4%, PC2 9%). Arrows join primary to metastatic samples; the three patients with both P1- and P2-dominant samples are highlighted. Generated by the trajectory analysis code (Additional file 2), with HCT83-4LM assigned to patient HCT83.

Additional file 23: Supplementary Figure S4. UMAP trajectories of the 63 organoid samples. (File format: PDF) UMAP of the same data, showing the same separation of P1- and P2-dominant samples and the same long trajectories for the three patients with both P1- and P2-dominant samples. Generated by the trajectory analysis code (Additional file 2), with HCT83-4LM assigned to patient HCT83.

Additional file 24: Supplementary Table S24. Per-sample P1/P2 isoform assignment and Splicing Index. (File format: XLSX) Four sheets: (1) per sample; (2) primary-to-metastasis patterns; (3) data source notes; (4) replication in the 10 public organoid arrays not used for discovery, with the 11-array sensitivity analysis including HCT55-1T. Shows that the isoform is largely preserved from primary to metastasis.

Additional file 25: Supplementary Table S23. TP53 status, MDM2 isoform and morphology for the 63 samples. (File format: XLSX) Two sheets: (1) per-sample three-way correspondence; (2) summary. Gives the per-specimen correspondence of TP53 status, MDM2 isoform and morphology-class fractions (re-classification with training-matched normalization).

Additional file 26: Supplementary Table S26. Per-patient and per-comparison detail for key marker genes. (File format: XLSX) Two sheets: (1) per-patient summary; (2) per-comparison detail; includes the SNORD116 cluster rows (data only; Section 3.3.1). Source data for Supplementary Figure S8.

Additional file 27: Supplementary Figure S8. Per-sample expression of key marker genes (table-format figure). (File format: PDF) Linear fold changes of the key marker genes (gastric-type, Paneth-cell, p53-target and intestinal markers, the SNORD116 cluster and EGR1) for each of the 33 TAC comparisons, shown numerically with the patient of origin of the tissue-2 (P2)-side samples, with per-patient summary statistics, showing between-patient heterogeneity that group means hide (including the per-patient contrast in EGR1). All values are identical to Additional file 26 (Supplementary Table S26).

Additional file 28: Supplementary Table S18. Per-sample and per-patient EGR1 expression. (File format: XLSX) Two sheets: (1) per analysis; (2) per-patient summary. Primary data for the per-patient EGR1 expression (Sections 3.1.3 and 3.3.1).

Additional file 29: Supplementary Table S37. P2_index and p53 target module output by TP53 functional class. (File format: XLSX) One sheet with two blocks (P2_index and the p53-target module), each of six rows (three contrasts under two gain-of-function definitions; 12 and 13 columns), and four notes, for the 374 samples with both P2_index and TP53 mutation data. Uses both a narrow (six hotspot residues) and a broad (TP53 Database release 21 transactivation class) definition of gain of function.

Additional file 30: Supplementary Figure S10. MDM2 inhibitor sensitivity by TP53 and MSI status in 935 GDSC2 cell lines. (File format: PDF) Sensitivity to MDM2 inhibitors stratified by TP53 mutation status (primary) and MSI status (secondary) across the 935-line GDSC2 panel. Statistical analysis code in Additional file 2.

Additional file 31: Supplementary Table S38. All comparisons from GDSC1/GDSC2 and DepMap. (File format: XLSX) One sheet, 14 columns × 23 data rows in five blocks: pan-cancer by TP53 status, colorectal by TP53 status, colorectal by MSI status, DepMap CRISPR MDM2 dependency, and the integrated sensitivity score. Cell line genotypes unified to DepMap 24Q4.

Additional file 32: Supplementary Table S40. nutlin-3 sensitivity and P1/P2 signature scores in 65 patient-derived organoids. (File format: XLSX) One sheet, 22 columns × 65 rows: sample ID, log(IC50), TP53 mutation and loss-of-function status, KRAS, BRAF and MDM2 mutation status, MSI status, ploidy and 13 GSVA scores.

Additional file 33: Supplementary Table S41. Stratified analyses of the same 65 organoids (eight blocks). (File format: XLSX) Eight numbered blocks (block 6 with two sub-blocks) plus notes 1-12: sensitivity by TP53 status, linear model with covariates, Spearman correlations (family = 13), partial correlation adjusted for TP53, gene set coverage, the 48 TP53-mutant lines only, leave-one-out, GSVA recomputed in the 48 lines, and other strata. Discloses the negative result (ρ = −0.241, FDR = 0.775).

Additional file 34: Supplementary Table S39. Drug-sensitivity analyses: oncoPredict results for the 198 drugs in GDSC2 and measured cell-line sensitivity and gene dependency. (File format: XLSX) Four sheets. S39: 14 columns × 198 rows: rank, drug, Spearman rho, P and FDR for P2, rho and FDR for P1, separation, Kruskal–Wallis P by CMS, predicted median IC50 per CMS class and n; nutlin-3a ranks first (ρ = −0.659, FDR = 8.4×10⁻⁷⁷). S39b: measured GDSC2 LN_IC50 versus the P1/P2 scores for 295 drug entries, unadjusted and as partial correlations adjusted for cancer type and TP53 status; nutlin-3a ranks first. S39c: DepMap CRISPR dependency of 17 target genes by TP53 status and versus the scores. S39d: S39b additionally adjusted for RAS and BRAF mutation status and within RAS/RAF wild-type lines.

Additional file 35: Supplementary Table S27. WNT pathway genes in P1 versus P2 samples. (File format: XLSX) Expression and sign consistency of WNT pathway genes toward the P1 or P2 side, with the distribution of the classical WNT targets.

Additional file 36: Supplementary Table S16. Probe-level signed linear fold changes across the 33 comparisons. (File format: XLSX) 67,528 probes × 37 columns: probe set ID, gene symbol, description, plus one linear fold-change column per comparison and the mean across comparisons (Mean linear FC). The primary data on which every gene-level analysis rests.

Additional file 37: Supplementary Table S7. IPA Upstream Regulator analysis across all 33 comparisons. (File format: XLSX) 45 columns including upstream regulator, assigned subtype, sign consistency, median/mean z-score, molecule type, predicted activation state, target molecules and mechanistic network, plus one z-score column per comparison; a second sheet lists the regulators with borderline median z (1.99–2.01).

Additional file 38: Supplementary Table S8. IPA Diseases and Bio Functions analysis across all 33 comparisons. (File format: XLSX) 44 columns: disease or function, assigned subtype, sign consistency, median/mean z-score, predicted activation state, number of molecules, molecules, plus one z-score column per comparison.

Additional file 39: Supplementary Table S9. IPA Toxicity Function analysis across all 33 comparisons. (File format: XLSX) 44 columns in the same layout as Additional file 38. Documents the repeated detection of elevated-ALP signals on the tissue 2 side, which meet the adoption criteria, and the AST, ALT and LDH elevation signals, which are in the same direction but below threshold.

Additional file 40: Supplementary Table S31. EnrichR Gene Ontology analysis (BP/CC/MF for each tissue). (File format: XLSX) Six sheets (BP, CC, MF for P1 and P2). Columns: rank, term with GO ID, overlap, P value, adjusted P value, odds ratio, combined score, significance and genes. An analysis platform independent of IPA.

Additional file 41: Supplementary Table S6. IPA Canonical Pathway analysis across all 33 comparisons. (File format: XLSX) 42 columns: pathway, assigned subtype, sign consistency, median/mean z-score, number of analyses, front-half/back-half rank, molecules, plus one z-score column per comparison. Full numerical basis for Table 3.

Additional file 42: Supplementary Table S34. CMS attribution broken down by functional module. (File format: XLSX) Eleven modules (with three MDM2-excluded sets in the GSVA block) × CMS class: median and mean per class, top class, Kruskal–Wallis P and pairwise BH-adjusted P (class sizes in the note under the title), with GSVA and ssGSEA giving the same class ordering. Specimen-level.

Additional file 43: Supplementary Figure S6. GSVA signature scores by CMS subtype in TCGA-COAD/READ and by mismatch-repair status in GSE39582. (File format: PDF) P1 and P2 signature scores across CMS subtypes and between dMMR and pMMR tumors; the full legend is given in the Supplementary Information (Additional file 12). Analysis code in Additional file 2.

Additional file 44: Supplementary Figure S7. Overall survival in TCGA-COAD/READ by signature score. (File format: PDF) Kaplan–Meier curves by the median P1 and P2 signature scores and the multivariable Cox model; the full legend is given in the Supplementary Information (Additional file 12). Analysis code in Additional file 2.

Additional file 45: Supplementary Figure S9. CMS specificity of the P1/P2 signatures (sensitivity analysis). (File format: PDF) (A) Gene overlap between the signatures and the CMScaller templates (at most three genes per CMS class). (B) Association strength of each functional submodule with each CMS class. Values in Additional file 42 (Kruskal–Wallis P) and in the figure legend in Additional file 12 (overlap counts); the plotting script is available from the corresponding authors on request.

Additional file 46: Supplementary Table S30. Immune checkpoint gene expression in P2 organoids. (File format: XLSX) Seven columns: gene symbol, protein, checkpoint function, mean/median fold change toward P2, consistency and targeted therapeutic. Consistencies range from 84.8% to 66.7%.

Additional file 47: Supplementary Table S32. IPA Regulator Effects analysis (causal cascades). (File format: XLSX) 12 columns including analysis, cascade ID, consistency score, rank within analysis, node/regulator/target counts, regulators, target molecules and diseases/functions. Traces the cascade from EGR1 through TP53/VEGFA to ALP elevation.

Additional file 48: Supplementary Table S19. IPA Networks analysis by comparison. (File format: XLSX) Six columns: analysis, network ID, score, focus molecules, top diseases and functions, molecules in network. Underlying data for Additional file 49: Supplementary Figure S14.

Additional file 49: Supplementary Figure S14. Integrated hub-molecule network based on the IPA Graphical Summary. (File format: JPG) Central hub molecules recurring across the six IPA analysis modules over all 33 two-group comparisons, integrated by the authors into a single network using Cytoscape version 3.10.4 (red = P1/tissue 1, blue = P2/tissue 2). Supplementary Figure S14; the full legend is given in Additional file 12. Underlying data in Additional files 50 and 51.

Additional file 50: Supplementary Table S20. IPA Graphical Summary hub molecules. (File format: XLSX) Four columns: molecule, degree, activation sign, assigned subtype. Makes the integrated network of Supplementary Figure S14 reproducible.

Additional file 51: Supplementary Table S21. IPA Graphical Summary directed relationships. (File format: XLSX) Three columns: from molecule, to molecule, activation sign. Lists every edge from which the Supplementary Figure S14 network was assembled.

Additional file 52: Supplementary Figure S5. Hierarchical clustering of the 63 samples by the 39-gene panel identified by IPA Biomarker Detection. (File format: PDF) Heatmap of the 39 genes (Manhattan distance, average linkage) with the MDM2 promoter usage of each sample; the full legend is given in the Supplementary Information (Additional file 12). Plotting script in Additional file 2.

Additional file 53: Supplementary Table S22. IPA Biomarker Detection 39-gene panel. (File format: XLSX) Eight columns including symbol, Entrez gene name, location, family, drugs, Affymetrix probe set, number of analyses detected and functional category (9/5/2/1/9/2/11 = 39). The gene set used for the clustering in Supplementary Figure S5.

Additional file 54: Supplementary Table S43. Key items of the biomarker and GO analyses. (File format: XLSX) Representative genes for five biomarker categories (p53 activation markers, gastric metaplasia markers, invasion and metastasis markers, DNA repair and survival, stemness and Wnt) and the top GO-MF, GO-CC and GO-BP terms.

Additional file 55: Supplementary Table S35. Immune deconvolution (quanTIseq, MCP-counter) and TIDE results. (File format: XLSX) Two sheets. (1) 12 columns × 22 rows for the two deconvolution methods. (2) 11 TIDE metrics × 9 columns. Stratification by P2 score tertiles (top 208 versus bottom 208).

Additional file 56: Supplementary Table S36. MSI-stratified sensitivity analysis of the immune findings. (File format: XLSX) One sheet, 12 columns × 33 rows: van Elteren test, MSS-only analysis, MSI-adjusted partial correlation and logistic regression. Shows that the cell composition findings survive MSI adjustment but the composite TIDE score does not.

Additional file 57: Supplementary Table S44. Comparison of the upstream and downstream aspects of MDM2 promoter regulation. (File format: XLSX) Five aspects (upstream regulators, promoter state, downstream consequences, the role of MDM2, and the causal relationship) contrasted between tissue 1 (P1-dominant) and tissue 2 (P2-dominant), summarizing the working model in which promoter usage reflects p53 activity while morphology and, in the independent biobank and tumors, metaplasia follow TP53 status.

Additional file 58: Supplementary Table S29. Mechanotransduction machinery in P1-dominant versus P2-dominant specimens. (File format: XLSX) 92 genes × 10 columns: mechano-module, gene symbol, gene name, higher in, mean linear fold change, sign consistency, DEG flag, IPA upstream subtype, IPA median z and concordance note.

Additional file 59: Supplementary Table S33. Comparison with representative organoid morphology and deep learning studies. (File format: XLSX) Five columns × five rows (this study plus four prior studies): study and year, samples, imaging and AI method, main output, and relation to this study.

Additional file 60: Supplementary Table S45. Comparison of expression-inferred physical properties and Tox functions. (File format: XLSX) Five physical aspects (physical state, principal structural molecules, internal stress, drug penetration and Tox toxicity functions) contrasted between tissue 1 (autonomous-proliferation type) and tissue 2 (environment-adaptive type).

Additional file 61: Supplementary Figure S11. Colibactin mutational signature burden versus the P1−P2 axis (Nunes cohort). (File format: PDF) (A-F) SBS88 and ID18 activity against the P1−P2 GSVA axis in 1,051 U-CAN samples, including the MSS-only sensitivity analysis and the partial Spearman summary. Test values in Additional file 63 (computed by R42 in Additional file 2); the plotting script is available from the corresponding authors on request.

Additional file 62: Supplementary Figure S12. Reproduction of the colibactin-P1/P2 association in TCGA-COAD/READ. (File format: PDF) Forest plot of Spearman correlations between SBS88/ID18 activity and P2_index or the P1−P2 GSVA axis in the 374 TCGA samples with both measurements, including the tests that were not significant. Values and plotting script in Additional file 2.

Additional file 63: Supplementary Table S42. All tests of colibactin signatures versus the P1−P2 axis (Nunes cohort). (File format: XLSX) One sheet, 8 columns in two blocks: six primary tests on all 1,051 samples and six sensitivity/adjusted tests, each BH-corrected within a family of six, with nine provenance notes. Discloses the tests that were not significant.

