## Additional files for "TP53 status aligns with morphology and metaplastic programs in colorectal cancer organoids, and MDM2 P2 promoter usage reflects p53 activity": Additional_file_01_Supplementary_Report.docx

### Additional file 1: Supplementary Report — full-length analyses and extended discussion

This Supplementary Report provides, unabridged, the full-length versions of the main-text sections that are presented in condensed form in the article. Sections R-1 to R-27 correspond to the main-text sections indicated in each heading and reproduce the full text of those sections. All figure, table, Supplementary Figure/Table, and Additional file numbers cited below refer to the article’s figures, tables, and Additional files; this Report contains no analyses beyond the full versions of those summarized in the article and introduces no new figures or tables. A reference list is provided at the end of this Report; citation numbers [1]–[150] are shared with the article’s reference list, [151]–[172] with the reference lists of Additional files 12 and 14 (references not cited in the article), and [173] and [174] denote the references cited only in this Report. Sections R-25 to R-27 give the full text of the results that the article presents in condensed form in Sections 3.2.1, 3.2.3 and 3.2.4 (p53 activity and MDM2 P2), Section 3.1.4 (morphology, TP53 status and chromosomal instability) and Sections 3.1.5 and 3.2.5 (replication in an independent organoid biobank); their methods are reproduced in Section R-2, and their discussion is extended in Sections R-18, R-23 and R-24. Subsection numbers of the form 4.2.1–4.2.4 and 4.5.1–4.5.3 refer to the full-length subsections of this Report (see the headings of Sections R-18, R-19 and R-22), not to the article.

### R-1. Full-length version of main text Section 1 (Background)

### 1. Background

Treatment of colorectal cancer (CRC) is becoming increasingly individualized according to molecular subtype, but implementable biomarkers that can be determined directly from biopsy specimens and that are linked to treatment selection remain limited. TP53 is mutated in most chromosomally unstable (CIN) CRCs, yet TP53 genotype is an imperfect proxy for p53 function, and although TP53 mutations are known to be unevenly distributed across the histological and molecular types of CRC, how far TP53 status itself, rather than the features that accompany it, relates to the morphology and lineage programs of the tumor is not well defined. This study asks how TP53 status, p53 activity and the usage ratio of the constitutive MDM2 promoter (P1) and the MDM2 promoter induced by wild-type p53 (P2) (the P1/P2 ratio) relate to two contrasting CRC phenotypes, and whether P2 usage could serve as a readout of p53 activity linked to MDM2-inhibitor therapy. The contrast between P1-dominant, largely TP53-mutant (autonomously proliferative, CIN/CMS2 type) and P2-dominant, largely TP53 wild-type (environment-adaptive, gastric-metaplasia type [acquisition of gastric mucous-cell-like traits by colorectal tumors; Section 3.3.1] with a signature-level resemblance to microsatellite-instability [MSI]-like tumors) identified in patient-derived organoids was examined through its expression signatures in two independent cohorts totaling 1,143 cases (TCGA-COAD/READ and GSE39582; the image-based morphological correspondence could not be tested in these cohorts, and a morphology gene program derived from it was examined in an independent organoid biobank (GSE294511) [Sections 3.1.5 and 3.2.5 of the main text], where its across-sample alignment with P2 was largely explained by TP53 status); the P1 signature score was not associated with prognosis independently of stage (Section 3.3.2), so no prognostic use is proposed. Below, we first describe the molecular heterogeneity of this disease and the background of MDM2 promoter biology, and then discuss the clinical implications of stratification by TP53 status and the P1/P2 ratio.

With an estimated 2.0 million new cases worldwide in 2024, colorectal cancer is the third most commonly diagnosed cancer [1]; in Japan, it has the highest incidence among gastrointestinal cancers. Its molecular heterogeneity is extremely high, and the consensus molecular subtype (CMS) classification proposed by Guinney et al. in 2015 [2] divided CRC into four subtypes—CMS1 (MSI immune), CMS2 (canonical), CMS3 (metabolic), and CMS4 (mesenchymal)—and showed that each subtype has a distinct prognosis and therapeutic responsiveness. However, much remains unresolved regarding the molecular driving mechanisms that determine why particular subtypes show dramatic phenotypic transformation (metaplasia) or distinctive physical properties.

MDM2 (Murine Double Minute 2) has long been known as the principal negative regulator of p53 [3,4], and its transcription is finely controlled by two major promoters—the constitutive promoter (P1) and the promoter induced by wild-type p53 (P2). Barak et al. (1993) [5] showed that wild-type p53 induces mdm2 expression; the P2 promoter carries p53 response elements (p53REs) and is directly induced by the activation of p53 [6,7]. P1, in contrast, drives p53-independent basal transcription and maintains the basal protein turnover of the cell. In mice, point mutations in the two p53 response elements of the Mdm2 P2 promoter show that this feedback is dispensable for p53 stability, development and longevity but limits p53 activity after DNA damage [8]. Phelps et al. (2003) [9] showed that P2 can also be activated in a p53-independent manner through multiple transcription factor response elements in estrogen receptor-positive breast cancer cells, revealing a complex mechanism of P2 regulation in cancer cells.

A window through which this molecular basis can be viewed from the side of the observable phenotype is organoid morphology. It is widely known that changes in cell state, typified by epithelial–mesenchymal transition (EMT), are accompanied by a change from epithelial luminal and budding structures to a mesenchymal, scattered morphology, and recent studies have reported attempts to quantify organoid morphology by image-based computational analysis and to link it to the underlying cell state and drug response (the deep-learning-based association of morphology with EMT state and drug response in breast cancer organoids [10]; quantification of the phenotypic landscape of normal intestinal organoid regeneration by high-content image-feature profiling [11]). These suggest that morphological differences may reflect the cell states and molecular programs that underlie them. From this perspective, the present study took as its starting point the question of what molecular axis underlies morphologically contrasting CRC organoids.

In this study, we studied two groups of CRC organoids that, despite carrying the same diagnosis, differ markedly in morphology and in their MDM2 P1/P2 usage ratio, and we aimed to elucidate their decisive differences through an integrative analysis combining deep-learning (VGG16) morphological classification of organoids, MDM2 Splicing Index analysis, gene expression profiling, multilayer IPA analysis (Ingenuity Pathway Analysis, a commercial knowledge-base tool that infers pathways and upstream regulators from expression changes; all 8 modules), EnrichR GO analysis (a gene-set enrichment analysis tool; GO MF/CC/BP, 2026 version), and TP53 targeted resequencing (the composition of each module and the versions of the reference databases are given in Section 2). Note that the patient-derived organoids analyzed in this study (21 patients, 63 samples, the HCT patient group) derive from the same patient-derived organoid biobank (the HCT series) as the cohort in which Okamoto et al. (2022) [12] reported inter-patient heterogeneity using AI-based 6-morphology typing, with largely overlapping patients. We consider the six-type morphological typing of that report to be a biologically meaningful classification that captures inter-patient heterogeneity. At the same time, we noted that these diverse organoid morphologies appeared to separate broadly into two groups — a group dominated by a compact glandular morphology and a group dominated by round (cystic–mucinous) morphology — and we undertook this study aiming to identify the molecular axis that defines this two-group separation. This study independently reanalyzes this biobank-matched cohort and complementarily extends the previous work in that it links organoid morphology to TP53 status and to a transcript-level molecular axis, the MDM2 mRNA isoforms (P1/P2), as well as to prognosis and therapeutic stratification.

The working hypothesis that motivated this study was that MDM2 P1/P2 promoter choice is not only a phenotypic consequence reflecting differences in genome instability pathways, but may also contribute to the organ identity (lineage) of cancer cells, the physical properties of the tumor tissue (inferred here from gene expression), and the mode of interaction with the microenvironment. The data instead give TP53 status the central position: morphology and, in the independent biobank and tumors, the gastric-metaplastic programs followed TP53 status, whereas P2 usage followed p53 activity (Section 3.1.3 and Sections 3.1.4, 3.1.5 and 3.2.5 of the main text), and no causal role was tested (Section 4.6). The morphologically exceptional cases were informative: in morphology, three of the four followed TP53 status rather than promoter usage (Section 3.1.3). The study aims to provide a candidate readout of p53 activity that could, after validation in biopsy tissue, refine the molecular discrimination and therapeutic stratification of colorectal cancer. It should be noted that gene dependency maps of patient-derived tumor organoids are already being assembled on a large scale [13]. What this study adds is not the dependency map itself, but a candidate locus-level readout of p53 activity (MDM2 P2 usage) that may refine TP53-based stratification of those dependencies.

Two further questions follow from the dual-promoter biology of MDM2. First, TP53 genotype is an imperfect proxy for p53 function [16]: wild-type tumors can lack p53 transcriptional activity, and whether p53-target expression predicts MDM2-inhibitor response beyond genotype is debated [17–19]; transcriptome-based classifiers likewise indicate that a proportion of TP53 wild-type tumors have lost p53 function [20,21], and a TP53-loss phenocopy signature (an expression state resembling that of TP53-mutant tumors in the absence of the mutation) is also present in many tumors without biallelic TP53 alterations [22]. Because P2 carries p53 response elements, its usage could reflect p53 activity directly at the locus of the drug target. Second, mucinous carcinomas carry fewer p53-pathway alterations [14], and in the iCMS classification TP53 mutations are associated with the chromosomally unstable iCMS2 class, whereas almost all MSI tumors fall into iCMS3 [15]; whether these associations reflect TP53 status itself or the features that accompany it has not been separated, and it is unknown what genome-wide axis organoid morphology follows and whether it corresponds to chromosomal instability or mucinous histology in patient tumors. This study therefore also asks whether P2 usage reflects p53 activity rather than TP53 genotype and relates to MDM2-inhibitor sensitivity (Section 3.2 of the main text), what genome-wide axis organoid morphology follows — chromosomal instability, TP53 status or the MDM2 axis (Section 3.1.4 of the main text) — and which expression programs follow TP53 status and which follow P2 usage (Sections 3.1.4, 3.1.5 and 3.2.5 of the main text).

### R-2. Full-length version of main text Section 2 (Methods)

### 2. Methods

#### Patient samples and generation of organoids

This study used colorectal cancer tissues (primary tumors, liver metastases, lung metastases, lymph node metastases, and ovarian metastases) obtained from patients at a hospital. Organoids were generated from a total of 63 samples collected from 21 patients [23]. Two arrays registered under the patient number HCT84 (one of the 63 specimens and one array not used for discovery) derive from patient HCT83: the originating laboratory's sample records list these specimens as HCT83-4LM and HCT83-6LMR, the GEO title is likewise HCT83-4LM, and both arrays cluster with HCT83-1T and HCT83-3LM by sex-chromosome genes (all male) and genome-wide expression (mutual nearest neighbors; pairwise correlations 0.39–0.74, above the 99th percentile of between-patient correlations, 0.36; R84). Specimen names follow the titles in the public database (GSE128213), and patient-level analyses assign these specimens to HCT83 (the only patient-level analysis that counts HCT83-4LM as a separate patient is the genome-wide survey of Section 3.1.4 of the main text; its recomputation with 21 patients is given under Genome-wide survey of morphology and TCGA histology, below); for six specimens the names differ from the array file names (HCT17-3LM, HCT33-4LMR, HCT38-5LM, HCT47-3LM, HCT50-4LM and HCT83-4LM), and HCT27-1T keeps its file name because its GSE128213 entry is another scan of the same specimen titled HCT27-1T-2 (Additional file 20: Supplementary Table S1, sheet 3). This cohort derives from the same patient-derived organoid biobank as the patient group in which Okamoto et al. (2022) [12] reported inter-patient heterogeneity by AI-based morphological typing, with largely overlapping patients (that report analyzed 72 organoid lines derived from 26 patients, versus 21 patients and 63 samples in the present study), and the present study constitutes an independent re-analysis. The establishment and characterization of this patient-derived organoid biobank (the HCT series) — its patient-matched primary–metastasis pair design, its bulk transcriptome (HTA 2.0) and single-cell RNA-seq profiling, and its mutational profile with APC (91%), TP53 (79%) and KRAS (56%) as the most frequently mutated genes — are described in detail in Okamoto et al. (2021) [24]. Sample names were assigned in the format HCT (Human Colorectal Tumor) followed by the patient number and the sampling site and occasion (e.g., HCT25-1T: patient 25, first sampling, primary tumor; LM denotes liver metastasis, LuM lung metastasis, LN lymph node metastasis, and Ov ovarian metastasis). According to the biobank's records, eight of the 63 specimens were obtained after chemotherapy (HCT25-7LMR, HCT25-8LMRR, HCT25-9LMRR and HCT25-10LMRR after chemotherapy; HCT31-4LMR with chemotherapy; HCT38-3LM, HCT38-5LM and HCT38-6LM after four courses of FOLFOX plus bevacizumab), whereas HCT41-4LMR and HCT41-5LuMRR were obtained without chemotherapy; the passage number recorded in the array file names ranged from 1 to 22 (median 3; recorded for 43 specimens).

#### Whole-transcriptome analysis by microarray

Expression analysis was performed on the 63 samples from 21 patients using the Affymetrix Human Transcriptome Array 2.0 (HTA2.0). For each sample, 50 organoids were pooled for total RNA extraction and hybridized to a single array; each array therefore represents the average of 50 organoids. Organoid harvesting, RNA preparation and array hybridization followed the protocol of the biobank-originating laboratory [24]: organoids were recovered in recovery solution (Corning), total RNA was extracted with an RNeasy Micro Kit (Qiagen) and quality-checked on an Agilent Bioanalyzer, and 100 ng of total RNA was used to prepare cRNA with a 3′ IVT PLUS Reagent Kit (Affymetrix) for hybridization to the HTA2.0 array [24]. HTA2.0 carries two types of probes: (1) PSR probes (Probe Selection Region), which are designed to cover exonic regions and detect exon-specific expression; and (2) JUC probes (Junction Probes), which detect the junctions between adjacent PSRs that span intronic regions and are suited to discriminating splicing variants. As with exon arrays [25], this design allows quantification not only of the expression level of the gene as a whole but also of the usage rate of individual exons (splicing patterns). Two preprocessing streams were used according to purpose. (i) Two-group comparison stream: values computed from the CEL files (the raw array intensity files) with the sst-RMA algorithm (RMA, a standard normalization of array intensities, with signal space transformation) in TAC (version 4.02, the array manufacturer's analysis software) were used for the 33 two-group comparisons (gene- and probe-level fold changes; Additional file 4: Supplementary Table S3), for the definition of differentially expressed genes (DEGs) based on them, and for the Splicing Index analysis of MDM2; the TAC comparison results were the input to all IPA analyses, including Biomarker Detection. This step was run in the TAC graphical interface, and this fact was recorded at the beginning of Code2. (ii) Per-specimen expression stream: the CEL files were normalized with the RMA algorithm (R package oligo [26]); the script that executes RMA normalization (oligo::read.celfiles() → oligo::rma()) is provided as Code2 (Supplementary Methods (Additional file 2)). Code4 is a script that converts probe set IDs into gene symbols and does not itself perform RMA normalization. The gene-symbol-level matrix thereby obtained (exprs_matrix_symbol.csv; 33,720 genes × 63 specimens) was used for the EMT score, PCA/UMAP, and the within-patient paired analysis, and the probe-set-level RMA values were used for the hierarchical clustering of the 39 genes. For the replication analyses (main-text Sections 3.1.2 and 3.2.1), 19 arrays of the same biobank not used for discovery were normalized together with the 63 discovery arrays by RMA, and identity of the 14 of them that match files public in GSE128213 (2 normal and 12 tumor) was confirmed by matching MD5 checksums; one unpublished tumor array (HCT55-1T) was added only in a sensitivity analysis.

#### Image classification by deep learning (VGG16 transfer learning)

Deep learning was used to classify organoid microscopy images, which were acquired by the procedure used for this biobank [12] (organoids embedded in Matrigel, a basement-membrane matrix gel, in 48-well plates, imaged with a stereomicroscope equipped with a monochrome charge-coupled device (CCD) camera from 4 days after plating). Because the number of training images was limited, we adopted transfer learning based on VGG16 [27]. The classifier was trained on a training set of 129 images (90 used for actual training and 39 as the validation split) obtained by random image-level splitting (with a fixed random seed) of a total of 194 manually labeled images of the three morphologies Type0, Type1, and Type5 (Type0 84, Type1 53, Type5 57); on the test set of 65 images not used for training (Type0 30, Type1 19, Type5 16), the overall accuracy was 98.5% (64/65) (the confusion matrix and confidence intervals are given in Section 3.1.1 and Table 1). Here, Type1 (a densely packed glandular type) and Type5 (a dispersed mucinous type) are the two opposite morphologies on which this study focuses, whereas Type0 is a reference category that belongs to neither. All subsequent specimen-level quantification of morphology is based on the fractions of the images of each specimen classified into these three classes (Section 3.1.1). The computations were performed in Python using the Keras and TensorFlow libraries on a GPU-equipped laboratory server. The complete set of scripts for training and inference is provided as Code1 (Supplementary Methods), and the classification performance, the confusion matrix, and the unit of the training/test split are presented in Section 3.1.1.

Using the classifier thus constructed, a total of 792 images of the organoids from the 63 samples were classified into the three types Type0, Type1, and Type5. Taking the class with the largest softmax output as the predicted class of each image, we calculated the Type0, Type1, and Type5 fractions of each sample (the proportion of images predicted as that class) and defined the non-Type1 fraction (the fraction of round morphology) as the sum of the Type0 and Type5 fractions. The initial application had not normalized the input images; because collation of the deposited code revealed a mismatch with the training-time preprocessing (pixel values divided by 255), all 792 images were re-classified with the same normalization as in training, and all morphology-class fractions in this manuscript are based on this re-classification (classification code: Code1b, Supplementary Methods; the course of events and the comparison of the two conditions are given in Section 3.1.2, and the limitations in Section 3.1.1). No specimen- or patient-level majority-vote labels were created; morphology was quantified as fractions. The training images for the classifier and the classified images are primary patient-derived data and are not attached as supplementary data (not publicly available; see the Availability of data and materials statement). (See Additional file 3: Supplementary Table S2.)

#### Two-group comparative expression analysis with TAC

Using Transcriptome Analysis Console (TAC) 4.02 from Thermo Fisher, we performed comparative expression analyses between the group of samples with MDM2 P1 dominance and the group of samples with P2 dominance. The sample composition of each comparison was designed on the basis of the initial morphological classification (before unification of the preprocessing; Section 3.1.2) and the isoform assignment, and was in effect organized by isoform (12 of the 14 samples placed on the P2 side were P2-dominant, and 46 of the 47 samples placed on the P1 side were P1-dominant). Hereafter the P1-side group of samples is called “tissue 1” (P1-dominant, autonomously proliferative) and the P2-side group “tissue 2” (P2-dominant, environment-adaptive), and the two sides of each comparison are written the tissue-1 side and the tissue-2 side (Type0, Type1, and Type5 are used only as the names of the image morphology classes, and the expression signatures and their scores are named P1/P2; beginning of Section 3). sst-RMA was used as the preprocessing algorithm. A total of 33 comparisons were performed. Each comparison carries a TAC analysis ID (analyses 49–109; the IDs are TAC-internal serial numbers and are non-consecutive), and “analysis N” in the main text refers to this ID. The combinations of samples used in each comparison are shown in Supplementary Table S3, which lists the sample composition of all 63 comparison sets: these 33 (analyses 49–109) plus the single-specimen split comparisons for the Splicing Index (analyses 110–144).

As quality control for differentially expressed genes, the quality of the CEL files was checked in TAC, and microarray files with a low-quality Labeling Controls Threshold were excluded. In order to dilute the influence of the individual patient's genetic background, only one sample from a given patient was used in each comparison, and common tendencies were extracted by a majority vote over multiple comparisons. For the definition of differentially expressed genes (DEGs), taking the heterogeneity between cases into account, we used as the primary criterion not statistical significance (P value or FDR) but the consistency (reproducibility) of the direction of change across the 33 two-group comparisons. Specifically, we defined as DEGs those genes for which (i) using the gene-level value obtained by averaging the linear fold changes of the multiple probes for the same gene in each comparison, the sign of the change was the same in at least 70% of the 33 comparisons, and (ii) the absolute value of the gene-level mean fold change was at least twofold (|log2 fold change| ≥ 1). For fold change we used the signed linear values output by TAC (positive = higher expression in tissue 1, negative = higher expression in tissue 2); for genes with multiple probe sets, the linear fold changes of the individual probes were averaged to give the gene-level value, and the log2-transformed value is also given. By these criteria, 1,890 genes (1,013 with higher expression in tissue 1 and 877 with higher expression in tissue 2) were identified as DEGs. Representative DEGs are shown by functional category in Table 2, and the quantitative values for all genes (linear fold change, log2 fold change, and consistency rate) are given in Additional file 8: Supplementary Table S4 (the complete gene-level DEG list). The consistency rates given alongside individual genes in the main text (e.g., 73% for OLFM4) are transparency indicators that report the observed reproducibility of each gene.

We confirmed the robustness of this consistency-based definition by (i) a threshold sensitivity analysis and (ii) a sign-randomization permutation test (Additional files 9, 10 and 11: Supplementary Table S5, Supplementary Figure S1, and Supplementary Figure S2; analysis code: Code5, Supplementary Methods). When the consistency-rate threshold (60–80%) and the |log2 fold change| threshold (1.5-, 2-, and 3-fold) were varied, the number of DEGs changed smoothly (1,890 genes at the adopted criteria of 70% and twofold), and the tissue 1 : tissue 2 ratio (approximately 54 : 46) and the representative markers were preserved under every setting. In addition, when the signs were randomized while preserving the absolute fold-change value of each comparison and all criteria were recomputed (1,000 iterations, seed 42), the number of genes satisfying both criteria by chance was on average only 11.3 (SD 3.31, maximum 23), so that the observed 1,890 genes far exceeded chance, with an empirical FDR of 0.60% and a permutation P < 1×10⁻³ (z = 567). This quantitatively supports the view that the observed directional agreement is unlikely to arise by chance even under the present definition, which does not use P values or FDR as the primary criterion. Note that the denominator of the sign consistency rate is the number of comparisons for which a value was obtained for each gene (the number of non-missing comparisons), which in this data set was 33 for all genes. The consistency rate was defined as the proportion of the majority direction, max(number of positive values, number of negative values)/number of non-missing comparisons, whereas assignment to the tissue-1 side or the tissue-2 side was made according to the sign of the gene-level mean fold change (7 of the 1,890 genes showed a discrepancy between these two conventions, and this is stated explicitly in the Notes of Supplementary Table S5). Aggregation by gene symbol was based on exact matching, and composite symbol rows (such as “MUC5AC; MUC5B”) were not assigned to either constituent gene.

Note that the rule for integrating probes into genes differs among analyses (the present DEG analysis uses the mean of the fold changes; the PCA/UMAP and the EMT score use the median of the expression values; and the external cohort validation adopts the probe with the largest variance), and therefore the rule is stated at the relevant point of each analysis. Moreover, the EMT score calculation (Code3) and the PCA/UMAP analysis (Code6) both use the same gene-level expression matrix based on the pd.hta.2.0 annotation (exprs_matrix_symbol.csv, in which multiple probe sets were integrated by the median of the expression values). The settings of these auxiliary analyses were as follows. The EMT score was defined as the mean of the gene-wise z-scores, computed across all 63 specimens, of 11 EMT/mesenchymal genes (CDH2, VIM, FN1, SPP1, VCAN, TGFB2, TGM2, LOXL4, CDH6, ALDH1A3, and ONECUT2), minus the corresponding mean for 2 epithelial marker genes (CDH1 and EPCAM) (Code3, Supplementary Methods). Within-patient paired expression comparisons were performed for the 17 patients having paired primary and metastatic samples, using a linear model with the duplicateCorrelation method of limma [70] with the patient as a block, and paired DEGs were defined by FDR < 0.05 and |log2 fold change| > 1.0 (Code6, Supplementary Methods). PCA and UMAP [71] used the top 3,000 genes by variance; PCA was computed with prcomp (scale. = TRUE) and UMAP with the umap package (n_neighbors = 15, random seed 42) (Code6, Supplementary Methods).

#### Splicing Index analysis of MDM2 mRNA isoforms with TAC

Using the Splicing Index (SI) algorithm implemented in TAC, we compared the mRNA structure of the MDM2 gene. The Splicing Index is defined by the following equation.

Splicing Index = (Exon 1 Cond.1 Intensity / Gene 1 Cond.1 Intensity) / (Exon 1 Cond.2 Intensity / Gene 1 Cond.2 Intensity)

This algorithm measures how much exon-specific expression differs between two conditions after the influence of the gene expression level has been removed. A positive SI value indicates that the RNA of that probe region is more abundant on the tissue-1 side (Condition 1), and a negative SI value indicates that the RNA is more abundant on the tissue-2 side (Condition 2).

SI > 0: the RNA of that probe region is relatively more abundant on the tissue-1 side (Condition 1). SI < 0: the RNA of that probe region is relatively more abundant on the tissue-2 side (Condition 2).

MDM2 has a total of 12 major exons (Liang et al. 2004 [28]). In humans, transcripts initiated from the P1 promoter start at exon 1 and skip exon 2. Under stress conditions, transcription is initiated from the P2 promoter, and exon 2 is used instead of exon 1. Exon 1 can be detected by three PSR probes (PSR12007995: genomic positions (hg19/GRCh37, the annotation loaded on the array) 69,201,971–69,202,001; PSR12007996: 69,202,002–69,202,052; PSR12007998: 69,202,062–69,202,243), and exon 2 can be detected by JUC12004243 and PSR12008003 (69,202,804–69,202,866). Each specimen's dominant promoter was assigned from the MDM2 Splicing Index — P1-dominant when the exon-1 probes were relatively enriched on its side of the comparison and P2-dominant when the exon-2 probes were — and the call was checked by single-specimen comparisons against a common reference specimen of the opposite type (HCT64-1T for P1-dominant and HCT25-1T for P2-dominant specimens), taken from the 33 two-group comparisons (analyses 49–109) and from 30 additional single-specimen split comparisons (analyses 110–144, three of which used other reference specimens; Supplementary Table S3).

A note on nomenclature: nutlin-3 is the racemate, and nutlin-3a is its active enantiomer; their mechanism of action — occupying the p53-binding pocket in the N-terminal domain of MDM2 — is the same, and this manuscript retains the drug-name notation of each data source. Note also that MDM2 exon 1 (the first exon of P1-derived mRNA) and exon 2 (the first exon of P2-derived mRNA) are both 5′-untranslated, and the translation start codon of full-length MDM2 (the first in-frame AUG) lies in exon 3, which is shared by both transcripts in the human [6] and murine [59] genes; the nutlin-binding site of full-length MDM2 (the N-terminal p53-binding domain) is therefore invariant with respect to promoter choice. The two transcripts nevertheless differ in their usage of translation initiation codons, and initiation at downstream AUG codons has been shown for the murine transcripts, in vitro and in cells, to yield N-terminally truncated MDM2 polypeptides that cannot bind p53 [59]; because our measurements were made at the RNA level, the composition of such protein isoforms was not assessed. In this study, the Splicing Index — a general algorithm of TAC — was applied to the probes of MDM2 exon 1 and exon 2 to compare toward which of tissue 1 and tissue 2 the relative amounts of mRNAs containing each exon are shifted; it does not detect the MDM2 splice variants reported in cancer that lack internal coding exons, losing the N-terminal p53-binding domain while retaining the C-terminal RING finger domain, such as MDM2-A, MDM2-B, and MDM2-C [28].

#### Ingenuity Pathway Analysis (IPA) and EnrichR GO analysis

Using Ingenuity Pathway Analysis (IPA; Qiagen) [72], we independently performed seven analyses — (1) Canonical Pathway analysis, (2) Upstream Analysis, (3) Disease and Bio Functions, (4) Tox Function, (5) Regulator Effects analysis, (6) Networks analysis, and (7) Graphical Summary — on each of the comparison datasets from all 33 pairs contrasting tissue 1 with tissue 2, and the results were aggregated on the basis of the frequency of appearance using the comparison function of IPA. Data were analyzed through the use of IPA (QIAGEN Inc., https://www.qiagenbioinformatics.com/products/ingenuity-pathway-analysis)[72]. As criteria for adopting only highly reproducible, robust findings in the main text (hereafter the “adoption criteria”), we required that (i) the sign consistency of the Activation z-score across the 33 analyses be at least 75%, and that (ii) the absolute value of the median Activation z-score across the 33 analyses be at least 2 (|median z| ≥ 2); these two criteria were applied to Upstream Analysis, which claims attribution to a tissue type on the basis of the effect size of activation. The listings of Canonical Pathways and of Disease and Bio Functions are descriptions based on the rank of the frequency of appearance and on the reproducibility of the direction of activation (items with |median z| < 2 are explicitly indicated as sub-threshold); the Tox Function findings were used only as corroborating evidence supporting the other modules; and for Regulator Effects analysis alone, only the results of the first 8 of the 33 analyses were integrated because of constraints on computational load. In addition, the lists of differentially expressed genes obtained from the two-group comparison analyses (TAC) (the top 400 genes each) were subjected to Gene Ontology (GO) enrichment analysis using the Enrichr R package [73] (GO Biological Process, GO Cellular Component, and GO Molecular Function, 2026 versions; adj. P < 0.05 after FDR correction by the Benjamini–Hochberg procedure). The detailed settings of each analysis, the aggregation scheme (first-half/second-half Rank, N/M consistency), the rationale for the design of the adoption criteria and their scope of application, and the sensitivity analysis for the choice of the adoption criteria (Additional file 13: Supplementary Table S10; Supplementary Results 10, Additional file 12) are described in the Supplementary Note (Additional file 14), Section 1.

#### Biomarker Detection

Using the Biomarker Detection function of IPA, candidate biomarker molecules discriminating the two tissue types (39 genes) were extracted. The hierarchical clustering of the 63 samples using these 39 genes as input was then carried out separately with an R script written by the authors (the Manhattan distance was used as the distance measure, with average linkage as the clustering method. The framework of hierarchical clustering and heatmap display follows Eisen et al. (1998) [74], but the Manhattan distance was used rather than the uncentered correlation that the original paper takes as standard, and no scaling of rows or columns was performed).

#### TP53 mutation analysis

All 63 samples were analyzed for TP53 mutations by targeted resequencing performed by the biobank-originating laboratory [24]: genomic DNA (extracted with a QIAamp DNA Mini Kit, Qiagen) was used for HaloPlex custom-panel library construction (Agilent) and sequenced on a MiSeq system (Illumina), with variant calling by GATK HaplotypeCaller and ANNOVAR annotation as described in the supplemental procedures of [24]. Multiple isoform transcripts of TP53 (NM_000546: p53α; NM_001126115: Δ133p53β; NM_001126118: Δ40p53β) were used as reference sequences, with conversion to canonical numbering (e.g., R136H on NM_001126118 corresponds to R175H). Variants were classified by the ACMG/AMP criteria [29] (standard guidelines for rating variant pathogenicity) with reference to the public variant databases ClinVar [30] and dbSNP [31]; intronic and untranslated-region variants, and variants also called in a patient's non-tumor tissue, were not counted as somatic mutations. Per-specimen variant tables were available for 35 of the 63 samples; for the others, status was taken from the per-organoid mutation maps published for the same biobank (Fig. 1A in ref. [24] and Fig. S5 in ref. [12]), and two samples of patient HCT25 that appear in neither were assigned the patient's mutation, which was present in all six of its other samples. Seven samples whose status could not be settled (a single published map without a variant table; a variant table showing only the P72R polymorphism, from sequencing runs that missed a known mutation in another patient, against a map showing a mutation; or a map without a mutation against TP53 mRNA at the level of truncating mutants) were treated as undetermined and excluded from analyses using TP53 status, and sensitivity analyses assigned them to either group; TP53 status was thus determined for 56 of the 63 samples (Supplementary Table S11, which gives the source of each call).

The identified mutations were classified into four categories — Pathogenic, Likely pathogenic, VUS (Variant of Uncertain Significance), and Benign — according to the ACMG/AMP criteria [29] with reference to the NCBI ClinVar [30] and dbSNP [31] databases. The correspondence between isoform-specific mutation numbering and the amino acid numbering on the canonical sequence (NM_000546/p53α) was converted taking into account the difference in the translation start site of each isoform (Δ133p53: +133; Δ40p53: +40). For example, R136H on NM_001126118 corresponds to R175H on the canonical sequence.

Additional file 5: Supplementary Table S11, which organizes, one sample per row, the mutation detection status, the mutation name (in both isoform and canonical notation), the mutation type, the ClinVar classification, and the functional impact of each sample, is provided with this paper. An overview of the analysis results is shown in Supplementary Table S11.

#### Signature validation and survival analysis in external independent cohorts

The external validity of the organoid-derived expression signatures identified in this study (the P1 signature, 21 genes; the P2 signature, 29 genes) was assessed in two independent large-scale public cohorts. First, we used the COAD/READ cohort of The Cancer Genome Atlas (TCGA). Of the 701 cases obtained from the Genomic Data Commons (GDC) using TCGAbiolinks, 647 primary tumors (sample type=‘01’) were analyzed, and 51 normal tissues, 2 recurrent specimens, and 1 metastatic specimen were excluded. An expression matrix was constructed from the RNA-seq data (STAR-Counts, TPM, transcripts per million, an abundance unit normalized for transcript length and sequencing depth), and gene IDs were converted from Ensembl IDs to gene symbols and Entrez IDs using org.Hs.eg.db. When multiple Ensembl rows corresponded to the same gene symbol or the same Entrez ID, they were collapsed to a single representative row, namely the row with the largest variance (rowVars). This convention was applied both to the symbol matrix used as input to Gene Set Variation Analysis (GSVA, which scores each sample for a gene set)/ssGSEA and to the Entrez matrix used as input to CMScaller, and the same convention was also used for the expression matrix of GSE39582 described below. Consensus molecular subtype (CMS) classification was performed with CMScaller[75] (Entrez ID, RNAseq=TRUE, FDR < 0.05); of the 647 aliquot rows (624 specimens), 574 rows could be classified, corresponding to 563 specimens after aggregation at the specimen level (averaging over the first 15 characters of the sample barcode) (CMS1: 96, CMS2: 167, CMS3: 96, CMS4: 204; among the 13 specimens with duplicate aliquots there was no discrepancy in CMS assignment). All TCGA analyses below were performed per specimen (one record per specimen, aliquots averaged); this convention was adopted for this study and for the companion study of the 15q11-q13 locus (T. Tsukui, R. Yao, and K. Tsuda, unpublished observations), which analyzes the same TCGA specimens.

Second, GSE39582 [76] (platform, Affymetrix HG-U133 Plus 2.0) was obtained from the Gene Expression Omnibus (GEO) using GEOquery. Of the 585 samples in total (566 colon cancers and 19 non-tumoral mucosa samples), the 19 non-tumoral samples were removed before GSVA scoring (17 of them carry an MMR annotation in GEO — 2 dMMR and 15 pMMR — so they are not removed by the MMR filter), and the 519 tumors with annotated mismatch repair (MMR) status (75 corresponding to dMMR/MSI-H and 444 corresponding to pMMR/MSS; the 47 tumors without MMR annotation were excluded) were analyzed (analysis code: R49, Supplementary Methods). On this microarray platform, 20 of the 21 genes of the P1 signature (95%) and 27 of the 29 genes of the P2 signature (93%) were detected, both detection rates being 90% or higher. The three genes that could not be detected were NPSR1 (P1 signature) and HULC and BBC3 (P2 signature). For NPSR1 and HULC, no corresponding probe set exists on GPL570. For BBC3 a corresponding probe set (211692_s_at) does exist, but that probe set is assigned to multiple genes (MIR3190///MIR3191///BBC3), and because the GEO platform annotation does not allow the signal to be unambiguously attributed to a single gene symbol, it was not recovered as BBC3 in the primary analysis. We therefore confirmed, as a sensitivity analysis, that the conclusions of the main tests were unchanged with a 28-gene version of the P2 signature in which 211692_s_at was attributed to BBC3. MIR3190 and MIR3191 are miRNA genes that overlap the BBC3 locus, and because mature miRNAs are not measured on this array, the signal of this probe set may be attributed to BBC3. For dMMR versus pMMR, the normal approximation for the P2 signature as a whole gave z = 8.56 → 8.24 and P = 1.14×10⁻¹⁷ → 1.73×10⁻¹⁶, and for the immune-excluded version P = 6.6×10⁻¹⁰ → 4.1×10⁻⁹; the Spearman correlation between the GSVA scores of the 27-gene and 28-gene versions was 0.97 or above in every case (analysis code: the R45 procedure applied to the 519 tumors in R49, Supplementary Methods).

The P1 (21-gene) and P2 (29-gene) signatures used for validation were selected, on the basis of the lists of differentially expressed genes (DEGs) obtained in the two-group comparison of this study and in light of the results of the IPA, EnrichR GO, and Biomarker Detection analyses, as representative genes of the functional modules that constitute the P1/P2 model. The P1 signature (CIN type) was composed of (i) intestinal epithelial identity and stemness (OLFM4, LGR5, SLC26A3, SLC9A3, DPP4, ANPEP, MGAM2, SLC7A5, NPSR1, CALB1, ALDH1A1, AKR1B10), (ii) an intestinal master transcription factor (HNF4A; its expression was consistently higher on the tissue-1 side, with a mean of approximately +2.0-fold and a sign consistency of 94%, and it was therefore adopted as the representative of intestinal-type identity. Note that its estimated activation in the IPA Upstream analysis does not meet our adoption criteria (Section 3.3.1). The intestinal homeobox factor CDX1 is also a canonical intestinal identity transcription factor, but in this cohort its bulk expression was shifted toward the tissue-2 side, contrary to the expectation for intestinal identity (mean linear FC −1.56, sign consistency 85%; Section 4.3), and its IPA-estimated activity was also below the threshold, so it was not adopted for this signature. HNF4A (+1.98) and CDX1 (−1.56) both have effect sizes short of two-fold, but they differ in that HNF4A is consistently shifted toward the tissue-1 side, which is the expected direction), and (iii) an autonomous cell-cycle program driven by the MYC/E2F/FOXM1 axis (FOXM1, MYBL2, MCM4, MCM5, MCM6, TOP2A, AURKB, HELLS). MSH2 and ABCB1, which were initially considered as auxiliary indicators of the MMR-proficient, CIN context, were excluded from the final signature used for scoring, because MSH2 is an MMR gene and using it to predict MSI status would be circular, and because the functional-axis assignment of ABCB1 (a drug efflux transporter) is ambiguous. The P1 signature therefore comprises 21 genes, and this exclusion applies to all of the GSVA scoring analyses in this manuscript, not only to the survival analysis. The P2 signature (resembling MSI-like/serrated tumors at the signature level only) was composed of (i) lineage plasticity and metaplasia (gastric type: CTSE, REN, TFF1, TFF3, MUC5B, MUC5AC, MUC6, MUC17, ANXA10; Paneth cell type: DEFA5, DEFA6), (ii) transcriptional targets of wild-type p53 (CDKN1A, DDB2, ZMAT3, FAS, TNFRSF10D, SPATA18, GDF15, BAX, BBC3) and MDM2 itself, (iii) immune checkpoint molecules involved in immune evasion (IDO1, PDCD1, LAG3, CTLA4, TIGIT, HAVCR2), and (iv) cancer-associated long non-coding RNAs (UCA1, HULC). In particular, the p53 target genes and MDM2 were included in the P2 signature on the basis that P2 is a p53-inducible promoter [6,7], so that the expression of the activated p53 targets could serve as a functional indicator of P2 promoter usage. Each gene corresponds to a mechanistic axis of the P1/P2 model, and by testing whether these axes were coordinately reproduced in independent cohorts, we evaluated the external validity of the expression programs linked to morphology and to MDM2 promoter usage. The complete gene lists of the signatures are those given in this paragraph, and the same sets are embedded in the deposited analysis code (Code8, Supplementary Methods).

The gene selection for both signatures followed four principles. (Principle 1: data-driven selection) Candidates were selected from the genes that showed high directional consistency in the two-group comparison of this study, and were in principle limited to those meeting the DEG criteria of this study (the same direction in 70% or more of the 33 comparisons, and |mean linear fold change| of two-fold or more). Of all 53 candidates considered (50 finally adopted genes = 21 P1 and 29 P2, plus 3 non-adopted genes = CDX1, MSH2, ABCB1), 42 genes meet these criteria (listed in Supplementary Table S4). The 11 genes that do not meet the criteria are the 10 genes that were adopted with the unmet criterion explicitly stated (next paragraph) and CDX1, which was not adopted (the paragraph after next). (Principle 2: mechanistic correspondence and consistency with prior knowledge) Among the DEGs, priority was given to genes that represent each functional axis constituting the P1/P2 model (for P1: intestinal identity, stemness, intestinal master transcription factor, and autonomous cell cycle; for P2: metaplasia/lineage plasticity, wild-type p53 targets, immune evasion, and cancer-associated lncRNAs) and for which the function in question is established in previous reports. (Principle 3: robustness) Findings that stood out in only a single specimen, and genes that could derive from stromal contamination, were not adopted, because they would compromise the generality of the signature. (Principle 4: avoidance of circularity) For the gene sets used for scoring, a separate operational exclusion was applied in order to avoid circularity (the paragraph after next). Under these principles, the signatures were constructed not by arbitrary choice of the authors but on the objective basis of the DEGs, reflecting the mechanistic hypothesis.

The 10 genes adopted despite not meeting the DEG criteria, and the rationale for each, are as follows. First, the six immune checkpoint genes (IDO1, PDCD1, LAG3, CTLA4, TIGIT, HAVCR2; mean linear fold change −0.38 to −0.77, consistency 67–85%) do not meet the effect-size criterion (two-fold). These molecules derive mainly from tumor-infiltrating lymphocytes, and in bulk expression their effect sizes are structurally compressed because of dilution by the epithelial component. We therefore adopted the reproducibility of direction, rather than the effect size, as the rationale for their inclusion. However, HAVCR2 has a sign consistency of 66.7% and thus does not meet the consistency criterion (70%) either, and we state this point explicitly as a limitation. Second, the wild-type p53 targets BAX (mean linear fold change −1.89, consistency 97%) and BBC3 (−1.09, 94%) have effect sizes slightly below two-fold, but their directional consistency is 93% or more, and they represent the p53-target axis, which links the signature to P2, a p53-inducible promoter [6,7]; they were therefore adopted. Third, the intestinal master transcription factor HNF4A (+1.98, 94%) also has an effect size slightly below two-fold, but was adopted for the same reason as the representative of the intestinal-type identity axis. Fourth, CALB1, an intestinal identity gene, has a consistency of 69.7% (23/33), slightly below the consistency criterion, but was adopted because of its large effect size (+338). For all 10 of these genes, the item that is not met (either effect size or consistency) is explicitly stated in the main text and in Supplementary Table S4.

On the other hand, three genes were considered as candidates but not adopted for the final signatures, for the following reasons. First, CDX1 is a canonical intestinal identity transcription factor and was mechanistically a strong candidate, but in this cohort its bulk expression was shifted toward the tissue-2 side, contrary to expectation (mean linear fold change −1.56, sign consistency 85%), and its estimated activity in the IPA Upstream analysis also did not reach the adoption threshold (|median z| ≥ 2), so it was not adopted (Section 3.3.1, Section 4.3). The reason we adopted HNF4A (+1.98), whose effect size is likewise below two-fold, but did not adopt CDX1, is that HNF4A is consistently shifted toward the tissue-1 side, which is the expected direction, whereas for CDX1 the direction itself is contrary to expectation. Second, MSH2 was initially considered as an auxiliary indicator of the MMR-proficient, CIN context, but because MSH2 is itself an MMR gene, predicting MSI status with a signature that contains it would be circular. It was therefore excluded from the final signature used for scoring (Principle 4). This exclusion is applied consistently to all of the GSVA scoring analyses in this manuscript, not only to the survival analysis, and as a result the P1 signature comprises 21 genes. Third, ABCB1 (a drug efflux transporter) was not adopted because its functional-axis assignment is ambiguous and it cannot be assigned uniquely to either intestinal identity or the autonomous cell cycle (Principle 2). Note that, by the same principle of avoiding circularity, MDM2 is excluded from every signature in the analyses that correlate MDM2 promoter usage (P2_index) with signature scores (Section 3.2.2).

The P1 and P2 signature scores of each sample were calculated as gene set enrichment scores using GSVA [78]. The association of the GSVA scores with CMS subtype (four groups) was evaluated by the Kruskal–Wallis test, and their association with dMMR/pMMR (MSI status, two groups) by the Wilcoxon rank-sum test. Because both groups in the dMMR/pMMR comparison of the P2 signature (GSE39582; dMMR 75, pMMR 444) exceed 50, wilcox.test used the normal approximation (immune-excluded P2 score, the primary score in the main text: P = 6.59×10⁻¹⁰; full 29-gene score: P = 1.14×10⁻¹⁷) (at this sample size, the minimum attainable P for the Wilcoxon rank-sum test is 1.1×10⁻⁴³ under the normal approximation with continuity correction). In running GSVA, kcdf = “Gaussian” was specified for the kernel estimation, because the input consists of continuous log expression values. In addition, because GSVA does not score gene sets with fewer than two genes, modules consisting of a single gene were not scored (Section 3.3.2). Furthermore, in order to confirm whether the correspondence between the signatures and the CMS classification is supported at the gene level as well, we evaluated the overlap between the DEGs of this study (Supplementary Table S4; 1,013 genes on the tissue-1 side and 877 genes on the tissue-2 side) and CMS class-specific marker genes by the one-sided Fisher's exact test (background 26,278 genes; the 8 tests of 4 classes × 2 directions were adjusted by the BH procedure). Two kinds of reference sets were used independently. (i) The CMS templates of CMScaller[75] (the full template comprises 529 genes = CMS1 126, CMS2 82, CMS3 84, CMS4 237; of these, the present analysis was restricted to the genes present in the expression matrix, using CMS1 123 genes, CMS2 77 genes, CMS3 82 genes, and CMS4 224 genes); and (ii) the top 200 genes of each class obtained by contrasting each class against the rest with the Wilcoxon rank-sum test in the 574 CMS-classified aliquot rows (563 specimens) of TCGA-COAD/READ (of which those present in the expression matrix were CMS1 182 genes, CMS2 172 genes, CMS3 169 genes, and CMS4 182 genes). The complete gene lists of both reference sets, and the genes overlapping with the DEGs of this study, are given in Additional file 15: Supplementary Table S28. The convention of the “top 200” for the latter was fixed before the results were inspected, and we also confirmed that the same tendency was preserved with the top 100 and the top 500. Neither reference set was used in defining the DEGs or the signatures of this study, so the query side and the reference side are independent. The results are shown in Supplementary Table S28.

Survival analysis was performed on 591 TCGA-COAD/READ patients with overall survival (OS) information, after removal of duplicates at the patient level. For each signature score, patients were dichotomized into high and low groups at the median, OS was compared by the Kaplan–Meier method, and the between-group difference was evaluated by the log-rank test (the Kaplan–Meier analysis by CMS subtype was performed with n = 533). Furthermore, univariate Cox regression [79], and multivariate Cox proportional hazards regression with the P1 and P2 scores entered simultaneously, were used to estimate the independent prognostic value of each score as a hazard ratio (HR) with a 95% confidence interval. All tests were two-sided, and P < 0.05 was considered statistically significant. The series of external validations was performed in R, using TCGAbiolinks[42] and GEOquery[77] for data acquisition, org.Hs.eg.db for ID conversion, CMScaller[75] for CMS classification, GSVA[78] for signature scoring, survival and survminer for survival analysis, and ggplot2 [173] for plotting. The analysis code for this external validation is provided as Code8, Supplementary Methods.

In addition, in order to strengthen methodological robustness, we performed the following sensitivity analyses: (i) splitting the signatures into functional submodules — P1 (identity, proliferation) and P2 (metaplasia, p53 targets, immune, lncRNA) — and scoring them separately; (ii) re-evaluating the association with MSI using a version of the P2 signature from which the immune checkpoint genes were excluded (a check for circularity; this immune-excluded version is the primary P2 score for the CMS and MMR associations in the main text); (iii) using the TP53 mutation status of TCGA (Masked Somatic Mutation, MAF) as a positive-control axis (expected from the p53-target content of the P2 signature); and (iv) in the survival analysis, adjusted Cox regression with age, stage, and MSI status as covariates, together with likelihood ratio tests of the incremental prognostic value over the clinical model (with the C-index also reported). Tumor purity (ESTIMATE; the tumor-cell fraction estimated from expression data) was not included as a covariate in the main analysis, and a sensitivity analysis adding it was performed separately. Note that, in order to avoid circularity, a version of the P1 signature from which MSH2 was excluded was used throughout the survival analyses of Section 3.3.2 (both the Kaplan–Meier and the Cox proportional hazards main analyses). For CMS classification, CMScaller was used, with log2(TPM+1) given as input and RNAseq=TRUE specified. Because that option internally performs a log2 transformation (pseudocount 0.25) and quantile normalization, a log transformation is applied twice in this analysis (confirmed by inspection of the implementation). CMScaller's nearest template prediction is based on rank-based correlation and is designed to be little affected by monotonic transformations, and this robustness was demonstrated in a sensitivity analysis: the CMS label concordance between reclassification with corrected inputs (linear TPM + RNAseq=TRUE, and log2(TPM+1) + RNAseq=FALSE) and the reported assignments was 96.3% and 94.2%, respectively, and the main tests (the Kruskal–Wallis tests of the P1 and P2 signature scores across CMS classes) retained equal or greater significance with the corrected inputs (for example, P = 5.0×10⁻¹⁵ and 1.4×10⁻²⁹ in the linear-TPM version). The CMS assignments of this manuscript are therefore robust to the choice of input scale, and we retain the reported assignments (a single run under the settings of this section) as the primary values (the sensitivity analysis code is R46, Supplementary Methods). Note also that the classification of CMScaller is based on nearest template prediction, and its P values and FDRs are given by permutation. Therefore, for specimens lying at the FDR = 0.05 boundary, the assignment of “classifiable” versus “unclassified (NA)” can switch from run to run. The CMS breakdown reported in this manuscript is the result of a single run, obtained mechanically from the same cache as the score matrix used for the main text and the figures and tables.

The analyses described in Section 3.3.2 of this section (analysis code: Code8, Supplementary Methods) were run in R 4.6.0 (x86_64-apple-darwin20, macOS 13.7). The versions of the main packages were GSVA 2.6.3, CMScaller 2.0.1, TCGAbiolinks 2.40.0, SummarizedExperiment 1.42.0, org.Hs.eg.db 3.23.1, AnnotationDbi 1.74.0, GEOquery 2.80.0, survival 3.8-9, survminer 0.5.2, limma 3.68.4, matrixStats 1.5.0, data.table 1.18.4, ggplot2 4.0.3, and dplyr 1.2.1. Code8 automatically writes out the sessionInfo() output at the end of each run, and this output is included in the Supplementary Methods as Code8_sessionInfo.txt. For the analysis in Section 3.2.2 (analysis code: Code9) as well, the sessionInfo() output obtained on the same computer is included in the Supplementary Methods as Code9_v3b_sessionInfo.txt: R 4.6.0 (as above), with UCSCXenaTools 1.7.0, GSVA 2.6.3, CMScaller 2.0.1, TCGAbiolinks 2.40.0, survival 3.8-9, survminer 0.5.2, data.table 1.18.4, matrixStats 1.5.0, ggplot2 4.0.3, dplyr 1.2.1, org.Hs.eg.db 3.23.1, AnnotationDbi 1.74.0, and GEOquery 2.80.0. The R execution environment was common to all analysis code in this study, including the analyses in Sections 3.2.2, 3.2.4 and 3.3.3 (Code12–Code15) and the discovery-cohort analyses (Code3–Code7) (the same computer, R 4.6.0; the bundled sessionInfo() records — Code8 [also covering Code11, Code12, Code14, Code16, Code18, Code19 and R30], Code9, R38, R39, R47, R47b, R47c, R48 and R49 — are listed in Section 5 of the README of the Supplementary Methods).

For the direct quantification of MDM2 P1/P2 promoter usage, we used the transcript-level expression values of TCGA-COAD/READ (UCSC Xena TOIL [43,44] — the uniformly re-quantified RNA-seq expression distributed by the public cancer-genomics data portal UCSC Xena — TcgaTargetGtex_rsem_isoform_tpm, GENCODE v23 transcript models, log2(TPM+0.001)), converted back to the linear scale. Each MDM2 transcript was classified on the basis of its transcription start site (TSS): upstream forms beginning at exon 1 were assigned to P1, and forms beginning at exon 2 to P2. The boundary between the two promoters was set on the basis of the known gene structure in which the p53-responsive promoter P2 lies within intron 1 (immediately upstream of exon 2) [6,59] and SNP309 (rs2279744, IVS1+309) is contained in this P2 region [7] (hg38, chr12, + strand). Transcripts with downstream, non-canonical first exons that correspond to neither P1 nor P2 were excluded from the analysis. In the GENCODE v23 transcript models used in this analysis, 36 transcripts are annotated for MDM2, and by this rule they fall into 8 P1-derived transcripts (TSS 68,808,172–68,808,464), 23 P2-derived transcripts (68,809,017–68,809,227), and 5 transcripts with downstream non-canonical first exons (68,813,602 onward). Of the 23 P2-assigned transcripts, 6 begin in exon 2, the P2 first exon (68,809,017–68,809,024), and 17 are GENCODE models whose annotated 5′ end lies at the start of exon 3 (68,809,212–68,809,227); the latter contain neither exon 1 nor exon 2, and they are assigned to P2 because their transcription start sites fall within the P2 coordinate window used here (68,809,017–68,809,227). The 5 excluded transcripts were counted in neither the numerator nor the denominator of P2_index. The transcript-level assignment (36 transcripts) is reconstructed from the rule above and printed at run time by the deposited analysis code (Code9, Supplementary Methods). For each tumor, the P2 usage measure was defined as P2_index = ΣTPM(P2)/ΣTPM(P1+P2) (0–1). CMS was called with CMScaller, and the P1/P2 signature functional module scores were computed with GSVA. Because P2_index is computed from MDM2 transcripts, MDM2 was excluded from all signatures correlated with P2_index in order to avoid circularity. Associations were tested with the Kruskal–Wallis test (CMS), the Wilcoxon test (MSI, TP53 mutation status), Spearman correlation (module scores), and the Kaplan–Meier method and Cox proportional hazards models (overall survival). In addition, to assess the robustness of the observed associations, we performed a sensitivity analysis restricted to the cases for which covariates were available. Tumor purity was the ESTIMATE-derived approximate purity (higher values indicate higher tumor purity; median 0.826, range 0.283–0.984 in this cohort; the same values as in the signature analyses, Code17), and MDM2 copy number was obtained from the gene-level copy number of the Genomic Data Commons (ASCAT3 [81], hg38). The ASCAT3 copy-number files are organized in units of tumor and matched normal pairs, and their column names are given as composite barcodes joining the two with a semicolon (for example, TCGA-AZ-4313-10A;TCGA-AZ-4313-01A). The retrieval conditions were data.category = “Copy Number Variation”, data.type = “Gene Level Copy Number”, and workflow.type = “ASCAT3”. Because the order of tumor and normal is not fixed, we split the composite barcode at the semicolon, selected the barcode whose sample code was 01 (Primary Solid Tumor), and aggregated per specimen, obtaining 575 specimens (COAD 422, READ 153). Note that an implementation that does not split the composite barcode and refers only to the sample code of the first barcode misses the 276 columns in which the tumor is listed second (COAD 194, READ 82), yielding 299 specimens (575 = 299 + 276). The retrieval script, provided as Code19 (Supplementary Methods), was written for this study and for the companion study of the 15q11-q13 locus (T. Tsukui, R. Yao, and K. Tsuda, unpublished observations). A total copy number of 5 or more was defined as high-level amplification. Using a multivariable linear model with standardized P2_index as the response variable and TP53 status, CMS, tumor purity, and MDM2 copy number as explanatory variables, we estimated the independent contribution of each factor. MSI was excluded from the covariates of this model and was examined only in univariate comparisons (Section 3.2.2), because only 29 specimens had all of the other covariates of this model available (TP53 status, CMS, tumor purity) and only 27 when MDM2 copy number was added as well, which would severely compromise the sample size of a complete-case analysis. For the association with the p53 target module, we computed a partial correlation adjusted for tumor purity and CMS by residualizing the ranks on these covariates. The 95% confidence intervals of the Spearman correlations were obtained by Fisher's z transformation, and the Benjamini–Hochberg false discovery rate (FDR) correction was applied to the multiple association tests.

#### Association with the colibactin mutational signature (the Nunes cohort and TCGA-COAD/READ)

To explore the relationship between the P1/P2 signatures and exogenous mutagen exposure in colorectal cancer, we used an independent cohort with both WGS and RNA-seq (Nunes et al. 2024; U-CAN, n = 1,063) [82]. Expression was taken from the log₂(TPM+1) matrix of ArrayExpress E-MTAB-12862, the marks of colibactin exposure (SBS88 and ID18, mutation patterns attributed to the genotoxin colibactin of pks-positive Escherichia coli) from the per-sample COSMIC-decomposed activities in Supplementary Table 13 of that paper, and the clinical annotation from Supplementary Table 1; the 1,051 specimens for which both were available were analyzed. Taking as the axis the difference between the GSVA scores of Type1_full and Type5_full (kcdf = “Gaussian”; the same gene sets as in the first part of this section), we performed Spearman rank correlation between the axis and the activities (t approximation because of ties), Fisher's exact test of the positivity rate of the activities (activity > 0) in the upper and lower tertile groups of the axis (351 specimens each), and the Wilcoxon rank-sum test (normal approximation because both groups were ≥ 50). As sensitivity analyses, we performed the same analysis restricted to MSS and a partial Spearman correlation by the rank-residual method (covariates: MSI status, stage, site, age). Each family of 6 tests was corrected by the Benjamini–Hochberg procedure (analysis code: R42, Supplementary Methods). We further examined whether the same association was observed in TCGA-COAD/READ, using the per-sample activities obtained by fitting the deposited somatic mutation data (maf_coadread.rds; 616 specimens, GRCh38) to the COSMIC reference (the v3.1 series, which includes SBS88 and ID18) with sigminer [83] (sig_fit). Two axes were used, P2_index (isoform quantification; the first part of this section) and P1−P2 GSVA, and tests of the same type were performed on the 374 specimens that could be matched to the activities by the first 12 characters of the barcode, with each block of 12 tests treated as a family for correction. The correspondence between the directions of the two axes was not assumed a priori but was interpreted only after the correlation between them had been measured (Spearman ρ = −0.156 between P2_index and the P1−P2 axis; analysis code: R43, Supplementary Methods). The clinical annotation (MSI, stage, site, age) was completed from the per-patient annotation derived from the GDC (analysis code: R44, Supplementary Methods).

The results were as follows. In the Nunes cohort the colibactin marks were slightly denser on the P2-dominant side of the axis (SBS88 Spearman rho = -0.085, P = 0.0059; ID18 rho = -0.104, P = 6.9x10-4; positivity in the P1-high versus P2-high tertiles 4 of 351 versus 16 of 351 for SBS88 and 20 of 351 versus 38 of 351 for ID18), the direction was retained when restricted to MSS, and partial correlation adjusted for MSI status, stage, site and age left ID18 significant (rho = -0.084, P = 0.0067) with SBS88 borderline (rho = -0.060, P = 0.053); all positive specimens were MSS (Supplementary Table S42, Supplementary Figure S11). In TCGA-COAD/READ only P2_index versus ID18 reproduced the direction (rho = +0.143, P = 0.0055, FDR = 0.044; Supplementary Figure S12). We then asked whether this shift runs through p53 activity, using the TP53 status, p53 activity score and hypermutation state of the 374 TCGA specimens (analysis R168, Supplementary Methods; all values in Supplementary Table S48, sheet P). It does not. The activities were unrelated to the p53 activity score in TP53 wild-type tumors (132 tumors: SBS88 rho = -0.059, P = 0.50; ID18 rho = -0.010, P = 0.91) and in the 91 non-hypermutated wild-type tumors that form the main set of Section 3.2.3 (rho = -0.088, P = 0.41; rho = -0.001, P = 0.99), and no adjustment (hypermutation, stromal score, log2 total MDM2) brought a relationship out. Dividing the wild-type tumors into those with and without evidence of p53 activity left the positivity rate unchanged (SBS88 3 of 57 versus 6 of 75, Fisher P = 0.73; ID18 7 of 57 versus 6 of 75, P = 0.56). The association of P2_index with ID18 was present in TP53-mutant tumors (rho = +0.209, P = 0.0011, 242 tumors) and absent in wild-type tumors (rho = +0.026, P = 0.77); adjusting the whole cohort for TP53 status left it at rho = +0.127 (P = 0.014), so the two readings are reported together. SBS88 positivity was in turn more common in TP53-mutant than wild-type tumors (42 of 242 versus 9 of 132, odds ratio 2.87, P = 0.0043) and uncommon in hypermutated tumors (2 of 59 versus 49 of 315, odds ratio 0.19, P = 0.012), which is consistent with the reported near-exclusivity of colibactin exposure and mismatch-repair deficiency. In the Nunes cohort the p53-target score itself was only weakly related to the marks in MSS specimens (SBS88 rho = +0.076, P = 0.028; ID18 rho = +0.095, P = 0.0063) and the relationship disappeared after the same covariate adjustment (rho = +0.021 and +0.023, P >= 0.51), while the metaplasia score was unrelated throughout (|rho| <= 0.04, P >= 0.23). Only 9 and 13 of the 132 wild-type specimens carry any activity, so these are statements about what was not seen at this size rather than evidence of no relationship.

#### Analysis of MDM2 inhibitor sensitivity, MDM2 gene dependency, and drug sensitivity prediction in cell line panels

To examine in independent pharmacogenomic data, indirectly (using TP53 status as a surrogate rather than promoter usage; MSI status was used only for background stratification, as MSI-H is not a proxy for P2, Section 3.2.2), the therapeutic implications of the P2-dominant group (wild-type TP53 with active p53; MSI-like only at the signature level) proposed in this study, we used the dose–response data (fitted_dose_response) and the cell line annotation (Cell_Lines_Details) of GDSC1 and GDSC2 from the Genomics of Drug Sensitivity in Cancer [49,50]. The drugs to be evaluated were identified by automatically matching all drug names in both databases (378 drugs in GDSC1 and 286 drugs in GDSC2) against a list of 13 known MDM2–p53 pathway drug names (including aliases) specified in the code. Four drugs matched: the MDM2–p53 interaction inhibitors nutlin-3a and Serdemetan, and the mutant p53 reactivators PRIMA-1MET and MIRA-1. Other MDM2–p53 pathway drugs for which clinical development has been attempted (RG7112, Idasanutlin (RG7388), AMG-232 (= navtemadlin/KRT-232), Milademetan (DS-3032b), HDM201 (siremadlin), SAR405838, and APG-115 (alrizomadlin)) were not listed in either database. LN_IC50 (smaller values indicate higher sensitivity) was used as the sensitivity measure, and when multiple curves were available for the same database × drug × cell line combination, they were aggregated to the median before testing.

The TP53 mutation status of the cell lines was determined from Model.csv and OmicsSomaticMutations.csv of the DepMap 24Q4 Public release [52,53] (DepMap is a public resource of cell-line expression, mutation and gene-dependency data) and joined to the GDSC cell lines via COSMIC ID. The determination was based on the VariantInfo column (not the VariantType column, which indicates the type of variant): cell lines carrying a non-synonymous variant were classified as mutant, and, among the cell lines for which a somatic mutation profile was available, those carrying no non-synonymous variant were classified as wild type. Cell lines for which no profile was available were not defaulted to wild type but were excluded from the analysis. MSI status was consolidated into two groups from the cell line annotation, with MSI-H and MSI treated as the MSI-High group and MSS and MSI-L as the MSS/MSI-L group. The analyses of colorectal cancer cell lines were restricted to the 47 lines whose lineage annotation was Colorectal or Bowel and whose TP53 status could be determined, and the analyses stratified by MSI were performed on the same set. Group comparisons used the Mann–Whitney U (Wilcoxon rank-sum) test, effect sizes used the Hodges–Lehmann estimator and its 95% confidence interval, and the Benjamini–Hochberg FDR correction was applied within each analysis block. A measure integrating multiple drugs was created by z-scoring LN_IC50 for each database × drug and taking the mean of these values for each cell line. The integration was restricted to the two MDM2 inhibitors that share a mechanism of action; the two reactivators, whose mechanism of action is the opposite, were not included. This measure is the mean of multiple measurements obtained for the same cell line and is not an integration of independent tests.

For genetic dependency, we used CRISPRGeneEffect.csv from the same DepMap 24Q4 Public release [52,53]. The gene effect of MDM2 (smaller values indicate greater dependency) was compared by TP53 status determined by the same rules as above. The analysis covered the 1,178 cell lines for which both the gene effect and TP53 status were available. The handling of tests and effect sizes was identical to that in the drug sensitivity analysis. Furthermore, to confirm whether the same direction is obtained in patient-derived models, away from cell lines that have undergone adaptation to culture, we independently re-analyzed publicly available drug sensitivity data for colorectal cancer organoids (analysis code: R38, Supplementary Methods). We used the primary data released by the authors of reference [13] on Figshare (doi:10.6084/m9.figshare.28339340); of the 256 organoid lines (as reported by the authors: 132 colorectal, 76 esophageal, 22 pancreatic, 20 ovarian, and 6 gastric), we analyzed the 65 colorectal organoids for which both the log(IC50) of the MDM2 inhibitor nutlin-3 and somatic mutation data were available. For log(IC50), smaller values indicate higher sensitivity. TP53 status was based primarily on TP53_mut in the mutation annotation table (48 mutant lines / 17 wild-type lines), and a split by TP53_LOF, which regards only truncating variants as mutant, was reported alongside it as a sensitivity analysis. Lines not appearing in the mutation annotation table were, by rule, excluded from the analysis rather than defaulted to wild type, but no line fell into this category. The group comparison in the main analysis (17 wild-type lines versus 48 mutant lines) used the exact Wilcoxon rank-sum test without ties. In contrast, in the sensitivity analysis by TP53_LOF one group comprised 51 lines, which falls outside the range in which the implementation used selects the exact test (both groups below 50), so that value is based on a normal approximation with continuity correction. Cliff's delta was used as the effect size, and the robustness of the result was confirmed with linear models that successively added KRAS mutation, MSI status, and MDM2 gain-of-function mutation. Signature scores were computed with GSVA (version 2.6.3) from the same expression matrix (255 lines × 15,764 genes), and the 13 sets that remained after excluding sets with fewer than 2 covered genes (which GSVA cannot score) were analyzed. Correlations were evaluated with the exact Spearman test without ties (exact Spearman, AS 89), and the Benjamini–Hochberg correction was applied treating the 13 sets as a family. Partial correlations adjusted for TP53 status were computed as the correlation between the residuals obtained by converting both variables to ranks and regressing out TP53 status. Note that the lncRNA module of P2 (2 genes) had zero coverage in this expression matrix, so that Type5_noimmune, which excludes the immune module, becomes the same gene set as the metaplasia plus p53 target core (Type5_core), and their ρ and P values coincide exactly. Therefore, of the 13 sets only 12 tests are independent, but even if the correction is redone with a family of 12, not a single significance call changes.

Furthermore, in order to evaluate drug sensitivity without preselecting the drugs of interest, we used oncoPredict (version 1.3.1) [57], which projects drug sensitivity from cell line panels onto patient specimens. For training we used 805 cell lines × 198 drugs from GDSC2 (the expression matrix RMA-normalized and log-transformed; the response matrix with IC50 returned to the linear scale), and as the projection target we used the expression matrix of 624 TCGA-COAD/READ specimens (aggregated per specimen by the same convention as in Section 3.3.2). Batch correction was performed with ComBat [60] (empirical Bayes), low-variance genes were removed as the bottom 20% on the raw data, the minimum number of training specimens per drug was 10, and a Box-Cox transformation was applied to the drug response on the training side (powerTransformPhenotype = TRUE). Associations between the predicted values and each subtype score were evaluated with Spearman rank correlation, and the Benjamini–Hochberg correction was applied over the 198 drugs. The computation was performed in 10 runs (nine runs of 20 drugs and one of 18), but the maximum absolute difference from the result of computing a single drug alone was 0, so the splitting did not affect the results.

To test drug associations on measured rather than predicted data, P1/P2 GSVA scores were also computed from GDSC2 cell-line expression (RMA) and related to measured GDSC2 LN_IC50 for all drug entries and to DepMap 24Q4 CRISPR gene effects of 17 target genes, by Spearman correlation and by rank-based partial correlation adjusted for cancer type (lineage) and TP53 status (R47/R47b) and, as a sensitivity analysis, additionally for RAS and BRAF mutation status (R47c; Supplementary Methods). Drug classes (the ATR/CHK1/WEE1, EGFR and MEK/ERK/BRAF inhibitors) were compared with the remaining drug entries (excluding these classes and nutlin-3a) by the two-sided Wilcoxon rank-sum test (exact where applicable) on the per-drug partial correlation coefficients (R47d; Supplementary Methods).

For the analysis of the immune microenvironment of P2 and the prediction of response to immune checkpoint inhibitors (ICIs), we used the 624 per-specimen TCGA-COAD/READ specimens (analysis code: Code13, Supplementary Methods). Stratification was by tertiles of the P2 signature score; the top 208 and bottom 208 specimens were compared as groups, the middle tertile was excluded from the group comparison, and correlations with the continuous axis were computed on all 624 specimens. Immune cell fractions were estimated with quanTIseq [61] and MCP-counter [62] of immunedeconv [63] (version 2.1.4), and only the cell types whose signs agreed between the two methods were treated as findings. xCell and EPIC failed to run and were therefore not used. Group comparisons used the Mann–Whitney test (normal approximation, continuity correction, tie correction), effect sizes used the Hodges–Lehmann estimator, and multiplicity was corrected within each method by the Benjamini–Hochberg procedure. To confirm that this was not a restatement of MSI status, we used three approaches in the 272 specimens for which MSI annotation was available: a van Elteren test stratified by MSI, a test restricted to MSS/MSI-Low, and a partial Spearman correlation controlling for an indicator variable of MSI-High. For ICI response prediction we used TIDEpy (version 1.3.8), a Python implementation provided by the original authors; the group comparison of the predicted Responder rate used Fisher's exact test, and the adjustment for MSI status used logistic regression.

For the analysis by TP53 functional class, we used the mutation annotation of the TP53 Database (formerly the IARC TP53 Database; currently maintained by the U.S. National Cancer Institute; release 21) [66,67] (analysis code: Code14, Supplementary Methods). The targets were the 374 TCGA-COAD/READ specimens for which both P2_index and TP53 mutation information were available (from the 380 specimens for which P2_index was available, the 6 specimens for which somatic mutation data were not available were removed by the same rule as in Section 3.2.2). The classification rules were declared before the analysis: wild type (WT) was defined as specimens in which no somatic TP53 mutation was detected in the Masked Somatic Mutation data, and loss-of-function (LOF) as truncating variants such as nonsense, frameshift, and splice-site mutations. Because no single operational definition of gain-of-function (GOF) is established in the literature, two definitions were used in parallel: definition A designated as GOF only missense variants at the 6 hotspot residues (R175H, G245S, R248Q/W, R249S, R273H/C, R282W) [68], and definition B designated as GOF all missense variants whose transactivation class in the TP53 Database was non-functional [69]. Missense variants that did not meet the GOF criteria were excluded from the three-class comparison. The comparison among the three classes used the Kruskal–Wallis test, and the three prespecified contrasts used the Mann–Whitney test and the Hodges–Lehmann estimator, with the Benjamini–Hochberg correction applied. P values that the analysis software returned as 0 because of underflow were recomputed from the per-specimen data and checked for consistency against the minimum attainable P at that sample size.

Note that ‘tumor purity’ in this manuscript denotes a single measure except in one analysis: in the external cohort validation (Code8) we used the value obtained by converting the ESTIMATE score into an approximate purity with the transformation of Yoshihara 2013 [80] (purity = cos(0.6049872018 + 0.0001467884 × ESTIMATE score)), and the multivariable model of the direct validation of MDM2 P1/P2 (Code21, with the covariate table of R16) used the same approximate purity (median 0.826, range 0.283–0.984; identical for all 624 specimens); only the purity-adjusted analyses of Section 3.2.3 of the main text used ABSOLUTE purity (inferred from copy-number data; R93). This transformation was calibrated on Affymetrix arrays, and its application to RNA-seq is an approximation. Higher values of this approximate purity indicate higher tumor purity. The purity-adjusted Cox regression in Section 3.3.2 also uses this approximate purity as a covariate, and wherever this manuscript writes ‘tumor purity (ESTIMATE)’ it refers to this approximate purity. The purity values are computed by Code17, which was written for this study and for the companion study of the 15q11-q13 locus (T. Tsukui, R. Yao, and K. Tsuda, unpublished observations). The approximate purity and the ESTIMATE score itself run in opposite directions: because the ESTIMATE score reflects the amount of stromal and immune components, higher values of that score indicate lower tumor purity. Because the approximate purity is cos(0.6049872018 + 0.0001467884 × ESTIMATE score), a strictly decreasing function of the ESTIMATE score over the observed range (the argument stays between 0 and π/2; values that would be negative are set to missing; P2_step0_recover_ESTIMATE_purity_v1.R), the two measures have a Spearman rank correlation of exactly −1 per aliquot, and specimen-level values, which are means over aliquots, are almost perfectly inversely ranked. Rank-based partial correlations therefore give the same results with either measure, but caution is required in that scale-dependent estimates such as Cox regression reverse direction.

#### p53 activity score

Activity of wild-type p53 was scored as the mean z-score of 12 direct p53 targets (CDKN1A, BAX, BBC3, TP53I3, SESN1, RRM2B, GDF15, FDXR, TIGAR, PHLDA3, SPATA18 and EDA2R) [37] (p53 activity score); MDM2 itself and three p53 targets that ranked high in the genome-wide exon-usage screen (FAS, ZMAT3 and TNFRSF10C) were not included, to limit circularity with the P2 index; five of the 12 scored genes (CDKN1A, BAX, BBC3, GDF15 and SPATA18) are shared with the P2 signature (Section 3.2.1 of the main text). Genes were z-scored across the 63 discovery specimens for organoids and within each cohort for TCGA and cell lines (10 of the 12 genes were available in the GDSC cell-line data). TP53 wild-type samples without p53 activity were defined as those with a p53 activity score at or below the 90th percentile of mutant samples. The array P2 index (organoid HTA2.0 arrays) is the mean of the exon-2 minus the mean of the exon-1 probe sets, each relative to the MDM2 transcript cluster (a transcript cluster is the gene-level group of probe sets on the HTA2.0 array; RMA; defined in Section 3.1.2), a measure on a different scale from the transcript-level ratio used for TCGA (P2_index) and, with the same formula, for cell lines (cell-line P2 index).

#### RNA-seq of nutlin-treated cells

From the bigWig files (indexed genome-wide coverage tracks) of GSE86221 (a SubSeries of GSE86222; HCT116 TP53 wild type and null, MCF7 and SJSA; DMSO or nutlin 10 µM for 12 h; Ion Torrent) [38], the region around MDM2 (hg19, chr12:69.1–69.3 Mb) was extracted, and the mean coverage of the P1-only exon 1 (69,201,956–69,202,271), the P2-only first exon (69,202,797–69,202,866), the shared ATG exon and the last exon was computed. Boundaries were checked against the Ensembl GRCh37 annotation and steps in coverage. Because the bigWig files are the sum of two replicates, the P2/P1 ratio was the primary measure, and induction of MDM2 and CDKN1A was confirmed with the gene-level count tables (two replicates per condition; R82).

#### p53 activity, protein levels and junction reads in TCGA tumors

In TCGA-COAD/READ, the p53 activity score was computed from TCGAbiolinks expression [42] and, as a sensitivity analysis, from UCSC Xena Toil gene TPM [43,44]. TP53 status was taken from the GDC masked somatic mutation calls (a tumor with any TP53 variant other than silent, intronic, untranslated-region or RNA-gene classes was counted as mutant, and tumors absent from the calls were excluded rather than counted as wild type; R16), and in the main analyses wild-type status was also confirmed in the MC3 mutation calls [45] (the TCGA pan-cancer consensus somatic-mutation call set). Hypermutation was defined as ≥ 400 non-synonymous mutations. Protein levels of BAX, p21 and p53 were taken from the TCGA Pan-Cancer Atlas reverse-phase protein array (RPPA; antibody-based protein quantification) data (UCSC Xena [43]); in the 69 non-hypermutated, MC3-confirmed wild-type tumors with RPPA data, BAX protein correlated with the p53 activity score (ρ = 0.48), P2_index (ρ = 0.33) and total MDM2 (ρ = 0.39), and its association with P2_index did not persist after adjustment for total MDM2 (partial ρ = 0.09, P = 0.46), as for the mRNA-based score (Additional file 7: Supplementary Table S48, sheet E; code r120_S243_paper1_rppa_v1.py, Supplementary Methods). MDM2 junction reads were counted from recount3 [46] (uniformly reprocessed public RNA-seq, which provides exon–exon junction counts) for the TCGA-COAD/READ tumors: reads joining exon 1 (P1) or exon 2 (the P2 first exon) to exon 3, with the P2 share = P2 / (P1 + P2) (R117d; Supplementary Table S48, sheet D).

#### P2 index of cell lines and drug response

The P2 index of cell lines was computed from DepMap 24Q2 transcript TPM [47,48] with the same transcription-start-site assignment as for TCGA, restricted to lines with sufficient MDM2 expression (TPM ≥ 5; R74). TP53 status was taken from GDSC (COSMIC) for analyses with GDSC (691 lines, 208 wild type) and from the DepMap mutation table (presence of protein-altering mutations) for the CRISPR and PRISM analyses (364 wild-type lines; the two definitions agreed for 686 of 691 lines). Drug response used nutlin-3a ln IC50 from GDSC1 and GDSC2 [49,50] and the PRISM secondary (19Q4) and primary (24Q2) screens [51] (pooled drug screens of barcoded cell lines); MDM2 dependency used DepMap CRISPR gene effects (Chronos scores, more negative meaning stronger dependency; DepMap 24Q4) [52–54]; rank partial correlations adjusted for the p53 activity score were also computed (R70 and R76). Nutlin-3 sensitivity (AUC) in CTRPv2 [55,56] (taken from the data files distributed with oncoPredict [57]) was analyzed in the same way (R93). MDM2 junction reads were also counted directly in the SG-NEx cell lines, which were sequenced with short reads and with Nanopore long reads (direct RNA, direct cDNA and PCR cDNA, including strand-specific PCR cDNA) [58]: only the MDM2 region of the public genome alignments was read, reads joining exon 1 (P1) or exon 2 (the P2 first exon) to exon 3 were counted as for TCGA (Section 2, p53 activity, protein levels and junction reads in TCGA tumors), and the six lines with 20 or more junction reads and a DepMap P2 index were compared (R137 and R142).

#### Genome-wide survey of morphology and TCGA histology

For the 39,438 sufficiently expressed transcript clusters, rank partial correlations with the non-Type1 fraction and with the P2 index (adjusted for scan period: the month of array scanning taken from the dates of the CEL files, six categories, as a proxy for processing batch) were computed, and clusters with FDR < 0.05 and the same direction in patient means were retained (patient means were computed for 22 patients, with HCT83-4LM [registered under patient number HCT84] counted as a separate patient; with HCT83-4LM assigned to HCT83 [21 patients], the 124 clusters associated with morphology only [44 on 20q] were unchanged, the clusters associated with the MDM2 axis only changed from 930 to 919, and the same arms were enriched). Exon usage was screened genome-wide in the same arrays: each exon (PSR) and junction (JUC) probe set of genes with a median log2 expression of at least 5 was expressed relative to its transcript cluster (probe-set minus transcript-cluster RMA value), keeping probe sets with a median of at least 4 (in either group for the comparison of promoter groups). In the genome-wide exon-usage screen, P2-dominant and P1-dominant specimens were compared on patient-by-group means (Welch's t test; Benjamini–Hochberg FDR < 0.05 and a difference of at least 1 log2 unit), with the same direction at FDR < 0.05 required in a specimen-level linear model adjusted for scan period (R68, r68_screen_v1.py; the probe sets referred to under p53 activity score and in Section 3.2.5 of the main text); in the survey of Section 3.1.4 of the main text, the same relative values (MDM2 excluded) were related to the non-Type1 fraction and to the P2 index with the covariate and criteria above (R77). Transcript clusters were assigned to chromosome arms by centromere position; arm-level enrichment was tested by one-sided Fisher's exact tests against all expressed clusters (42 arms; Benjamini–Hochberg), and arm-level mean expression was used as a proxy for copy number. Genes on the cystic–mucinous and compact glandular sides were tested for bias toward ten prespecified cytoskeleton and adhesion GO terms by one-sided Fisher's exact tests. The morphology gene score was computed by z-scoring, in primary TCGA-COAD/READ tumors (Toil gene TPM; TPM ≥ 1 in at least half of tumors), the genes of the 141 transcript clusters associated with morphology (morphology only, or morphology and the MDM2 axis in the same direction) and subtracting the mean of the genes higher on the compact glandular side from that of the genes higher on the cystic–mucinous side. Histology was taken from the histological_type field of the TCGA Pan-Cancer clinical table (TCGA-CDR) [32]: tumors recorded as colon or rectal mucinous adenocarcinoma (43; the pathological diagnosis recorded by TCGA, not re-assessed here) were counted as mucinous carcinoma, and the three tumors recorded as discrepant were excluded (372 tumors; r101.py, r101c.py). Molecular subtypes were taken from the TCGA molecular subtype table of UCSC Xena [43] (TCGASubtype.20170308) [33]. In this table the MSI (HM-indel) subtype is one category, and it was used as the MSI covariate. Whether morphology and expression programs follow TP53 status or the P2 measure was tested with rank linear models (partial F test for the added term, which asks whether that term improves the fit) in the discovery organoids (56 specimens with determined TP53 status), the GSE294511 organoids with published TP53 status (88) and TCGA-COAD/READ primary tumors with MC3 TP53 calls (358), with 20q and aneuploidy (arm-level expression proxies: within each organoid cohort, an arm score was the mean z-score of the expressed transcript clusters [median log2 RMA ≥ 5] on the arm minus the sample's genome-wide mean, over the 39 arms with at least 40 clusters and with the clusters of the p53-score genes and of 30 known p53 targets excluded; 20q was the 20q arm score and the aneuploidy index the standard deviation of a sample's 39 arm scores [r144_organoid_arm_p53_v1.py]; in TCGA, 20q was the mean z-score of the expressed 20q genes) or 20q and MSI (335 tumors with a molecular subtype) as covariates; in a sensitivity analysis in TCGA (TP53 calls of Supplementary Table S47; 327 tumors), BRAF mutation (any non-silent BRAF mutation in the MC3 calls, not restricted to V600E; 49 of 352 tumors with MC3 data) and the CpG island methylator phenotype (CIMP; widespread hypermethylation of CpG islands) status were added (called by us because the subtype table gives no CIMP call for CRC; CIMP-high when, for three or more of the five markers of the panel of Weisenberger et al. [35] (CACNA1G, IGF2, NEUROG1, RUNX3 and SOCS1), the median β value of the 450K DNA-methylation array probes in the marker's CpG island exceeded 0.30; the CIMP calls were computed by this rule from the GDC 450K β values for this study and the companion analysis of the 15q11-q13 locus [T. Tsukui, R. Yao, and K. Tsuda, unpublished observations], and the call table is included in the code archive as R152_TCGA_CIMP_calls.csv), and tumors that were not of the MSI subtype, BRAF wild-type and CIMP-low were compared by Mann–Whitney tests and with adjustment for 20q; program scores were the mean within-cohort z-scores of previously defined gene sets (the P1 and P2 signature components, the p53 activity score, a nine-gene gastric marker set chosen by the authors and fixed in the discovery cohort before the GSE294511 analysis — the gastric genes of the P2 signature with TFF2 and VSIG1 in place of TFF3 and MUC5B (CTSE, TFF1, TFF2, MUC5AC, MUC6, ANXA10, REN, MUC17 and VSIG1) — and the SNORD116 cluster, scored over the 25 HTA2.0 transcript clusters annotated as SNORD116; in TCGA only genes expressed in at least half of tumors were used, which left seven of the 11 metaplasia genes and four of the nine gastric genes). The metaplasia genes of the P2 signature are its nine gastric-type genes (CTSE, REN, TFF1, TFF3, MUC5B, MUC5AC, MUC6, MUC17 and ANXA10) and the Paneth-cell genes DEFA5 and DEFA6; gastric metaplasia in this study denotes the expression of these gastric-type genes and was not assessed histologically or by immunostaining. In the discovery organoids, the genome-wide comparison of TP53 status and promoter usage used gene-level values (transcript clusters mapped to gene symbols and averaged; 19,676 genes with log2 expression of at least 4 in at least one of the 56 specimens with determined TP53 status); each gene was regressed on TP53 status, on promoter usage and on both, with cluster-robust standard errors by patient (which allow for correlation among specimens of the same patient) and Benjamini–Hochberg correction across genes. These analyses were added after the per-specimen TP53 calls had been finalized (post hoc), and the program analysis, the CIMP sensitivity analysis and the genome-wide comparison were reproduced by independent re-implementations (Supplementary Table S48, sheet H).

#### Independent organoid biobank (GSE294511)

For replication in independent patients, we used the public HTA2.0 arrays of a biobank of organoids from colorectal liver metastases established at another institution (GSE294511; 119 organoids from 50 patients and 50 matched tumor tissues) [36], together with the per-organoid TP53 status, H&E-based morphology (cystic, mixed or solid) and drug sensitivity scores (DSS; higher = more sensitive; idasanutlin and the 23 other drugs common to both drug libraries of that study) published with it. CEL files were processed as for the discovery cohort (oligo RMA; organoids and tissues normalized separately), and the array P2 index (Section 3.1.2 of the main text), the p53 activity score and a morphology gene score were computed with definitions fixed in the discovery cohort. The morphology gene score is the mean z-score of the 44 transcript clusters higher on the non-Type1 side (not the 44 morphology-only clusters on 20q) minus that of the 97 higher on the Type1 side, among the 141 clusters associated with morphology in the genome-wide survey (morphology only, or both axes in the same direction; Section 3.1.4 of the main text). Total MDM2 was the MDM2 transcript cluster (TC12000606.hg.1). Because 17 of the 141 clusters were also associated with the MDM2 isoform, the score was recomputed from the 124 clusters associated with morphology only, and because the organoids come from fewer patients, the linear models of drug sensitivity were refitted with standard errors clustered by patient and the partial correlations repeated on patient means (sensitivity analyses added after the prespecified analyses). The analysis plan, including the primary test (one-sided Spearman correlation between the morphology gene score and the P2 index, with patient-mean and patient random-intercept analyses, and a within-patient comparison in patients whose H&E morphology differed between organoids), was fixed before the data were examined, and the key values were recomputed independently from the raw files. The values are given in Supplementary Table S48 (sheet F).

#### Statistical analysis

All tests were two-sided unless stated otherwise (the Fisher's exact tests of DEG–CMS marker overlap, chromosome-arm enrichment and GO-term bias, the patient-level permutation test and the primary test in the independent organoid biobank were one-sided). Group comparisons used the Wilcoxon rank-sum (Mann–Whitney) test or, for more than two groups, the Kruskal–Wallis test; correlations used Spearman's ρ, with rank-based partial correlation where covariates were adjusted for; and multiple testing was controlled by the Benjamini–Hochberg procedure within each family of tests. The specimen was the primary unit of analysis, because specimens from the same patient could differ in morphology and MDM2 isoform (HCT38, HCT41 and HCT67) and even in TP53 status (HCT67); because several specimens of the discovery cohort derive from the same patient, the morphology–isoform comparison was also evaluated at the patient level as a sensitivity analysis, with a linear mixed model of the Type1 fraction and a binomial mixed model of image counts, each with a patient random intercept (likelihood-ratio tests; R package lme4 [84]), and with a permutation test in which isoform labels were permuted between patients (R48, Supplementary Methods); in the independent organoid biobank (Sections 3.1.5 and 3.2.5 of the main text), linear models with a patient random intercept were fitted by maximum likelihood in Python, and standard errors clustered by patient (CR1) were used where stated. Analyses were performed in R 4.6.0 and Python; package versions are recorded in the sessionInfo() files in Supplementary Methods (Additional file 2). Script identifiers cited in this manuscript (CodeN and RN) refer to the files indexed in the README of Supplementary Methods.

#### Use of large language models

The use of a large language model in the preparation of this study is disclosed in the Methods section of the article (Use of large language models); the statement is not repeated here.

### R-3. Full-length version of main text Section 3.1.1 (Performance of deep-learning morphological classification)

#### Performance of deep-learning morphological classification of organoids (Figure 1, Table 1)

We first define the terms used from this section onward. “Type1 (compact glandular type),” “Type5 (dispersed mucinous type),” and the reference category “Type0” are the morphological classes that the deep-learning classifier assigns to individual organoid images. “Tissue 1” and “tissue 2,” by contrast, denote a contrast drawn at the group level between the MDM2 P1-dominant and the P2-dominant specimen groups (Section 2). We therefore use Type0/Type1/Type5 for the image-level morphological classes, P1 signature/P2 signature (P1 score/P2 score) for the expression signatures and their scores, and tissue 1/tissue 2 for between-group comparisons and biological interpretation. The P1 and P2 signatures are gene sets defined from the expression comparisons between the tissue-1-side and tissue-2-side sample groups (Section 2); they are written with the identifiers Type1_* and Type5_* in the deposited code and primary outputs (for example, the full P2 signature = Type5_full). The morphological classes other than Type1 (Type5 + Type0) are collectively called the cystic–mucinous “round morphology” (Section 3.1.2). The quantitative morphological measures “Type1 fraction,” “Type0 fraction,” and “Type5 fraction” are, for each specimen, the proportion of its images that the classifier assigned to that morphological class (Section 2), and the “non-Type1 fraction” is the sum of the Type0 and Type5 fractions (that is, the fraction of round morphology). “Type1-dominant” and “non-Type1-dominant” describe the relative magnitude of these fractions and are not per-specimen majority-vote labels. The notations in the supplementary tables and figures (Type1 fraction, non-Type1 fraction, Type0/Type1/Type5 fractions) refer to these same morphology-class fractions, which are quantities distinct from the P1/P2 expression signatures. The correspondence between the morphological classes and tissue 1/tissue 2 is not perfect (Section 3.1.2). The Type numbers are inherited from the serial numbers that the experimenters assigned when sorting morphologies visually in the morphological typing of Okamoto et al. (2022) [12], which analyzed a largely overlapping patient group from the same biobank. This is why the numbers not used in this study (Type2-4) are absent; neither the magnitude nor the continuity of the numbers carries any meaning.

##### Architecture of the classifier and the training and test sets

The architecture, training conditions, and input-image preprocessing of the VGG16 transfer-learning classifier are described in Section 2 (image classification by deep learning). Here we report the performance of that classifier on a test set that was not used for training. The training conditions, established from the training scripts and training logs themselves, were as follows. VGG16 was initialized with ImageNet-pretrained weights, with the fully connected part replaced by Flatten-Dense 256 (ReLU)-Dense 3 (softmax) and all layers trainable. Input images were 224×224 pixels, with pixel values normalized by division by 255. Optimization used SGD (learning rate 0.001) with a batch size of 5 for up to 100 epochs; 30% of the training set was held out as a validation split, and training was stopped by EarlyStopping (patience 7). No data augmentation was applied, and the random seed was fixed only for the training/test split, not for the training itself. In the surviving training logs, several runs with different early-stopping epochs all reached a test accuracy of 64/65 or higher (of the 5 surviving execution records, 4 gave 64/65 and 1 gave 65/65). The complete training and inference scripts are provided as Code1 (Supplementary Methods).

##### Classification performance and confusion matrix (Table 1)

Table 1 gives the overall accuracy on the test set (65 images), the class-wise sensitivity (recall), precision, and F1 score, together with the confusion matrix for the 3 classes (Type0, Type1, and Type5) as observed class × predicted class. Only 1 image was misclassified (an observed Type0 predicted as Type1), and misclassification between the two tumor-side classes (Type1 and Type5) was 0. The values in Table 1 were tabulated mechanically from the predicted and true labels written out in the same run as the training (the output of Code1).

##### Design of the training/test split (feasibility of a patient-level split)

This cohort consists of 63 specimens derived from 21 patients, and individual patients contributed multiple specimens and multiple images. The training/test split was a random split at the image level (Code1); no blocking at the specimen or patient level was performed. Images from the same specimen and the same patient can therefore appear in both the training set and the test set, so the test accuracy may be optimistically biased by within-patient correlation. We have not evaluated the accuracy under a re-split at the patient level, and the external validity of this classifier (generalization across institutions and patients) is unverified (Section 4.6).

##### Application to 792 images from 63 specimens and per-specimen morphology fractions

We applied the trained classifier to 792 organoid images from the 63 specimens and, from the predicted class of each image (the argmax of the softmax output), calculated the Type0/Type1/Type5 fractions for each specimen, defined as the proportion of images predicted as that class (Supplementary Table S2). The input images were normalized by dividing pixel values by 255, as in training (the initial application had not applied this normalization; the course of events and the comparison of the two conditions are given in Sections 3.1.1 and 3.1.2). The per-specimen fractions are given in Supplementary Table S2. The 194 images used for training and evaluation were chosen as images with highly reliable manual labels. Of these, 49 derive from 9 of the 63 specimens analyzed here and therefore overlap with the 792 images (established by exhaustive matching of file names; 6.2% of the 792 images). The remaining 145 images derive from specimens outside these 63, including 81 images of organoids derived from normal tissue, and do not overlap. Claims about classification performance are confined to the evaluation on the held-out test set (Section 3.1.1), which is unaffected by this overlap.

##### Limitations of this classifier and reporting standards

This classifier has seven limitations. First, the images used for training and testing derive from a single institution and a single imaging condition, and we have not validated the classifier on an external dataset. Second, the morphological labels were assigned manually by the authors, and we have not evaluated independent inter-observer agreement (for example, Cohen's kappa). Third, the test set is small (65 images), and the 95% confidence interval for the overall accuracy of 98.5% is correspondingly wide (Wilson method, 91.8–99.7%; note to Table 1). Fourth, because the training/test split was at the image level (Section 3.1.1), the accuracy may be optimistically biased by within-patient correlation. Fifth, because the output of this classifier is the starting point of every downstream group assignment, misclassification could propagate to the downstream analyses through the group assignment of a specimen. Sixth, collation of the deposited code revealed that the preprocessing used when the classifier was initially applied to the 792 images differed from that used during training: pixel values were divided by 255 during training but were not normalized at the time of application. We therefore re-classified all 792 images with the same normalization as in training, and all morphology-class fractions in this manuscript are based on that re-classification (the course of events and the comparison of the two conditions are given in Section 3.1.2; the re-classification code and per-image outputs are provided in the Supplementary Methods, and the per-specimen fractions under both conditions in Supplementary Table S46 (Additional file 16)). The outputs of the old condition were reproduced exactly, for all 792 images, against the records of the original run, and the re-classification agreed better with the manual labels (49/49 versus 45/49 for the 49 images overlapping the training/evaluation set, and 72/79 versus 63/79 for the 79 images whose file names carry manual labels). The separation of P1- and P2-dominant specimens by Type1 fraction held under both conditions, whereas the assignment between Type0 and Type5 within the cystic spectrum was sensitive to the preprocessing condition (Sections 3.1.2 and 4.1). Note that the comparison design of the expression analyses (DEG, IPA, and others) is isoform-based (Section 2) and does not depend on the preprocessing condition of the morphological classification. Seventh, the per-specimen Type fractions were calculated from a median of 10 images per specimen (range 2–48; 28 of the 63 specimens have fewer than 10 images), so the fractions of specimens with few images carry a large sampling error (Supplementary Table S2). Among the 4 specimens whose morphology deviates most from the typical pattern of their group (Section 3.1.2), the number of images ranged from 4 to 28, and the fraction for HCT41-1T rests on 4 images. Although this classification is based on single-time-point imaging, time-lapse observation in the same biobank reported that 19 of 21 organoids maintained stable morphology over 3 days [12], making it unlikely that short-term morphological fluctuation substantially confounds the classification. In light of these points, the reporting in this section follows the relevant items of the reporting standards for medical AI (CLAIM [85] and TRIPOD+AI [86]).

### R-4. Full-length version of main text Section 3.1.2 (Correspondence between MDM2 mRNA isoforms and morphology)

#### Correspondence between MDM2 mRNA isoforms and morphology (Figure 1)

In the 63 specimens classified by the deep-learning classifier (VGG16 transfer learning; classification performance and confusion matrix in Section 3.1.1 and Table 1), we performed Splicing Index (SI) analysis of the MDM2 gene across 33 two-group comparisons. SI values for exon 1 were positive on the tissue-1 (P1-dominant) side: the representative probe PSR12007998 was positive in all 33 comparisons (SI +2.12 to +18.50), and the other exon 1 probes (PSR12007995 and PSR12007996) were also positive in every comparison except for PSR12007995 in analysis 108 (Supplementary Table S12). Organoids of tissue 1 therefore preferentially use the P1 promoter and produce transcripts containing exon 1 (Figure 1A–C).

SI values for exon 2, by contrast, were negative on the tissue-2 (P2-dominant) side in all 33 comparisons (representative probe JUC12004243: SI range −4.77 to −569.76; PSR12008003 behaved likewise), indicating that organoids of tissue 2 preferentially use the P2 promoter and produce transcripts containing exon 2 (Figure 1A–C). Within the genomic structure — taking TAC analysis 49 as a representative example — the signs of the SI values are completely reversed between the regions corresponding to exon 1 (PSR12007995, PSR12007996, and PSR12007998) and those corresponding to exon 2 (JUC12004243 and PSR12008003), which directly shows that P1 and P2 promoter usage is shifted in opposite directions between tissue 1 and tissue 2 (Figure 1B, which also shows the 30 single-specimen split comparisons; SI values for all 63 comparison sets are given in Additional file 17: Supplementary Table S12). The isoform axis was found through exploration: TAC comparisons were first made between the two morphological groups, the MDM2 Splicing Index emerged there as a consistent feature, and later comparisons were made between isoform groups. Two of the 33 comparisons (analyses 49 and 52) retain the morphological grouping of that phase — HCT41-1T on the P1 side and HCT31-4LMR and HCT71-3LM on the P2 side — whereas the sides of the other 31 agree with the isoform assignment (Supplementary Table S3). The dominant promoter of each specimen (Supplementary Table S2) was taken from the original exon-level analysis. Single-specimen split SI against a common reference of the opposite type (HCT64-1T or HCT25-1T) confirmed the assignment for 53 specimens (45 of the 50 P1-dominant and 8 of the 13 P2-dominant specimens), all with the expected signs for the representative probes (PSR12007998 positive and JUC12004243 negative on the P1 side; individual probes deviated in four of these comparisons — PSR12007995 in analysis 108, PSR12008003 in analysis 115 and PSR12007996 in analyses 122 and 129); the remaining 10 specimens were the two common references themselves (HCT64-1T and HCT25-1T) and eight specimens compared only in multi-specimen comparisons or against another single specimen (analyses 49–54, 63, 110, 112 and 113; Figure 1E; Supplementary Tables S3 and S12). In total, opposite-sign values among the other exon probes occurred in only four of the 63 comparison sets (the deviations noted above).

Morphology and MDM2 isoform corresponded clearly at the specimen level (quantified below); the correspondence was established through the following course. In the initial application (before unification of the preprocessing), the specimens separated into a Type1-dominant group and a Type5-dominant group, which corresponded well to P1 and P2 dominance, respectively (this classification is the partition used in the exploratory comparisons that preceded the isoform grouping; Section 2). Because collation of the deposited code revealed the preprocessing mismatch (Section 3.1.1), all 792 images were re-classified with the same normalization as in training; the outputs of the old condition were reproduced for all 792 images, and the re-classification agreed better with the manual labels. Under re-classification, many of the cystic images that had been assigned to Type5 under the old condition moved to the Type0 class (of the 185 images assigned to Type0 by re-classification, the old calls were Type5 for 97, Type1 for 83, and Type0 for 5), and when the images of P2-dominant specimens assigned to Type0 were inspected visually, their morphology was not the compact glandular type (Type1) but cystic, close to Type5 (Section 4.1). The correspondence of morphology is therefore best captured as a two-group structure — “Type1 (compact glandular)” versus “non-Type1 (round morphology: a cystic–mucinous spectrum comprising Type0/Type5)” — and all quantification below is based on the per-specimen fractions from the re-classification (majority-vote labels are not used, because their outcome depends on the choice of rule). The 50 P1-dominant specimens had high Type1 fractions (median 0.826 [interquartile range 0.54–1.00] versus 0.444 [0.33–0.50] in the 13 P2-dominant specimens; Wilcoxon rank-sum test P = 1.1×10⁻³), and the P2-dominant specimens had high non-Type1 fractions (median 0.556 versus 0.174; AUC 0.79 for identifying P2 by the non-Type1 fraction; Type0 fraction: median 0.429 versus 0.095, P = 0.013; Type5 fraction: 0.111 versus 0.000, P = 0.036; Figure 1D; per-specimen fractions in Supplementary Table S2 and the comparison of the two conditions in Supplementary Table S46). Because specimens from the same patient are not independent and the 13 P2-dominant specimens derive from only six patients (HCT27, HCT33 and HCT64, whose specimens are all P2-dominant, and HCT38, HCT41 and HCT67, each contributing one P2-dominant specimen), we repeated the comparison at the patient level as a sensitivity analysis (R48, Supplementary Methods; mixed models fitted with the R package lme4 [84]). The difference was in the same direction but did not reach significance: in a linear mixed model of the Type1 fraction with a patient random intercept, the P2-minus-P1 difference was −0.21 (95% CI −0.42 to +0.02; likelihood-ratio P = 0.069; intraclass correlation 0.29); in a binomial mixed model of image counts with patient and specimen random intercepts, the odds ratio was 0.31 (95% CI 0.08–1.15; P = 0.089); and for patient × isoform means the medians were 0.725 versus 0.404 (18 versus 6 units; exact Wilcoxon P = 0.20). Excluding the three patients with specimens of both types, the mean Type1 fractions of the three exclusively P2 patients (0.35, 0.36, 0.44) were lower than the median of the 15 exclusively P1 patients (0.77; exact Wilcoxon P = 0.076; permutation of isoform labels between patients (20,000 random permutations, seed 1; 816 distinct arrangements are possible in total), one-sided P = 0.035; linear mixed model P = 0.041). All patient-level values assign HCT83-4LM to patient HCT83 (Section 2; R86); with HCT84 counted as a separate patient, the corresponding values were P = 0.093, 0.10, 0.20, 0.064, 0.035 and 0.053. The morphology–isoform correspondence is therefore established at the specimen level and held in the three exclusively P2 patients; after adjustment for patient it was in the same direction but not significant (linear mixed-model P = 0.069), and it requires confirmation in a larger number of patients, particularly P2-dominant patients. At this stage, however, it could not be determined whether morphology followed promoter usage itself or TP53 status, which largely coincided with promoter usage (Section 3.1.3); Sections 3.1.4 and 3.1.5 of the main text address this question and suggest that morphology follows TP53 status rather than promoter usage. Okamoto et al. (2022) [12] independently performed an AI-based 6-morphology typing analysis of CRC organoids in a cohort derived from the same biobank with largely overlapping patients (the HCT patient group), so the results of the two studies can be compared directly (Section 4.4). We first describe the specimens whose morphology deviates from the typical pattern of their group, three of which followed TP53 status rather than promoter usage (Sections 3.1.2 and 3.1.3), and then, in Sections 3.1.3, 3.2.2, 3.2.4 and 3.3, examine the expression programs of the two groups and the results of independent external validation.

##### Specimens deviating from the typical morphology of their group

The 4 specimens that deviate most from the typical pattern of their group on the axis of morphology-class fractions — the 2 P1-dominant specimens with the lowest Type1 fractions (HCT71-3LM 0.07 and HCT31-4LMR 0.13) and the 2 P2-dominant specimens with the highest Type1 fractions (HCT41-1T 1.00 and HCT38-3LM 0.71) — showed dissociation between morphology and isoform. Of note, the patients from whom these 4 specimens derive (HCT31, HCT38, HCT41 and HCT71), together with HCT67, the patient showing an intra-patient P1→P2 switch (Supplementary Report Section 4.2.1), are all among the patients in whom the independent morphological typing of the same-biobank cohort [12] reported organoids of both type A and type B within different lesions of a single patient (HCT38, 41, 27, 67, 31 and 71); this independently supports the interpretation that the morphology–isoform discordance reflects real intra-patient morphological admixture rather than classifier error. In HCT38-3LM a cancer-cell-intrinsic epithelial–mesenchymal transition (EMT) program was activated specifically in that specimen, separately from the central markers on which this study focuses (gastric metaplasia and p53 targets). Specifically, there was a cadherin switch to N-cadherin (CDH2) and CDH6, together with increases in vimentin (VIM), fibronectin (FN1), osteopontin (SPP1), versican (VCAN), TGFB2, LOXL4, and TGM2, accompanied by increases in ALDH1A3 (stemness) and ONECUT2 (lineage plasticity). None of these changes was seen in the other comparisons; they were specific to HCT38-3LM (Additional file 18: Supplementary Table S13; detailed profiles including the other 3 deviating cases are given in Additional file 19: Supplementary Table S14). Because stromal fibroblast markers (COL1A1, SPARC, FAP, and others) did not rise in parallel, we interpret this as partial EMT of the cancer cells themselves rather than as stromal contamination. Modification of morphology accompanying EMT and dedifferentiation may be one reason for the dissociation from MDM2 promoter usage, and could constitute the molecular background of the morphology–isoform discordance in this case. We treat EMT here as a modifying factor that helps explain morphologically deviating cases and do not test its causal contribution; causal validation with manipulation of the EMT state remains a task for future work.

HCT31-4LMR and HCT71-3LM, by contrast, showed no separate, independent program and remained at an intermediate profile in which the P2 signature was uniformly attenuated. HCT41-1T, the only primary tumor among the 4 cases, was included in the tissue-1-side reference group in the gene expression comparisons, so it was difficult to evaluate individually; it could, however, be evaluated by Splicing Index comparison based on a single-specimen split. When the TAC Splicing Index (MDM2 exon 1/exon 2) was evaluated in this way, the molecular type (promoter usage) of these 4 deviating cases agreed with the isoform assignment (Figure 1E). HCT31-4LMR robustly showed P1 with exon 1 dominance, within the SI range of concordant P1 specimens; HCT38-3LM robustly showed P2 with exon 2 dominance; and HCT71-3LM and HCT41-1T both showed intermediate SI values, with weak exon 1 and weak exon 2 dominance, respectively. Promoter usage in these 4 cases was therefore robust (HCT31-4LMR and HCT38-3LM) or borderline (HCT71-3LM and HCT41-1T), and in morphology three of the four (HCT38-3LM, HCT41-1T and HCT71-3LM) followed TP53 status rather than promoter usage (Section 3.1.3). These are exploratory findings based on a small number of cases, some of them single specimens, and confirmation requires validation in a large cohort.

Furthermore, in a within-patient paired analysis using the gene expression matrix of all 63 specimens (RMA normalization) (17 patients; primary–metastasis pairs; limma duplicateCorrelation method), no consistent differentially expressed genes (DEGs) were identified in the metastases (within-patient correlation coefficient 0.301; under the criteria of FDR < 0.05 and |log2FC| > 1.0, there were 0 genes both for increases and for decreases); nevertheless, EMT/mesenchymal genes (VCAN, SPP1, TGM2, FN1, CDH6, TGFB2, and others, 17 in total) increased in a consistent direction in the metastases (log2FC > 0 for all 17), and intestinal identity markers (OLFM4, LGR5, SLC26A3, and FOXM1) showed a consistent tendency to decrease (exploratory analysis; none significant after FDR correction).

When we calculated the EMT score of each specimen (the mean z-score of the EMT/mesenchymal genes minus the mean z-score of the epithelial markers), the EMT score was significantly higher in the metastatic specimens than in the primary specimens (metastasis median +0.130 versus primary median −0.590; per-specimen Wilcoxon rank-sum test P = 0.014. Because multiple specimens from the same patient are included, averaging per patient and performing a paired Wilcoxon signed-rank test also yields significance in the same direction (17 patients, P = 0.004)). In addition, the difference in EMT score between the P2-dominant and P1-dominant specimens was not significant (in all 63 specimens, median +0.130 for the 13 P2-dominant specimens versus median −0.396 for the 50 P1-dominant specimens; Wilcoxon P = 0.079) (Additional file 20: Supplementary Table S1; generating code: Code3, Supplementary Methods). Note that the actual distribution overlaid in Figure 9B is that of the 59 specimens with the 4 specimens whose morphology deviates markedly from the typical pattern of their group (Section 3.1.2) shown separately and grouped by sampling site rather than by isoform, because that is the axis the score follows (median −0.56 for the 19 primary specimens and +0.05 for the 40 metastatic specimens); because both the grouping and the target set differ, these medians do not agree with the values in this section. Sampling site is balanced between the two isoform groups (metastases: 34 of 50 P1-dominant and 9 of 13 P2-dominant specimens), so it does not account for the isoform contrast: within primary specimens the isoform difference was absent (medians −0.380 in the 4 P2-dominant versus −0.590 in the 16 P1-dominant specimens, exact P = 0.89), within metastases it rested on two specimens (medians +0.359 versus −0.210, P = 0.033; P = 0.20 after removing HCT38-3LM and HCT67-4LMR), at the patient level the 3 exclusively P2 patients did not differ from the 15 exclusively P1 patients (medians +0.129 versus −0.359, exact P = 0.65), and the score did not differ by TP53 status either (medians +0.130 in the 14 wild-type versus −0.439 in the 42 mutant specimens, P = 0.17). The EMT score therefore follows the sampling site rather than the two poles and is treated as a separate axis of progression (Additional file 7: Supplementary Table S48, sheet L). This consistency of direction suggests a tendency toward partial EMT-like reprogramming during the metastatic process, and is consistent as a background for the intrinsic EMT observed in HCT38-3LM (described above).

This interpretation of partial EMT is also consistent with the finding that the EMT-suppressive microRNA miR-200 family (MIR200C, MIR200B, and MIR141) showed a consistent tendency to be relatively high in the P2-dominant group (a partial EMT state in which ZEB1/2 are not completely suppressed even under elevated miR-200 but are maintained at low levels; exploratory, Additional file 21: Supplementary Table S15). This directionality was also confirmed quantitatively by the mean linear FC of the 33 paired TAC comparisons (negative = higher in the P2-dominant group). While the EMT-suppressive miR-200 family (MIR200C −1.40, MIR200B −0.85, MIR141 −0.77; all in the same direction in 25 of the 33 comparisons (76%)) was higher on the tissue-2 side, their targets ZEB1 (−0.90, 29/33 (88%)) and ZEB2 (−0.99, 28/33 (85%)) and the EMT transcription factor SNAI2/SLUG (−1.15, 32/33 (97%)) also showed high sign consistency in the tissue-2 direction, but the effect sizes all remained below 2-fold (below the DEG threshold). In contrast, the EMT transcription factor SNAI1 was rather higher on the tissue-1 side (+1.88, with 27/33 in the tissue-1 direction (82%), approximately 1.9-fold), so the direction of expression was split among the EMT transcription factors (for SNAI3, the consistency was 51.5% (16 of the 33 comparisons in the tissue-1 direction and 17 in the tissue-2 direction) and the mean linear FC was −0.021, and we detected no bias in either direction). The mesenchymal markers CDH2 (−0.86, 82%) and VIM (−0.85, 70%) were mildly higher on the tissue-2 side, whereas for the epithelial marker CDH1 the consistency was 60.6% and the mean linear FC was +0.300, and we detected no clear between-group difference (Additional file 21: Supplementary Table S15).

The heterogeneity of the patterns of metastatic change among patients is high (no paired DEGs were identified), and confirmation of the above tendencies requires validation in a large prospective cohort.

Across all 63 specimens, HCT38-3LM had the highest EMT score (4.94; 1st among all specimens), quantitatively corroborating the outstanding degree of intrinsic EMT activation in this specimen (Supplementary Table S1). In principal component analysis (PCA) and UMAP of the RMA-normalized expression matrix of all 63 specimens, P1-dominant specimens (blue) and P2-dominant specimens (red) separated in expression space; when the primary-to-metastasis trajectories of the same patient were drawn as arrows, the patients with both P1- and P2-dominant specimens (HCT38, HCT41, and HCT67) followed trajectories in directions markedly different from those of the other patients (Additional files 22 and 23: Supplementary Figure S3 and Supplementary Figure S4; generating code: Code6, Supplementary Methods).

The EMT and mesenchymal genes listed in this subsection (CDH2, CDH6, VIM, FN1, SPP1, VCAN, TGFB2, LOXL4, TGM2, ALDH1A3, ONECUT2, and others) are not a fixed list selected in advance to represent a mechanistic axis, as the P1/P2 signatures were. They are a descriptive enumeration, grounded in the known biology of EMT, of the program activated specifically in the single specimen HCT38-3LM and of the genes that increased consistently in the metastatic direction. They are therefore exploratory findings and are distinguished from the selected signatures.

##### Replication in organoid arrays not used for discovery

To test the P1/P2 separation outside the discovery set, we used the organoid arrays of the same biobank that had not been used for discovery and are identical to files public in GSE128213: 10 tumor arrays from seven patients, all of them other specimens from patients in the discovery cohort (the two arrays with the same titles as discovery specimens of HCT33 were excluded as other arrays of the same specimens), normalized together with the 63 discovery arrays (RMA). A per-specimen MDM2 P2 index computed from these values (mean of the exon-2 minus mean of the exon-1 probe sets, each relative to the MDM2 transcript cluster) separated the discovery specimens without overlap (P1-dominant maximum 0.78, P2-dominant minimum 1.49) and correlated with the P2 signature score (23 immune-excluded genes; Spearman ρ = 0.61, P = 1.0×10⁻⁷). In the 10 replication arrays, the P2 index again correlated with the P2 signature (ρ = 0.82, P = 0.0038; with the p53 targets removed, ρ = 0.68, P = 0.029; patient-level means, ρ = 0.82, P = 0.023, n = 7), and the three specimens called P2-dominant at the discovery threshold (HCT31-1T, HCT38-9LMR and HCT64-5LM) had higher P2 scores than the seven called P1-dominant (median +1.22 versus −0.17; Mann–Whitney P = 0.017). The P1 signature was not associated with the P2 index in either set (discovery ρ = −0.23, P = 0.066; replication ρ = −0.18, P = 0.63). Because all replication specimens come from patients already in the discovery cohort, this is replication in independent arrays and specimens rather than in independent patients. Adding one unpublished array (HCT55-1T, the only patient not in the discovery cohort) to give 11 arrays gave the same direction (ρ = 0.75, P = 0.007; Mann–Whitney P = 0.012) (Additional file 24: Supplementary Table S24, sheet 4; R85 and R89). Replication in organoids from independent patients is reported in Sections 3.1.5 and 3.2.5 of the main text.

### R-5. Full-length version of main text Section 3.3.1 (Tissue 1)

#### Tissue 1: genes higher in the P1-dominant, mostly TP53-mutant specimens (intestinal identity and proliferation; CIN-like expression features)

##### Overall picture of the gene expression profile

Here we list the genes highly expressed on the tissue-1 side detected consistently across all 33 sets (i.e., varying in the predicted direction in 70% or more of the comparison sets, which is the DEG criterion of this study). CALB1 alone, at 69.7% (23/33), falls slightly below the criterion, but it is listed together because its effect size is large. OLFM4 (avg FC +1,252, 73% consistent), SLC26A3 (+782, 88%), ALDH1A1 (+272, 94%), SLC9A3 (+199, 94%), NPSR1 (+516, 97%), MGAM2 (+87, 91%), AKR1B10 (+54, 91%), DPP4 (+25, 94%), LGR5 (+13, 82%), and CALB1 (+338, 69.7%) were identified as highly reproducible tissue-1-side markers (Table 2; values by analysis in Additional file 36: Supplementary Table S16; quantitative values at the gene level for all DEGs in Supplementary Table S4, the full gene-level DEG list). By functional category, we identified intestinal absorptive epithelial markers (SLC9A3/NHE3, SLC26A3/DRA, SLC6A20, SLC7A5/LAT1, ABCB1, CA12, DPP4, MGAM2, and ANPEP), metabolic enzymes (ALDH1A1, AKR1B10, GSTM4, UGT2A3, ADH1C, PYGB, SOAT1, and PLBD1), proliferation factors (FOXM1, MYBL2, MCM4, ATAD2, HELLS, TOP2A, AURKB, STMN1, and MSH2), and intestinal stem cell markers (OLFM4, LGR5, and LRIG1). The proliferation- and mitosis-related genes were biased toward the tissue-1 side with high sign consistency: MYBL2 (mean linear FC +9.66, 91% of the 33 comparisons), TOP2A (+6.54, 94%), FOXM1 (+4.58, 88%), AURKA (+3.24, 94%), BUB1 (+3.07, 94%), PLK1 (+2.79, 97%), E2F1 (+2.10, 94%), and MKI67 (+2.00, 91%) all showed tissue-1-side dominance. This is the transcript-level readout of the MYC/E2F/FOXM1 axis identified as an activating factor in the Upstream analysis (Section 3.3.1), and it shows that activation of the proliferation hub is corroborated not only by predicted activity but also by measured mRNA.

The list of differentially expressed genes of tissue 1 showed a profile centered on the absorptive and metabolic functions intrinsic to the large intestine. SLC9A3 (NHE3) and SLC26A3 (DRA), which mediate electrolyte and water transport, are the major Na+/H+ exchange and Cl−/HCO3− exchange transporters in the intestine, and their high expression indicates that the tissue is actively absorbing water and electrolytes. CA12 (carbonic anhydrase) helped supply the H+ and HCO3− required by these transporters.

High expression of MGAM2 (maltase-glucoamylase) and DPP4 (dipeptidyl peptidase 4), digestive enzymes of the microvillar brush border, corroborated that this tissue is a highly differentiated intestinal epithelium actively carrying out the terminal digestion of carbohydrates and proteins. Genes of the retinoid and aldehyde metabolic systems, represented by ALDH1A1 and AKR1B10, were also markedly enriched, indicating an extremely high capacity for metabolic defense against oxidative stress and xenobiotics.

Of particular note, LGR5 and OLFM4, stem cell markers of the colonic crypt base, were highly expressed (the direction and sign consistency of the WNT pathway genes as a whole are given in Additional file 35: Supplementary Table S27). OLFM4 was highly expressed (avg FC +1,252; positive in 73% of the 33 comparisons; however, the median was +4.0, and extreme overexpression in a small number of patients raises the mean). Okamoto et al. [12], who analyzed an overlapping patient group from the same biobank as this study, also independently confirmed high expression of intestinal stem cell markers such as LGR5 and OLFM4 in Type 1 PDOs (Figure 5D of that paper), which supports the reproducibility of the present finding. LGR5 is a target gene of Wnt signaling and, as Barker et al. showed [92], a definitive marker of intestinal stem cells. As van der Flier et al. reported [91], OLFM4 is also highly expressed in the LGR5-positive stem cell population, consistent with tissue 1 maintaining a crypt base columnar cell-type stem cell program. Furthermore, analyzing patient-matched primary–metastasis pairs from the same biobank, Okamoto et al. (2021) [24] identified OLFM4 as the gene most strongly correlating with a stem-like cell cluster, showed that OLFM4-positive cells were required for the efficient growth of primary-derived organoids but dispensable for metastasis-derived organoids, and reported that the expression of both differentiated-cell markers and intestinal stem cell markers was decreased in metastasis-derived organoids. The high OLFM4 expression observed in tissue 1, and its skew toward a small number of patients (see above), are consistent with this independently described inter-lesion and inter-patient heterogeneity of the OLFM4-associated stem-like cluster.

XIST (X-inactive specific transcript) showed a high avg FC of +410, but it was positive in only 67% of the 33 comparisons (median +0.8), and the mean derives from high values in 2 of 3 transcript clusters. XIST is a sex chromosome-linked lncRNA involved in X chromosome inactivation, so we interpreted this as variation biased by sex (female patients) and excluded it from interpretation as a biological marker. The high expression of NPSR1 (Neuropeptide S Receptor 1; avg FC +516, 97% consistent) may reflect activation of enteric nervous system pathways or the presence of ectopic neuropeptide signaling in the tumor. CALB1 (calbindin 1; avg FC +338, 69.7% consistent) is a calcium-binding protein that functions as a differentiation marker of the intestinal epithelium, consistent with the well-differentiated intestinal epithelial phenotype of tissue 1. The epigenetic regulators ATAD2, ATAD5, HELLS, DNMT1, SUZ12, and BMI1 were also simultaneously highly expressed, suggesting the establishment of a proliferative program through chromatin reorganization. High expression of non-coding RNAs (DUXAP10, FAM127B, and LINC01296) was reproducibly confirmed in multiple comparisons, suggesting that these lncRNAs may be involved in post-transcriptional regulation in tissue 1.

##### Summary of the molecular pathway landscape (IPA and EnrichR analyses; details in Supplementary Note Section 2)

The details of the molecular pathway landscape of tissue 1 (IPA and EnrichR analyses) are described in Supplementary Note Section 2; here we present a summary. In IPA Canonical Pathway analysis, cell cycle checkpoints, DNA synthesis/replication, the major DNA repair pathways (HDR, NHEJ, and BER), ribosome biogenesis, and aerobic metabolic pathways (e.g., oxidative phosphorylation and the TCA cycle) were simultaneously enriched among the top ranks, headed by “Processing of Capped Intron-Containing Pre-mRNA,” which ranked first overall (present in 33/33 analyses); this is the pathway profile typical of the CIN pathway (CMS2) type (Table 3). In Upstream Analysis, 121 factors (101 molecules and 20 chemicals) met the adoption criteria as activating factors in the tissue-1 direction, headed by MYC (median z = +5.76, consistent in 32/33) and including E2F, FOXM1, MDM4, and AURKB, indicating the establishment of an autonomous cell cycle-driving program that accelerates every step from the G1/S transition to the execution of M phase (the full list is in Additional file 37: Supplementary Table S7). In contrast, the intestinal identity factor CDX1 had a sign consistency of 93% but an effect size below the threshold (median z = +1.61), and for HNF4A the direction of activation was not determined (sign consistency 52%); the confirmatory basis for the retention of intestinal identity in tissue 1 therefore rests on the expression findings for the intestinal markers (Section 3.3.1). Disease & Bio Functions converged on DNA repair, survival of tumor cells, and glandular formation (the glandular-formation items Abnormal morphology of gland and Colorectal adenoma were below the effect-size threshold; Supplementary Note Section 2.3), and Tox Function on a proliferative toxicity signature headed by Nephritis (median z = +2.47; Additional files 38 and 39: Supplementary Tables S8 and S9), while EnrichR GO analysis showed enrichment of amino acid transport, DNA replication, and apico-basal polarity of the intestinal epithelium (Supplementary Results 2; the full lists of GO terms are given in Additional file 40: Supplementary Table S31). That is, in tissue 1 the simultaneous maximization of the proliferative machinery and the dominance of the MYC/E2F axis were compatible with the retention of intestinal identity.

### R-6. Full-length version of main text Section 3.3.1 (Tissue 2)

#### Tissue 2: genes higher in the P2-dominant, mostly TP53 wild-type specimens (p53 targets, metaplasia and inflammation)

##### Overall picture of the gene expression profile: p53 targets and gastric metaplasia

As genes consistently upregulated in tissue 2 (P2-dominant), we identified MDM2 (mean linear fold change 7.0-fold, median 6.0-fold, same direction in all 33 comparisons; one of the top DEGs; Table 4 and Supplementary Table S16), wild-type p53 target genes (CDKN1A/p21, DDB2, ZMAT3, FAS, TNFRSF10D, SPATA18, GDF15, BAX, BBC3), gastric metaplasia markers (CTSE/cathepsin E, REN/renin, TFF1, TFF3, ANXA10), small-intestinal Paneth cell markers (DEFA5, DEFA6), gel-forming mucins (MUC5B, MUC5AC, MUC17, MUC6), inflammation and hypoxia markers (CXCL14, IL33, DUOX2, TXNIP, CA9), invasion markers (MMP7, MSLN/mesothelin, PLAUR, LAMA3, LAMC2), long non-coding RNAs (HULC, UCA1, H19), an immune marker (B2M/β2-microglobulin), a secreted ER-related protein (ERP27), and an organic anion transporter (SLCO1B3) (Table 2; values for each individual analysis in Supplementary Table S16; gene-level quantitative values for all DEGs in Supplementary Table S4, the full gene-level DEG list; and the individual fold changes and consistency rates of the 10 wild-type p53 targets including MDM2 in Table 4).

The genes listed here for each functional category (Table 2) are examples of DEGs with high consistency and large fold change that represent that category; they are not a definitive list covering all DEGs (for all gene-level DEGs see Supplementary Table S4, and for all probe-level DEGs see Supplementary Table S16). In contrast, the 10 genes of the wild-type TP53 activation signature in Table 4 (TNFRSF10D, GDF15, DDB2, ZMAT3, MDM2, SPATA18, FAS, CDKN1A, BAX, BBC3) constitute a dedicated list selected by explicit inclusion criteria. The criteria were threefold: (i) a directional consistency of 90% or more across the 33 comparisons in this study—8 of the 10 genes also meet the DEG criterion of |mean linear fold change| of 2-fold or more, while BAX (−1.89, consistency rate 97%) and BBC3 (−1.09, 94%) do not reach a 2-fold effect size but were included because their direction is highly reproducible and they are established direct p53 targets; (ii) establishment in previous reports as direct transcriptional targets of wild-type p53 (based on the target-gene census of Fischer [37]; all 10 genes are among the 116 high-confidence p53 target genes of that census); and (iii) serving as an indicator of wild-type p53 activity, on the basis that P2 is a p53-inducible promoter [6,7]. MDM2 itself was also included in this signature because transcription from the P2 promoter is one of the p53 targets. In Table 4 the 10 genes are arranged in descending order of |mean linear fold change| (GDF15, TNFRSF10D, ZMAT3, DDB2, MDM2, SPATA18, FAS, CDKN1A, BAX, BBC3).

The list of differentially expressed genes for tissue 2, in descending order of mean fold change, comprised CTSE (−1,900.6), REN (−1,281.1), SNORD116-17/-19 (−662.4), SNORD116-15 (−568.1), SNORD116-18 (−556.1), UCA1 (−546.7), MUC17 (−477.6), TFF1 (−408.2), DEFA5 (−381.0), SNORD116-24 (−298.6), and HULC (−263.6), showing a molecular landscape entirely different from that of tissue 1.

Among the top differentially expressed genes, REN (renin; avg FC −1,281), which ranks second after CTSE (avg FC −1,901), is particularly noteworthy. Renin is normally produced by juxtaglomerular cells of the kidney and by mucous neck cells of the gastric mucosa and is essentially undetectable in the colon. Its marked expression is the clearest evidence that in tissue 2 the colonic differentiation program is attenuated and a gastric mucosa-like gene expression profile has been acquired. The simultaneous high expression of REN and AGT (angiotensinogen, a tissue-2-side hub molecule in the IPA Graphical Summary; Supplementary Note Section 4.4) indicates that components of the renin–angiotensin system co-occur within the tumor, although local function (angiotensin production) was not tested.

CTSE (cathepsin E) is an aspartic protease highly expressed in gastric chief cells and neck cells and is established as one of the signature genes specifically expressed in intestinal metaplasia of the stomach; its expression in colorectal cancer is consistent with gastric metaplasia. ANXA10 (annexin A10) has been reported by Bae et al. [93] as an immunohistochemical marker of colorectal cancer arising from the serrated pathway (for CIMP-P CRC, specificity 96.2% but sensitivity 39.7%), which is consistent with a serrated or MSI-like pathway origin of tissue 2 [94] (an expression-based inference).

TFF1 (trefoil factor 1) and TFF3 are mucosal protective factors secreted from the gastric surface epithelium (foveolar cells) and, as shown by Hoffmann [95], form the mucosal barrier in concert with the MUC family. Their expression in colorectal cancer indicates transdifferentiation (metaplasia) toward the gastric type.

DEFA5 and DEFA6 (α-defensin 5/6) are normally potent antimicrobial peptides secreted by Paneth cells of the small intestine. Their high expression in colorectal cancer indicates that, within intestinal metaplasia, an extremely peculiar transformation—small-intestinal Paneth cell-type differentiation—has occurred. In colorectal cancer, expression of these Paneth cell α-defensins has been reported to be potentially associated with prognosis (DEFA5 with favorable and DEFA6 with poor prognosis) [96]. In the mucin-producing system, multiple gel-forming mucins—MUC5B, MUC5AC, MUC17, and MUC6—were simultaneously highly expressed. MUC5B and MUC5AC are mucins characteristic of gastric surface mucous cells and neck mucous cells, whereas MUC17 is an intestinal-type mucin; their simultaneous expression indicates the complexity of metaplasia as a mixture of gastric and intestinal types. The mucus pool formed by these mucins may contribute to the highly viscoelastic, gel-like physical properties inferred for tissue 2 (inferred from expression; no mechanical measurements were performed; Supplementary Note Section 5.3). In this pattern, upregulation of gastric-type mucins (MUC5AC, MUC6) and gastric-type markers (TFF1, CTSE) coexists with retention of the colonic-type mucin MUC2 (in this study as well, the two-group directional consistency of MUC2 was only 48% [16 of 33 comparisons higher in tissue 1] and showed no significant between-group difference). This is directionally concordant with the established pathological finding that serrated lesions such as hyperplastic polyps show gastric pyloric gland-type differentiation, expressing MUC5AC, TFF1, and MUC6 while retaining MUC2 [97], and it supports the view that the gastric metaplasia of tissue 2 is continuous with the gastric differentiation program of the serrated pathway. More recently, a framework has been proposed in which this gastric metaplasia itself could be the point of origin of the serrated tumorigenesis pathway [98]. The primary study underlying that framework showed, by single-cell and spatial analysis of human colorectal polyps, that serrated polyps arise from differentiated cells via gastric metaplasia [99]. The gastric metaplasia signature observed in tissue 2 is directionally consistent with this framework.

##### Non-coding RNAs

Several of the top tissue-2 DEGs were C/D box snoRNAs of the SNORD116 cluster of the imprinted 15q11-q13 locus (Section 3.3.1; probe-level values in Additional file 36: Supplementary Table S16); as a continuous score the cluster followed neither P2 usage nor, consistently, TP53 status (Section 3.1.4 of the main text). The locus is analyzed in the companion study on the 15q11-q13 locus (T. Tsukui, R. Yao, and K. Tsuda, unpublished observations) and is not interpreted here.

Among non-coding RNAs, numerous long non-coding RNAs (lncRNAs) such as HULC (hepatocellular carcinoma up-regulated long non-coding RNA), UCA1, H19, NEAT1, and SNHG1 were also identified. These have been described as competing endogenous RNAs (ceRNAs) that sponge multiple microRNAs and may thereby act as an epigenetic amplification mechanism that indirectly increases the expression of genes involved in proliferation, invasion, and EMT.

##### Summary of the molecular pathway landscape (IPA and EnrichR analyses; details in Supplementary Note Section 3)

The details of the molecular pathway landscape of tissue 2 (IPA and EnrichR analyses) are given in Supplementary Note Section 3, and a summary is presented here. In the IPA Canonical Pathway analysis, Pathogen Induced Cytokine Storm Signaling (sign consistency 100%, median z = −3.32), extracellular matrix assembly and degradation, inflammatory signaling such as IL-17 and NF-κB, Colorectal Cancer Metastasis Signaling, HIF1α, Ferroptosis, and CGAS-STING were among the top-ranked tissue-2 pathways; this profile of inflammation, invasion, and hypoxic adaptation is diametrically opposite to that of tissue 1 (Table 3 and Additional file 41: Supplementary Table S6; of these, IL-17, Colorectal Cancer Metastasis, HIF1α, Ferroptosis, and CGAS-STING were below the effect-size threshold [|median z| < 2]; Supplementary Note Section 3.1). In the Upstream Analysis, NUPR1 (median z = −8.59, sign consistency 94%) and TP53 (−7.14, 100%) headed the list, and TGFB1/2/3, SMAD3, IL1B, TNF, HIF1A, CTNNB1, and MAPK3, among others, were identified as activating factors in the tissue-2 direction that met the adoption criteria (Supplementary Table S7). The most important finding is that wild-type TP53 ranks in the top class of activated upstream regulators; this, together with EGR1 (−1.95, 91%; below the threshold), is consistent with transcriptional induction of the MDM2 P2 promoter (Section 3.1.2 and Figure 1). Note that EGR1 mRNA expression varied widely among patients (Additional file 27: Supplementary Figure S8 and Additional file 28: Supplementary Table S18); whether this reflects patient-to-patient differences in the inputs to P2 activation was not tested. In Disease & Bio Functions, “Misalignment of chromosomes” ranked first, and DNA damage, apoptosis, necrosis, and senescence signals accumulated simultaneously (Supplementary Table S8; items such as Misalignment of chromosomes are likely inferred from the lower expression in tissue 2 of mitotic and checkpoint genes that are more highly expressed in tissue 1 and do not indicate chromosomal instability in tissue 2; Supplementary Note Section 3.3), while in Tox Functions, increased ALP (−2.12, 94%) was the top signal meeting the adoption criteria (Supplementary Table S9). In the EnrichR GO analysis, extracellular vesicle, exosome, and secretory granule lumen (CC) and Endopeptidase Inhibitor Activity (MF) were the most highly enriched terms (Supplementary Results 4). In short, in tissue 2, inflammation, invasion, hypoxic adaptation, and the acquisition of a secretory cellular architecture proceeded simultaneously under the dominance of wild-type p53 and stress-response factors.

##### Expression of immune checkpoint molecules (exploratory)

For the seven immune checkpoint molecules with directionally consistent higher expression on the tissue-2 side (TIGIT, CTLA4, PDCD1, IDO1, LAG3, IDO2, HAVCR2), the sign consistency in the tissue-2 direction was TIGIT 84.8%, CTLA4 84.8%, PDCD1 81.8%, IDO1 81.8%, LAG3 75.8%, IDO2 75.8%, and HAVCR2 66.7% (Figure 8B; Additional file 46: Supplementary Table S30). However, all seven molecules had mean linear fold changes of −0.38 to −0.77 and thus did not reach the DEG effect-size criterion (2-fold). HAVCR2 also fails to meet the consistency criterion (70%). Because these molecules derive mainly from tumor-infiltrating lymphocytes and their effect sizes are structurally compressed in bulk expression owing to dilution by the epithelial component, the findings in this section are exploratory, based on the reproducibility of direction rather than on effect size (for the same reason as the basis for adoption described in Section 2). Of these, the six molecules other than IDO2 (IDO1, PDCD1, LAG3, CTLA4, TIGIT, and HAVCR2) were adopted as the immune module of the P2 signature (details of the adoption criteria in Section 2 and in Section 4.5 of this Report; Supplementary Table S4). Note that this finding does not propose the P2-dominant group itself as a population indicated for immune checkpoint inhibitors (Section 3.3.3 and Supplementary Report Section 4.5.2).

### R-7. Full-length version of main text Section 3.3.1 (Integrated IPA analyses and the 39-gene panel)

#### Integrated IPA analyses and the 39-gene biomarker panel

##### Summary of the integrated IPA analyses (Regulator Effects, Networks, and Graphical Summary; details in Supplementary Note Section 4)

The details of the IPA Regulator Effects analysis, Networks analysis, and Graphical Summary are given in Supplementary Note Section 4; here we present a summary. In the Regulator Effects analysis (integrating the first 8 of the 33 analyses; Section 2; all cascades are listed in Additional file 47: Supplementary Table S32), the highest-scoring causal cascade (Rank1, Consistency Score 25.93) showed a chain running from upstream regulators such as EGR1, BMP4, and HIF1A through targets such as TP53 and VEGFA to increased ALP (a computational cascade; whether EGR1 drives P2 promoter usage or the ALP signal was not tested). The Rank2 cascade (Score 25.02) showed bidirectional regulation of MDM2, TP53, CDKN1A, and BAX by mitotic regulators such as ANLN and the AURK family, and a group of 11 recurrent upstream regulators including ANLN, AURK, CKAP2L, Eldr, LIN9, NEDD8, and PPM1A recurred across all 2,442 entries in Supplementary Table S32 (Supplementary Results 5). In the Networks analysis, MDM2 coexisted in the same protein–protein interaction module with the factors responsible for its degradation (FBXW7), phosphorylation control (CSNK2A1/2), and nuclear export (XPO1) (Additional file 48: Supplementary Table S19), and Cell Death and Survival and RNA Post-Transcriptional Modification were the most frequently observed network function categories. In the Graphical Summary (Additional file 49: Supplementary Figure S14), in which the top factors of each module were integrated using Cytoscape (version 3.10.4) [157], 32 factors such as FOXM1, MYC, MYBL2, E2F1/E2F2/E2F3, AURKB, and PLK1 in tissue 1 and 33 factors such as TP53, CDKN1A, RBL1, RBL2, IL1B, TGFB1, EGF, TNF, NUPR1, and AKT1 in tissue 2 were located at the central hubs (the full lists of the hub molecules and their directed relationships are given in Additional file 50: Supplementary Table S20 and Additional file 51: Supplementary Table S21). This contrasts the autonomously proliferative hub structure of tissue 1 with the hub structure of tissue 2, in which p53-induced apoptotic signaling and MDM2/AKT/EGF-dependent survival signaling are antagonistic. The RB-E2F axis was reciprocally regulated in the two types (tissue 1: RBL1/RBL2 inactivation → E2F derepression; tissue 2: RBL1/RBL2 activation → E2F suppression).

##### Separation of 63 samples into two clusters by the 39-gene panel identified with IPA Biomarker Detection

In the IPA Biomarker Detection analysis, 39 genes were identified as candidate biomarkers distinguishing tissue 1 from tissue 2. Using these 39 genes as input, we performed hierarchical clustering based on the Manhattan distance and the average linkage method with an R script that we implemented ourselves, and the 63 samples from 21 patients were divided into two clusters (51 and 12 specimens) (Supplementary Figure S5; the origin, function, and detection frequency of the 39 genes are given in Additional file 53: Supplementary Table S22; analysis code: Code7, Supplementary Methods). The final merge height of the dendrogram was 57.93, clearly exceeding the preceding merge height of 48.66, and the cophenetic correlation was 0.833. This hierarchical clustering is not a function of IPA but a separate analysis; however, because the 39 genes were selected by IPA Biomarker Detection from the same P1-versus-P2 comparisons, the 62/63 agreement reported below is a within-sample (non-independent) result. The two resulting clusters corresponded well with MDM2 promoter usage (P1-dominant/P2-dominant), agreeing in 62 of the 63 specimens (98.4%). The only exception was HCT41-1T, which was classified as P2-dominant yet fell into the P1-dominant cluster. Furthermore, of the four specimens whose morphology deviates from the typical pattern of their group (Section 3.1.2), three fell into the cluster corresponding to promoter usage rather than to morphology (the exception being HCT41-1T), consistent with this panel reading out the P1/P2 axis rather than morphology. That is, this 39-gene panel, although not independent of the grouping, is consistent with the morphology–isoform correspondence (Section 3.1.2).

The genes in this panel were not chosen subjectively by the authors; they were extracted by the IPA Biomarker Detection function as candidate molecules discriminating the two groups (tissue 1 and tissue 2) in the same P1-versus-P2 comparisons. The rationale for “why these 39 genes” therefore lies in the fact that the algorithm ranked and extracted them automatically according to their contribution to discrimination. The functional category classification presented below is a post hoc organization of the 39 extracted genes by the authors according to biological function; it is an interpretation of the results, not a criterion for selection.

The 39 genes identified are classified by functional category as follows. The seven categories below partition the 39 genes without overlap, with 9 genes in category 1, 5 in category 2, 2 in category 3, 1 in category 4, 9 in category 5, 2 in category 6, and 11 in category 7, totaling 39. Gene symbols follow the output of IPA Biomarker Detection (Supplementary Table S22), and where they differ from the row notation on the expression matrix side this is noted. The category to which each gene belongs is shown in the functional category column of Supplementary Table S22.

(Category 1: direct target genes of wild-type TP53 (9 genes)) Nine genes—CDKN1A (p21), SPATA18, FAS, TNFRSF10B (DR5), TNFRSF10C (DcR1), ZMAT3, TRIM22, DDB2, and MDM2—were included in the biomarker panel. Notably, 6 of the 10 genes of the wild-type TP53 activation signature shown in Table 4 (CDKN1A, DDB2, ZMAT3, MDM2, SPATA18, FAS) were identified again by IPA, a different analytical platform. The three analyses—EnrichR GO analysis, DEG analysis, and IPA Biomarker Detection—thus converge on the same biological signal of wild-type TP53 activation (Figure 8); because all three derive from the same P1-versus-P2 comparisons, this is concordance across platforms rather than independent validation.

(Category 2: invasion- and extracellular matrix-related genes (5 genes)) Five genes—MSLN (mesothelin), MMP7, SERPINB5 (maspin), COL17A1, and DCBLD2—were included. MMP7 (100% consistent; Section 3.3.1) and MSLN are both tissue-2-side-dominant DEGs with high consistency, and their reappearance in the biomarker panel indicates the molecular centrality of the invasive characteristics of tissue 2. MSLN is already used as a clinical tumor marker in peritoneal mesothelioma, pancreatic cancer, and ovarian cancer [102], and the significance of its high expression in P2-dominant colorectal cancer specimens (including any affinity for a metastatic target organ) was not tested in this study.

(Category 3: gastric metaplasia markers (2 genes)) CTSE (avg FC −1,901) and REN (avg FC −1,281) were included. That the two major markers of gastric metaplasia—one of the central phenotypes of tissue 2—reappeared in the biomarker panel (selected from the same comparisons, so this is not independent confirmation) suggests that the gastric-metaplasia program can be read from the expression signature; whether it corresponds to histopathologically diagnosed gastric metaplasia remains to be examined.

(Category 4: DNA damage response gene (1 gene)) PRKDC (DNA-PKcs; the catalytic subunit of DNA-dependent protein kinase) was included. It is a central factor responsible for non-homologous end joining (NHEJ) of DNA double-strand breaks. This gene is, however, expressed in the tissue-1 direction (mean linear fold change +8.02, sign consistency rate 94%; Supplementary Table S4), consistent with the accumulation of HDR and NHEJ among the top ranks in the Canonical Pathway analysis of tissue 1 (Section 3.3.1) and with its assignment to tissue 1 in Table 2 and in Additional file 54: Supplementary Table S43. A biomarker panel selects genes that contribute to separating the two tissue types, and inclusion in the panel does not imply attribution to the tissue-2 side.

(Category 5: immune- and cell signaling-related genes (9 genes)) The nine genes included NCR3LG1 (B7-H6), the ligand of the NK cell receptor NKp30; TNFRSF19 (TROY), a TNFR-type receptor in Wnt signaling; and HCP5, an lncRNA in the 6p21 MHC region (all genes and annotations are given in Supplementary Table S22).

(Category 6: two genes with unresolved annotation) These are two rows in which the annotation on the expression matrix side (Affymetrix HTA2.0) and that on the IPA Biomarker Detection output side (PYGB, RNF24) do not agree; in this study they are not assigned to a functional category and are treated as two genes with unresolved annotation (details of the probe set IDs and annotation correspondence are given in the notes to Supplementary Table S22). Because the clustering (Supplementary Figure S5) uses expression values only, this discrepancy in annotation does not affect the separation result.

(Category 7: other function-related genes (11 genes)) Eleven genes distributed across diverse functions—the methylation cycle (AHCY), glycosylation, lipid metabolism, DNA repair-related functions, MAPK dephosphorylation, and amino acid transport—were included (all genes and annotations are given in Supplementary Table S22).

We summarize four salient features of this biomarker panel. First, 6 (60%) of the 10 genes of the wild-type TP53 activation signature downstream of the MDM2 P2 pathway were re-identified by the panel. Second, direct markers of gastric metaplasia (CTSE, REN) were included in the panel, showing that the panel contains the gastric-metaplastic program at the level of specific genes (not an independent confirmation, because the panel was selected from the same comparisons). Third, MSLN, which already has a track record as a clinical tumor marker, is included, which could facilitate future evaluation in clinical samples (not tested here). Fourth, for two genes of the panel (TC20001170.hg.1, TC20001393.hg.1) the annotations on the expression matrix side and the IPA side did not agree, and gene assignment could not be determined. This arises because the transcript cluster annotation of HTA2.0 differs by version and source; it is a practical constraint that must be checked when this panel is transferred to other cohorts.

A panel size of 39 genes is of a scale that could in principle be adapted to an RT-qPCR multiplex assay or a targeted NGS panel. However, because the panel was selected from the same comparisons, it summarizes the p53-active/metaplastic contrast of these specimens rather than providing an independent classifier, and this is an exploratory analysis in a 21-patient cohort. Although the between-group differences of the related P1/P2 signatures were reproduced in TCGA-COAD/READ (n = 624) and GSE39582 (n = 519) (Section 3.3.2; discriminative performance was not evaluated), an assay derived from this 39-gene panel would require independent validation in biopsy tissue as well as external cross-validation in independent cohorts and prospective validation.

### R-8. Full-length version of main text Section 3.1.3 (TP53 mutation analysis)

#### TP53 mutation analysis: correspondence with morphology and MDM2 isoform

##### Overview of the TP53 mutations identified

Targeted resequencing of TP53 (status determined for 56 of the 63 samples; Section 2) identified, in the samples with variant tables, the mutations listed below (Supplementary Table S11). The pathogenic mutations were R273H (R114H/R141H in our data; rs28934576) and R248W (R116W; rs121912651) at DNA-contact residue hotspots, R175H (R136H; rs28934578) at a structural hotspot, the early stop codon R213* (R174*; rs397516436), and the loss-of-function mutations S127F (rs730881999), R158H (R119H; rs587782144), and R282W (R150W; rs28934574). As likely pathogenic mutations, M237I (M105I; rs587782664) and I255del (I123del; NM_000546) were detected. These are consistent with the TP53 mutations encountered in routine clinical practice for colorectal cancer and are also reported in the IARC TP53 database [66]. The published mutation maps showed further TP53 mutations, truncating in HCT37 and HCT50 and non-truncating in HCT38, HCT45, HCT47, HCT60 and HCT83, without the amino-acid change.

The benign polymorphism P72R (P33R; rs1042522) was detected in multiple samples. Because this codon 72 polymorphism is frequent in the Japanese population (minor [Pro72] allele frequency 0.35 in the Tohoku Medical Megabank panel; dbSNP rs1042522 [31]) and no effect on p53 function is recognized, it was not treated as a pathogenic mutation. In patient HCT67, the isoform-level calls S215R, L179V, S193R, P157A, and others were also made in the patient's normal liver or in other patients, and the ANNOVAR annotation of the same samples contained no corresponding coding change (only intronic and 5′-UTR variants besides P72R); they were therefore not treated as somatic mutations, and R175H was the only somatic coding mutation of this patient, detected in the primary tumor and the liver metastasis but not in the recurrent liver metastasis HCT67-4LMR.

##### Three-way correspondence among morphology, MDM2 isoform, and TP53 mutation status

We examined how the TP53 mutation status of each patient corresponded to the MDM2 isoform (P1/P2) and to the organoid morphology (Type1 fraction; Type1 versus non-Type1) (Additional file 25: Supplementary Table S23, compiled from Supplementary Tables S11 and S2). The following clear patterns emerged.

Distribution of TP53 mutations in the P1-dominant group. Among the 18 patients with P1 isoform dominance, 15 carried TP53 mutations (HCT17, HCT25, HCT26, HCT31, HCT37, HCT38, HCT41, HCT43, HCT45, HCT47, HCT50, HCT51, HCT60, HCT67, HCT83), including, in the samples with variant tables, the pathogenic or likely pathogenic mutations R273H, R248W, R175H, R213*, S127F, R158H, R282W, M237I, and I255del; two were wild type (HCT59, which carries only the variant of uncertain significance F113V together with the benign polymorphism P72R, and HCT71), and one was undetermined (HCT69). Here the per-patient call follows the rule that a patient is classified as mutant when a pathogenic or likely pathogenic mutation is detected in at least one specimen from that patient — a rule that matters for HCT67, whose recurrent liver metastasis HCT67-4LMR carried none — and the patient is assigned to the P1 or P2 group by the isoform of the majority of its specimens (per-specimen calls in Supplementary Table S23). Thus, TP53 mutations largely coincide with the P1 group (patient level, Fisher P = 0.0088; among specimens with determined status, 40 of 43 P1-dominant and 2 of 13 P2-dominant, P = 1.7×10⁻⁷), although they are not a requirement for it. All of the detected mutations with a known amino-acid change impair wild-type p53 transactivation (loss-of-function or dominant-negative [87]; the map-derived mutations were reported only as truncating or non-truncating), as is typically observed in the CIN (chromosomal instability) pathway of colorectal cancer (some hotspot missense variants have also been proposed to exert allele-specific gain-of-function activities [68]), and in these cases the loss of wild-type p53 function is mechanistically consistent with MDM2 remaining at the constitutive basal expression driven by the P1 promoter, because the p53 required to activate the MDM2 P2 promoter through its p53 response elements [88] is absent (the two TP53-mutant, P2-dominant specimens, which were p53-active, are the exceptions examined in Section 3.2.1 of the main text). The two wild-type P1-dominant patients suggest that P1 dominance may also arise through routes other than TP53 mutation (for example, failure of P2 induction by EGR1 or similar factors); p53 was inactive in both specimens of HCT59 (Section 3.2.1 of the main text).

Retention of wild-type TP53 in P2-dominant patients. In contrast, in the patients with P2 isoform dominance (HCT27, HCT33, HCT64), no pathogenic mutation was detected and only the benign polymorphism (P72R/P33R) was confirmed (effectively wild-type TP53); that is, at the patient level (majority isoform) all three P2 patients were TP53 wild-type, whereas at the specimen level the P2 specimens HCT41-1T and HCT38-3LM, from P1-dominant patients, carried TP53 mutations yet were p53-active (Section 3.2.1 of the main text), and the P2 specimen HCT67-4LMR carried none (Supplementary Table S11). The retention of wild-type p53 is consistent with tumorigenesis through routes that do not pass through TP53 mutation, such as MSI-like or serrated-pathway tumorigenesis; this is a signature-level resemblance rather than an MSI status, because MSI was not measured and one of the three P2 patients (HCT27) was microsatellite-stable (Section 4.6). In the mutation analysis accompanying the CMS classification of Guinney et al. [2], TP53 mutations—unlike other cancer drivers—were not enriched in CMS1 (the MSI immune subtype) despite its hypermutated background, and our data are consistent with this observation. Wild-type p53 may be persistently activated in response to inflammatory stress (TNF, IL1B and ROS, inferred from expression and IPA prediction rather than measured), which would induce transcription from the MDM2 P2 promoter. For two of the P1-dominant patients (HCT31 and HCT71), only a single specimen was obtained; in each, the specimen was P1-dominant yet had a low Type1 fraction and Type5 as its largest class fraction (HCT31-4LMR: Type1 fraction 0.13, Type5 fraction 0.50; HCT71-3LM: 0.07 and 0.82), that is, they are specimens deviating from the typical pattern of their group on the axis of morphology-class fractions (2 of the 4 deviating cases; Section 3.1.2; Supplementary Table S23). Because the group is assigned by isoform, both are counted in the P1-dominant group above (HCT31 as mutant, carrying R273H and R248W; HCT71 as wild type), so that the 18 P1-dominant and 3 P2-dominant patients account for all 21 patients. TP53 status also aligned with morphology: mutant specimens had a higher Type1 fraction than wild-type specimens (median 0.854 versus 0.422, P = 6.1×10⁻⁴; Figure 2E and Figure 1D), to a degree similar to promoter usage (Section 3.1.2), and three of the four morphology-deviating specimens (HCT38-3LM and HCT41-1T, P2-dominant but TP53-mutant and Type1-leaning; HCT71-3LM, P1-dominant but wild-type and non-Type1) followed TP53 status rather than promoter usage. In this cohort, therefore, the morphology–isoform correspondence cannot be separated from TP53 status by these descriptive comparisons alone; Sections 3.1.4 and 3.1.5 of the main text separate the two by adjusted analyses and in an independent biobank.

##### Consistency of the tissue-2 traits across the three P2 patients

By mapping the fold changes of each TAC two-group comparison onto the sample composition of the comparison design, we evaluated the molecular behavior of the tissue-2/P2 patients (HCT27, HCT33, HCT64) at the patient level (Additional file 24: Supplementary Table S24; per-patient and per-comparison values in Additional file 26: Supplementary Table S26; Additional file 27: Supplementary Figure S8). MDM2, CTSE, REN, and MMP7 were consistently higher on the tissue-2 side in all three patients (a common basis), and these characterize the core traits of tissue 2. EGR1, by contrast, varied among the patients: it was consistently higher on the tissue-2 side in HCT27, leaned toward tissue 2 in HCT64 and was higher on the tissue-1 side in HCT33 (Additional file 28: Supplementary Table S18; Supplementary Note Section 3.2). This is an observation in three patients.

### R-9. Full-length version of main text Section 3.3.2 (External validation: cohorts)

#### External validation and direct quantification in independent cohorts

We externally validated the organoid DEG signatures of this study using two independent large public cohorts, TCGA-COAD/READ (n = 624 primary tumors) and GSE39582 (519 tumors with MMR status annotation among 585 samples, the 19 non-tumoral mucosa samples removed; 75 dMMR and 444 pMMR) (analysis code: Code8 and R49, Supplementary Methods). P1 and P2 signature scores were calculated by GSVA [78], and their associations with CMS subtype and MSI status were evaluated.

##### Validation of the signatures in the TCGA-COAD/READ cohort

From the RNA-seq data of 647 aliquot rows (624 specimens) (STAR-Counts, tpm_unstrand) we constructed a log2(TPM+1) matrix and performed CMS classification with CMScaller [75] (Entrez ID, RNAseq=TRUE, FDR < 0.05), which was possible for 574 rows. Averaging multiple aliquots of the same specimen gave 563 specimens (CMS1: 96, CMS2: 167, CMS3: 96, CMS4: 204; a further 61 unclassifiable cases were excluded from this comparison), and the analyses below are performed at this per-specimen level. In the comparison of GSVA scores across CMS classes (Kruskal–Wallis test), both the P1 signature score and the P2 signature score differed significantly among the CMS classes (P = 7.2×10⁻¹⁵ and P = 3.3×10⁻²⁸, respectively; for the P2 score without the immune checkpoint genes, which is the primary score in the main text and is shown in Supplementary Figure S6B, P = 2.54×10⁻³⁰, highest in CMS3 [median 0.397 versus 0.202 in CMS1, BH-adjusted P = 1.95×10⁻⁴]). However, the class showing the highest value differs when the signature is decomposed into functional modules (the module definitions are given in Section 2, “External validation of the signatures in independent cohorts and survival analysis”; the breakdown of all modules is in Additional file 42: Supplementary Table S34). The median of the whole P1 signature (21 genes) was highest in CMS1, but the difference from CMS2 was not significant (0.167 versus 0.107, BH-adjusted Wilcoxon P = 0.349), and both were significantly higher than CMS3 and CMS4. For the differentiation core (13 genes), which excludes the proliferation module (8 genes), CMS2 was highest and significantly higher than CMS1 (0.150 versus 0.012, P = 0.038; Kruskal–Wallis P = 1.0×10⁻⁶). Conversely, for the proliferation module alone, CMS1 was highest and the difference from CMS2 was significant (0.387 versus 0.101, P = 1.1×10⁻⁴). That is, the whole P1 score leans toward CMS1 because MSI-type tumors are highly proliferative, whereas the lineage trait of intestinal identity corresponds to CMS2. A similar structure is seen for the P2 signature. For the whole signature (29 genes), CMS1 and CMS3 were equivalent, with no difference between them (0.281 versus 0.281, P = 0.929), whereas for the core (metaplasia plus p53 targets, 21 genes), which excludes the immune/lncRNA module (6 immune checkpoint genes plus 2 cancer-associated lncRNA genes), CMS3 was highest and significantly higher than CMS1 (0.435 versus 0.256, P = 1.6×10⁻³; Kruskal–Wallis P = 5.3×10⁻³¹). For the gastric/Paneth metaplasia module alone (11 genes) this difference was even larger (CMS3 0.478 versus CMS1 0.204, P = 5.9×10⁻⁶), whereas for the immune module alone CMS1 was conversely highest (0.673 versus CMS3 −0.183, P = 1.6×10⁻¹³). When the same specimens and the same gene sets were scored by ssGSEA [101], the ranking of the highest class given above was concordant for all 11 modules that were scored (the identity-TF module of P1 consists of HNF4A alone and is therefore not scored as a module by either GSVA or ssGSEA; Section 3.3.2) (at the per-aliquot level before aggregation, only the P1 intestinal identity module was discordant, but this was resolved at the per-specimen level; with either method the difference between CMS1 and CMS2 is not significant). However, because ssGSEA scores have a smaller variance and therefore lower statistical power for the contrast between CMS1 and CMS3, the values of the pairwise comparisons were taken from GSVA and ssGSEA was used only to confirm the concordance of the rankings (Supplementary Table S34). Taken together, these results show that the signatures identified from 21 patients' organoids are consistent with clinical CMS subtypes in a large independent cohort of 624 specimens (563 of which were CMS-classifiable), and that this correspondence takes the form of different functional modules corresponding to different classes rather than of an assignment to a single class.

##### Validation of the signatures in the GSE39582 cohort

We applied GSVA to the microarray data of GSE39582 and compared the 519 tumors with MMR status annotation. The P2 signature score was significantly higher in dMMR (equivalent to MSI-H) (Wilcoxon rank-sum test (normal approximation) P = 1.14×10⁻¹⁷; n = 519; for the immune-excluded P2 score, the primary score shown in Supplementary Figure S6C, P = 6.59×10⁻¹⁰), consistent in an independent cohort with an MSI-associated expression profile of the P2 signature (not of P2 promoter usage, which was not associated with MSI status; Section 3.2.2 and Section 3.1.4 of the main text). The P1 signature score was significantly higher in pMMR (P = 0.017). This effect size is smaller than the dMMR skew of the P2 score, but that is because the pMMR cohort contains a mixture of CMS2 (P1-like), CMS3, and CMS4, so that the signal is partially diluted by CMS3/CMS4; it is consistent with the CMS-wise distribution of the P1 score in TCGA (P = 7.2×10⁻¹⁵; in the differentiation core, CMS2 is highest; Section 3.3.2).

### R-10. Full-length version of main text Section 3.3.2 (P1 signature score and prognosis)

##### P1 signature score and prognosis: Cox proportional hazards analysis

We performed survival analysis using the TCGA-COAD/READ cohort (n = 591 patients with OS information, after per-patient deduplication; the P1 signature is the 21-gene version, from which MSH2 was excluded to avoid circularity, and this version is used in all analyses in this manuscript).

**Kaplan–Meier analysis**

In Kaplan–Meier analysis with a two-group split by the median P1 signature score (high group: P1-like, 295 cases / low group: P2-like, 296 cases), a trend toward prolonged OS was seen in the high group, but it did not reach the significance level (log-rank P = 0.114; Supplementary Figure S7A). Median survival was not reached in the high (P1-like) group (lower bound of the 95% CI = 3,042 days) and was 1,910 days in the low (P2-like) group (95% CI 1,661 days–not reached). A two-group split by the P2 signature score (high group: P2-like, 295 cases / low group: P1-like, 296 cases) also showed no significant difference (log-rank P = 0.104; Supplementary Figure S7B; median survival: high group 3,042 days (95% CI 2,532 days–not reached), low group 2,003 days (95% CI 1,711 days–not reached)). In the CMS-wise KM analysis, overall survival differed significantly among the CMS subtypes (log-rank P = 0.0072; n = 533; CMS1 91 cases, CMS2 158 cases, CMS3 92 cases, and CMS4 192 cases; Supplementary Figure S7C). CMS2 and CMS3 did not reach median survival within the observation period (events: 27 of 158 for CMS2 and 8 of 92 for CMS3), whereas CMS1 was 2,134 days and CMS4 was the shortest at 1,881 days. The poor prognosis of CMS4 is consistent with the previously reported CMS prognostic classification [2].

**Cox proportional hazards analysis**

In univariate Cox analysis the P1 score showed a trend toward association with a reduced risk of death, but it did not reach the significance level (HR per 1 SD = 0.841, 95% CI: 0.703–1.004, P = 0.056). In multivariate Cox analysis entering the P1 and P2 scores simultaneously, the P1 score likewise showed a trend toward association with a favorable prognosis but did not reach the significance level (mutually adjusted for the P2 score; HR per 1 SD = 0.852, 95% CI: 0.713–1.019, P = 0.080; Supplementary Figure S7D). The P2 score was also not significant in the multivariate analysis (HR per 1 SD = 0.893, 95% CI: 0.747–1.066, P = 0.210). The differentiation core of the P1 signature (13 genes, excluding the proliferation module) is consistent with CMS2, the canonical/WNT type; the highest whole score is in CMS1, but its difference from CMS2 is not significant (P = 0.349; Section 3.3.2). The relatively favorable prognosis of CMS2 is consistent with the present analysis and with previous reports [2].

The multivariate analysis above is based on mutual adjustment between the P1 and P2 scores and does not include adjustment for established clinical factors. We therefore performed adjusted Cox regression with age, stage, and MSI status as covariates; after this adjustment, no independent contribution of the P1 score was observed (likelihood ratio test for incremental prognostic value over the clinical model P = 0.65; C-index = 0.785; complete cases n = 214), and the association with prognosis was attributable mainly to stage. In a sensitivity analysis that added tumor purity (ESTIMATE) as a covariate, no independent contribution of the P1 score was observed either (HR = 0.97, 95% CI 0.67–1.40; likelihood ratio test P = 0.88; C-index = 0.790; complete cases n = 214). In module-wise univariate Cox analysis, the prognostic signal was carried mainly by the proliferation module (HR per 1 SD = 0.82, P = 0.029), whereas the differentiation (identity) module was not significant (P = 0.37). Therefore, while this signature is associated with molecular subtype, MSI, and TP53 mutation status, positioning it as a prognostic factor independent of established clinical factors will require larger adjusted analyses. To confirm that this finding does not derive from a loss of statistical power caused by missing covariates, we performed sensitivity analyses with different covariate sets. With age alone (n = 591, 122 events), HR = 0.84 (95% CI 0.71–1.01) and likelihood ratio test P = 0.063; with age plus stage (n = 477, 79 events), HR = 0.93 (0.75–1.15), P = 0.51; and with age plus stage plus MSI (n = 214, 36 events), HR = 1.08 (0.79–1.48), P = 0.65; the incremental prognostic value was not significant in any of these sets. The association between the P1 score and prognosis is attenuated at the point at which stage is adjusted for (HR 0.84 → 0.93), and the changes brought about by the further addition of MSI and tumor purity were slight. That is, the non-significant results of this section are not explained by a loss of power due to missing MSI information (available in only 252 of 591 cases).

### R-11. Full-length version of main text Section 3.3.2 (Specificity and robustness of the signatures)

##### Specificity and robustness of the signatures and their mechanistic validity (sensitivity analyses)

To confirm that the association between the signatures and the CMS classification is not an artifact arising from gene overlap, we examined the overlap between the signature genes and the CMS calling templates of CMScaller; the overlap remained at a maximum of 3 genes for each CMS class and each signature side (even when summed per class, the maximum is 4 genes for CMS3). The overlapping genes were, against the CMS1 template (126 genes), 2 genes on the tissue-1 side (CALB1, DPP4) and 1 gene on the tissue-2 side (CTSE); against the CMS2 template (82 genes), 2 genes on the tissue-1 side (OLFM4, SLC9A3) and 0 on the tissue-2 side; against the CMS3 template (84 genes), 1 gene on the tissue-1 side (OLFM4) and 3 genes on the tissue-2 side (CTSE, TFF1, TFF3); and against the CMS4 template (237 genes), 0 genes on both the tissue-1 and the tissue-2 side. The overlap is limited to intestinal absorptive epithelium markers (CALB1, DPP4, OLFM4, SLC9A3) and gastric metaplasia markers (CTSE, TFF1, TFF3); the 8 genes of the P1 proliferation module and the 10 p53 target genes, 6 immune checkpoint genes, and 2 cancer-associated lncRNA genes of P2 do not overlap with any CMS template by even a single gene. That is, the correspondence between the signatures of this study and the CMS classification did not arise mechanically from a sharing of gene sets (Additional file 45: Supplementary Figure S9: sensitivity analysis; generating code: make_FigureS9_v3.py). Furthermore, every scored functional submodule was significantly associated with CMS on its own (identity P = 3.1×10⁻⁷, proliferation P = 3.3×10⁻¹⁴, metaplasia P = 1.2×10⁻²⁹, p53 targets P = 3.1×10⁻¹⁶, immune P = 1.2×10⁻²⁶, lncRNA P = 2.3×10⁻²¹; all at the per-specimen level, 563 specimens), showing that the signal does not depend on a single module. Note that no module score was calculated for the identity-TF module of P1, which consists of HNF4A alone, because GSVA by design requires a set of two or more genes (HNF4A itself is included in the whole P1 signature).

To examine the possibility that the association between the P2 signature and MSI derives from the definitional overlap between immune-related genes and MSI-H (circularity), we reanalyzed GSE39582 using versions from which the immune checkpoint genes were excluded. Even so, the P2 score was significantly higher in dMMR (immune-excluded version P = 6.59×10⁻¹⁰; core version of metaplasia plus p53 targets P = 2.63×10⁻¹¹), showing that the association between the P2 signature and MSI does not depend on circularity. For this reason, the immune-excluded version is used as the primary P2 score for the CMS and MMR associations in the main text (Kruskal–Wallis P across CMS = 2.54×10⁻³⁰; Additional file 42: Supplementary Table S34).

As a positive control expected from the p53-target content of the P2 signature (rather than an independent validation), we used the TP53 mutation status in TCGA (Masked Somatic Mutation). The P2 score, and in particular the p53 target module, was strongly associated with TP53 mutation status (whole P2 signature P = 1.8×10⁻⁴¹; p53 target module P = 5.4×10⁻⁴⁷). In terms of direction, the median p53 target module score was significantly higher in TP53 wild type (wild type +0.48 versus mutant −0.41), and the whole P2 signature showed the same direction (wild type +0.25 versus mutant −0.23). These comparisons are restricted to the 578 specimens for which somatic mutation data were available, comprising 234 wild-type and 344 mutant specimens; cases for which somatic mutation data were not available were not treated as wild type and were excluded from the analysis. Given that expression of the p53 target group reflects functional wild-type p53 activity, this is consistent with p53 driving its targets, including P2 [6,7]. In contrast, the P1 score was slightly higher in TP53 mutants, but the effect size is small (median: wild type −0.077 versus mutant +0.026, Wilcoxon P = 0.0094). This difference is an order of magnitude smaller than that of the p53 target module (wild type +0.48 versus mutant −0.41) and does not change the positioning of P1 as a p53-independent autonomous proliferation program. These sensitivity analyses support the view that the validity of the signatures with respect to molecular classification, MSI, and TP53 is robust and does not rest on circularity or on artifacts of gene overlap.

### R-12. Full-length version of main text Section 3.2.2 (Direct quantification of MDM2 P1/P2 promoter usage in an independent cohort)

##### Direct quantification of MDM2 P1/P2 promoter usage in an independent cohort

In order to examine P1/P2 promoter usage directly in an independent cohort and on an independent platform, without relying on the downstream signature of this classifier (the exon 1 versus exon 2 Splicing Index), we quantified MDM2 P2 promoter usage (P2_index = ΣTPM(P2)/ΣTPM(P1+P2)) directly at the transcript level in TCGA-COAD/READ (624 cases) (the analysis code is provided as Code9, and the code that generates its input expression matrix and covariate table as Code16; both are in the Supplementary Methods). P2_index was associated with TP53 mutation status (Wilcoxon P = 1.0×10⁻⁵), and its median was significantly higher in TP53 wild type (wild type 0.481 versus mutant 0.404; 132 wild-type and 242 mutant specimens). Here “wild type” refers to cases in which no functional mutation was detected in the Masked Somatic Mutation data, and non-detection of a mutation is not distinguished from true wild type. In addition, the 6 specimens for which somatic mutation data were not available were, by the same rule as in Section 3.3.2, not treated as wild type and were excluded from this comparison (374 specimens analyzed). This is consistent with the mechanistic hypothesis that functional wild-type p53 drives the p53-responsive promoter P2. In the same Toil recompute, adjacent normal colorectal mucosa used mainly P1 (median P2_index 0.289, n = 51), and both wild-type and mutant tumors were above it (P = 1.1×10⁻¹² and 3.5×10⁻⁶); in 32 patient-matched pairs the tumor had the higher P2_index in 23 (Wilcoxon P = 6.1×10⁻⁴) (Additional file 7: Supplementary Table S48, sheet I; code r160_S264_normal_baseline_P2_index_v1.py, Supplementary Methods). Furthermore, P2_index correlated positively with the p53 target module excluding MDM2 (Spearman ρ = +0.248, P = 1.1×10⁻⁶, n = 380), and also showed significant positive correlations with the whole P2 signature (excluding MDM2; ρ = +0.153, P = 2.8×10⁻³) and with the gastric/Paneth metaplasia module (ρ = +0.162, P = 1.6×10⁻³; unadjusted — after adjustment for TP53 status the metaplasia genes did not follow P2_index, Section 3.1.4 of the main text) (all at the per-specimen level, 380 specimens. When multiple specimens were available from the same patient, only the first specimen in ascending barcode order was used, and all 380 specimens are primary tumors (sample type 01). Because P2_index is calculated from MDM2 transcripts, MDM2 is excluded from the modules with which the correlations are taken). In contrast, no correlation with the P1 signature was observed (ρ = −0.053, P = 0.30, n = 380), consistent with the positioning of P1 as a p53-independent autonomous proliferation program.

P2_index also differed significantly among the CMS subtypes (Kruskal–Wallis P = 1.48×10⁻⁴, n = 342; Figure 5). CMS calling by CMScaller is permutation-based, so the assignment of borderline cases can change between runs; all values in this section are therefore computed from the fixed specimen-level CMS calls that are also used for the other CMS analyses of this manuscript. However, this difference derives from CMS4 being the lowest. After Benjamini–Hochberg correction, only CMS1 versus CMS4 (medians 0.466 versus 0.390, P = 0.031) and CMS2 versus CMS4 (0.465 versus 0.390, P = 1.3×10⁻⁴) were significant, and no significant differences were found among CMS1, CMS2, and CMS3 (all P ≥ 0.217). The medians, P values, and n values were all calculated from the same per-specimen set of 342 specimens. The association with MSI status was not significant (P = 0.19; 8 MSI-H versus 24 MSS/MSI-L specimens, 32 specimens in total. Because this is limited to cases for which both P2_index and MSI annotation were available, statistical power is poor and this non-significance does not mean the absence of an association); this is compatible both with MSI annotation being available for only a subset of cases and with the P2 signature resembling the MSI-like/serrated program only at the signature level, P2 dominance not being equivalent to MSI. No association with overall survival was observed (the 368 specimens for which both P2_index and overall survival information were available were analyzed; Kaplan–Meier analysis with a split at the median gave log-rank P = 0.82; univariate Cox gave HR per 1 SD = 1.06, P = 0.57. The SD of P2_index is 0.169, which converts to HR = 1.41 per unit), which is consistent with the finding of this study that prognostic independence was not observed after adjustment for stage (Section 3.3.2).

These associations were robust to multiple-testing correction, and the positive correlations with the p53 target module, the whole P2 signature, and the gastric metaplasia module all remained significant after FDR correction by the Benjamini–Hochberg procedure, which was applied to the three correlation tests (maximum FDR = 2.8×10⁻³). The 95% confidence intervals of the effect sizes also support the directionality (p53 targets ρ = +0.248, 95% CI +0.15 to +0.34), and only for the P1 signature did the confidence interval cross 0 and remain non-significant (ρ = −0.053, 95% CI −0.15 to +0.05). Furthermore, in a sensitivity analysis restricted to the cases for which the covariates (TP53 status, CMS, tumor purity, MDM2 copy number) were available, the association between P2_index and TP53 status was independently significant in a multivariate linear model adjusted for the other factors (standardized partial regression coefficient for TP53 wild type = +0.580, 95% CI +0.357 to +0.803, P = 5.3×10⁻⁷, n = 328). In the same model, tumor purity showed a positive contribution (+1.316, P = 0.0092), and none of the CMS levels was significant (CMS2 +0.139, P = 0.42; CMS3 −0.240, P = 0.20; CMS4 −0.192, P = 0.22). The positive correlation with the p53 target module was also maintained in a partial correlation adjusted for tumor purity and CMS (Spearman partial correlation ρ = +0.253, P = 2.3×10⁻⁶, n = 342). MSI was not included among the covariates because only 29 specimens have all of the other covariates of this model available, and only 27 when MDM2 copy number is added (the 32 specimens of the univariate comparison above are the specimens for which both P2_index and MSI annotation were available; excluding the 3 specimens for which TP53 mutation status or a CMS call was not available leaves 29, and further excluding the 2 specimens for which MDM2 copy number was not available leaves 27; Section 2). This is a limitation of the present analysis, and confounding by MSI has not been completely excluded. As for MDM2 copy number, in addition to showing no independent contribution in the same model (standardized partial regression coefficient = −0.011, P = 0.797), stratified analysis also yielded no evidence of confounding. Specifically, no rank correlation was observed between P2_index and total MDM2 copy number (Spearman ρ = −0.038, P = 0.467, n = 370), and binary stratification defining a total copy number of 5 or more as high-level amplification showed no difference in the median P2_index (18 high-level amplified specimens 0.406 versus 352 non-amplified specimens 0.429, Wilcoxon P = 0.949). The interaction between TP53 status and amplification was also not significant (P = 0.057), and the frequency of high-level amplification was 8 of 127 TP53 wild-type specimens (6.3%) and 10 of 237 mutant specimens (4.2%), with no skew between the two groups. Taken together, the conclusion that the observed associations are not explained by gene amplification—which in astrocytic tumors preferentially drives transcription from P1 [90]—was supported both by covariate adjustment and by stratification. It should be noted that P2_index is the promoter usage computed directly from the TPM of MDM2 transcripts (based on transcript-level estimated quantification) and that the effect sizes are moderate; nevertheless, the directionality with respect to TP53 status and to the p53 target and P2 programs was reproduced consistently with the mechanistic hypothesis. Overall, these results directly show, without recourse to the signatures, that MDM2 promoter usage itself is linked, with modest effect sizes, to TP53 status and to the p53 target and P2 programs in an independent cohort.

Because P2_index rests on transcript-level estimated quantification, we also counted promoter usage directly from splice-junction reads in the same tumors, without transcript-level estimation (recount3 [46]; P1, exon 1 to exon 3, chr12:68,808,492→68,809,207 and 68,808,481→68,809,207; P2, exon 2 to exon 3, 68,809,087→68,809,207; junction P2 share = P2/(P1 + P2) per tumor; Additional file 7: Supplementary Table S48, sheet D; code R117_S242_recount3_MDM2_junctions_v1.R and r117d_S242_crc_junction_v1.py, Supplementary Methods). TCGA-COAD/READ has few MDM2 junction reads (median 7 per tumor; interquartile range 4–12), so the per-tumor share is coarse, but it was again higher in TP53 wild-type tumors (medians 0.316 in 131 wild-type versus 0.000 in 231 mutant tumors with at least one read, Wilcoxon P = 2.8×10⁻¹⁶; in the 126 tumors with at least 10 reads, 0.392 versus 0.100, P = 2.6×10⁻¹²; shares pooled within each group, 0.375 versus 0.136). In the tumors with at least 10 reads, the junction P2 share agreed with P2_index (Spearman ρ = 0.64, P = 7.0×10⁻¹⁶; ρ = 0.23 over all tumors with at least one read). The difference between wild-type and mutant tumors therefore does not depend on transcript-level estimation. The junction reads were too few to test the correlation with the p53 activity score. In cell lines with full-length reads the same counting gave nearly the same ranking: in six SG-NEx cell lines sequenced with short and Nanopore long reads [58], the junction P2 share ranked the lines in nearly the same order as the DepMap P2 index (Spearman ρ = 0.94 with short reads and 0.83 with long reads; 1.00 for the cDNA-type long-read protocols), and over all SG-NEx long reads, reads beginning in exon 2 carried the exon 2→exon 3 junction (948 of 961), whereas those beginning in exon 1 carried the exon 1→exon 3 junction (2,131 of 2,289) (Supplementary Table S48, sheet G; code r137_sgnex_mdm2_v1.py and r142_sgnex_p2j_vs_ccle_v1.py, Supplementary Methods).

### R-13. Full-length version of main text Section 3.2.4 (MDM2 inhibitor sensitivity and MDM2 dependency)

##### TP53-dependent MDM2 inhibitor sensitivity and MDM2 dependency in cell line panels and organoids

Using wild-type TP53—a feature associated with the P2-dominant group—as a surrogate marker (stratification by MSI status is reported only as a background observation; MSI-H is not a proxy for P2, Section 3.2.2), we tested sensitivity to MDM2 inhibitors in cell line panels (an indirect test: TP53-dependent sensitivity to MDM2 inhibitors is well established, so these contrasts provide context rather than validation of promoter usage) (Additional file 30: Supplementary Figure S10: GDSC2 sensitivity; the statistical analysis code for this section is Code12, Supplementary Methods). We mechanically scanned the dose-response data of GDSC1 (378 drugs) and GDSC2 (286 drugs); only four compounds could be mapped as agents targeting the MDM2–p53 pathway: the MDM2 inhibitors nutlin-3a and Serdemetan, and the mutant p53 reactivators PRIMA-1MET and MIRA-1. Other MDM2 inhibitors for which clinical development has been attempted (RG7112, Idasanutlin, AMG-232 (= navtemadlin/KRT-232), Milademetan, HDM201, SAR405838, APG-115) are listed in neither GDSC1 nor GDSC2 and therefore could not be included in this validation. Cell line genotype information was unified to the DepMap 24Q4 Public release [52,53], and all values below were calculated on that same release.

For the MDM2 inhibitor nutlin-3a, TP53 wild-type cell lines were significantly more sensitive than mutant lines. In the pan-cancer setting, the median LN_IC50 was 2.672 in wild-type (n = 297) versus 5.167 in mutant (n = 638) (Mann–Whitney P = 1.5×10⁻⁶¹, Hodges–Lehmann estimator −2.383, 95% CI −2.622 to −2.135, Benjamini–Hochberg-adjusted FDR = 8.7×10⁻⁶¹), and independent evaluation of the same drug in GDSC1 gave the same direction (wild-type 1.981 (n = 282) versus mutant 4.101 (n = 618), P = 1.6×10⁻⁵⁸, Hodges–Lehmann −1.921, FDR = 4.7×10⁻⁵⁸). The difference was preserved when restricted to colorectal cancer cell lines: in GDSC2, wild-type 1.546 (n = 14) versus mutant 6.073 (n = 33) (P = 3.3×10⁻⁷, Hodges–Lehmann −4.210, FDR = 2.0×10⁻⁶), and in GDSC1, wild-type 2.188 (n = 12) versus mutant 4.612 (n = 32) (P = 3.7×10⁻⁵, FDR = 1.1×10⁻⁴). Serdemetan, an MDM2-binding agent with a different mechanism of action (Section 3.2.4 of the main text), was also significant in the same direction in both databases (pan-cancer: GDSC2 P = 2.9×10⁻⁴, GDSC1 P = 2.1×10⁻³; colorectal: GDSC2 P = 3.7×10⁻³, GDSC1 P = 6.9×10⁻³). For LN_IC50, smaller values indicate greater sensitivity.

We also constructed an integrated index combining the two drugs. LN_IC50 values were z-transformed for each database × drug combination, and their mean for each cell line was taken as the integrated sensitivity score. Integration was restricted to the two MDM2 inhibitors, which share a mechanism of action; the two reactivators, whose mechanism is the opposite, were not included. Of the 975 cell lines, 892 had all four measurements (2 drugs × 2 databases) available (44 of 47 colorectal cancer lines). With this score as well, TP53 wild-type lines were significantly more sensitive: pan-cancer median −0.555 (n = 299) versus +0.269 (n = 642) (P = 4.0×10⁻⁴⁶, Hodges–Lehmann −0.786, 95% CI −0.883 to −0.688), and colorectal −0.567 (n = 14) versus +0.839 (n = 33) (P = 4.3×10⁻⁶, Hodges–Lehmann −1.393, 95% CI −1.687 to −1.018). Note that this score averages multiple measurements obtained for the same cell line; it does not combine independent tests.

When colorectal cancer cell lines were compared by MSI status, both MDM2 inhibitors were significantly more sensitive on the MSI-High side (nutlin-3a: 2.489 in 15 MSI-High lines versus 5.862 in 32 MSS/MSI-L lines, P = 5.2×10⁻³, FDR = 1.5×10⁻², Hodges–Lehmann −3.048, 95% CI −4.028 to −0.551. Serdemetan: in GDSC2, 3.878 in 15 MSI-High lines versus 4.628 in 31 lines, P = 7.0×10⁻³, FDR = 1.5×10⁻²; in GDSC1, 3.091 in 14 MSI-High lines versus 3.710 in 30 lines, P = 7.3×10⁻³, FDR = 1.5×10⁻²). The integrated sensitivity score showed the same direction (−0.431 in 15 MSI-High lines versus +0.791 in 32 MSS/MSI-L lines, P = 6.5×10⁻⁴, Hodges–Lehmann −1.038, 95% CI −1.437 to −0.567). However, for nutlin-3a in GDSC1 the FDR did not fall below the threshold, and we did not detect a difference in the comparison by MSI status (2.683 in 14 lines versus 4.477 in 30 lines, P = 5.1×10⁻², FDR = 6.1×10⁻²). Among the reactivators, PRIMA-1MET showed greater sensitivity on the MSI-High side (4.659 in 15 lines versus 5.540 in 32 lines, P = 2.6×10⁻², FDR = 3.9×10⁻²), whereas for MIRA-1 we did not detect a difference (5.569 in 15 lines versus 6.035 in 31 lines, P = 0.26). The direction for PRIMA-1MET, namely greater sensitivity on the MSI-High side, where cases retaining wild-type TP53 are more frequent, is inconsistent with its presumed mechanism of reactivating mutant p53. Because these data cannot identify the reason, we do not use it as mechanistic support and record it only as an observation.

Independently of drug sensitivity, the same direction was confirmed for genetic dependency. In the DepMap CRISPR knockout screen (1,178 cell lines), the gene effect of MDM2 was significantly lower in TP53 wild-type cell lines, that is, dependency was higher. The median was −1.097 (n = 388) versus −0.321 (n = 790) in the pan-cancer setting (P = 8.9×10⁻⁹⁶, Hodges–Lehmann −0.758, 95% CI −0.809 to −0.705) and −1.389 (n = 12) versus −0.338 (n = 51) in colorectal cancer (P = 1.3×10⁻⁷, Hodges–Lehmann −1.024, 95% CI −1.223 to −0.733). For gene effect, smaller values indicate greater dependency. Whereas drug sensitivity is influenced by the properties of individual compounds, this analysis measures the requirement for the MDM2 gene itself, and it independently supports the interpretation that cells retaining wild-type p53 are more dependent on MDM2.

We add a note on the rules used to determine TP53 mutation status. In the DepMap 24Q4 release, 1,174 cell lines carried non-synonymous mutations in TP53, whereas 0 cell lines carried only synonymous mutations. Accordingly, the rule that treats cell lines with only synonymous mutations as wild-type and the rule that treats any mutation as mutant regardless of its type coincide completely in this release, and the results are unchanged whichever is adopted. In addition, the 176 cell lines for which no mutation profile was available were not defaulted to wild-type but were excluded from the analysis. Taken together, these large-scale cell line data are consistent with the established TP53-dependent sensitivity to MDM2 inhibitors and with our hypothesis that the P2-dominant group, which retains wild-type p53, is a candidate target population for these agents; they do not by themselves test promoter usage (GDSC [49,50]). Furthermore, in a biobank comprising 256 patient-derived tumor organoids (of which 162 completed genome-wide CRISPR screening, 85 of them colorectal), MDM2 dependency in colorectal organoids has been reported to be significantly higher in TP53 wild-type (effect size 0.463, adjusted P = 0.001), and HRAS and MDM2 gain-of-function events to correlate with nutlin-3a sensitivity (Pearson correlation coefficient 0.49; nutlin-3a is the name used in that report, whereas the drug-response data it released, which we re-analyze below, are labeled nutlin-3; see the note on nomenclature in Section 2) [13]. That is, the TP53 dependence described in this subsection is also observed in patient-derived models that are removed from cell lines adapted to culture. However, the same report also observed MDM2 dependency in colorectal organoids carrying the KRAS G12D mutation (effect size −0.576, adjusted P = 0.01; 4 of the 5 relevant cases are TP53 wild-type), so MDM2 dependency is not a property specific to TP53 wild-type. A list of all contrasts is shown in Additional file 31: Supplementary Table S38. In this study, we independently reanalyzed the primary data of that report. Among the colorectal organoids released by that biobank, in the 65 lines for which both the log(IC50) of nutlin-3 and somatic mutation data were available, the median log(IC50) was 1.386 in the 17 TP53 wild-type lines versus 4.283 in the 48 mutant lines, indicating that wild-type lines were significantly more sensitive (exact Wilcoxon rank-sum test P = 7.4×10⁻⁸, Cliff's delta = −0.804). The direction and magnitude were preserved when KRAS mutation, MSI status, and MDM2 gain-of-function mutation were added sequentially (regression coefficient for wild-type β = −2.515 (univariable) to −1.959 (after adjustment for the three covariates), P = 7.8×10⁻¹¹ to 3.3×10⁻⁸). When TP53 was split by truncating mutations alone, the difference was blunted (3.986 in the 51 non-LOF lines versus 4.445 in the 14 LOF lines, P = 0.022 by normal approximation), which we interpret as reflecting the admixture of missense-mutant lines on the non-LOF side. Furthermore, our P2 signature score showed an association in these 65 lines whereby higher scores corresponded to greater nutlin-3 sensitivity (p53 target module excluding the MDM2 gene: Spearman ρ = −0.539, exact Spearman test P = 5.3×10⁻⁶, Benjamini–Hochberg-adjusted FDR = 5.1×10⁻⁵, family = 13). A partial correlation adjusted for TP53 status retained an association in the same direction, but it did not fall below the significance threshold after multiple-testing correction (ρ = −0.331, raw P = 0.0071, BH-adjusted FDR = 0.056). That is, the P2 axis constructed in the discovery cohort (63 organoid samples from 21 patients) was associated with the efficacy of MDM2 inhibitors in the 65 patient-derived organoids of Herranz-Ors et al., but within these 65 organoids this association could not be shown to be independent of TP53 mutation status. In GDSC2 cell lines, by contrast, the association of the P2 score with measured nutlin-3a sensitivity persisted after adjustment for cancer type and TP53 status (partial ρ = −0.179, P = 6.5×10⁻⁷; Section 3.2.4), although the effect was small (Section 4.6). Per-line values are shown in Additional file 32: Supplementary Table S40, and the correlations for each signature in Additional file 33: Supplementary Table S41.

### R-14. Full-length version of main text Section 3.3.3 (Immune microenvironment and ICI response prediction)

##### The immune microenvironment of the P2-dominant group and prediction of response to immune checkpoint inhibitors

We tested, in TCGA-COAD/READ (624 specimens on a per-specimen basis), the immune microenvironment predicted from the wild-type TP53 status and MSI-like signature features that characterize the P2-dominant group. Stratification used tertiles of the P2 signature score, and we compared the top 208 specimens (P2-high group) with the bottom 208 specimens (P2-low group) (the middle tertile was excluded from the between-group comparison, and correlations with the continuous axis were calculated in the 624 specimens). Deconvolution was performed with two methods, quanTIseq and MCP-counter, and only cell types whose signs agreed between the two methods were treated as findings. Testing was by Mann–Whitney (normal approximation with continuity and tie corrections), the effect size was the Hodges–Lehmann estimator, and multiplicity was handled by within-method Benjamini–Hochberg correction (Additional file 55: Supplementary Table S35). In the P2-high group, NK cells (MCP-counter: Hodges–Lehmann = +0.210, P = 2.0×10⁻²¹), the cytotoxicity score (+0.432, P = 1.5×10⁻²²), B cells (+0.507, P = 2.3×10⁻¹⁰), and T cells (+0.357, P = 1.5×10⁻¹⁴) were significantly higher. The CD8-positive T cell fraction was significantly higher by quanTIseq (+0.0014, P = 1.3×10⁻¹⁰), whereas with MCP-counter the sign was in the same direction but we did not detect a significant difference (P = 0.121). Note that for neutrophils the signs did not agree between the two methods, and for myeloid dendritic cells (quanTIseq) the median was 0 in both the high and low groups so that no direction could be determined; neither is treated as a finding. Because the MCP-counter Macrophage/Monocyte and Monocyte estimates were identical in all 624 specimens, one of them was removed from the family for multiple comparisons. xCell and EPIC failed to run, so this analysis is based on those two methods. The analysis script is deposited as Code13 (Supplementary Methods).

To confirm that these findings are not a restatement of MSI status, we performed a sensitivity analysis stratified by MSI in the 272 specimens with MSI annotation available (38 MSI-High, 234 MSS/MSI-Low) (Additional file 56: Supplementary Table S36). NK cells, the cytotoxicity score, B cells, and T cells (MCP-counter) and CD8-positive T cells (quanTIseq) were significant in the van Elteren test stratified by MSI (P = 3.1×10⁻⁵ to 1.1×10⁻³), in tests restricted to MSS/MSI-Low (P = 6.3×10⁻⁵ to 2.0×10⁻³), and in partial Spearman correlations controlling for the MSI-High indicator variable (ρ = +0.179 to +0.284, P = 3.1×10⁻³ to 2.0×10⁻⁶). Therefore, the finding that immune cell infiltration and cytotoxicity are high in P2 is not a restatement of MSI status. Note that the coverage of MSI annotation differs among analysis pipelines. The adjusted Cox model in Section 3.3.2 (252 of 591 cases) obtained MSI annotation from the subtype table returned by TCGAquery_subtype in TCGAbiolinks, the 272 specimens in this section from the paper_MSI_status column of the GDC clinical table, and the univariable comparison in Section 3.2.2 (32 of 380 specimens) from the MSI column of the input covariate table for Code9 (generated by Code16); each analysis was restricted to specimens with MSI annotation available.

By contrast, prediction of response to immune checkpoint inhibitors (ICIs) cannot be claimed with the same strength. In response prediction by TIDE (TIDEpy 1.3.8), across all 624 specimens the predicted Responder rate was 51.0% (106/208) in the P2-high group, higher than the 31.7% (66/208) in the P2-low group (Fisher exact test P = 9.8×10⁻⁵, odds ratio 2.24). However, when restricted to the 173 top- and bottom-tertile specimens with MSI annotation available, the rates were 53.7% (66/123) versus 48.0% (24/50), and we did not detect a significant difference (odds ratio 1.25, P = 0.50). This loss is not explained by MSI confounding. In a logistic regression that added MSI-High as a covariate, the odds ratio for P2-high in fact increased (1.61, 95% CI 0.81–3.20, P = 0.18); the effect had already become small at the point of restricting to the subset with MSI annotation, before adjustment. Indeed, the presence or absence of MSI annotation is strongly associated with the P2 score: MSI was known in 24.0% (50/208) of the P2-low group, 47.6% (99/208) of the middle tertile, and 59.1% (123/208) of the P2-high group. We therefore treat the TIDE response prediction as an exploratory finding that is not reproduced in the population with MSI annotation, and this study does not claim that P2 is an indicated population for ICIs.

Two features of these data do not permit a simple interpretation. First, the TIDE score is lower on the P2-high side (the direction in which response is expected; Hodges–Lehmann = −0.412, P = 9.9×10⁻⁶), whereas its component T cell dysfunction (Dysfunction) is higher on the P2-high side (+0.864, P = 6.1×10⁻²¹). TIDE is designed to switch between weighting dysfunction and exclusion according to the level of cytotoxic T cells, and on the P2-high side the low values of T cell exclusion (−0.761), myeloid-derived suppressor cells (−0.061), and M2 tumor-associated macrophages (−0.035) all contribute to the low TIDE score. The interpretation that response is expected because dysfunction is low is therefore not available. Second, the TIDE-predicted response rate was lower in MSI-High (34.2%, 13/38) than in MSS/MSI-Low (54.7%, 128/234), which is the opposite direction from the clinical knowledge that MSI-High colorectal cancer responds well to ICIs. The input to TIDE is only the pre-treatment tumor expression profile; MSI status, tumor mutational burden, and PD-L1 expression are all absent from the input. TIDE is an index that combines a T cell dysfunction signature, derived from genes whose association with survival is attenuated under cytotoxic T cell infiltration, with a T cell exclusion signature derived from immunosuppressive cells, namely cancer-associated fibroblasts, myeloid-derived suppressor cells, and M2 macrophages. The original paper [64] showed that in a first-line melanoma cohort it predicted response more accurately than PD-L1 expression level or tumor mutational burden. That is, TIDE is not designed to substitute for or reproduce the clinical indication criterion of MSI-High. These two points indicate that the findings of this section cannot be extrapolated to decisions about ICI indication. Note that the TIDE web server (tide.dfci.harvard.edu) has been shut down (the successor site, the Cancer Immunology Data Engine, carries a TIDE Server Retired notice), and in this study we used TIDEpy (1.3.8), the Python implementation released under GPLv3 by the same group. Because TIDEpy is not a reimplementation of the method but the original implementation itself and has no citation of its own, citation is by two reports, the original paper by Jiang et al. [64] and the TIDE platform paper by Fu et al. [65] (bibliographic details in Supplementary Results 13).

A note on the definition of stratification. The stratification used in this section (tertiles of the P2 signature score) is not the only possible axis. Tertiles of the GSVA difference P2 − P1, an axis used in the companion study of the 15q11-q13 locus (T. Tsukui, R. Yao, and K. Tsuda, unpublished observations), rank the same specimens similarly, but the membership of the middle tertile differs considerably between the two axes. Each stratification is internally consistent, but when this report is read side by side with a report that uses the P2 − P1 axis, P2-high cannot be read as referring to the same set of specimens.

### R-15. Full-length version of main text Section 3.2.2 (TP53 functional class and MDM2 P2 promoter usage)

##### TP53 functional class and MDM2 P2 promoter usage

In Section 3.2.2 we treated TP53 as a binary variable (mutated / not mutated). To increase this resolution, we used the mutation annotation of the TP53 Database (formerly the IARC TP53 Database; now maintained by the US National Cancer Institute; release 21) [66,67] to test whether P2_index and p53 target module output are ordered in a stepwise manner by the functional class of the somatic mutation (TCGA-COAD/READ, 374 specimens for which both P2_index and TP53 mutation information were available, using the same exclusion rules as in Section 3.2.2; the analysis code is Code14, Supplementary Methods). The classification rules were declared before the analysis. Wild-type (WT) specimens are those in which no somatic mutation in TP53 was detected in the Masked Somatic Mutation data, and loss-of-function (LOF) specimens are those with truncating mutations such as nonsense, frameshift, and splice-site mutations. Here too, by the same rule as in Sections 3.2.2 and 3.3.2, non-detection of a mutation is not distinguished from true wild-type, and specimens for which no somatic mutation data were available were not treated as wild-type but were excluded from the analysis.

For gain-of-function (GOF), because the operational definition in the literature is not settled on a single form, we designed the analysis to run two definitions in parallel and to check whether the conclusion depends on the definition. Definition A is the conservative definition, in which only missense mutations at the 6 hotspot residues whose mutation frequency in human cancers is outstandingly high and for which gain of function has been repeatedly discussed (R175H, G245S, R248Q/W, R249S, R273H/C, R282W) [68] are counted as GOF. Definition B is the broader definition, which includes on the GOF side all missense mutations whose transactivation class in the TP53 Database is non-functional. This classification derives from a high-resolution missense mutation analysis in yeast that comprehensively measured the transactivation capacity of p53 mutants [69]. It should be noted that what definition B captures directly is missense mutations that have lost transcriptional activity, not gain of function itself. Under either definition the determination of WT and LOF is identical; only the boundary of GOF differs. Missense mutations not qualifying as GOF were excluded from the three-class comparison. The breakdown of the 374 target specimens is 132 wild-type, 68 truncating, and 169 missense (93 hotspot, 76 non-hotspot), 369 in total; the remaining 5 specimens are mutants that are neither truncating nor missense and are therefore outside the three-class comparison under either definition. Definition A yielded 293 specimens across the three classes (WT 132, LOF 68, GOF 93), and definition B yielded 358 (WT 132, LOF 68, GOF 158). When we checked the agreement of the two definitions specimen by specimen, definition B moved into GOF 65 of the 76 non-hotspot missense specimens that definition A left unclassified, and there was not a single specimen in which the GOF and LOF assignments were interchanged.

P2_index differed significantly among the three classes (Kruskal–Wallis, definition A P = 1.3×10⁻⁵, definition B P = 3.9×10⁻⁶), with medians in the order wild-type 0.481, GOF 0.423 (definition A) / 0.422 (definition B), and LOF 0.364. With the three prespecified contrasts and Benjamini–Hochberg correction, under definition A wild-type versus LOF (Hodges–Lehmann estimator +0.126, P = 1.7×10⁻⁶, FDR = 5.0×10⁻⁶), wild-type versus GOF (+0.053, P = 0.030, FDR = 0.030), and LOF versus GOF (−0.069, P = 0.015, FDR = 0.022) were all significant. Definition B gave the same directions: wild-type versus LOF (+0.126, P = 1.7×10⁻⁶), wild-type versus GOF (+0.065, P = 1.6×10⁻³, FDR = 2.4×10⁻³), and LOF versus GOF (−0.060, P = 0.013). That is, P2_index shows a gradient that is lowest in truncating mutants, intermediate in missense mutants, and highest in wild-type, and this ordering was preserved under both GOF definitions.

By contrast, p53 target module output differed markedly between wild-type and mutant, but showed no step within the mutant group. The medians are wild-type +0.487, LOF −0.400, and GOF −0.397 (definition A) / −0.437 (definition B). Wild-type versus LOF (Hodges–Lehmann +0.723, P = 5.9×10⁻¹⁷) and wild-type versus GOF (definition A +0.780, P = 2.4×10⁻²²; definition B +0.806, P = 1.3×10⁻²⁸) were all strongly significant, whereas for LOF versus GOF we did not detect a significant difference (definition A +0.030, P = 0.52; definition B +0.046, P = 0.24). Note that 4 of the P values returned by the analysis software had been recorded as 0 because of numerical underflow, so the values given above are those recalculated from the specimen-level data by Mann–Whitney (normal approximation with continuity and tie corrections). All recalculated values are larger than the minimum attainable P that could be reached if complete separation occurred at the corresponding sample size (for example, 1.1×10⁻⁴⁸ for n = 132 versus 158), which is consistent with the attainable range of the test.

These results indicate that P2_index carries information that is not explained by p53 target pathway output alone, because the p53 target module does not distinguish within the mutant group whereas P2_index distinguishes truncating from missense mutations. However, this finding cannot be interpreted as a GOF-specific effect. Comparing hotspot (93 specimens) and non-hotspot (76 specimens) missense mutations within the missense group, we did not detect a significant difference (P2_index P = 0.365, p53 target module P = 0.316), and, as noted above, widening the boundary of GOF does not change the conclusion. What is observed is a gradient of truncating < missense < wild-type, not an effect specific to gain of function. It is possible that the loss of the p53 protein itself in truncating mutants, as against the persistence of a full-length protein lacking transcriptional activity in missense mutants, corresponds to the intermediate level of P2 promoter usage, but these data cannot identify this mechanism. The results are shown in Additional file 29: Supplementary Table S37.

### R-16. Full-length version of main text Sections 3.2.4 and 3.3.3 (Drug-sensitivity prediction; CMS marker overlap)

##### Prediction of drug sensitivity from cell line pharmacogenomics

In Section 3.2.4 we compared cell line sensitivity for four drugs selected in advance. Because this view extracts only the drugs consistent with the hypothesis, it cannot tell us whether the characteristics of the P2-dominant group emerge independently of how the drugs are chosen. We therefore used oncoPredict [57], which projects sensitivity from cell line panels onto patient specimens, to predict the sensitivity of the 624 TCGA-COAD/READ specimens for all 198 drugs listed in GDSC2, and obtained the correlation with the P2 score for each drug (the analysis code is Code15, Supplementary Methods). Training used 805 cell lines × 198 drugs from GDSC2, and the expression matrix was aggregated per specimen under the same conventions as in Section 3.3.2.

Of the 198 drugs, the one most strongly correlated with the P2 score was nutlin-3a (Spearman ρ = −0.659, P = 4.2×10⁻⁷⁹, Benjamini–Hochberg-adjusted FDR = 8.4×10⁻⁷⁷, n = 624). A negative correlation means that the higher the score, the lower the predicted IC50, that is, the greater the sensitivity. Nutlin-3a ranked first among these drugs in ascending order of ρ, and it correlated in the opposite direction with the P1 score (ρ = +0.107, FDR = 1.4×10⁻²). Nutlin-3a also ranked first on a separation index that quantifies the degree to which the directions diverge between the P2 and P1 scores (0.766). The predicted nutlin-3a sensitivity also differed across CMS classes (Kruskal–Wallis P = 3.4×10⁻²¹; the 563 specimens with a CMS call available from CMScaller (CMS1 96, CMS2 167, CMS3 96, CMS4 204); this set is identical to the per-specimen CMS calls used in Section 3.3.2 and Supplementary Results 11), with the median predicted IC50 lowest in CMS1 (48.2) and highest in CMS2 (183.3). Of the 198 drugs, 195 reached FDR < 0.05 across CMS classes.

This analysis has an important limitation. Of the 198 drugs, 171 reached FDR < 0.05 in the direction of higher P2 score corresponding to greater sensitivity. That is, the P2 score correlates not with particular drugs but with drug sensitivity in general, and it is likely to include general axes such as proliferation and differentiation. Therefore, the smallness of the P value obtained for nutlin-3a cannot in itself be read as evidence specific to MDM2 inhibitors. What should be used for interpretation is the rank. That the correlation is the strongest across all drugs, and that the degree to which the directions diverge between the P2 and P1 scores is the largest, are not explained by a general gradient of sensitivity alone. Moreover, because the P2 signature contains canonical p53 target genes (e.g., CDKN1A, BAX, BBC3), a top rank for nutlin-3a is expected from its composition and is not by itself independent evidence for promoter usage; the TP53-adjusted analyses below address this. Note that among the MDM2 inhibitors, Serdemetan is not included in the oncoPredict training set (the 198 drugs of GDSC2), so the MDM2–p53 pathway drugs that could be evaluated in this analysis were the three drugs nutlin-3a, PRIMA-1MET, and MIRA-1. Predicted values were computed in 10 batches (nine batches of 20 drugs and one of 18), but the maximum absolute difference from results computed for a single drug alone was 0, so the batching did not affect the results. The full results for the 198 drugs are shown in Additional file 34: Supplementary Table S39. Because oncoPredict yields predictions rather than measurements, the same question was then tested on measured data (R47/R47b, Supplementary Methods): P1 and P2 scores computed from GDSC2 cell-line expression were correlated with measured LN_IC50 for all 295 GDSC2 drug entries (DRUG_IDs; they correspond to the 286 GDSC2 drug names of Section 2, some drugs being represented by two entries; up to 763 lines), adjusting for cancer type and TP53 status. Nutlin-3a again ranked first of 295 on the P2 score (partial ρ = −0.371 adjusted for cancer type; −0.179 after further adjustment for TP53, P = 6.5×10⁻⁷), and in DepMap CRISPR screens the P2 score tracked MDM2 dependency independently of lineage and TP53 (partial ρ = −0.165, P = 2.2×10⁻⁴). On the P1 side, all seven inhibitors of the replication-stress checkpoint kinases ATR, CHK1 and WEE1 showed higher sensitivity with higher P1 scores (median partial ρ = −0.077 versus approximately 0 for other drugs; P = 4.3×10⁻⁷), a direction compatible with the proposed reliance of TP53-deficient, chromosomally unstable cells on the G2/M checkpoint (not tested here); EGFR inhibitors showed only a weak shift shared by both scores, and 5-FU, oxaliplatin and irinotecan showed no P1-specific association. MEK/ERK/BRAF inhibitors unexpectedly correlated with the P2 score (median partial ρ = −0.125). Although the P2 score was itself higher in RAS- and BRAF-mutant lines, this association was not explained by the mutation spectrum: for the MEK inhibitors, dabrafenib and the ERK inhibitor SCH772984 it persisted after further adjustment for RAS and BRAF mutation status (R47c; e.g., trametinib partial ρ = −0.123, P = 6.8×10⁻⁴) and within RAS/RAF wild-type lines (trametinib −0.122, n = 513), whereas for PLX-4720 and the other ERK inhibitors it did not; under this adjustment nutlin-3a remained first (partial ρ = −0.178, P = 7.9×10⁻⁷) and the P1 association of the checkpoint inhibitors was unchanged. These cell-line effects are small, the 44 colorectal lines with expression data were too few on their own for significance, and the only drug-response data released for the organoid biobank [13] are for nutlin-3; these results are exploratory (Additional file 34: Supplementary Table S39, sheets S39b–S39d).

##### Overlap test between DEGs and CMS class-specific marker genes

Using the one-sided Fisher exact test described in Section 2 (background of 26,278 genes; the 8 tests from 4 classes × 2 directions were adjusted by the Benjamini–Hochberg procedure), we evaluated the overlap between the DEGs of this study (Supplementary Table S4; 1,013 genes on the tissue-1 side and 877 genes on the tissue-2 side) and CMS class-specific marker genes. When the reference sets were the top 200 genes per class derived empirically from the 563 CMS-classified TCGA-COAD/READ specimens, the tissue-1 side DEGs were specifically enriched in CMS2 markers (OR = 3.92, BH-FDR = 8.7×10⁻⁷), were not significant for the other three classes, and had 0 genes overlapping with CMS4 markers. The tissue-2 side DEGs showed the strongest enrichment in CMS3 (metabolic) markers (OR = 5.91, BH-FDR = 2.0×10⁻¹¹) and were also significantly enriched in CMS1 (MSI immune) markers (OR = 2.63, BH-FDR = 3.2×10⁻³). The 28 genes overlapping on the CMS3 side are dominated by secretory, mucinous, and metabolic systems. When the reference sets were replaced by the CMS templates of CMScaller, enrichment of the tissue-2 side DEGs in CMS3 was again the strongest among the four classes (OR = 18.21), whereas the tissue-1 side DEGs overlapped the CMS1–CMS3 templates to a similar degree (OR = 3.09–3.77), and the CMS2 template was overlapped more strongly by the tissue-2-side (OR = 9.68) than by the tissue-1-side DEGs (OR = 3.75). The full results are shown in Supplementary Table S28. That is, separately from the assignment at the score (pathway) level (Section 3.3.2), at the gene level as well the P1-dominant group corresponds to CMS2, and the P2-dominant group corresponds to CMS3 as its center while also carrying CMS1 (Section 4.1 and Supplementary Report Section 4.2.2). Removing from the query the P2 signature genes and a metaplasia/intestinal marker panel (47 genes after removing duplicates) hardly attenuated the CMS3 enrichment (OR = 5.91 → 5.30; BH-FDR = 2.0×10⁻¹¹ → 2.0×10⁻⁹), so it is not an artifact of the signature definitions (Supplementary Note Section 5.2). In the exploratory colibactin analysis (Section 2), SBS88 and ID18 activities were only slightly shifted toward the P2-dominant side of the P1−P2 GSVA axis in the Nunes cohort (weak correlations; Spearman ρ = −0.085 and −0.104, n = 1,051; retained in MSS tumors), whereas in TCGA-COAD/READ only P2_index × ID18 was significant in the same direction (ρ = +0.143, FDR = 0.044, n = 374) (Section 4.6).

### R-17. Full-length version of main text Section 4.1 (TP53 status, morphology and MDM2 isoforms)

#### 4.1 TP53 status, morphology and MDM2 isoforms: the central finding of this study

The isoform difference is supported by three concordant observations from the 33 comparisons: (1) across the 33 Splicing Index comparisons, exon 1 (representative probe PSR12007998) was enriched on the tissue-1 side and exon 2 (representative probe JUC12004243) on the tissue-2 side in every comparison (values, and the exceptions for individual probes, in Section 3.1.2 and Figure 1B); (2) an approximately 7-fold elevation of MDM2 mRNA expression in tissue 2 (mean linear fold change 7.0-fold, median 6.0-fold, in the same direction in all 33 comparisons; Table 4); and (3) Upstream analysis identified wild-type TP53 as a top-class activator on the tissue-2 side (median z = −7.14, sign consistency 100%), and EGR1 (a transcription factor that can regulate MDM2 expression in a context-dependent manner [100]) also showed high sign consistency in the same direction (median z = −1.95, sign consistency 91%; the effect size was below the adopted threshold) (details in Supplementary Note Section 3.2). In the discovery organoids (21 patients, 63 specimens), we first found that morphology corresponded to MDM2 promoter usage specimen by specimen, but TP53 status coincided with promoter usage (40 of the 42 mutant specimens were P1-dominant), and when the two were compared, morphology followed TP53 status to a degree similar to promoter usage, three of the four morphology-deviating specimens followed TP53 status rather than promoter usage, and within the narrow range in which the two axes disagreed (five specimens), P2 dominance added no detectable contribution to morphology beyond TP53 status (a post hoc test with little power, added after the per-specimen TP53 calls had been finalized; Sections 3.1.3 and 3.1.4 of the main text); like the morphology–isoform association, the TP53–morphology association in this cohort was established at the specimen level (patient-level sensitivity analysis in Section R-26). In an independent biobank (GSE294511 [36]; 88 organoids with published TP53 status), where it held on patient means, and in TCGA tumors, the morphology genes again followed TP53 status beyond 20q, and the across-sample alignment of morphology with P2 in that biobank was largely explained by TP53 status (Section 3.1.5 of the main text). We therefore interpret the central finding as reflecting the alignment of morphology with TP53 status: TP53-mutant organoids tend to be compact and glandular with chromosomal-instability expression features, and TP53 wild-type organoids round (cystic–mucinous), with gastric-metaplastic genes following TP53 status in the independent biobank and in tumors, while MDM2 P2 usage, which tracks p53 activity (the evidence that it does so beyond genotype resting mainly on the independent biobank and two TP53-mutant P2-dominant specimens; Sections 3.1.4 and 3.2 of the main text), reflects the p53 state rather than determining morphology. Within TCGA wild-type tumors the morphology genes were unrelated to p53 activity (ρ = −0.08, P = 0.37; 119 tumors; Supplementary Table S48, sheet H2), so morphology follows TP53 genotype, or the pathway it marks, rather than p53 function. Whether TP53 loss itself drives the morphological change was not tested, and TP53 mutation is entangled with chromosomal instability, although adjustment for 20q, aneuploidy and MSI (and, in tumors, BRAF mutation and CIMP) did not remove the association (among non-MSI, BRAF wild-type, CIMP-low tumors, however, the 20q-adjusted morphology difference was only marginal; Section R-26). Because it rests on molecular axes (TP53 status and MDM2 P1/P2 isoform usage) that are independent of the morphological typing of Okamoto et al. [12], who analyzed an overlapping patient group from the same biobank, the observation is not a mere reproduction of that earlier study but a complementary new finding (Section 4.4). Furthermore, the DEG lists of this study overlapped significantly with the CMS class-specific markers of Guinney et al. (2015) [2] (the tissue-1-side DEGs were enriched in CMS2 (OR = 3.92), whereas the tissue-2-side DEGs were most strongly enriched in CMS3 (OR = 5.91) and were also significantly enriched in CMS1 (OR = 2.63); details in Section 3.3.3 and Supplementary Table S28), showing that the present findings are consistent with published molecular classifications. In addition, external analyses using TCGA (n = 624) and GSE39582 (n = 519) reproduced the CMS and mismatch-repair associations of the signatures, whereas the P1 score was not associated with prognosis independently of stage (details in Section 3.3.2). A note on the interpretation of morphology is warranted here. In the initial application (before unification of the preprocessing), the specimens separated into a Type1-dominant and a Type5-dominant group; under re-classification with training-matched preprocessing (Section 3.1.2; Supplementary Table S46), the images on the cystic side were assigned across the two classes Type0 and Type5, and the images of P2-dominant specimens assigned to Type0 showed, on visual inspection, cystic morphology close to Type5. This similarity is not merely apparent. When the P2-dominant specimens were divided into those whose Type0 fraction exceeded their Type5 fraction (7 specimens) and those with the reverse (5 specimens; HCT41-1T, with neither class, was excluded) and their expression profiles were compared, no molecularly distinct subgroup was detected: the two sides were intermixed within specimens of the same patients (HCT27 and HCT33); they did not coincide with the sub-clusters within P2 defined by the 39-gene panel; there was no consistent difference in the principal indices including the gastric metaplasia markers (CTSE, REN, TFF1) and the EMT score (of 10 indices, only LGR5 had nominal P < 0.05, which was not significant after correction for multiple comparisons); and in a genome-wide search the number of nominally different genes (28) was within the null distribution obtained by permuting the group labels (an exploratory analysis with a small number of specimens). That is, the Type0-leaning and Type5-leaning specimens are not distinguished — not only in appearance but also in molecular-biological state — and can be treated as morphological variation within a single P2 program. Type0 is the class that the 6-morphology classification of Okamoto et al. defined as a differentiated type with a single lumen, and the fact that the morphology of the P2 specimens of this cohort is assigned across the two classes Type5 and Type0 can be interpreted coherently as a morphological manifestation of the differentiated, secretory traits of the P2 group (gastric metaplasia and mucus production; Section 4.2). The core of the correspondence is thus the separation into two groups — a P1 group dominated by the compact glandular type (Type1) and a P2 group lacking Type1 dominance and dominated by round (cystic–mucinous; Type0/Type5) morphology — and this separation does not depend on the choice of image preprocessing (pooled non-Type1 fraction: 0.33 versus 0.30 in P1 specimens and 0.62 versus 0.59 in P2 specimens [before versus after unification]). This two-group structure does not negate the 6-morphology typing of the same biobank (above); rather, it indicates that behind that morphological diversity there exists a higher-level two-group structure that aligns with TP53 status (Type1-leaning in TP53-mutant and non-Type1 in TP53 wild-type specimens), and the fine classification (6 types) and the coarse classification (2 groups) form complementary hierarchies (Section 4.4). On the TP53 wild-type, largely P2-dominant side, morphology is best treated not as the single class Type5 but as a non-Type1 (round morphology) spectrum.

### R-18. Full-length version of main text Sections 4.2 and 4.2.1 (MDM2 P2 usage as a readout of p53 activity; subsections 4.2.1–4.2.3 below follow the full-length numbering and are condensed into main text Sections 4.2 and 4.2.1, subsection 4.2.2 below corresponding to main text Section 4.2.1)

#### 4.2 MDM2 P2 usage as a readout of p53 activity: a working model

Our data support the change in MDM2 isoforms as a consequence and readout of p53 activity (Sections 3.1.4 and 3.2 of the main text); whether promoter usage itself feeds back on cellular programs was not tested (the upstream and downstream factors of the two tissue types are contrasted in Additional file 57: Supplementary Table S44). In tissue 1 (P1-dominant), mutation or loss of the TP53 gene accompanying progression of the CIN pathway (or p53 inactivity in wild-type cases; Section 3.2.1 of the main text) would be expected to abolish p53-dependent P2 activation (consequence), leaving MDM2 predominantly transcribed from P1, while the MYC/FOXM1/E2F proliferative program and glandular architecture accompany TP53 status and chromosomal instability rather than promoter usage (Section 3.1.4 of the main text). Public nascent-transcription data qualify this picture. In ENCODE PRO-cap (signal tracks from the ENCODE portal [39,40], the Encyclopedia of DNA Elements, a public functional-genomics resource), which maps the start sites of capped nascent RNA at single-nucleotide resolution [174], P2 accounted for about half of MDM2 transcription initiation in six normal colon and stomach samples (median 0.506; range 0.421–0.657) and for 0.565 in Caco-2 cells, which carry a homozygous truncating TP53 mutation (Cellosaurus [41]), whereas P2 accounted for only 0.173 of MDM2 mRNA in the same line in DepMap (a different dataset; Additional file 7: sheet K). Basal initiation at P2 therefore does not appear to require p53; beyond inducing P2, p53 may also act by turning P2 initiation into productive, mRNA-yielding transcription. Because PRO-cap also captures paused and unstable transcripts, and only normal tissue and one p53-deficient line could be examined, this remains a hypothesis. In tissue 2 (P2-dominant), wild-type p53 is active and induces the P2 promoter (consequence); chronic inflammation (TNF, IL1B), DNA damage, and oxidative stress are possible triggers suggested by expression and IPA prediction (not measured). In this working model, P2 activation co-occurs with suppression of p53-induced apoptosis and with a complex lineage conversion comprising TGFβ/SMAD3-mediated partial EMT and activation of a gastric-type transcriptional program that follows TP53 status, whose temporal order relative to P2 induction is not established (Section 4.3). Regarding the EMT that arises in this process, Nieto et al. (2016) [105] have shown that EMT proceeds not as a dichotomous conversion but as multiple dynamic intermediate states between the epithelial and mesenchymal types. The high expression of the invasion markers MMP7 (100% consistent), MSLN, and PLAUR observed in tissue 2 is consistent in direction with this EMT activation. This activation of the mesenchymal program is compatible with the expression data of this study: in a comparison in which the EMT scores of all 63 specimens were matched at the patient level, metastatic specimens showed significantly higher EMT scores than primary specimens (metastasis median +0.130 versus primary median −0.590 across all 63 specimens, per-specimen Wilcoxon rank-sum test P = 0.014; patient-level paired Wilcoxon signed-rank test P = 0.004, 17 patients, two-sided), whereas the difference between P2 and P1 cases was not significant (P = 0.079; Section 3.1.2 and Supplementary Table S1). This indicates that the literature findings described above agree in direction with the quantitative observations in the present cohort (although this is an exploratory finding based on a small number of cases). Furthermore, as Mani et al. showed [109], EMT confers stem cell-like properties on cancer cells. The acquisition of gastric metaplasia observed in tissue 2 coexists within the same subtype with the plasticization of cell identity induced by EMT, and the apoptosis evasion associated with MDM2 P2 induction is hypothesized to support the cell survival required for this process (Section 4.3). In the working model of this study, MDM2 promoter usage is positioned as a readout of p53 activity along the mutation-accumulation pathway of cancer evolution. However, according to the integrated model of this study (Supplementary Report Section 4.2.2), P1/P2 choice is interpreted not as a fixed, irreversible state but as a context-dependent output that is specified, with upstream TP53 status and p53 activation as the branch point, according to the balance between the genomic background (TP53 mutation or deletion) and the metastatic microenvironment (chronic inflammation, DNA damage, oxidative stress). From this viewpoint, the P1 (primary) → P2 (metastasis) pattern observed in patients HCT38 and HCT67 in this study (HCT41, whose primary specimen was P2-dominant, is addressed below) can also be understood as a condition-dependent transition in which P2 was induced by the addition of environmental signals that activate the P2 promoter during the metastatic process (signals that may act through active p53; the two P2-dominant specimens that carried TP53 mutations were also p53-active [Section 3.2.1 of the main text], and a non-p53-dependent contribution via EGR1 or other factors is speculative and was not tested). It should be noted that, because this study is based on cross-sectional observations comparing primary tumors and metastases, although the direction of the transition observed in the two patients for whom directionality could be assigned (HCT38 and HCT67) was P1→P2 (of the three patients with both P1- and P2-dominant specimens, HCT41 was not subjected to directional assignment because its primary specimen HCT41-1T showed a borderline Splicing Index value with weak exon 2 dominance; Section 3.1.2), this does not establish that this directionality is unidirectional and irreversible (that is, that the reverse P2→P1 transition cannot occur); determining the reversibility and directionality of promoter choice will require longitudinal sampling or validation by functional perturbation (Section 4.6).

##### 4.2.1 Mechanistic support from TP53 mutation status

The TP53 targeted resequencing analysis of this study provided genomic-level support consistent with the working model of Section 4.2, in which TP53 status determines whether p53-dependent P2 activation can occur. TP53 mutations (in the samples with variant tables, pathogenic mutations such as R273H, R248W, R175H, R213*, and S127F) were found in 15 of the 17 patients with tissue 1 (P1-dominant) whose status was determined (the remaining two, HCT59 and HCT71, retained wild-type TP53, indicating that TP53 mutation is closely associated with the P1 phenotype but not a prerequisite for it). All of those with a known amino-acid change are loss-of-function or dominant-negative mutations with respect to wild-type transactivation (the map-derived mutations were reported only as truncating or non-truncating), as is typical of the CIN pathway of colorectal cancer [103] (the hotspot missense variants R175H, R248W, and R273H have also been reported to exert gain-of-function activities [103]), and they are expected to render the p53 protein unable to bind to the p53RE on the P2 promoter of MDM2, so that transcription of MDM2 mRNA would depend mainly on constitutive, basal expression from the P1 promoter without strong induction via P2. We propose this as one molecular basis of the tissue-1/P1 phenotype in the TP53-mutant cases (two of the 18 P1-dominant patients were TP53 wild-type and one was undetermined; p53 was inactive in both specimens of HCT59; Additional file 25: Supplementary Table S23; Section 3.2.1 of the main text).

In contrast, no Pathogenic mutation was detected in the patients with tissue 2 (P2-dominant; HCT27, HCT33, and HCT64), and TP53 was wild-type (no pathogenic variant, although p53 function was not assayed; at the patient level, by majority isoform; at the specimen level the P2 specimens HCT41-1T and HCT38-3LM carried TP53 mutations yet were p53-active, whereas HCT67-4LMR carried none; Supplementary Table S11). TP53 mutation is uncommon in MSI-type colorectal cancer, which arises through mismatch-repair deficiency with a hypermutated genome [104], and TP53 mutations are not enriched in the MSI-rich CMS1 subtype [2]; P2 dominance was, however, not confined to MSI tumors, because HCT27, one of the three P2 patients, was microsatellite-stable with retained mismatch-repair protein expression according to the biobank's records (Section 4.6), and P2_index was not associated with MSI status in TCGA (Section 3.2.2). Wild-type p53 is inferred to be persistently activated in response to environmental stresses in tissue-2 organoids—chronic inflammation (high CXCL14 expression, with TNF and IL1B predicted by IPA as activated upstream regulators; not measured), DNA damage (γH2AX nuclear foci predicted by IPA; not measured), and hypoxia (high CA9 expression, with activation of the HIF1α pathway in IPA)—which would induce the P2 promoter. We propose this as one molecular basis of the tissue-2/P2 phenotype.

These findings show that the MDM2 P1/P2 isoform difference is supported at the transcript level (Splicing Index), is placed by IPA Networks analysis in a protein-interaction module with MDM2 regulators (a knowledge-based inference, not a measurement), and co-varies with organoid morphology (VGG16 image classification), which, however, follows TP53 status. TP53 mutation status is closely aligned with this model: all three P2-dominant patients were TP53 wild-type at the patient level (with the specimen-level exceptions HCT41-1T and HCT38-3LM noted above), TP53 mutations were found in 15 of the 17 P1-dominant patients with determined status (the per-specimen three-way correspondence is given in Supplementary Table S23), and in adjusted analyses morphology followed TP53 status rather than promoter usage (Sections 3.1.4 and 3.1.5 of the main text). More than three decades after the p53–MDM2 autoregulatory loop was established in the early 1990s [3,5], this study provides a framework for examining its clinical relevance in an actual disease model, colorectal cancer patient-derived organoids. In HCT67, the primary tumor (HCT67-1T) and the liver metastasis (HCT67-3LM), which carried the pathogenic mutation R175H, showed P1 dominance and Type1-dominant morphology (EMT scores −0.62 and −0.14), whereas the recurrent liver metastasis (HCT67-4LMR) switched to P2 dominance and non-Type1 (cystic–mucinous) morphology (Type1 fraction 0.12), and an intrapatient P1→P2 switch in which the EMT score rose to +4.50 was observed (Figure 9D). R175H, detected in the primary tumor and the liver metastasis (and in a later recurrence outside the analyzed set), was not detected in HCT67-4LMR, which carried only the benign polymorphism P72R. The switch to P2 therefore occurred in a lesion without a detectable TP53 mutation, which is consistent with p53-dependent P2 induction; because the P2 promoter can also be activated in a p53-independent manner through multiple transcription factor response elements [9], P2 induction by EGR1, which showed directionally consistent but sub-threshold activity in our IPA Upstream analysis, in response to signals of the metastatic microenvironment may also have contributed, although this is speculative and was not tested. This is consistent with the integrated model of this study (Supplementary Report Section 4.2.2), in which P1/P2 choice is not fixed but is a context-dependent output; P2 activation in the P2-dominant specimens accompanied active p53 (including the two P2-dominant specimens with TP53 mutations; Section 3.2.1 of the main text), and a non-p53-dependent route via EGR1 or other factors remains a speculative possibility that was not tested.

Speculatively, this observed P1→P2 transition from the primary tumor and liver metastasis (P1-dominant) to the recurrent liver metastasis (P2-dominant) might also coincide with a transition in physical properties, in which the cells move from a relatively stiff primary site to the softer hepatic microenvironment [106]; whether the mechanical environment of the metastatic site influences p53 activity or P2 usage was not examined.

##### 4.2.2 A unified model of the four observational layers (Figure 9)

These four observational layers—TP53 mutation status, MDM2 promoter choice (P1/P2), the downstream programs, and organoid morphology—tend to co-vary along the P1–P2 contrast (Figure 9A), and the data place TP53 status at their center, but they separate into two readouts rather than a single axis: the MDM2 axis reflects p53 activity, whereas organoid morphology follows TP53 status beyond expression-inferred 20q gain and aneuploidy (Supplementary Report Section 4.2.3). At the [Tissue 1/P1] pole, mutation or deletion of TP53, or p53 inactivity in TP53 wild-type specimens (CIN/CMS2 type; TP53 mutation in 15 of the 17 P1-dominant patients with determined status, Supplementary Report Section 4.2.1) is associated with basal P1 transcription of MDM2 without p53-dependent P2 activation, and the MYC–FOXM1–E2F proliferative hub is maximized, while the epithelial character is retained and a compact glandular morphology is adopted. At the [Tissue 2/P2] pole, transcriptionally active wild-type p53 (possibly sustained by chronic inflammation and oxidative stress, which were inferred rather than measured; a signature-level resemblance to the MSI-like/serrated program rather than microsatellite instability, Section 4.6; a composite subtype centered on CMS3 at the gene level and in the P2 core score, with CMS1 features in its immune module and CMS4-like features only at the pathway level) is associated with induction of the p53-inducible P2 promoter, and partial EMT mediated by TGFβ/SMAD3 (a hybrid E/M state that does not reach complete mesenchymal conversion, in which ZEB1/2 and SNAI2 are not fully repressed but coexist at low levels while the miR-200 family is elevated), and metaplasia toward the gastric type and the small-intestinal Paneth cell type proceed in parallel (higher SNORD116 expression was seen in the P2-dominant comparisons but was not associated with P2, nor consistently with TP53 status, as continuous scores; Section 3.1.4 of the main text), (the EMT score itself does not separate the two poles: the P2-versus-P1 difference was not significant [P = 0.079], and although the highest scores were in the metastatic specimen HCT38-3LM [4.94] and the P2-dominant recurrent liver metastasis HCT67-4LMR [+4.50], sampling site is balanced between the two isoform groups and the difference was absent within primary specimens [P = 0.89] and rested on those two specimens within metastases [P = 0.20 after their removal]; the score follows the sampling site instead [P = 0.014] and is treated as a separate axis of progression; Section 3.1.2) and a mucinous, cystic and dispersed morphology (non-Type1; the high viscoelasticity of the mucus gel is inferred from gel-forming mucin expression) is adopted. In our working model, the bias in MDM2 isoforms is a consequence and readout of p53 activity; whether it feeds back on the proliferative, differentiation, and physical characteristics was not tested (beginning of Section 4.2). However, morphology is only a partial correlate of the transcript-level MDM2 P1/P2 state: it follows TP53 status beyond 20q and aneuploidy, corresponds at the level of transcript clusters to 20q, correlates only weakly with the MDM2 axis (ρ = 0.33), and overlaps with P1/P2 because TP53 status and promoter usage largely coincide (Supplementary Report Section 4.2.3; on the TP53 wild-type, largely P2-dominant side it appears as a non-Type1 [cystic–mucinous] spectrum; Section 4.1).

However, this axis is not a deterministic dichotomy but a continuous spectrum (the EMT score gradient in Figure 9B), and exceptions in which morphology and isoform diverge also exist. The EMT score is the value obtained by subtracting the mean z-score of epithelial markers (CDH1, EPCAM, etc.) from the mean z-score of mesenchymal markers (VIM, FN1, CDH2, SPP1, etc.); the more negative the value, the stronger the correspondence with epithelial characteristics (high E-cadherin expression, maintenance of LGR5-positive intestinal stemness, compact glandular morphology), and the more positive the value, the stronger the correspondence with mesenchymal transition (high VIM, FN1, and CDH2 expression), activation of invasion markers (MMP7, MSLN, PLAUR), and a dispersed mucinous morphology (Supplementary Table S1). For example, in HCT38-3LM, independently of the central markers of this study (gastric metaplasia and p53 targets), a cancer cell-intrinsic EMT program (a cadherin switch to CDH2/CDH6, elevation of VIM, FN1, and SPP1, etc.) was comprehensively activated specifically in this specimen (Section 3.1.2 and Supplementary Table S13), indicating that the EMT score axis may capture an additional dimension of plasticity that is partly independent of the morphological axis. In morphology, HCT38-3LM (P2-dominant but TP53-mutant and Type1-leaning) is one of the three deviating specimens that followed TP53 status rather than promoter usage (Section 3.1.3).

A detailed biological interpretation of the two tissue types under this unified model is provided in Supplementary Note Section 5, and only the key points are stated here. In tissue-1 cases that have lost p53 function through TP53 mutation (15 of the 17 P1-dominant patients with determined status), induction of P2 would not occur and MDM2 would be stably expressed at basal levels from the constitutive P1 promoter; this can be understood as a state that acts cooperatively with the autonomous proliferation program driven by the MYC/FOXM1/E2F axis (however, the data of this study only show co-variation between MDM2 expression and proliferation-related gene sets, and do not show functional causality; Supplementary Note Section 5.1). In contrast, tissue 2 (P2-dominant) is interpreted as a dynamic equilibrium in which the MDM2 P2 promoter is induced in a p53-dependent manner in response to sustained activation of wild-type TP53 (the top-ranked factor in the Upstream analysis), which would suppress p53-dependent apoptosis; the seemingly contradictory feature that p53 target genes, MDM2 itself, and in addition AKT1 and EGF signaling are simultaneously located at hubs of the Graphical Summary is interpreted as antagonism between p53-induced apoptotic signaling and MDM2/AKT/EGF-dependent survival signaling (an inference from bulk expression and IPA prediction; co-occurrence within the same cells was not tested) (Supplementary Note Section 5.2). As for the positioning as a molecular subtype, whereas the P1-dominant group corresponds to CMS2, the P2-dominant group is a composite subtype in which the lineage and differentiation traits (gastric metaplasia, secretion, metabolism) correspond to CMS3 and the immune-evasion traits correspond to CMS1; this dual assignment was consistent across two readouts, the score level and the gene level (Sections 3.3.2 and 3.3.3, Supplementary Tables S34 and S28; details, including verification that this is not circularity arising from the signature definitions used in this study, are given in Supplementary Note Section 5.2).

##### 4.2.3 P2 reflects p53 activity, and morphology follows TP53 status

The results of Section 3.2 of the main text make the meaning of P2, at the center of the working model, concrete. Among the 56 discovery specimens with determined TP53 status, wild-type P2-dominant specimens were p53-active compared with mutants (AUC 0.99), none having a score at or below the 90th percentile of mutants, whereas two of the three wild-type P1-dominant specimens, both from HCT59, did; the only mutant specimens with clearly positive scores were the two P2-dominant ones (HCT41-1T and HCT38-3LM). P2 was induced more strongly than P1 by nutlin-mediated p53 activation and was not induced in the absence of p53. P2 dominance therefore marks a state in which p53 is actually active, not the TP53 wild-type genotype as such. In tumor tissue, P2 tracked p53 activity in non-hypermutated tumors confirmed as wild type, though not beyond total MDM2 (only weakly, and depending on expression quantification, across all wild-type tumors), and in cell lines the P2 index tracked MDM2 dependency beyond both the p53 activity score and total MDM2, and nutlin-3a sensitivity beyond the p53 activity score but only weakly beyond total MDM2 (Section 3.2.4 of the main text); in an independent organoid biobank (GSE294511), idasanutlin sensitivity followed the P2 index but could not be separated from total MDM2 (Section 3.2.5 of the main text). To the debate over whether p53-target signatures predict MDM2 inhibitor sensitivity [17] or merely reflect misclassified genotypes [18], this study adds that lines without p53 activity were less sensitive even among lines confirmed as wild type by DepMap mutation data, and that P2 largely reflects p53 activity while carrying some information beyond the p53 activity score in GDSC and CRISPR (this independence was not seen in PRISM). An advantage of P2 is that it can be read as a single ratio at the locus of the drug target rather than as a combination of many target genes. Morphology, in contrast, correlated only weakly with P2 (ρ = 0.33); at the level of transcript clusters it corresponded to chromosomal instability centered on 20q (inferred from arm-level expression), whereas the MDM2 axis corresponded to p53 targets, and when TP53 status was compared with promoter usage, morphology followed TP53 status beyond 20q and aneuploidy, and P1/P2 usage did not contribute beyond TP53 status (Section 3.1.4 of the main text). The P2 signature also separated into two parts: its p53-target genes (MDM2 excluded) followed P2 usage beyond TP53 status in both organoid cohorts, whereas its gastric-metaplastic genes, like morphology, followed TP53 status in the independent biobank and in tumors and not P2 (Section 3.1.4 of the main text). The tissue-1/tissue-2 contrast (Figure 9A) can thus be reread as the contrast between TP53-mutant tumors with chromosomal-instability features (P1 dominance and p53 inactivity with expression-inferred 20q and 8q gains) and TP53 wild-type tumors (P2 dominance and p53 activity). The TP53 wild-type side does not mean microsatellite-unstable: P2_index did not differ among the CIN, GS and MSI subtypes and was not associated with MSI status (Section 3.2.2 and Section 3.1.4 of the main text). That the organoid morphology genes identified mucinous carcinomas in patient tumors indicates that organoid morphology retains properties aligned with pathological histology.

### R-19. Full-length version of the MDM2-bidirectionality sentence of main text Section 4.2 (Bidirectionality of MDM2; subsection 4.2.4 in the full-length numbering)

#### 4.2.4 Bidirectionality of MDM2

The bidirectional control between MDM2 and the mitotic checkpoint indicated by the Rank 2 cascade identified in the IPA Regulator Effects analysis (Section 3.3.1) (mitotic regulators such as ANLN and the AURK family → MDM2, TP53, CDKN1A, BAX, etc.) suggests, at the level of the IPA causal inference network, a bidirectional relationship in which MDM2 is an upstream regulator of p53 and at the same time a downstream target of mitotic checkpoint factors. However, because of computational constraints this module is based on the integration of only the first 8 of the 33 analyses (Sections 2 and 3.3.1), and the criteria for sign consistency and effect size that this study used for the other modules were not applied. This item is therefore positioned not as a confirmatory claim but as the presentation of a hypothesis. It is suggested that, when normal mitosis has been completed, these upstream factors are suppressed and MDM2 maintains p53 in a low-activity state so that proliferation continues, whereas when mitotic abnormalities occur, RASSF6/AURK/ANLN are activated and destabilize MDM2; the system may thus function as a quality-control mechanism in which stabilization of p53 leads to elimination of the cell through CDKN1A/BAX. Details of the 11 recurrent upstream regulators that appear repeatedly in multiple entries (including the pathway of Cullin-RING-type E3 ligase activation by NEDD8) are given in Supplementary Note Section 4.1. The NEDDylation pathway of NEDD8 is essential for activation of Cullin-RING-type E3 ligase complexes; MDM2 has also been reported to act as a NEDD8 E3 ligase for p53, and neddylation of p53 inhibits its transcriptional activity without markedly affecting its stability [155].

The IPA Networks analysis further supports this bidirectionality model. The fact that MDM2 co-occurs in the same protein-protein interaction module with the factors responsible for its degradation, phosphorylation modification, and nuclear export (Section 3.3.1) indicates that the production, modification, transport, and degradation of MDM2 are regulated in an integrated manner. This finding is consistent with a functional model in which MDM2 acts as a sensor molecule that detects the completion status of cell division and modulates p53 activity. However, this too is an interpretation based on co-occurrence relationships in a protein-protein interaction network, and is a working hypothesis that requires functional validation.

### R-20. Full-length version of main text Section 4.3 (Lineage plasticity in the TP53 wild-type, P2-dominant specimens)

#### 4.3 Lineage plasticity in the TP53 wild-type, P2-dominant specimens

An exploratory observation of this study is that, in the TP53 wild-type, P2-dominant specimens (tissue 2), attenuated intestinal identity and acquired gastric metaplasia co-occur, within the same subtype, with a partial EMT program. Whether these are causally linked cannot be tested with the design of this study; what follows is presented as a working hypothesis that may explain this co-occurrence, and is not an established mechanism.

Quintanal-Villalonga et al. argued [115] that lineage plasticity is a common pathway to therapeutic resistance. In the injured mouse colon, wild-type p53 terminates the damage-induced regenerative, fetal-like state, and p53-deficient crypts remain locked in it [116]; whether the gastric-type program of the p53-active, P2-dominant specimens relates to such injury-associated states is not established. The loss of cell identity observed in tissue 2 could lead to alteration of therapeutic targets for cytotoxic and molecularly targeted agents. The order of progression of lineage conversion may be organized into three stages: (1) p53 activation by inflammatory cytokines and ROS → induction of MDM2 P2; (2) TGFB1/SMAD3-driven EMT during the period of survival reprieve afforded by MDM2 (Thiery et al. 2009 [110]; Kalluri and Weinberg 2009 [111]); and (3) activation of gastric metaplasia, which follows TP53 status in the independent biobank and in tumors (Section 3.1.4 of the main text). Here we present it as a co-occurrence model that makes no claim about temporal order. Likewise, for the second stage, the EMT score did not differ significantly between P2 and P1 specimens in this cohort (P = 0.079), and confounding by sampling site (metastatic specimens) is possible (Section 3.1.2). Following the classification of Kalluri and Weinberg (2009) [111], and consistent with the EMT research guidelines of Yang et al. (2020) [112], the EMT of the second stage corresponds to “Type 3 EMT” (EMT involved in cancer invasion and metastasis), and is positioned as the stage at which cancer cells acquire migratory and invasive capacity together with stem cell-like properties (Mani et al. 2008 [109]). As Nieto et al. (2016) [105] show, this EMT proceeds not as a complete mesenchymal conversion but as a partial EMT, and the intermediate state between the epithelial and mesenchymal types is considered to make metaplasia into multiple cell lineages—gastric type and Paneth cell type—simultaneously possible. Induction of the MDM2 P2 promoter may contribute to this process, in that apoptosis suppression by MDM2 P2 induction may support the cell survival required for lineage conversion (this study did not perform perturbation experiments, and this interpretation requires functional validation; Section 4.6).

The derepression arm of this lineage conversion is supported, in the expression data of the present cohort, by a general difference in nuclear chromatin repression machinery. Specifically, the B-type lamins (LMNB1, LMNB2), the lamin B receptor (LBR), LAP2 (TMPO), and BANF1, which are involved in tethering lamina-associated domains (LADs) to the nuclear envelope, the H3K9 methylation writers (EHMT2/G9a, SUV39H1, SETDB1) and HP1β (CBX1), and Polycomb repressive complex 2 (EZH2, EED, SUZ12) were all coordinately highly expressed on the tissue-1 side and relatively decreased on the tissue-2 side (Additional file 58: Supplementary Table S29). In light of the framework in which nuclear lamina tension and heterochromatin silence lineage-determining gene loci as LADs, and in which their decrease permits derepression of loci and lineage plasticity [106–108], the activation of gastric metaplasia genes observed in tissue 2 is consistent with a general decrease in this repressive machinery. However, because these factors (LMNB1, EZH2, DNMT1, ATAD2, etc.) are coupled to proliferation, an aspect reflecting the high proliferative capacity of tissue 1 cannot be excluded, and causal separation of mechanical factors from proliferative state is difficult with the present data. We therefore restrict this finding to the driver-independent interpretation that “in tissue 2 the nuclear chromatin repression machinery is generally decreased, providing a permissive chromatin basis for the observed derepression of lineage loci” (an exploratory finding; direct validation by chromatin immunoprecipitation or Hi-C is required).

Note that this “loss of the colorectal type” is clearly captured as a gradient of the master transcription factor HNF4A (mean approximately +2.0-fold on the tissue-1 side, sign consistency 94%) and of LGR5. In contrast, the mean expression difference of CDX2 between the two groups is very small and almost flat (sign consistency 51.5%), and although CDX1 shows a mild direction of being higher on the tissue-2 side (mean linear FC −1.56, sign consistency 85%), its effect size is less than twofold and does not reach the DEG threshold. Therefore, the attenuation of intestinal identity is understood not as a complete loss of bulk expression of either CDX1 or CDX2, but mainly as an attenuation of the colorectal-type program captured by the HNF4A gradient, together with permissive changes at the chromatin level.

Here, the MDM2 isoform axis (P1/P2 choice) and the EMT axis may be linked through p53 as a shared upstream node (proposal of a working model). First, p53 activity (lost through TP53 mutation or through p53 inactivity in wild-type cases; Section 3.2.1 of the main text) may lie upstream of both axes. Wild-type p53 induces the P2 promoter through p53 response elements (the P2_index in this study is high in TP53 wild-type cases, Section 3.2.2). Wild-type p53 is also known to restrain EMT by inducing microRNAs of the miR-200 family (and miR-192) that repress the EMT-inducing transcription factors ZEB1/2 [113,114]; on this axis alone, loss of p53 would be expected to favor EMT, which is opposite to the observed tendency (EMT-related features somewhat more frequent on the TP53 wild-type P2 side, although the P2-versus-P1 difference in EMT score was not significant [P = 0.079]; sampling site is balanced between the two isoform groups, the difference was absent within primary specimens [P = 0.89] and rested on two specimens within metastases [P = 0.20 after their removal], and the score follows the sampling site instead, so it is treated as a separate axis of progression; Section 3.1.2). We therefore do not attribute the bifurcation between “P1 choice without EMT features” and “P2 choice with partial EMT” to p53-mediated EMT suppression; rather, we propose that p53 activity acts as the branch point through the P2-induction arm described next. The EMT transcription factor SNAI1 was higher on the TP53-mutant tissue-1 side (mean FC +1.88, 82% consistent; Supplementary Table S15), but this is not explained by the p53–miR-200 axis, because p53 inhibition did not induce SNAI1 or SNAI2 in the study that established this axis [113]. Second, in cells in which wild-type p53 has been activated, MDM2 P2 induction grants a reprieve from apoptosis and maintains cell survival, while that same p53 activity does not allow EMT to proceed to complete mesenchymal conversion, keeping it in a partial EMT state by partially repressing ZEB1/2 through the miR-200 family [113,114] (consistent with the observation in this study that, even under elevated miR-200 in the P2-dominant group, ZEB1/2 and SNAI2/SLUG were consistently in the higher direction while their effect sizes were modest; Additional file 21: Supplementary Table S15). That is, autonomous proliferation without EMT features in the P1-dominant group (TP53-mutant, or wild-type with inactive p53), and apoptotic reprieve, a tendency toward partial EMT, and multilineage metaplasia in the P2-dominant group (wild-type with active p53), may be linked with p53 activity as the branch point. The observed morphology (compact glandular versus mucinous cystic), by contrast, is not an output of this model but follows TP53 status beyond expression-inferred 20q gain and aneuploidy, overlapping with P1/P2 because TP53 status and promoter usage largely coincide (Supplementary Report Section 4.2.3). In this working hypothesis, MDM2 P2 induction might function not merely as a temporal prerequisite for EMT but as a node that, by directing p53 activity toward survival, renders the progression of partial EMT compatible with cell survival. However, this model is a hypothesis derived from the correlative data of this study (TP53 mutation status, P2_index, p53 target modules, miR-200/ZEB1/2, and the TGFβ/SMAD3 pathway), and each causal relationship within the model—in particular reverse feedback from EMT transcription factors or the TGFβ system to MDM2 promoter choice, and the possibility that a difference in translational efficiency between the P1 and P2 isoforms specifies the EMT threshold through the strength of p53 repression—requires functional validation including promoter-specific manipulation and TP53 perturbation (Section 4.6).

### R-21. Full-length version of main text Section 4.4 (Comparison with previous studies)

#### 4.4 Comparison with previous studies

We compare the findings of this study with existing CRC organoid studies and with morphology × AI studies. van de Wetering et al. (2015) [117] described the diverse patient-specific organoid morphologies (from cystic to densely packed), but did not link morphology to molecular subtypes or to therapeutic stratification. Fujii et al. (2016) [118] showed, in a large CRC organoid library, the stepwise loss of niche factor requirements accompanying carcinogenic progression, and Betge et al. (2022) [119] described drug-induced phenotypic landscapes on a large scale; they thus related CRC organoid phenotypes to niche-factor requirements and to drug responses, respectively. Zhao et al. (2021) [10] linked EMT status and drug response by morphology × deep learning in breast cancer organoids, and Lukonin et al. (2020) [11] quantified the phenotypic landscape of normal intestinal organoid regeneration by high-content image-feature profiling. In colorectal cancer as well, there is a report that identified cystic and solid morphological subtypes by image-based profiling of bright-field images, related them to organoid viability and apoptosis by deep learning, and proposed the cystic subtype, a relapse phenotype with intestinal stem cell signatures, as a candidate biomarker for diagnosis and prognosis [120]. This study extends to colorectal cancer these frameworks that link morphology and molecular state by image-based analysis, and is novel in linking morphology to TP53 status and to a transcript-level readout of p53 activity (MDM2 P1/P2), and to therapeutic stratification. A detailed comparison, including the mutual complementarity of an independent analysis using a cohort shared with Okamoto et al. (2022) [12], is given in Supplementary Results 6 of the Supplementary Information and in a comparison table (Additional file 59: Supplementary Table S33). In addition, Okamoto et al. (2021) [24], which reported the establishment of the same biobank, described a cell-composition axis — a reduction of OLFM4-associated stem-like clusters in metastatic lesions — on the basis of patient-matched primary–metastasis comparisons; morphological typing (Okamoto 2022 [12]), cellular composition (Okamoto 2021 [24]) and the alignment of morphology with TP53 status together with a promoter-level p53 readout (MDM2 P1/P2; this study) thus constitute three independent, mutually complementary analytical axes built on the same material. In addition, the original morphological-typing study [12] reported that PDO morphology is not correlated with the clinicopathological features of the original tumors and showed no apparent correlation with the APC/TP53/KRAS mutation profile. In our specimens, by contrast, TP53-mutant organoids had a higher Type1 fraction, beyond 20q and aneuploidy, and in an independent biobank the morphology gene score followed TP53 status (Section 3.1.3 and Sections 3.1.4 and 3.1.5 of the main text); the difference may reflect our two-class summary (Type1 versus non-Type1) of their morphological types and the form of the comparison — a continuous per-specimen fraction against TP53 status, rather than categorical types against a mutation profile. In mouse Apc-mutant intestinal tumor organoids, which were round and cystic when Trp53 was wild-type or deleted, homozygous mutant p53 (R270H) induced complex glandular (tubular) structures associated with invasiveness, and in nine human colorectal cancers, tumors with TP53 mutations around codon 273 (six) had higher gland-branching scores than TP53 wild-type tumors (three) [121]; our human organoid data agree in direction (compact glandular Type1 morphology in TP53-mutant organoids), although they cannot distinguish the effects of mutant p53 from those of p53 loss. Our associations were observed at the specimen level, the unit at which morphology, isoform and TP53 status varied within patients (Sections 3.1.2 and 3.1.3), and the TP53–morphology association was replicated across patients in the larger independent biobank (Section 3.1.5 of the main text and Section 4.1). Furthermore, the novelty of this study is differentiated from previous reports along three axes. First, although the basic biology of the dual P1/P2 promoters of MDM2 (P1 is constitutive, whereas P2 is p53-responsive and is regulated by response elements for multiple transcription factors) is established [6,9,59], we found no report that used MDM2 promoter usage as a locus-level readout of p53 activity in CRC. In fact, in oral cancer it has been reported that induction of the P2 transcript correlates with stabilization of wild-type p53 [122], which is consistent, by extrapolation, with the correspondence of tissue 2 (P2-dominant, TP53 wild-type) in this study. It has also been reported that, in CRC, MDM2 amplification does not correlate with SNP309 or with TP53 mutation status [123], which supports the independence of the “promoter choice” axis of this study from the known amplification and SNP axes. Second, recent findings on lineage plasticity in CRC center on a fetal progenitor state and suppression of non-intestinal lineages by PROX1 [124], and on maintenance of colonic epithelial identity by HNF4A and ATRX [125]. Note that the framework itself, which positions gastric metaplasia as the starting point of the serrated tumorigenesis pathway, has already been proposed [98], and the primary study on which it is based also reached the same conclusion from single-cell analysis of human colorectal polyps [99]. The gastric metaplasia signature that this study showed in tissue 2 is not a novel observation. The novelty of this study lies not in the existence of metaplasia, but in showing that the metaplastic program follows TP53 status (in the independent biobank and in tumors) rather than MDM2 promoter usage, extending to organoids the lower frequency of p53-pathway alterations reported in mucinous colorectal carcinomas [14]. Third, whereas existing classifications such as CMS [2], CRIS [126], and iCMS [15] are all based on multigene signatures, this study is methodologically orthogonal in that it unifies a CMS2-like type (tissue 1) and a type centered on CMS3 with CMS1-like immune features (tissue 2; CMS4-like features only at the pathway level) through TP53 status and a single locus-level readout of p53 activity (MDM2 P1/P2). These claims of novelty rest on the fact that the molecular axis addressed and the analytical approach used are independent of existing classifications.

### R-22. Full-length version of main text Section 4.5 (Clinical Implications; subsections 4.5.1–4.5.3 below follow the full-length numbering and are condensed into main text Section 4.5)

#### 4.5 Clinical Implications

Before discussing therapeutic targets, we summarize the implications of the physical material properties and of drug accessibility (details in Supplementary Note Section 5.3). The contrast in material properties between the “dense solid” of tissue 1 (solid stress arising from high cell density) and the “highly viscoelastic gel” of tissue 2 (a mucus barrier of MUC5B, MUC5AC, and MUC17 together with high interstitial fluid pressure) (Additional file 60: Supplementary Table S45) — physical states inferred from gene expression, not measured in this study — may limit drug accessibility in both, although through different modes [127,128]. In tissue 2, vascular normalization by anti-VEGF therapy [129] becomes an important perspective for combination strategies that improve the intratumoral penetration of subsequent treatments.

For the 39-gene panel identified by IPA Biomarker Detection analysis (Section 3.3.3; Supplementary Figure S5), the partition obtained by hierarchical clustering agreed with MDM2 promoter usage (P1 versus P2) in 62 of 63 specimens (methods and breakdown in Section 3.3.3; because the panel was selected from the same P1-versus-P2 comparisons, this is a within-sample, non-independent agreement). The wild-type TP53 target genes contained in this panel (nine genes including CDKN1A, FAS, and ZMAT3; Section 3.3.3), the gastric metaplasia markers (CTSE, REN), and the invasion markers (MSLN, MMP7) summarize the p53-active/metaplastic contrast of these specimens rather than providing an independent classifier; an assay derived from them (an RT-qPCR multiplex assay of a selected panel of 5–10 genes, or a targeted NGS panel) would require independent validation in biopsy tissue. Because organoid morphology followed TP53 status rather than promoter usage (Section 3.1.3 and Sections 3.1.4 and 3.1.5 of the main text), morphometric analysis with the deep-learning classifier (98.5% test accuracy) could at most serve as a pre-screen for TP53 status rather than for promoter usage in settings where molecular assays are unavailable (the TP53 wild-type side is best assessed not as Type5 alone but as non-Type1 [cystic–mucinous]; Sections 3.1.2 and 4.1). Conversely, quantifying the MDM2 isoform ratio (the P1/P2 ratio) by RNA-seq or RT-qPCR could serve as a candidate readout of p53 activity obtainable from a tumor biopsy (not yet validated in biopsy tissue); because the P1 score was not associated with prognosis independently of stage (Section 3.3.2), a prognostic use is not proposed. Because P2 tumors retain wild-type p53, release of p53 function by MDM2 inhibitors could induce apoptosis (the support from the Upstream analysis and the current status of clinical development are given in Supplementary Report Section 4.5.2). Two exploratory therapeutic hypotheses are consistent with the drug-response data: MDM2 inhibitors for the P2-dominant group (Section 3.2.4 and Supplementary Report Section 4.5.2) and ATR/CHK1/WEE1 checkpoint inhibitors for the P1-dominant group (Section 3.2.4); the other candidates below, suggested only by IPA upstream z-scores or expression patterns, remain hypotheses. Furthermore, the detection of the XPO1 inhibitor eltanexor (median z = −4.59, sign consistency 97%) as an upstream regulator in the tissue-2 direction in the IPA Upstream analysis (selinexor, although also an XPO1 inhibitor, had a median z = −1.22, an effect size below the threshold, and the two are distinct compounds; Supplementary Table S7) suggests, from two IPA modules applied to the same comparisons (Networks analysis and Upstream analysis), that a therapeutic strategy of nuclear retention of MDM2 leading to p53 stabilization through XPO1 inhibition is a hypothesis worth testing. The expression of TNFRSF10B (DR5) and TNFRSF10C (DcR1) identified by IPA Biomarker Detection is consistent with engagement of the TRAIL (TNF-related apoptosis-inducing ligand) receptor pathway (pathway activity was not measured) and raises the hypothesis that P2 tumors could be sensitive to TRAIL-based agents. The efficacy of immune checkpoint inhibitors in MSI-H colorectal cancer, shown by Le DT et al. [133], is directionally in line with the NK cell signaling found in tissue 2 (CGAS-STING was below the effect-size threshold; Supplementary Note Section 3.1). In addition, as described above, seven immune checkpoint molecules (TIGIT, CTLA4, PDCD1, IDO1, LAG3, IDO2, HAVCR2) were consistently more highly expressed in tissue 2 in terms of direction (sign consistency 66.7–84.8%; the effect sizes were all below the DEG criteria, and this finding is exploratory, based on the reproducibility of direction; Section 3.3.3 and Supplementary Table S30), and these resemble the immunosuppressive tumor microenvironment of the MSI-H type (a signature-level resemblance; Section 4.6). Combination strategies with IDO inhibitors (e.g., epacadostat) or LAG-3/TIGIT inhibitors, in addition to anti-PD-1/PD-L1 antibodies, are hypothetical therapeutic options. Of note, these individual drug candidates are all computational suggestions based on IPA z-scores and expression patterns; given the limitation described in Section 3.2.4—that a drug-sensitivity score can correlate with drug sensitivity in general rather than with any specific agent—the same caution applies to interpreting their specificity for individual agents. Sections 4.5.1–4.5.3 below present specific therapeutic strategies for each tissue type.

We also applied explicit criteria to the selection of this list of immune checkpoint molecules (Supplementary Table S30 and Figure 8 panel B). First, among the genes more highly expressed in tissue 2, we took as candidates those corresponding to clinically druggable immune checkpoint or immunosuppressive molecules (the PD-1/PD-L1 axis, CTLA-4, LAG-3, TIGIT, TIM-3, the IDO pathway, and other targets of approved drugs or of agents in clinical development). Next, for these candidates we calculated the sign consistency in the tissue-2 direction across the 33 two-group comparisons and presented the seven molecules with directionally consistent higher expression (exploratory; below the DEG effect-size criterion) (TIGIT, CTLA4, PDCD1, IDO1, LAG3, IDO2, HAVCR2) (Figure 8 panel B is ordered by sign consistency; the data are in Supplementary Table S30). Of these, six molecules—IDO1, PDCD1, LAG3, CTLA4, TIGIT, and HAVCR2—were adopted for the P1/P2 signatures (Section 2) on the basis of representativeness and of the degree to which they are established as targets in previous reports, whereas IDO2 was excluded from the scoring signature because its immune-evasion function is represented by the closely related IDO1. That is, the seven molecules in Figure 8 are defined as “druggable checkpoint molecules with high sign consistency,” and the six molecules in the signature as “those among them with high representativeness and a well-established target status,” so that the list is constructed by a two-step set of criteria.

##### 4.5.1 Therapeutic strategy for tissue 1 (P1, autonomous-proliferation type)

Tissue 1 corresponds to the CIN type and the CMS2 subtype, and the following established strategies apply according to their own biomarkers (none was tested against the P1/P2 axis in patients). In standard chemotherapy, 5-FU (fluorouracil)-based agents and oxaliplatin, which target the high mitotic rate driven by FOXM1, MYC, and E2F, are expected to work as standard therapy, although neither their predicted nor their measured sensitivity was P1-specific (Section 3.2.4). In RAS/BRAF wild-type, left-sided primary tumors, anti-EGFR antibodies (cetuximab, panitumumab) may be considered. Because ERBB2 showed high sign consistency in the tissue-1 direction in the Upstream Analysis (the effect size was below the adoption threshold; Supplementary Note Section 2.2 and Supplementary Table S7), anti-HER2 therapy (trastuzumab plus tucatinib [130], or trastuzumab deruxtecan) may also be considered in cases showing HER2 amplification. The marked upregulation of CDK1/2, PLK1, and AURKA/B also suggests possible applicability of mitotic kinase inhibitors targeting these molecules (PLK1 inhibitors, Aurora inhibitors, etc.), although in cell lines the P1-associated sensitivity was clearest for ATR/CHK1/WEE1 checkpoint inhibitors rather than for PLK1 or Aurora kinase inhibitors (Section 3.2.4).

As a recurrence risk, persistence of the LGR5- and OLFM4-positive stem cell population is a concern. If LGR5-positive stem cells persist even after the glandular architecture has been destroyed by treatment, they become the source of minimal residual disease (MRD). Combination with stem cell-targeted therapy such as an LGR5-ADC (antibody–drug conjugate) may be effective in preventing recurrence.

##### 4.5.2 Therapeutic strategy for tissue 2 (P2, environment-adaptive type)

Tissue 2 resembles the MSI-like/serrated program at the signature level (MSI was not measured, and P2 dominance does not itself imply MSI-High/dMMR; Section 4.6) and corresponds to a composite type centered on CMS3 at the gene level and in the P2 core score, with CMS1 traits in its immune module and CMS4 traits only at the pathway level; the following multilayered therapeutic strategy is considered to be necessary. As for immunotherapy, when MSI-High/dMMR is present, immune checkpoint inhibitors show high efficacy [133]. In first-line treatment of metastatic cases, nivolumab plus ipilimumab prolonged progression-free survival compared with chemotherapy [134], and pembrolizumab monotherapy remains a standard of care, with its benefit over chemotherapy sustained at more than 5 years of follow-up [135]. In addition, the addition of atezolizumab in adjuvant therapy for stage III dMMR colon cancer [136] and nonoperative management of dMMR tumors with an anti-PD-1 antibody (dostarlimab) [137] have also been reported, and the treatment framework for dMMR colorectal cancer has changed substantially in recent years. The identification of STAT4 as an activated upstream regulator in the Upstream Analysis, with IFNG also in the same direction although its effect size was below the adoption threshold (the values for each regulator are given in Supplementary Table S7), and the enrichment of Natural Killer Cell Signaling in the Canonical Pathway analysis (Table 3) are consistent with an antitumor immune response (inferred from expression). However, this implication rests on the established indication of MSI-High/dMMR [133], and it does not show that the P1/P2 axis of this study itself predicts response to ICI. As shown in Section 3.3.3, although TIDE-based response prediction showed a high predicted responder rate in the P2-high group, this was not reproduced in the population for which MSI annotation was available, and the TIDE predicted response rate was in fact lower on the MSI-High side. Therefore, the statements on immunotherapy in this section are a reaffirmation of the known indication for P2 cases that carry MSI-High, and they do not propose P2 as a new ICI-eligible population.

With regard to breaching the physical barrier, normalization of the tumor vasculature by anti-VEGF therapy (e.g., bevacizumab) may lower IFP and improve the intratumoral penetration of subsequent chemotherapy and immunotherapy. Against the mucus barrier (MUC5B/MUC5AC), pretreatment with the mucolytic agent NAC (N-acetylcysteine) could function as an adjuvant. Indeed, Cantero-Recasens et al. showed that colorectal cancer cells increase mucin (MUC5AC) secretion in response to chemotherapy, forming a physical barrier that impedes cellular drug uptake, and that suppression of mucin secretion restores the sensitivity of patient-derived organoids to 5-fluorouracil plus irinotecan by 40-fold [138], supporting the view that control of the mucus gel is a rational target for improving drug accessibility in tissue-2 type tumors.

With regard to direct targeting of the MDM2 P2 axis, MDM2 inhibitors (e.g., milademetan/DS-3032b) [139] may be more effective in tissue 2. Because p53 is retained in the wild-type state in tissue 2, release of p53 function by MDM2 inhibition could induce apoptosis. In contrast, little efficacy can be expected in the TP53-mutant cases of tissue 1, although P1-dominant cases that retain wild-type TP53 mostly lacked p53 activity (Section 3.2.1 of the main text), so candidates may be narrowed to TP53 wild-type, P2-dominant cases (Supplementary Report Section 4.5.3). The fact that milademetan (DS-3032b) was listed as an upstream regulator in the IPA Upstream Analysis points in the same direction. We confirmed in the primary IPA export that this regulator met the adoption criteria (sign consistency of 75% or more and |median z| ≥ 2): sign consistency 100%, 31/31; a z value was computed in 31 of the 33 analyses; median z = −2.74 (Supplementary Table S7). Furthermore, this therapeutic hypothesis is consistent with independent pharmacogenomic data (the GDSC2 cell line panel). TP53 wild-type colorectal cancer cell lines were significantly more sensitive to MDM2 inhibitors (nutlin-3a, Serdemetan) (MSI-High lines were also more sensitive, but MSI-H is not a proxy for P2; Sections 3.2.2 and 3.2.4), and these independent large-scale cell line data are consistent with the hypothesis of this study that the P2-dominant group, which retains wild-type p53, is a candidate target population for MDM2 inhibitors (GDSC [49,50]); sensitivity is determined mainly by TP53 function, and whether the P1/P2 ratio adds information beyond TP53 status is untested (Section 4.6).

This therapeutic implication has become more concrete in light of recent clinical data. Milademetan was evaluated in a phase II trial (MANTRA-2, 2025) in TP53 wild-type, MDM2-amplified advanced solid tumors, and a best overall response of 19.4% (6/31; confirmed objective response rate, 3.2% [1/31]) and a median progression-free survival of 3.5 months were reported—antitumor activity that is limited for a single agent but nonetheless real [140]. The P2-dominant group in this study retains TP53 wild-type status while overexpressing MDM2 mRNA approximately 7-fold through P2 promoter-driven transcription (mean linear fold change 7.0-fold, in the same direction in all 33 comparisons; Table 4). The P2-dominant group shares wild-type p53 and high MDM2 expression with the MANTRA-2 target population, although it is not defined by MDM2 amplification; it is therefore a candidate, not an established, target population for MDM2 inhibitors. At the same time, the limited single-agent response rate suggests the need for patient-selection biomarkers that go beyond the single indicator of TP53 wild-type status. MDM2 P1/P2 promoter usage (P2_index) could serve as an additional patient-selection indicator that directly captures the functional driving mode of MDM2 (a P2-dependent wild-type p53 axis) in addition to TP53 mutation status. It should be noted that clinical development of this class is in a difficult phase as of the time of writing (October 2026). For milademetan, the phase III trial in dedifferentiated liposarcoma (MANTRA) did not meet its primary endpoint [141], and development was halted; brigimadlin (BI 907828) was discontinued in 2025 [142] (Brightline-1, a randomized phase II/III trial [143]); siremadlin (HDM201) development in acute myeloid leukemia was discontinued [144]; and idasanutlin (RG7388) also failed to improve overall survival in a phase III trial (MIRROS) in relapsed/refractory acute myeloid leukemia [145,146]. Navtemadlin (KRT-232, formerly AMG-232) remains in phase III in myelofibrosis (the ongoing POIESIS trial; the earlier phase III BOREAS trial did not meet its primary endpoint of spleen volume reduction) [144], and every agent of this class remains at the stage of clinical development [146]. These trials share the feature that they select patients by a single genomic indicator, either “TP53 wild-type” or “MDM2 amplification,” and an indicator such as P2_index, which captures the driving mode of MDM2 itself at the transcriptional level, would make patient selection possible along an axis that these trials did not use. Sonkin has also reported that expression signatures based on TP53 target genes do not predict response to MDM2 inhibitors in TP53 wild-type tumors and that they in effect function merely as a surrogate for TP53 mutation status [18]. P2_index differs from these in that it measures upstream promoter usage itself rather than downstream target expression; however, because P2_index is itself strongly associated with TP53 status (Section 3.2.2), whether it has response-predictive ability independent of TP53 status was not tested in this study, and prospective validation in cohorts with response data is required (Section 4.6). MDM2 degraders, which can overcome the feedback induction of MDM2 that limits MDM2 inhibitors, have shown more sustained p53 pathway activation and deep tumor regressions in TP53 wild-type Merkel cell carcinoma models [147], and would face the same question of patient selection.

Furthermore, the molecular characteristics of the P2-dominant group provide a rationale for combination therapy. First, in TP53 wild-type colorectal cancer with MAPK pathway activation, the combination of a MEK inhibitor and an MDM2 inhibitor has shown synergistic antitumor effects accompanied by p53 pathway activation in patient-derived xenograft models [148]. The P2-dominant group shows the RAF/MAP Kinase Cascade in the IPA Canonical Pathway analysis and activation of MAPK3 (ERK1) in the Upstream analysis (Section 3.3.1), and it is therefore a candidate for this combination strategy. Second, MDM2/MDMX inhibition has been reported to act synergistically with anti-PD-1 immunotherapy in wild-type p53 tumors [149]. The P2-dominant group retains wild-type p53 and showed directionally consistent higher expression of immune checkpoint molecules (IDO1, PDCD1, LAG3, CTLA4, TIGIT, HAVCR2; exploratory); MDM2-inhibitor-plus-ICI combinations are rational to test in TP53 wild-type tumors, but whether the P2-dominant group specifically benefits was not examined.

With regard to epigenetic targeting, and only as an exploratory hypothesis, 5-azacytidine and HDAC inhibitors (e.g., vorinostat) might be considered in relation to the altered DNA methylation and histone modification that may underlie epigenetic plasticity (not tested in this study; Section 4.3). In the Upstream Analysis, the DNMT inhibitor decitabine (median z = −4.54, sign consistency 100%) and the HDAC inhibitor vorinostat (−2.17, 97%) were both detected as upstream regulators in the tissue-2 direction that met the adoption criteria. 5-azacytidine was in the same direction, but its effect size was below the threshold (−1.95, 88%). Because decitabine and 5-azacytidine are distinct compounds, we describe them separately here.

##### 4.5.3 Therapeutic stratification using the MDM2 P1/P2 ratio as a biomarker

A clinical proposal of this study is a stratification algorithm that uses the P1/P2 promoter usage ratio of MDM2 (the P1/P2 ratio) as a candidate biomarker for treatment selection (hypothesis-generating). As a concrete measurement method, 5'-UTR sequence analysis of MDM2 transcripts by RNA sequencing (P1-derived transcripts contain exon 1, whereas P2-derived transcripts do not) is practical. (Note that the measurement method and the stratification algorithm proposed in this section are conceptual proposals, and they are to be validated prospectively in a dedicated translational study with functional drug-response data and a prospective design.)

P1-dominant tumors (high P1/P2 ratio; tissue-1 type): TP53 mutation is expected in most cases (15 of the 17 P1-dominant patients with determined status in the discovery cohort; the two wild-type patients, HCT59 and HCT71, are exceptions (Supplementary Table S23); because p53 was inactive in both specimens of HCT59 (Section 3.2.1 of the main text) and wild-type cell lines without p53 activity were less sensitive to nutlin-3a (Section 3.2.4 of the main text), MDM2 inhibitor candidates may be narrowed to TP53 wild-type, P2-dominant tumors — the P1/P2 ratio being a candidate axis for selection that reflects p53 activity, not only TP53 genotype, although in bulk tumors it did not add to total MDM2 expression [Section 3.2.3 of the main text]), and 5-FU-based chemotherapy, oxaliplatin, and EGFR inhibitors (in RAS wild-type cases) remain the standard options (neither the predicted nor the measured sensitivity to 5-FU or oxaliplatin was P1-specific; Section 3.2.4). Enrollment in trials of ATR/CHK1/WEE1 checkpoint inhibitors, for which the P1-associated sensitivity in cell lines was clearest (Section 3.2.4), and of CDK and PLK1 inhibitors should also be considered. When a KRAS G12C mutation is present, sotorasib (960 mg) plus panitumumab prolonged progression-free survival compared with standard care in chemotherapy-refractory disease [131], and when a BRAF V600E mutation is present, first-line encorafenib plus cetuximab plus chemotherapy improved the objective response rate compared with standard of care [132] (the latter tends to co-occur with MSI-High and the serrated pathway, and therefore also requires consideration in the tissue-2 type). P2-dominant tumors (low P1/P2 ratio; tissue-2 type): TP53 wild-type status with active p53 is expected (MSI-High is not implied: P2_index was not associated with MSI status in TCGA; Section 3.2.2); in cases in which MSI-High/dMMR is confirmed, immune checkpoint inhibitors should be considered as first-line therapy on the basis of the known indication (this study does not propose the P1/P2 ratio itself as an indicator of ICI eligibility; Section 3.3.3), and a combination of breaching the physical barrier (anti-VEGF therapy), MDM2 inhibitors, and epigenetic therapy is a hypothesis that could be tested. These therapeutic stratifications are, at present, implications based on sensitivity in a cell line panel (GDSC2) and on molecular correspondence in public cohorts, and they require prospective validation with patient-level treatment response data. In the exceptional cases in which morphology and MDM2 isoform diverged, the P1/P2 state remained definable by the exon-specific Splicing Index, and in morphology three of the four followed TP53 status rather than promoter usage (Sections 3.1.2 and 3.1.3). As future directions, the following are important: (i) standardization of methods for measuring the P1/P2 ratio by RT-qPCR or targeted NGS and validation of outcome and treatment response in prospective cohorts; (ii) direct evaluation of the responsiveness of P2-dominant (TP53 wild-type) tumors to MDM2 inhibitors (as of the time of writing, navtemadlin, KRT-232, remains in phase III; Supplementary Report Section 4.5.2); and (iii) development of a companion diagnostic that integrates TP53 status and promoter usage, in which organoid morphology, which followed TP53 status, could at most serve as a pre-screen for TP53 status rather than for promoter usage.

### R-23. Full-length version of main text Section 4.6 (Novelty and limitations)

#### 4.6 Novelty and limitations of this study

The novelty of this study can be summarized in the following four points. (1) To our knowledge, the first observation in a human CRC organoid cohort that organoid morphology follows TP53 status (previously described in mouse intestinal tumor organoids and a small human series [121]) — compact and glandular in TP53-mutant and round, cystic–mucinous in wild-type organoids — beyond 20q and aneuploidy, in the discovery cohort (56 specimens from 20 patients; P = 0.013, specimen level) and in an independent biobank (GSE294511; P = 6.1×10⁻⁴; also on unadjusted patient means), with the morphology gene program identifying mucinous carcinoma and following TP53 status beyond 20q, MSI, BRAF mutation and CIMP in patient tumors (Section 3.1.3 and Sections 3.1.4 and 3.1.5 of the main text; in tumors, mucinous histology has been linked to fewer p53-pathway alterations [14]); the correspondence of morphology with MDM2 mRNA isoforms at the specimen level, in 21 patients and 63 samples (a P1 group dominated by Type1 [compact glandular] morphology and a non-Type1 [cystic–mucinous] P2 group: median Type1 fraction 0.826 versus 0.444, P = 1.1×10⁻³; Section 3.1.2), is consistent with this alignment with TP53 status (the size of the present cohort is limited, and confirmation requires replication in a large cohort; see the latter part of Section 4.6). The external analyses in independent cohorts — the CMS-wise distribution of module-specific GSVA scores, dMMR/pMMR discrimination, and enrichment analysis of DEGs against CMS class-specific markers (Sections 3.3.2 and 3.3.3) — concern the expression signatures; the image-based morphological correspondence itself was examined externally through a morphology gene program in an independent organoid biobank (GSE294511; Section 3.1.5 of the main text), where its across-sample alignment with P2 was largely explained by TP53 status, and in TCGA tumors. (2) Direct quantification of promoter usage itself (P2_index) in TCGA, where it tracked TP53 status and a truncating < missense < wild-type gradient not seen in p53-target output (Section 3.2.2). (3) Evidence that, among TP53 wild-type samples, P2 dominance marks a state in which p53 is transcriptionally active — P2 was induced in a p53-dependent manner by nutlin, tracked p53 activity in non-hypermutated wild-type tumor tissue (though not beyond total MDM2) and cell lines, and in cell lines tracked MDM2 dependency beyond both p53 activity and total MDM2, and nutlin-3a sensitivity beyond p53 activity but only weakly beyond total MDM2 (Section 3.2 of the main text). (4) The separation of the P2 signature into a p53-target part that follows P2 usage beyond TP53 status in organoids and a gastric-metaplastic part that, like morphology, follows TP53 status (in the independent biobank and in tumors) and not P2 (Section 3.1.4 of the main text). Other contributions include the reproduction of the CMS and mismatch-repair associations of the signatures in two independent cohorts (n = 1,143 cases in total; the P1 score was not associated with prognosis independently of stage, Section 3.3.2) and an unbiased drug analysis in which nutlin-3a ranked first for the P2 score on both predicted and measured sensitivity and ATR/CHK1/WEE1 inhibitors tracked the P1 score (Section 3.2.4). The following contributions are exploratory: (i) Construction of a multilayered comparative framework integrating all eight IPA analysis modules with EnrichR GO (integrated figure = Additional file 49: Supplementary Figure S14; Section 3.3.1). (ii) Presentation of the hypothesis (an IPA-based inference not tested experimentally) that the molecular origin of the clinical-chemistry toxicity indices (only ALP meets the adoption criteria; AST, ALT, and LDH have effect sizes below the threshold and are presented alongside only as supporting evidence; Supplementary Note Sections 3.4 and 5.2) lies in necrotic cell breakdown accompanying the antagonism between p53-induced apoptotic signaling and MDM2 P2-dependent survival signaling. (iii) A working model linking the expression-inferred physical properties of tissue 1 (“dense solid”) and tissue 2 (“highly viscoelastic gel”), which were not measured, to jamming/unjamming transitions of cell collectives [150] and extending this to the therapeutic implication of drug accessibility to the tumor (morphology itself follows TP53 status rather than the MDM2 axis; Supplementary Report Section 4.2.3).

On the other hand, this study has the following limitations. It consists mainly of bioinformatic analysis, and we did not perform cell-functional experiments that switch MDM2 isoforms. The cohort size (21 patients, 63 samples) is too small for a comprehensive clinical outcome analysis, but the associations of the signatures were reproduced in the independent cohorts TCGA-COAD/READ (n = 624) and GSE39582 (n = 519) (Section 3.3.2). Note, however, that these external validations tested between-group differences in the signature scores; discriminative performance (AUC, sensitivity, and specificity) was not evaluated. Compared with a biobank that has assembled patient-derived tumor organoids on a large scale (256 lines, of which 162 lines underwent genome-wide CRISPR), the scale of this study is small [13]. However, that biobank did not address promoter choice, isoforms, or morphological classification, and the MDM2 P1/P2 axis presented in this study is complementary not in scale but in the axis that is measured. Note that, although the P1 signature score showed a trend toward association with favorable overall survival in multivariate Cox analysis (with mutual adjustment for the P2 score), it did not reach the significance level (HR = 0.852 per 1 SD, P = 0.080), and it did not show independence in a multivariate model that additionally included age, stage, and MSI (Section 3.3.2). This loss of independence is interpreted as arising from the fact that the prognostic signal is carried mainly by the proliferation module (HR = 0.82 per 1 SD, P = 0.029), together with the structural correlation whereby clinical stage itself partly reflects the proliferative and invasive state of the tumor. That is, the P1/P2 signatures are associated with the biological properties of the tumor (CIN-type high proliferation versus inflammation and metaplasia with a signature-level MSI-like resemblance; the metaplastic genes follow TP53 status rather than P2, Section 3.1.4 of the main text), but its association with prognosis appears as co-variation with stage rather than as an independent prognostic factor. Consistent with this interpretation, P2_index, which directly quantifies P1/P2 usage as a continuous variable, also showed no association with overall survival (KM P = 0.82; Section 3.2.2). In the future, validation in prospective cohorts and exploration of subgroup effects by stage-stratified analysis are considered to be useful.

The HTA2.0 microarray is limited to exon-level resolution. MSI/MMR status was not determined in this study; the biobank's records available to us document it for one patient only — HCT27, one of the three P2 patients, was microsatellite-stable at six markers (BAT25, BAT26, BAT40, D2S123, D5S346, D17S250) with retained MLH1, MSH2, MSH6 and PMS2 expression — so the resemblance of the P2 signature to MSI-like tumors (inferred from the dMMR association of the P2 signature in GSE39582 and its immune profile) is a signature-level resemblance rather than MSI status and requires direct confirmation. Prior chemotherapy (eight specimens, one of them P2-dominant: HCT38-3LM) and passage number (1 to 22) were not adjusted for (Section 2), and the public arrays of Sections 3.1.2 and 3.2.1 are other specimens from patients of the discovery cohort; the independent biobank (GSE294511) of Sections 3.1.5 and 3.2.5 of the main text comprised liver metastases only and recorded morphology on H&E sections rather than with our classifier. We found no public bright-field organoid images with matched expression data, so the image classifier itself was not tested on external images; only the morphology gene score derived from it was.

In addition, the validation of MDM2 inhibitor sensitivity using the GDSC cell line panel does not directly identify the P2-dominant group; it is a cell line-level analysis that uses TP53 status as a surrogate index (stratification by MSI status is only a background observation, and MSI-H is not a proxy for P2; Section 3.2.2), and direct validation of the association between P2_index (promoter usage) in clinical specimens and drug response remains a future task. For the four cases whose morphology deviates from the typical pattern of their group (HCT31-4LMR, HCT38-3LM, HCT41-1T, HCT71-3LM; Section 3.1.2), additional factors such as epigenetic regulation of the MDM2 promoter, copy number variation, and post-translational modification need to be examined. In addition, the P1/P2 axis of this study has only been mapped onto the existing CMS classification, and we have not mapped it onto the IMF classification [15], which combines the intrinsic epithelial subtypes based on single-cell analysis (iCMS2/iCMS3) with the microenvironment and fibrosis. In particular, the metaplastic and inflammatory traits shown by the P2-dominant group may overlap with the description of iCMS3, and validating where this axis is positioned within the iCMS/IMF framework remains a future task. Furthermore, it has already been reported that a TP53 target gene signature alone cannot predict MDM2 inhibitor response in TP53 wild-type tumors [18], and whether P2_index has predictive ability independent of TP53 mutation status is likewise untested (Supplementary Report Section 4.5.2): the TP53-adjusted association of the P2 signature with MDM2 inhibitor sensitivity was significant but small in GDSC2 cell lines (Section 3.2.4) and in the same direction but not significant after correction in the 65 organoids of Herranz-Ors et al. (Section 3.2.4), and neither analysis used P2_index. Finally, the morphology–isoform correspondence was significant at the specimen level but, with the P2-dominant specimens deriving from six patients, not after adjustment for patient (linear mixed model P = 0.069; Section 3.1.2), so it requires confirmation in more patients. Because the isoform axis emerged from comparisons that began with the morphological groups, and two of the 33 comparisons (analyses 49 and 52) retain that morphological grouping, the morphology–isoform correspondence is not fully independent; the single-specimen split SI against a common reference, which does not use morphology, and the four morphology-deviating specimens provide partial independent support (Section 3.1.2). Note that the terminology for histological types and lesion names in this manuscript follows the WHO classification of tumors of the digestive system. In the sixth edition, published in 2026, recognition of serrated polyps advanced in the chapter on colorectal tumors, and they were divided into three types: hyperplastic polyp, sessile serrated lesion (SSL), and traditional serrated adenoma (TSA) [34]. This manuscript does not grade the dysplasia of individual precursor lesions, and the lesion and pathway names used are the serrated pathway, serrated polyp, serrated lesion, and hyperplastic polyp; we do not use the old terms that were replaced in the sixth edition (such as SSA/P). Therefore, this revision does not require any change to the descriptions in this manuscript.

Further limitations concern the p53-activity and genome-wide analyses (Sections 3.1.4 and 3.2 of the main text) and the composition of the discovery cohort. Because the P2-dominant group was defined partly through p53-target differences, the separation of wild-type P2-dominant from mutant specimens by the p53 activity score (AUC 0.99) may partly reflect circularity (Section 3.2.1 of the main text). Variant tables were available for only 35 of the 63 discovery samples; the others were called from published mutation maps of the same biobank, and seven samples remained undetermined (Section 2). Because TP53 status and promoter usage largely coincided in the discovery cohort, the organoid analyses there cannot fully separate p53 activity from TP53 genotype; the conclusion that morphology follows TP53 status rather than promoter usage rests on adjusted analyses in 56 specimens (specimen level only), the independent biobank and bulk tumors (Sections 3.1.4 and 3.1.5 of the main text), was reached in analyses added after the per-specimen TP53 calls had been finalized, is observational, and cannot exclude that TP53 mutation marks the chromosomal-instability pathway more broadly than the 20q, aneuploidy, MSI, BRAF and CIMP covariates capture; within MSI tumors, TP53 status was not associated with the morphology genes, and among non-MSI, BRAF wild-type, CIMP-low tumors the morphology difference was marginal after adjustment for 20q. Organoids lack stroma and immune cells and are grown in defined media; culture, dissociation and passaging could themselves modulate p53 activity and morphology, so p53 activity measured in organoids may not equal that in the tumor of origin. Chromosome-arm gains in the organoids were inferred from arm-level mean expression, not measured by copy-number profiling, and chromosomal instability was not measured directly. The predictive value of P2 beyond p53 activity is limited (on p53-target expression as a predictor of MDM2-inhibitor response, see [17–19]): in cell lines, the P2 index remained associated with nutlin-3a sensitivity and MDM2 dependency after adjustment for the p53 activity score in GDSC and CRISPR but not in PRISM (Section 3.2.4 of the main text), the association was not significant for nutlin-3 in CTRPv2, and in the 65 organoids of Herranz-Ors et al. the P2 signature was significantly associated with nutlin-3 sensitivity but not after adjustment for TP53 status (FDR = 0.056; Section 3.2.4); the association of P2 usage itself with MDM2 inhibitor sensitivity was examined in cell lines and in 14 TP53 wild-type organoids of an independent biobank (GSE294511), where it could not be separated from total MDM2 (Section 3.2.5 of the main text), but not in patients; MDM2 dependency was examined only in cell lines. In TCGA, the correlation between P2 and p53 activity across all wild-type tumors depended on expression quantification (ρ = 0.03–0.19) and was positive with both quantifications when restricted to non-hypermutated tumors whose wild-type status was confirmed in the second mutation call set; P2_index was also inversely correlated with an immune-cell marker (PTPRC; ρ = −0.47 in colorectal wild-type tumors) and may be influenced by tissue composition. P2_index and the cell-line P2 index are estimated from transcript quantification; direct junction counts agreed with them in TCGA tumors with 10 or more junction reads and in six cell lines with long reads (Section 3.2.2), but transcription start sites were not mapped in tumors or organoids. Generalization beyond colorectal cancer is outside the scope of this article. The proportion of wild-type tumors judged p53-inactive depends on the classifier: 55% here, with a threshold relative to mutant tumors; 9% with the conservative phenocopy threshold of Fito-Lopez et al. [20]; and 99% with the classifier of Li et al. [21], which is trained against adjacent normal tissue — although the continuous measures agree in direction (Section 3.2.3 of the main text). Adjacent normal mucosa uses mainly P1, and TP53-mutant tumors also exceeded it (Section 3.2.2 of the main text), so P2 usage in tumors also has inputs other than p53 activity. The public nutlin RNA-seq coverage is the sum of two replicates, so per-replicate ratios and tests were not obtained. Finally, the discovery cohort combines primary tumors with liver, lung, lymph node and ovarian metastases, and sampling site was not adjusted for in the P1–P2 comparisons; site-related differences, including the higher EMT scores of metastases (Section 3.1.2), may therefore confound part of the tissue-1/tissue-2 contrast.

In addition, in the reanalysis of 65 patient-derived organoid lines (Herranz-Ors et al.) presented in Section 3.2.4, the association between the P2 signature and nutlin-3 sensitivity remained in the same direction even in partial correlation adjusted for TP53 status, but it did not survive multiple-testing correction (ρ = −0.331, raw P = 0.0071, BH-adjusted FDR = 0.056). Because the p53 target module of P2 is a set of p53 target genes, its score is by definition lowered in TP53-mutant specimens. To break this circularity in principle, we performed a stratified analysis restricted to the 48 TP53-mutant lines, and although an association in the same direction was observed (ρ = −0.241, exact Spearman test P = 0.099, Benjamini–Hochberg-adjusted FDR = 0.775), it did not fall below the significance level. Because gene set scores are calculated by GSVA as relative quantities with respect to the sample set, we also performed a sensitivity analysis in which the scores were recalculated using only those 48 lines, but the conclusion that the association is in the same direction and does not fall below the significance level was unchanged (ρ = −0.281, P = 0.054, FDR = 0.564; Supplementary Table S41). With this sample size and effect size, statistical power at the two-sided 5% level is limited, and this result does not indicate the absence of an association. Therefore, this study shows that the P2 signature is associated with MDM2 inhibitor sensitivity, but it does not claim that this association is based on information independent of TP53 mutation status. Settling this point will require a prospective organoid cohort with a sufficient sample size in which TP53 status is matched. In addition, the colibactin analysis (Additional files 61, 62 and 63: Supplementary Figures S11 and S12, Supplementary Table S42) has the following limitations. (i) The axis in the Nunes cohort is the GSVA of gene-level expression (Type1_full − Type5_full), not the MDM2 P2_index itself (isoform quantification), because the public expression matrix is at the gene level (the coverage of Type5_full is 27/29 genes, and the Type5_lncRNA module (2 genes) could not be scored). (ii) The activities of SBS88 and ID18 are zero-inflated (3–8% positive), so the effect sizes are limited to rank correlations and positivity rates. (iii) TCGA-COAD/READ is mainly WES, and both ID18 (insertions and deletions) and SBS88 are detected with lower sensitivity than in WGS (11.0% and 13.6% activity-positive); only one test, P2_index × ID18, was significant in the direction of replication, and because this depends on how the axis is taken, we position it as an exploratory partial replication. In addition, in TCGA, even after supplementing per-patient MSI annotation from public GDC-derived annotation, only 32 of the 374 specimens for which both the axis and the activity were available had an annotation (MSS 24), so the MSS-restricted, covariate-adjusted sensitivity analysis performed in the Nunes cohort could not be carried out in an informative form (in the 24 MSS specimens, all tests had FDR = 1). (iv) All of these are cross-sectional associations, and the temporal relationship between the timing of exposure and the expression state cannot be evaluated. In the future, we plan to test whether TP53 loss itself drives the morphological change and whether MDM2 P2 usage improves on total MDM2 expression for selecting patients for MDM2 inhibitors, in prospective cohorts.

### R-24. Full-length version of main text Section 5 (Conclusions)

### 5. Conclusions

In this study, we compared in detail, by multilayered omics analysis, two types of CRC tissue that carry the same diagnostic name of colorectal cancer yet have diametrically opposite biological properties (tissue 1: CIN-type, MDM2 P1-dominant; tissue 2: MDM2 P2-dominant with active wild-type p53 and a signature-level resemblance to the MSI-like/serrated type). Morphological classification by deep learning (VGG16, test accuracy 98.5% (64/65)) corresponded with MDM2 isoform usage specimen by specimen — the P1 group was dominated by Type1 (compact glandular) morphology and the P2 group by non-Type1 (cystic–mucinous) morphology (median Type1 fraction 0.826 versus 0.444, P = 1.1×10⁻³ at the specimen level) — and, when compared with TP53 status, which largely coincided with promoter usage, morphology followed TP53 status beyond 20q and aneuploidy, also in an independent organoid biobank and in tumors (Sections 3.1.4 and 3.1.5 of the main text), whereas promoter usage reflects p53 activity.

[Tissue 1] is a tumor type, inferred from gene expression to be densely solid, that retains colorectal stemness (LGR5, OLFM4) and intestinal absorptive epithelial identity and is characterized by autonomous cell cycle acceleration through the FOXM1-MYC-E2F axis, comprehensive accumulation of pre-mRNA/rRNA processing, DNA replication, and oxidative phosphorylation, and maintenance of apico-basal polarity, in which MDM2 is predominantly transcribed from the constitutive P1 promoter (for details, see Table 5 and Section 3.3.1).

[Tissue 2] is a secretory, mucin-rich adaptive tumor (high viscoelasticity is inferred from the expression of gel-forming mucins; no mechanical measurements were performed) that shows lineage conversion toward gastric metaplasia (CTSE, REN, TFF1/3, ANXA10), the small-intestinal Paneth cell type (DEFA5/6), and high expression of gel-forming mucins (MUC5B/5AC/6), and shows MDM2 P2 dominance accompanying transcriptionally active wild-type p53. It is accompanied by simultaneous high expression of 10 p53 target genes (all ≥91% consistent) and by directionally consistent higher expression of 7 immune checkpoint molecules (exploratory; 6 of which were adopted in the P2 signature; the effect sizes of all 7 did not meet the DEG criteria, and the reproducibility of direction is the basis for their adoption; HAVCR2 has a consistency rate of 66.7% and does not meet the consistency criterion either) (Figure 8; for details, see Table 5 and Section 3.3.1). A 39-gene panel that well separates the two tissue types was identified (Supplementary Figure S5; the cluster assignment agreed with MDM2 promoter usage in 62 of 63 specimens, a within-sample, non-independent agreement because the genes were selected from the same comparisons); it includes wild-type TP53 target genes, gastric metaplasia markers (CTSE, REN), and MSLN, and, because it was selected from the same comparisons, it summarizes the p53-active/metaplastic contrast rather than providing an independent classifier, so any RT-qPCR/NGS assay derived from it would require independent validation.

In independent external validation using TCGA-COAD/READ (n = 624) and GSE39582 (n = 519; 1,143 cases in total), the dMMR predominance of the P2 signature, the highest value in CMS2 of the differentiation core of the P1 signature (13 genes, excluding the proliferation module), were observed in the external cohorts (Section 3.3.2); the known CMS-wise survival differences were also observed (Section 3.3.2), but they serve as a positive control rather than as a validation of the present findings. In multivariate Cox analysis, the P1 score remained only at a trend toward association with favorable prognosis under adjustment for the P2 score and did not show independence after adjustment for stage (Section 3.3.2). In addition, direct quantification of the MDM2 P2_index was significantly higher in TP53 wild-type cases (Section 3.2.2); in the cell line panel and in DepMap CRISPR screening, wild-type TP53 lines were more sensitive to MDM2 inhibitors and more MDM2-dependent (in colorectal cancer cell lines, high sensitivity was also seen on the MSI-High side; Section 3.2.4); and in comprehensive sensitivity prediction across the 198 drugs included in GDSC2, nutlin-3a ranked first both in correlation with the P2 score and in separation (a top rank expected in part from the p53-target content of the P2 signature) (to be interpreted as a rank rather than as the absolute smallness of the P value), and first of 295 drug entries on measured cell-line sensitivity after adjustment for cancer type and TP53, whereas ATR/CHK1/WEE1 checkpoint inhibitors tracked the P1 score (Section 3.2.4). Furthermore, in an independent patient-derived colorectal organoid biobank of 65 lines, TP53 wild-type lines were also significantly more sensitive to nutlin-3 (Section 3.2.4). These results are consistent with an alignment of CRC phenotypes with TP53 status and with established TP53-dependent MDM2 inhibitor sensitivity; whether the P1/P2 axis adds predictive value beyond TP53 status in patients remains to be tested.

The main contributions of this study can be summarized as follows: first, the alignment of colorectal cancer organoid morphology with TP53 status — compact glandular (Type1) in TP53-mutant and round, cystic–mucinous in wild-type organoids — beyond 20q and aneuploidy, in 56 discovery specimens (specimen level), in an independent organoid biobank (GSE294511; also on unadjusted patient means) and in tumors, with which the specimen-level association, in 21 patients and 63 samples, between morphology and MDM2 mRNA isoforms (the P1 group Type1-dominant and the P2 group non-Type1) is consistent; second, the proposal of MDM2 P1/P2 promoter usage as a transcript-level readout of p53 activity, whose p53-target program followed P2 beyond TP53 status, whereas the gastric-metaplastic program followed TP53 status (in the independent biobank and in tumors); and third, external validation in two independent cohorts (n = 1,143 cases in total), in which the organoid-derived P1/P2 signatures tracked CMS and mismatch-repair status, and promoter usage itself (P2_index, TCGA) tracked TP53 status (for details of the novelty, see Section 4.6). In addition, wild-type P2-dominant organoid specimens were p53-active (AUC 0.99 against mutants), as were the two P2-dominant specimens that carried TP53 mutations, so that P2 dominance identified the state in which p53 is actually active, P2 was induced in a p53-dependent manner by nutlin-mediated p53 activation, P2 tracked p53 activity in non-hypermutated wild-type tumors (though not beyond total MDM2) and in cell lines, and among TP53 wild-type cell lines P2 tracked MDM2 dependency beyond both p53 activity and total MDM2, and nutlin-3a sensitivity beyond p53 activity but only weakly beyond total MDM2 (Section 3.2 of the main text); organoid morphology followed TP53 status beyond expression-inferred 20q, and its gene program identified mucinous carcinoma in patient tumors (Section 3.1.4 of the main text). In an independent biobank of 119 organoids from 50 patients (GSE294511), the morphology program again followed TP53 status, its across-sample alignment with P2 being largely explained by TP53 status, and in TP53 wild-type organoids P2 tended to remain associated with p53 activity after adjustment for total MDM2 (Sections 3.1.5 and 3.2.5 of the main text). The P1/P2 ratio reflects p53 activity, not only TP53 genotype; whether it improves on MDM2 expression itself for selecting tumors for MDM2 inhibitors — suggested for p53 activity in the independent organoids (not in the discovery organoids) and for MDM2 dependency in cell lines, but weak or absent for nutlin-3a sensitivity in cell lines, idasanutlin sensitivity in an independent organoid biobank (GSE294511) and p53 activity in bulk tumors — requires prospective validation.

### R-25. Full-length version of main text Sections 3.2.1, 3.2.3 and 3.2.4 (MDM2 P2 promoter usage and p53 activity)

#### P2 dominance marks a state in which p53 is actually active (Figures 3, 4, 6 and 7)

In Section 3.1.3, TP53 status and promoter usage largely coincided. Because the P2 signature contains p53 targets (Section 3.3.2), we measured whether p53 is actually active, rather than TP53 genotype, directly from 12 direct p53 targets (p53 activity score; Section 2).

##### P2-dominant organoid specimens are p53-active

Among the 56 discovery specimens with determined TP53 status, the median p53 activity score was −0.30 in 42 TP53-mutant specimens, −0.35 in three wild-type P1-dominant specimens and +1.45 in 11 wild-type P2-dominant specimens (Figure 3A). Wild-type P2-dominant specimens were p53-active compared with mutants (P = 6.8×10⁻⁷, AUC 0.99; at the patient level [means per patient and group], P = 5.2×10⁻⁴), and none of them had a score at or below the 90th percentile of mutants, whereas two of the three wild-type P1-dominant specimens, both from HCT59 (variant of uncertain significance F113V), did (wild-type P2-dominant versus wild-type P1-dominant specimens, P = 0.005). Because TP53 status and promoter usage largely coincided (P1-dominant, 40 mutant and 3 wild-type; P2-dominant, 2 mutant and 11 wild-type; Section 3.1.3), the organoids cannot separate p53 activity from TP53 genotype. Two observations nevertheless favor activity: the only mutant specimens with clearly positive scores were the two P2-dominant ones (HCT41-1T, +1.09; HCT38-3LM, +0.60; both above the 90th percentile of mutant specimens), and across the 56 specimens the P2 index correlated with the p53 activity score (ρ = 0.53, P = 2.6×10⁻⁵). Within the 14 wild-type specimens, the P2 index and the p53 activity score were correlated (ρ = 0.69, P = 0.0065; Figure 3B), but total MDM2 expression correlated to a similar degree (ρ = 0.71), and neither P2 dominance nor the continuous P2 index added significantly to total MDM2 (R² 0.50 to 0.59, P = 0.15; 0.50 to 0.52, P = 0.50; Additional file 7: Supplementary Table S48). In each of the three patients with both P1 and P2 specimens (HCT38, HCT41 and HCT67), the P2 specimen had the highest p53 activity score (Figure 3C). The p53 suppressors MDM4, PPM1D and USP7 did not differ significantly between groups. Assigning the seven specimens with undetermined TP53 status (all P1-dominant) to the mutant group left these results unchanged; assigning them to the wild-type group gave 10 wild-type P1-dominant specimens, eight at the mutant level (wild-type P2-dominant versus P1-dominant, P = 1.2×10⁻⁴; P2 dominance beyond total MDM2, P = 0.021; TP53 calls in Additional file 5: Supplementary Table S11; values in R148_results.csv in Additional file 2). Because the P2 signature shares genes such as CDKN1A, BAX and BBC3 with the p53 activity score, and the P2-dominant group was defined in part through differences in p53 target expression, the separation may partly reflect circularity.

In the 14 public HTA2.0 arrays not used for discovery (2 normal and 12 tumor; identical to GSE128213 files), the P2 index and the p53 activity score were again strongly correlated (ρ = 0.89, P = 1.6×10⁻⁵; tumors alone, ρ = 0.86, P = 3.3×10⁻⁴; Figure 3D). The 12 tumor arrays include two arrays with the same titles as discovery specimens of patient HCT33 (HCT33-4LMR and HCT33-5LMR; considered other arrays of the same specimens and therefore excluded from the replication in Section 3.1.2); both had a high P2 index and a high p53 activity score, as in the discovery cohort, where HCT33 was P2-dominant. The correspondence with the P2 signature in the same 10 tumor arrays (Section 3.1.2) is shown in Figure 3E.

##### P2 is selectively induced in a p53-dependent manner

To test whether P2 usage results from p53 activity, we counted promoter-specific reads at the MDM2 locus in public RNA-seq of cells treated with nutlin (10 µM, 12 h) [38] (Figure 4A; Section 2). In TP53 wild-type HCT116 cells, nutlin increased reads on the P2-only first exon 29.5-fold, whereas reads on the P1-only exon 1 increased only 2.8-fold, and the P2/P1 ratio rose 10.5-fold, from 0.28 to 2.91 (Figure 4B). In TP53-null HCT116 cells, P1 and P2 barely changed (1.13-fold and 1.11-fold), and neither did the P2/P1 ratio (0.156 and 0.153). MCF7 (P2/P1 ratio 0.11→14.4) and SJSA (0.23→4.73) cells showed the same direction, and under DMSO P1 reads predominated in every cell line. In gene-level counts, induction of MDM2 and CDKN1A was seen only in TP53 wild-type cells (Figure 4C). When p53 is active, MDM2 transcription shifts strongly toward P2, so that P2 usage becomes a readout of p53 transcriptional activity; this confirms, on the measures used in this study, the known status of P2 as a direct target carrying p53 response elements [5,6]. At the level of transcription initiation, ENCODE PRO-cap of six normal colon and stomach tissues gave a median P2 share of 0.506 (range 0.421–0.657), and TP53-mutant Caco-2 cells 0.565, whereas the P2 share of MDM2 mRNA for the same line in DepMap was 0.173 (Section 2; Additional file 7: sheet K); thus initiation at P2 was not reduced in this TP53-mutant line (interpretation in Section 4.2).

##### P2 and p53 activity in tumor tissue

In TCGA-COAD/READ, P2_index showed a gradient across TP53 functional classes (Section 3.2.2; Figure 6A). Across all 132 TP53 wild-type tumors, the correlation between P2_index and the p53 activity score was weak and depended on how expression was quantified (ρ = 0.19, P = 0.029 with TCGAbiolinks expression; ρ = 0.03, P = 0.73 with the Toil recompute; Additional file 7: Supplementary Table S48). In the 82 tumors that were not hypermutated and were confirmed as wild type in the MC3 mutation calls, P2_index and the p53 activity score were clearly correlated (ρ = 0.42, P = 1.0×10⁻⁴; ρ = 0.26, P = 0.016 with the Toil recompute), and the p53 activity score was higher in the middle and upper P2_index tertiles than in the lower tertile (medians 0.11, 0.72 and 0.64; upper versus lower P = 8.5×10⁻⁵; Figure 6B). In the same 82 tumors, however, total MDM2 expression (the sum of all MDM2 transcripts) tracked the p53 activity score more closely (ρ = 0.57), and adding P2_index to it did not improve the fit (R² 0.323 to 0.324, P = 0.71; P = 0.19 with the Toil recompute; Additional file 7: Supplementary Table S48); in bulk tumors, P2 usage did not carry information on p53 activity beyond the amount of MDM2 transcript. Across 12 combinations of TP53 call, hypermutation definition and expression source, ρ ranged from 0.03 to 0.42, and the P2_index added no significant positive information beyond total MDM2 in any of them (Additional file 7: Supplementary Table S48, sheet N). The same held at the protein level: in the 69 of these 82 tumors with reverse-phase protein array data (TCGA Pan-Cancer Atlas RPPA, obtained via UCSC Xena [43]), BAX protein, a p53 target, correlated with the p53 activity score (ρ = 0.48, P = 3.0×10⁻⁵), P2_index (ρ = 0.33, P = 0.0059) and total MDM2 (ρ = 0.39, P = 9.3×10⁻⁴), but its association with P2_index did not persist after adjustment for total MDM2 (partial ρ = 0.09, P = 0.46; Additional file 7: Supplementary Table S48). The 12-target p53 activity score also agreed with a published transcriptome-based classifier of p53 function. In the 118 wild-type tumors with a TP53-loss phenocopy score (Fito-Lopez et al. [20]; an elastic-net model on 217 genes, six of which are among our 12 targets, whereas MDM2 is not), the two measures were inversely correlated (ρ = −0.49, P = 2.0×10⁻⁸), the p53 activity score discriminated phenocopy status with an AUC of 0.80, and 10 of the 11 tumors called phenocopies were among the 65 tumors with a score at or below the 90th percentile of mutant tumors. The classifier of Li et al. [21], trained against normal tissue, called 104 of the 105 wild-type tumors with a prediction (99%) functionally deficient (Additional file 7: sheet J). P2_index also correlated with the phenocopy score (ρ = −0.31, P = 7.5×10⁻⁴) but, as for the p53 activity score above, not after adjustment for total MDM2 (partial ρ = −0.09, P = 0.36) (Additional file 7: Supplementary Table S48, sheet J). Why some TP53 wild-type tumors lack p53 activity is beyond the scope of this paper and is the subject of a companion study on p53 activity in TP53 wild-type tumors (T. Tsukui, R. Yao, and K. Tsuda, unpublished observations). P2_index was inversely correlated with an immune-cell marker (PTPRC; ρ = −0.47 in wild-type tumors) but only weakly with tumor purity (ρ = 0.12), and bulk P2_index may be influenced by tissue composition. Generalization beyond colorectal cancer is outside the scope of this article (Section 4.6).

##### P2 tracks MDM2 dependency and, less specifically, MDM2 inhibitor sensitivity

From DepMap transcript quantification [47,48] we computed the P2 index of 1,517 cell lines; in the 691 lines with known TP53 status it was higher in wild-type lines (median 0.231 versus 0.104, P = 1.8×10⁻³³) and correlated with the p53 activity score among the 208 wild-type lines (ρ = 0.57, P = 1.4×10⁻¹⁹; Figure 7A). In six of the 1,517 lines that were also sequenced with short and Nanopore long reads [58], the P2 share counted directly from MDM2 junction reads ranked the lines in nearly the same order as this transcript-based P2 index (Spearman ρ = 0.94 with short reads and 0.83 with long reads; lowest in the TP53-mutant line K562 and highest in HepG2), and over all SG-NEx long reads, reads beginning in exon 2, the P2 first exon, carried the P2 junction (948 of 961), whereas those beginning in exon 1 carried the P1 junction (2,131 of 2,289) (Additional file 7: Supplementary Table S48). In TP53 wild-type lines, a higher P2 index was associated with greater sensitivity to nutlin-3a (GDSC1, 198 lines, ρ = −0.26, P = 1.7×10⁻⁴) [49,50], and the association persisted after adjustment for the p53 activity score (partial correlation −0.19, P = 0.009; GDSC2, −0.15, P = 0.034; Figure 7B). MDM2 dependency in CRISPR knockout screens [52,54] was also greater in wild-type lines with a higher P2 index (364 lines, ρ = −0.33, P = 8.1×10⁻¹¹; tertile medians −0.77, −1.19 and −1.24; mutant lines −0.32) and remained, weakly, after adjustment for the p53 activity score (partial correlation −0.12, P = 0.022; Figure 7C). Total MDM2 expression, however, was associated with nutlin-3a sensitivity about as strongly as the P2 index (GDSC1 ρ = −0.26; GDSC2 ρ = −0.17), and after adjustment for total MDM2 the association of the P2 index weakened (GDSC1 partial correlation −0.15, P = 0.035, and −0.13, P = 0.077 after adjustment for both total MDM2 and the p53 activity score; GDSC2 −0.05, P = 0.43), as it did for nutlin-3 in CTRPv2 (161 wild-type lines [55,56]; −0.02, P = 0.76). For MDM2 dependency, by contrast, the P2 index remained associated after adjustment for total MDM2 (ρ = −0.29 for total MDM2; partial correlation of the P2 index −0.27, P = 1.1×10⁻⁷; −0.11, P = 0.041 after adjustment for both; Additional file 7: Supplementary Table S48).

This relationship was specific to the mechanism of action. In the PRISM drug screens [51], all eight agents that block the MDM2–p53 interaction and release p53 (idasanutlin, AMG-232, CGM097, SAR405838, RG7112, milademetan, nutlin-3 and siremadlin) were in the negative direction; in the primary screen (24Q2), seven reached P < 0.05 (ρ = −0.17 to −0.30; siremadlin, ρ = −0.11, P = 0.055), and in the secondary screen the association was strongest for idasanutlin (ρ = −0.31 and −0.34). Drugs with other mechanisms (RITA, serdemetan, SP-141 and others) showed no association (Figure 7D). In PRISM, however, the association largely disappeared after adjustment for the p53 activity score. Most of the information in P2 thus overlaps with p53 activity, and the part beyond it was seen only in GDSC and CRISPR. TP53 wild-type lines without p53 activity (about 30%; in this analysis the p53 activity score used the 10 genes with available expression) were less sensitive to nutlin-3a (GDSC1: median ln IC50 3.12 versus 1.74, P = 3.8×10⁻⁷), and this held in lines confirmed as wild type by DepMap mutation data alone, so it is not explained by the misclassification of mutant lines as wild type that has been raised as a concern [18]. For nutlin-3 in CTRPv2, the association within wild-type lines did not reach significance (161 lines, ρ = −0.14, P = 0.07; Additional file 7: Supplementary Table S48). Whether P2 usage tracks MDM2-inhibitor sensitivity in organoids from independent patients is examined in Section 3.2.5.

### R-26. Full-length version of main text Section 3.1.4 (Morphology, TP53 status and chromosomal instability)

#### Morphology and, in the independent biobank and tumors, the metaplastic genes follow TP53 status rather than promoter usage, and the MDM2 axis follows p53 activity (Figure 2)

Because organoid morphology corresponded to MDM2 promoter usage (Section 3.1.2) and promoter usage largely coincided with TP53 status (Section 3.1.3), we asked whether morphology follows promoter usage or TP53 status (this paragraph and the next). Morphology and the MDM2 axis were weakly correlated (non-Type1 fraction and P2 index, ρ = 0.33). To disentangle them, we matched 39,438 sufficiently expressed transcript clusters against both axes by rank partial correlation (Section 2). Of the 124 clusters associated with morphology only, 44 lay on the long arm of chromosome 20 (20q) (background 1.9%, P = 6.5×10⁻⁴⁴), all higher on the compact glandular side. The 930 clusters associated with the MDM2 axis only were centered on p53 targets (FAS, DDB2, RPS27L, ZMAT3 and others; 83% remained after adjustment for the p53 activity score) and were enriched on 8p, 8q and 4q, and weakly on 16p (Figure 2A). In mean expression by chromosome arm, the expression features of chromosomally unstable colorectal cancer (expression-inferred gains of 20q and 8q; losses of 8p, 18q and 4q) [89] aligned with the compact glandular, P1-dominant side, and the three P2-dominant patients lacked 8q and 20q gains. The correlations of arm-level mean expression with the two axes were of nearly the same direction and magnitude (20q: morphology −0.34, P2 index −0.39; 8q: −0.30, −0.46; 8p: +0.29, +0.31; Figure 2B). Thus, at the level of transcript clusters, morphology corresponded preferentially to 20q and the MDM2 axis to p53 targets, whereas at the arm level both axes aligned with the same chromosomal-instability pattern; their overlap can be explained by the dichotomy between chromosomally unstable and stable tumors. Genes higher on the cystic–mucinous side were enriched for four of the ten prespecified cytoskeleton and adhesion GO terms (actin cytoskeleton, 5/42 genes, odds ratio 6.1, q = 0.021; adherens junction, 5/42, odds ratio 3.7, q = 0.038; cell adhesion, 8/42, q = 0.048; cytoskeleton, 9/42, q = 0.048), the recurring genes being VCL, AHNAK, TRIOBP, FYN and CDC42BPB, with no enrichment among the 73 genes on the opposite side (Figure 2C). No exon usage corresponded to morphology.

Here and below, the organoid morphology genes are the 141 transcript clusters whose expression tracked morphology — the 124 associated with morphology only and the 17 associated with morphology and the MDM2 axis in the same direction — and the morphology gene score is the difference in expression between their two sides (Section 2; each cluster, with its correlations, chromosome arm and use in TCGA, is listed in Additional file 7: Supplementary Table S48, sheet O, and the gene symbols of each side are listed together at the end of that sheet, O3). They are a data-driven summary of the morphological axis rather than a predefined pathway, and the two sides differ in character. The 44 clusters higher on the cystic–mucinous (non-Type1) side are almost all protein coding (41), none lies on 20q, and they include genes of the actin cytoskeleton, cell adhesion and membrane trafficking (VCL, AHNAK, TRIOBP, FYN, CDC42BPB, INF2, ARHGAP5, PLEKHG3 and MICALL1) and the tetraspanins TSPAN14 and TSPAN15. The 97 clusters higher on the compact glandular (Type1) side include 49 on 20q, whose expression rises with 20q gain — among them genes of RNA processing and translation (RALY, PRPF6, EIF2S2 and EIF6), the proteasome (PSMA7), mitochondria (ATP5E), nuclear transport (CSE1L and RAE1) and glycosylation in the endoplasmic reticulum (DPM1, RPN2 and PIGU) — and 21 without a gene annotation. The score thus contrasts a cytoskeleton–adhesion program with the expression consequence of 20q gain, and because no exon usage corresponded to morphology, it marks the expression state that accompanies a morphology rather than genes shown to cause it.

The organoid morphology genes also corresponded to histology in patient tumors. In 372 primary TCGA-COAD/READ tumors, the difference in expression between genes higher on the cystic–mucinous side (39) and those higher on the compact glandular side (64) (morphology gene score) distinguished mucinous carcinomas (43 tumors; AUC 0.83, P = 1.4×10⁻¹²; Figure 2D), and 97 of the 103 genes showed the same direction as in organoids. This correspondence was not explained by MSI (AUC 0.87, P = 6.8×10⁻⁸ in chromosomally unstable tumors alone); the score ran opposite to the amount of 20q (ρ = −0.52) and was unrelated to the p53 activity score within TP53 wild-type tumors (ρ = −0.08, P = 0.37; 119 tumors; Supplementary Table S48, sheet H2), but it was higher in TP53 wild-type than in mutant tumors (medians 0.67 versus −0.31, P = 2.5×10⁻¹⁶; 358 tumors with MC3 calls; Figure 2E), within the CIN (P = 3.0×10⁻⁵) and GS subtypes (P = 0.006) but not within the MSI subtype (P = 0.37), and after adjustment for 20q and MSI (P = 1.5×10⁻⁴; 335 tumors); mucinous carcinomas were more frequent among wild-type tumors (26 of 119 versus 16 of 239, P = 7.8×10⁻⁵), in line with the lower frequency of p53-pathway alterations reported in mucinous colorectal carcinomas [14]. In contrast, P2_index did not differ between mucinous and other carcinomas (0.413 versus 0.428, P = 0.35) or among molecular subtypes (CIN, GS and MSI; P = 0.11), and after adjustment for TP53 status, 20q and MSI it ran opposite to the morphology gene score (P = 1.9×10⁻¹¹), possibly reflecting tissue composition (Section 3.2.3). In the discovery organoids, likewise, the Type1 fraction differed by TP53 status after adjustment for 20q and aneuploidy (P = 0.013), and P2 dominance did not contribute beyond TP53 status and 20q (P = 0.97; Additional file 7: Supplementary Table S48), a test with little power because the two axes disagreed in only five of the 56 specimens; in the patient-level sensitivity analysis (patient means; 19 patients with uniform TP53 status, five wild type), P = 0.13. To ask which expression programs follow TP53 status and which follow P2 usage, we scored previously defined gene sets in the discovery organoids, the GSE294511 organoids (Sections 3.1.5 and 3.2.5) and TCGA, and compared TP53 status adjusted for the P2 measure with the P2 measure adjusted for TP53 status (rank regression; Section 2; Figure 2F; Additional file 7: Supplementary Table S48). With TP53 status adjusted for the P2 measure, the morphology gene score and the image-based non-Type1 fraction followed TP53 status (discovery P = 0.0063 and 0.013 in these models without covariates, distinct from the covariate-adjusted P = 0.013 above; GSE294511 P = 6.1×10⁻⁵; TCGA P = 3.1×10⁻¹⁹), as did the gastric-metaplasia genes of the P2 signature and the nine-gene gastric set in GSE294511 (P = 0.0051 and 6.0×10⁻⁴) and TCGA (P = 9.9×10⁻¹⁰ and 3.0×10⁻⁶), but not in the discovery organoids (56 specimens, 14 wild type; P = 0.32 and 0.64); in TCGA, after further adjustment for 20q and MSI, the metaplasia genes still followed TP53 status (P = 3.2×10⁻⁵), whereas the nine-gene gastric set did not (P = 0.18; Supplementary Table S48, sheet H2, covariate-adjusted columns). With the TP53 calls of Additional file 6: Supplementary Table S47 (327 tumors, 113 wild type), further adjustment for BRAF mutation and for the CpG island methylator phenotype (CIMP, called from the 450K arrays; Section 2) left the TP53 coefficients essentially unchanged (morphology gene score, +0.110 before and +0.109 after adding CIMP, P = 5.8×10⁻⁵; metaplasia genes, +0.136 and +0.134, P = 1.8×10⁻⁴), whereas the nine-gene gastric set was associated with CIMP-high status (P = 7.3×10⁻⁶) and not with TP53 status (P = 0.24); among tumors that were not of the MSI subtype, BRAF wild-type and CIMP-low (202 tumors, 47 wild type), the morphology gene score and the metaplasia genes remained higher in wild-type tumors (Mann–Whitney P = 5.8×10⁻⁵ and 4.8×10⁻⁴), although after adjustment for 20q the morphology difference was marginal (P = 0.069; metaplasia genes P = 0.012). None of the morphology, metaplasia, gastric, P1 or SNORD116 scores was significantly positively associated with the P2 measure after adjustment for TP53 status or within wild-type samples (in the discovery organoids the coefficients were positive but not significant, and in TCGA several were negative). The p53-target genes of the P2 signature (MDM2 itself excluded), by contrast, followed the P2 measure beyond TP53 status in both organoid cohorts (P = 5.8×10⁻⁴ and 2.7×10⁻⁵; within mutant organoids, ρ = 0.38 and 0.33; for the 12-gene p53 activity score, P = 0.094 and 0.012), whereas in bulk tumors they ran opposite to P2_index after adjustment for TP53 status (P = 0.0028; within TP53-mutant tumors the P2_index was inversely correlated with the 12-gene p53 activity score, ρ = −0.347, P = 3.7×10⁻⁸). These results held at a false discovery rate below 0.05 across the programs of each cohort. The P1 signature did not follow TP53 status consistently across cohorts, and the SNORD116 cluster was not associated with P2 in either organoid cohort nor consistently with TP53 status (discovery P = 0.92; GSE294511, slightly lower in wild type, P = 0.038; not measurable in poly(A) RNA-seq). The organoid morphology program is thus reproduced in patient tumors as the pathological histotype of mucinous carcinoma and follows TP53 status beyond chromosomal instability, as do the metaplasia genes in tumors, whereas the MDM2 axis follows p53 activity in organoids.

In the discovery organoids, we also asked whether the genome-wide expression differences between tissue 1 and tissue 2 (the 1,890 DEGs) follow TP53 status or promoter usage. Because TP53 status and promoter usage disagreed in only five of the 56 specimens with determined status, from four patients (HCT38-3LM and HCT41-1T, P2-dominant and TP53-mutant; HCT59-3LM, HCT59-4LM and HCT71-3LM, P1-dominant and wild type), this comparison has little power to separate the two and is exploratory. Across 19,676 expressed genes, the expression differences by TP53 status and by promoter usage were closely correlated (Spearman ρ = 0.885). Of the 1,890 DEGs, the 1,600 represented in the gene-level data (815 higher in tissue 1 and 785 in tissue 2) kept their direction in 92.7% when compared by promoter usage and in 94.3% when compared by TP53 status, and in models with both terms (patient-clustered robust standard errors; Section 2) the median DEG retained 70% of its promoter-usage coefficient and 52% of its TP53 coefficient; genome-wide, 55 genes followed TP53 status beyond promoter usage and 145 followed promoter usage beyond TP53 status (q < 0.05), and among the DEGs only 14 and 33, respectively. Of the 13 p53-target genes whose expression predicted sensitivity to the p53–MDM2 inhibitors NVP-CFC218 and NVP-CGM097 in cancer cell lines, better than TP53 status alone, and to NVP-CGM097 in patient-derived xenografts [17], 8 were among the 145 genes following promoter usage beyond TP53 status (BAX, DDB2, MDM2, RPS27L, RRM2B, TNFRSF10B, XPC and ZMAT3; one-sided Fisher's exact P = 8.9×10⁻¹⁵; 7 of 12 without MDM2, P = 7.5×10⁻¹³) and none among the 55 following TP53 status (Figure 2G). Because promoter usage was called from MDM2 exons and P2 itself is p53-inducible, this enrichment is not independent of the grouping; it agrees, however, with the program-level result above, whereas the bulk of the DEG program could not be assigned to either axis in this cohort (Additional file 7: Supplementary Table S48, sheet H).

### R-27. Full-length version of main text Sections 3.1.5 and 3.2.5 (Replication in an independent organoid biobank)

#### Replication in an independent organoid biobank (GSE294511)

##### Morphology and P1/P2 usage

To test the main correspondences in independent patients, we analyzed the public HTA2.0 arrays of 119 organoids from 50 patients with colorectal liver metastases, established at another institution (Kryeziu et al. [36]; GEO GSE294511), under an analysis plan fixed before the data were examined (Section 2). Organoids whose morphology gene score lay further on the non-Type1 side had a higher P2 index (Spearman ρ = 0.22, one-sided P = 0.0085; 0.22 without the 20q genes), also when the score was built only from the 124 clusters associated with morphology alone (ρ = 0.23, one-sided P = 0.0057). Patient means were correlated (ρ = 0.30, P = 0.037, 50 patients), but not significantly without the 20q genes (ρ = 0.23, P = 0.11). In a model with a patient random intercept, the slope was not significant (−0.05, P = 0.68), the between-patient variance being 4.6 times the residual variance; among the 34 patients with more than one organoid, the morphology gene score and the P2 index moved in the same direction in 16, in opposite directions in 17 and were tied in one, and among the 15 patients whose H&E morphology differed between organoids, the more solid organoids had the higher P2 index in 9 and the lower in 6 (sign test P = 0.61). The correspondence therefore held across samples, was weaker between patients and was not detected within patients, whereas in the discovery cohort, from which the score was derived, the gene score (unlike the image-based fractions; Section 3.1.2) was also associated with P2 within patients (random-intercept slope +0.46, P = 0.0019). The H&E-based morphology categories published with the biobank (cystic 32, mixed 50 and solid 37 organoids) did not differ in P2 index (medians −1.03, −0.80 and −0.66; Kruskal–Wallis P = 0.38; random-intercept model P = 0.56), and the morphology gene score tended to be higher in solid organoids (P = 0.064). The two classifications overlap only partly: the difference between CDX2 and KRT20 mRNA, whose proteins distinguished cystic from solid organoids in that study, was unrelated to the non-Type1 fraction in the discovery cohort (ρ = 0.08, P = 0.55), but in the biobank it was lower toward the P2 index (ρ = −0.40, P = 8.9×10⁻⁶) and toward the non-Type1 side of the morphology gene score (ρ = −0.47). The 20q genes, higher on the Type1 side, again moved opposite to the rest of the morphology gene score (ρ = −0.63; discovery −0.88). Morphology followed TP53 status in this biobank as well: among the 88 organoids with published TP53 status, the morphology gene score lay further on the non-Type1 side in wild-type organoids (medians 1.11 versus −0.19, P = 1.9×10⁻⁵; Figure 2E; patient means, P = 0.0029, 49 patients with uniform status), also after adjustment for 20q and aneuploidy (P = 6.1×10⁻⁴), and the P2 index did not contribute beyond TP53 status, 20q and aneuploidy (P = 0.51); within TP53 groups the P2 index was unrelated to the score (wild type ρ = 0.27, P = 0.35; mutant ρ = 0.07). Given these within-group correlations, the across-sample correspondence between morphology and P2 described above is largely explained by TP53 status (Additional file 7: Supplementary Table S48).

##### p53 activity and idasanutlin sensitivity

The P2 index was higher in TP53 wild-type than in mutant organoids (medians 1.18 versus −0.89, P = 6.9×10⁻⁴) and, within the 14 wild-type organoids, was strongly correlated with the p53 activity score (ρ = 0.91, P = 5.5×10⁻⁶; discovery 0.69; mutant organoids, ρ = 0.05). Total MDM2 also correlated with the score (ρ = 0.87), but the P2 index remained associated after adjustment for it (partial ρ = 0.61, P = 0.028; patient means, 0.66, P = 0.053, 10 patients). The authors of the biobank reported that the MDM2 inhibitor idasanutlin was more active in TP53 wild-type organoids [36]; among the wild-type organoids, sensitivity followed the P2 index (ρ = 0.70, P = 0.0049): the six with the lowest P2 index were insensitive (scores 0–9.1) and the eight with a P2 index above 0.3 were sensitive (19.9–31.4). Across the 87 organoids with known TP53 status and idasanutlin sensitivity score (49 patients), the P2 index was associated with sensitivity after adjustment for TP53 status (coefficient +3.17, P = 1.6×10⁻⁸; 0.0022 with standard errors clustered by patient), and after further adjustment for the p53 activity score and the mean sensitivity to the 23 other drugs the coefficient was +1.85 (P = 0.0013; clustered, 0.069); P2-leaning organoids were also somewhat more sensitive to some other drugs (gemcitabine ρ = 0.33, SN-38 ρ = 0.27). Among wild-type organoids, however, total MDM2 tracked idasanutlin sensitivity as closely (ρ = 0.69), and after adjustment for it the P2 index was no longer associated (partial ρ = 0.26, P = 0.39); in the regression across the 87 organoids, total MDM2 remained (+4.80, clustered P = 0.006), whereas the P2 index was weaker (+1.59, P = 0.033; clustered, 0.16). Thus, in organoids from independent patients, P2 tended to remain associated with p53 activity after adjustment for total MDM2 (P = 0.028 across organoids, 0.053 on patient means), whereas for MDM2-inhibitor sensitivity it could not be distinguished from total MDM2 among the few wild-type organoids, similar to the cell-line results for nutlin-3a (Section 3.2.4).

##### Exon usage and the direction of gene expression

The exon- and junction-level differences that accompanied P2 usage in the genome-wide exon-usage screen of the discovery cohort (39 probe sets in 11 genes, excluding the probe set used to define the P2 index; including FAS, ZMAT3, TNFRSF10C and ACTA2) all changed in the same direction with the P2 index (39 of 39, P = 3.6×10⁻¹²; 38 of 38 without the MDM2 probe set). Of the 1,890 DEGs, 65% changed in the same direction with the P2 index (P = 1.5×10⁻⁴⁰; 175 of the 187 with |ρ| > 0.3), as did 31 of the 36 genes of the 39-gene panel identifiable by symbol (panel score, ρ = 0.47, P = 6.4×10⁻⁸), and the gastric-metaplasia score was again higher toward P2 (ρ = 0.22, P = 0.017; discovery 0.23). In the 50 matched tumor tissues, by contrast, the P2 index was higher than in the organoids (medians 0.35 versus −0.96) and its ranking agreed poorly with that of the organoids (ρ = 0.15, P = 0.29), although within tissues the morphology gene score and the P2 index were strongly correlated (ρ = 0.70); bulk tissue contains stroma and immune cells (Section 3.2.3).
