## Additional files for "TP53 status aligns with morphology and metaplastic programs in colorectal cancer organoids, and MDM2 P2 promoter usage reflects p53 activity": Additional_file_12_Supplementary_Information.docx

*Detailed data that complement the principal findings of the main text. Supplementary sections that support statements in the main text are referred to from there; the others give additional detail. Separately from this Supplementary Information, an independent supplementary document, the Supplementary Note ("Supplementary Note: Molecular landscape of the two organoid groups"; Additional file 14), contains the details (methods, results, and discussion) of the IPA and EnrichR molecular pathway landscape summarized in the main text; it is submitted together with this Supplementary Information (Additional file 12) and the Supplementary Tables. The figure, table, supplementary table, supplementary figure, and reference numbers cited in the Supplementary Note are those of the main text; references not cited in the main text are listed at the end of each document that cites them, with numbering continuing from the main text. Where this Supplementary Information refers to "Supplementary Note Section N," a section of that document is meant. In addition, an independent supplementary document, the Supplementary Report (Additional file 1), provides the unabridged full-length versions of the main-text sections presented in condensed form in the article, and is likewise submitted together with these files. As in the main text, “tissue 1” and “tissue 2” denote the MDM2 P1-dominant and P2-dominant specimen groups, which largely coincide with TP53-mutant and TP53 wild-type specimens, respectively; the descriptive labels “autonomous-proliferation type” and “environment-adaptive type” summarize their expression programs and do not imply that promoter usage determines them.*

### **Supplementary Results 1. IPA Tox Function analysis (tissue 1: organ-spanning activation topped by nephritis)**

*Corresponds to Section 3.3.1 of the main text and Supplementary Note Section 2.4.*

In the IPA Tox Function analysis, the following biological signals accumulated as the toxicity profile of tissue 1. The 1st-ranked term was "Nephritis" (30/33 consistent, first-half Rank1), and "Glomerulonephritis" followed in 5th place (32/33 consistent). These are signals related to the inflammatory and immune responses of renal tissue. WNT, BMP, and Notch signaling are shared by kidney development and intestinal development, and the appearance of the Nephritis signal in colorectal cancer is interpreted as reflecting the tumor cells' use of these shared developmental signals. Indeed, the Nephritis gene set contains TGFB1, MDM2, TP53, and AGT. In this study, all of these are principal molecules on the tissue-2 side, and the fact that the Tox Function gene sets contain the principal molecules of both tissue types across the board is one reason why high scores arise across modules. The membership of TGFB1, MDM2, TP53, and AGT in the Nephritis gene set is corroborative only. Section 3.3.1 and Table 5 of the main text read Nephritis, together with Hyperplasia of kidney cells, as part of the proliferative signature of tissue 1 on the basis of the direction of its z-score (predicted activation in the tissue-1 direction; median z = +2.47), not on the basis of these member genes. Also accumulated were Hyperplasia of kidney cells (6th place; 25/27 consistent), Hyperplasia of mesangial cells (7th place; 29/31 consistent), Nephromegaly, Hypertrophy of kidney, Familial congenital heart disease, Regeneration of cardiac muscle, Congenital heart disease, Enlargement of cardiomyocytes, Fibrosis of myocardium, Hepatic steatosis (a fatty-liver-related signal), and Ventricular septal defect. Of these signals, only Nephritis (median z = +2.47) and Hyperplasia of kidney cells (+2.00) meet the adoption criteria (sign consistency ≥ 75% and |median z| ≥ 2); the others, including Glomerulonephritis (+1.47), are below threshold in effect size or sign consistency (Supplementary Note Section 2.4; Table 5 of the main text), and, like all Tox Function findings, these signals serve only as corroborating evidence. The accumulation of kidney and heart hypertrophy and hyperplasia signals may reflect aberrant activation within the tumor of WNT/β-catenin and BMP signaling, which are indispensable for the development of these organs. As Clevers reviewed in Cell (2006) [151], WNT signaling controls development and self-renewal in multiple tissues, including the intestine, and the tumor is interpreted as hijacking these developmental signals and exploiting them for proliferation. (Supplementary Table S9: IPA Tox Function analysis)

### **Supplementary Results 2. EnrichR GO analysis (tissue 1: amino acid transport, DNA replication, and apico-basal polarity)**

*Corresponds to Section 3.3.1 of the main text and Supplementary Note Section 2.5.*

In the GO Biological Process (BP) analysis, six significantly enriched terms satisfying adj. P < 0.05 were identified. L-leucine Transport (adj. P = 0.031), Branched-chain Amino Acid Transport (adj. P = 0.031), DNA Metabolic Process (adj. P = 0.031), Double-strand Break Repair via Break-Induced Replication (adj. P = 0.034), L-alpha-amino Acid Transmembrane Transport (adj. P = 0.050), and Epithelial Fluid Transport (adj. P = 0.050) occupied the top ranks. The enrichment of amino acid transport processes (L-leucine Transport and Branched-chain Amino Acid Transport), together with the amino acid transporter activities in the GO MF analysis, reflects the essential function of the intestinal absorptive epithelium. The enrichment of DNA Metabolic Process and Double-strand Break Repair via Break-Induced Replication indicates heightened DNA replication and repair activity accompanying a high proliferation rate. The enrichment of Epithelial Fluid Transport indicates a function in maintaining intestinal fluid homeostasis.

In the GO Cellular Component (CC) analysis, five significantly enriched terms satisfying adj. P < 0.05 were identified. Basolateral Plasma Membrane (adj. P = 0.013), External Side of Apical Plasma Membrane (adj. P = 0.013), Extracellular Exosome (adj. P = 0.019), Extracellular Vesicle (adj. P = 0.019), and CMG Complex (adj. P = 0.030) occupied the top ranks. The simultaneous enrichment of Basolateral Plasma Membrane and External Side of Apical Plasma Membrane indicates that the apico-basal polarity characteristic of intestinal absorptive epithelial cells — the clear separation of the luminal side (apical membrane) from the basement-membrane side (basolateral membrane) — is maintained in tissue 1, supporting the retention of a functional intestinal epithelial identity. The enrichment of CMG Complex (the DNA replication-initiating helicase complex composed of Cdc45-MCM-GINS [152]) supports the high proliferative capacity of tissue 1 from the standpoint of the DNA replication machinery.

In the GO Molecular Function (MF) analysis, four significantly enriched terms satisfying adj. P < 0.05 were identified. Branched-chain Amino Acid Transmembrane Transporter Activity (adj. P = 0.0063), Nucleotide Transmembrane Transporter Activity (adj. P = 0.0063), Single-stranded DNA Helicase Activity (adj. P = 0.0086), and L-leucine Transmembrane Transporter Activity (adj. P = 0.016) occupied the top ranks. The enrichment of transporter activities for branched-chain amino acids (leucine, isoleucine, and valine) corresponds directly to actual transporter genes such as SLC7A5/LAT1, SLC7A8, and SLC43A1, which are included among the tissue-1-side DEGs, and reflects active amino acid uptake in intestinal absorptive cells. The enrichment of Single-stranded DNA Helicase Activity indicates the functional activity of DNA replication helicases such as MCM4, MCM5, and MCM6, which are included among the tissue-1-side high-expression genes, and is consistent with high proliferative activity. (Supplementary Table S31: EnrichR GO analysis; tissue 1)

### **Supplementary Results 3. IPA Tox Function analysis (tissue 2: necrotic cell breakdown suggested by elevated ALP, with AST, ALT, and LDH in the same direction below threshold)**

*Corresponds to Section 3.3.1 of the main text and Supplementary Note Section 3.4.*

In the IPA Tox Function analysis, the following clinical-chemistry signals accumulated as the toxicity profile of tissue 2. Increased Levels of Alkaline Phosphatase (ALP) was detected independently multiple times (the top signal), and Increased Levels of AST and ALT (4 entries each), Increased release of LDH, Necrosis of liver, Steatohepatitis, Viral hepatitis, Liver metastasis, and Inflammation of liver also accumulated. Of these, only the ALP signals meet the adoption criteria (sign consistency ≥ 75% and |median z| ≥ 2; Increased Levels of ALP, median z = −2.12, sign consistency 94%); AST (−1.44), ALT (−1.45), and LDH (−1.50) are consistent in direction but below threshold in effect size, and the necrosis- and hepatitis-related signals are also below threshold (Supplementary Note Section 3.4; Table 5 of the main text). These are predictions by IPA from expression data, not measurements, and, like all Tox Function findings, they serve only as corroborating evidence. Taken together, the profile is consistent with, but does not demonstrate, leakage of cytoplasmic enzymes from necrotic tumor cells, that is, necrotic cell breakdown accompanying the antagonism between p53-induced apoptotic signaling and MDM2-dependent survival signaling. The accumulation of liver-related toxicity signals (all below threshold) is interpreted cautiously as a possible reflection of damage to the metastatic site; because organoids lack stroma and immune cells, it is not interpreted as damage to the tumor microenvironment. (Supplementary Table S9: IPA Tox Function analysis)

### **Supplementary Results 4. EnrichR GO analysis (tissue 2: extracellular vesicles, secretory granule structures, and endopeptidase inhibition)**

*Corresponds to Section 3.3.1 of the main text and Supplementary Note Section 3.5.*

In the GO Biological Process (BP) analysis, 20 significantly enriched terms were identified (adj. P < 0.05). The top term was Regulation of Epidermal Cell Differentiation (adj. P = 0.0003; Combined Score=399.1; BMP4, SULT2B1, ERRFI1, KLF7, SFN, ABCA12, ZFP36L1), suggesting that tissue 2 expresses genes associated with the epidermal cell differentiation program. This may reflect a "skin-type metaplasia-like" program in addition to gastric metaplasia and is consistent with the composite plasticity of tissue 2 inferred in this study (not tested at the single-cell level). Proteolysis (adj. P = 0.022; MMP7, ERAP2, MMP1, TMPRSS4, PCSK7, KLK6, ADAM19, CAPN9, CTSS, and others) indicates enrichment of proteases involved in invasion and ECM remodeling. Maintenance of Gastrointestinal Epithelium (adj. P = 0.025; VSIG1, TFF3, TFF1, TLR4, MUC6) indicates that the gastrointestinal epithelium maintenance program in tissue 2 operates through gastric metaplasia markers (TFF1, TFF3, MUC6). Skin Development (adj. P = 0.029; ERRFI1, SCEL, ANXA1, ITGB4, ADAM9, TXNIP, ABCA12, TGM3), Phospholipid Efflux (adj. P = 0.029; ABCA1, APOA2, ABCA12, ABCG1), Negative Regulation of Complement Activation (adj. P = 0.032; SERPING1, C4BPA, CLU, CD55), Sulfation (adj. P = 0.036; SULT1C3, SULT2B1, SULT1B1, SULT1C2), Cellular Response to Hypoxia (adj. P = 0.038; FABP1, BNIP3L, EGLN3, ACAA2, BNIP3, MDM2, NDRG1, ZFP36L1), Intracellular Sterol Transport (adj. P = 0.038), and Regulation of Cell Migration (adj. P = 0.032), among others, were also significantly enriched. The direct inclusion of MDM2 in the hypoxia-response gene set—given that HIF-1α is known to bind MDM2 directly and modulate p53 function [153]—is consistent with a possible axis of hypoxia → HIF1α → transcriptional induction of MDM2 P2 (not tested in this study). Such p53-independent regulation of P2 is not limited to the hypoxia response and has also been reported in adult astrocytoma [90], indicating that the P2 promoter can be activated in a context-dependent manner through multiple pathways.

In the GO Cellular Component (CC) analysis, 14 significantly enriched terms were identified (adj. P < 0.05). The highest significance by far was shown by Extracellular Vesicle (adj. P ≈ 0; CS=98.4) and Extracellular Exosome (adj. P ≈ 0; CS=80.9), suggesting increased production of tumor-derived extracellular vesicles in tissue 2 [154] (vesicle production was not measured). These were followed by Golgi Lumen (adj. P = 0.0005; 11 genes: ERO1A, VCAN, DEFA6, MUC17, DEFA5, DEFB1, MUC5B, HSPG2, MUC20, MUC5AC, MUC6), Tertiary Granule Lumen (adj. P = 0.0007; 8 genes: TCN1, TIMP2, CTSH, GGH, QSOX1, YPEL5, B2M, CTSS), Platelet Alpha Granule (adj. P = 0.003), Vesicle (adj. P = 0.003), Intracellular Organelle Lumen (adj. P = 0.003), Tertiary Granule (adj. P = 0.003), Endoplasmic Reticulum Lumen (adj. P = 0.004), Secretory Granule Lumen (adj. P = 0.005), Platelet Alpha Granule Lumen (adj. P = 0.008), Apical Dendrite (adj. P = 0.016), Extracellular Membrane-Bounded Organelle (adj. P = 0.021), and Specific Granule Lumen (adj. P = 0.023). The genes enriched in Golgi Lumen directly include gastric-type and Paneth-cell-type markers such as DEFA5, DEFA6, MUC5B, MUC5AC, MUC17, and MUC6, consistent with secretion of these proteins via the Golgi apparatus. The enrichment of Tertiary Granule Lumen (neutrophil tertiary granules) reflects secretory and granule-associated genes expressed by the epithelial cells themselves (such as TCN1, CTSH, and B2M) that overlap with the neutrophil-granule annotation, and the Neutrophil degranulation identified in the IPA analysis likely has the same origin (because organoids lack immune cells, neither indicates neutrophil degranulation or inflammation of the tumor microenvironment). This suggests a hybrid, metaplasia-like expression pattern with features of multiple organs and cell types (stomach, neutrophils, platelets, and pancreatic acinar cells), inferred from gene-set enrichment (the neutrophil and platelet features reflect overlap of epithelial secretory genes with these annotations).

In the GO Molecular Function (MF) analysis, 11 significantly enriched terms were identified (adj. P < 0.05). The top term was Endopeptidase Inhibitor Activity (adj. P = 2.0×10⁻⁵; CS=114.0; 15 factors: SPINK1, SERPINA1, ITIH2, SERPINE2, SERPINF1, TFPI2, CST7, SORL1, SERPINB5, CST4, CST1, SLPI, TIMP2, SERPING1, SERPINI1), indicating that serine protease inhibitors centered on the serpin family are highly enriched in tissue 2. Cholesterol Binding (adj. P = 0.0004; SULT2B1, ABCA1, GRAMD1B, NPC1, PTCH1, APOA2, OSBPL1A, ABCG1, PROM2) and Sterol Binding (adj. P = 0.0015) followed, and molecular functions related to cholesterol metabolism and lipid homeostasis were highly enriched. The enrichment of Death Receptor Activity (adj. P = 0.009; TNFRSF19, TNFRSF10B, FAS, EDA2R) indicates high activity of apoptotic signaling through TNF superfamily receptors, consistent with p53-dependent cell death signaling in tissue 2. Serine-type Endopeptidase Inhibitor Activity (adj. P = 0.005), Endopeptidase Activity (adj. P = 0.010), Serine-type Endopeptidase Activity (adj. P = 0.011), Aryl Sulfotransferase Activity (adj. P = 0.032; SULT1C3, SULT1B1, SULT1C2), and Tumor Necrosis Factor Receptor Activity (adj. P = 0.037; TNFRSF19, FAS, EDA2R) were also significantly enriched. (Supplementary Table S31: EnrichR GO analysis, tissue 2)

### **Supplementary Results 5. IPA Regulator Effects analysis (details of all causal cascades)**

*Corresponds to Section 3.3.3 of the main text and Supplementary Note Section 4.1.*

Because of computational constraints, this Regulator Effects analysis is based on the integration of the first 8 analyses (analyses 49, 50, 51, 52, 53, 54, 59, and 63) of the 33 analyses integrated by the other IPA modules (see the Methods of the main text). The highest-scoring causal cascade (Rank1, Consistency Score 25.93, analysis 50) was: BMP4/EGR1/HIF1A/NF-κB/TGFβ3/WNT1-3A (23 upstream regulators) → TP53/VEGFA/TGFBR1/NOTCH1/TNFRSF11B/TNFRSF19 (11 target molecules) → increased ALP. The inclusion of EGR1 in this cascade is noteworthy. EGR1 is a transcription factor that can regulate MDM2 expression in a context-dependent manner via the EGR1-binding sequence in the MDM2 promoter and can act in either direction, activation or repression; for example, it has been reported to repress MDM2 transcription in head and neck squamous cell carcinoma (demonstrated by ChIP and promoter assays) [100] (the P2 promoter itself can be activated through multiple transcription factor response elements [9]; whether the effect is direct or indirect remains unresolved; see Supplementary Note Section 3.2 for details). Therefore, this cascade suggests that P2 usage, a readout of p53 activity, and the increase in ALP co-vary in these specimens, possibly under shared upstream regulators that include EGR1; this is a computational association, and a causal relationship between EGR1 and P2 usage was not tested. In the IPA Regulator Effects Rank2 cascade (Score 25.02), six mitotic regulators—ANLN, the AURK family, PRDM4, RASSF6, TERF2, and TOPBP1—were identified as functioning as upstream regulators of 11 molecules: MDM2, TP53, CDKN1A, BAX, CASP7, CDK4, CDKN1B, GDF15, PTEN, and TNFRSF10A/B. This is consistent with a bidirectional regulatory relationship in which MDM2 is an upstream regulator of p53 and, at the same time, a downstream target of mitotic checkpoint factors (a knowledge-base prediction). The IPA Regulator Effects Rank10 cascade (Score 17.64) indicated a high-leverage group of proliferation regulators (6 → 63 molecules) in which CKAP2L, Eldr, LIN9, TOR1AIP1, OGDH, and USP47 (6 factors) collectively control 63 molecules, including FOXM1, PLK1, MYBL2, and AURKB. Furthermore, a cross-sectional analysis of all 2,442 entries in Supplementary Table S32 confirmed that 11 factors—ANLN, the AURK family, CKAP2L, Eldr, LIN9, TOR1AIP1, RASSF6, NEDD8, PPM1A, TP53COR1, and POLR2M—appeared repeatedly as upstream regulators in multiple entries. The NEDD8 conjugation (NEDDylation) pathway is essential for the activation of Cullin-RING-type E3 ligase complexes; MDM2 has also been reported to act as a NEDD8 E3 ligase for p53, and neddylation of p53 inhibits its transcriptional activity without markedly affecting its stability [155]. PPM1A regulates inflammatory stress kinases as a phosphatase of p38 MAPK and JNK. These 11 factors appeared consistently as upstream regulators in multiple IPA analyses (which share specimens and are not independent) (see Supplementary Note Section 4.1 for details). (Supplementary Table S32: IPA Regulator Effects analysis)

### **Supplementary Results 6. Detailed comparison with prior studies (van de Wetering 2015, Zhao 2021, Lukonin 2020, Okamoto 2022, and others)**

*Corresponds to Section 4.4 of the main text.*

Here we compare the findings of this study with existing CRC organoid studies.

**Comparison with morphological classification frameworks. van de Wetering et al. (2015) [117] established a living organoid biobank from 20 consecutive CRC patients (22 tumor organoid cultures) and were the first to show systematically that tumor organoids present with a range of patient-specific morphologies, from thin-walled cystic structures to compact organoids devoid of a lumen. By H&E staining, they also confirmed that this "cystic versus solid" morphological organization is largely preserved from the primary tumor to the organoid. Fujii et al. (2016) [118] constructed a library of 55 CRC tumor organoid lines and showed that each line could be classified on the basis of its gene expression signature and that the histological grade and differentiation capacity of the original tumor were recapitulated in xenografts as well. Type1 of this study (compact glandular type) corresponds broadly to the "solid" category of van de Wetering 2015, and the non-Type1 classes (Type5 + Type0; the cystic–mucinous round morphology) to the "cystic" category; the novelty of this study, absent from the prior work, is that we further linked this morphological axis to TP53 status and to a transcript-level readout of p53 activity, the MDM2 P1/P2 isoforms (Type1-dominant specimens were mostly P1-dominant and non-Type1-dominant specimens mostly P2-dominant, and morphology followed TP53 status rather than promoter usage; quantified as morphology-class fractions, Sections 3.1.2 and 3.1.4 of the main text).**

**Comparison with AI-based morphological typing. Betge et al. (2022) [119] analyzed more than 5 million CRC organoid images by high-throughput image profiling and identified two principal axes of morphological variation: "size" (correlated with IGF1R signaling) and "cystic versus solid architecture" (correlated with the LGR5-positive stem cell state). This "solid morphology ↔ LGR5+ stemness" correspondence agrees with our finding that the intestinal stem cell markers OLFM4 and LGR5 [91,92] are highly expressed in the tissue-1 organoids of this study. In contrast, the P2-dominant, mostly TP53 wild-type group of this study is not merely "cystic" but exhibits a qualitatively distinct cell-biological state, namely transdifferentiation toward the gastric type (TFF1, CTSE, REN) and the Paneth cell type (DEFA5, DEFA6), and thus has a complex morphological phenotype that goes beyond the dichotomy of Betge 2022. Okamoto et al. (2022) [12] established a 6-morphology typing of CRC organoids based on visual inspection and AI (machine learning) and demonstrated inter-patient heterogeneity, but the quantitative correspondence with molecular subtypes remained unanalyzed. The VGG16-based classification of this study (test accuracy 98.5% [64/65; Wilson 95% CI 91.8–99.7%] [27]; Section 3.1.1 of the main text) is consistent with that direction while adding new dimensions: integration with TP53 status and with MDM2 isoform information as a readout of p53 activity. Of particular note, the patient cohort analyzed by Okamoto et al. (2022) [12] is described with patient IDs such as HCT38, HCT41, HCT27, HCT67, HCT31, and HCT71, which are identical to the patient IDs of this study. That is, the two studies performed independent analyses of largely overlapping patient groups derived from the same patient-derived organoid biobank (the HCT series) and provide mutually complementary findings. Furthermore, Huang et al. (2024) [120] identified cystic and solid morphological subtypes of CRC organoids by image-based profiling of bright-field images, related them to viability and apoptosis by deep learning, and proposed that the cystic subtype, a relapse phenotype accompanied by an intestinal stem cell signature, may serve as a diagnostic and prognostic biomarker. However, this "cystic ↔ intestinal stemness" correspondence is in the opposite direction to the "solid (Type1) ↔ LGR5/OLFM4-positive stemness" shown by Betge 2022 [119] and by this study; this likely reflects differences among studies in the operational definitions of cystic/solid and in the composition of the intestinal stem cell signature, or possibly a non-monotonic relationship between lumen formation and stemness (exploratory interpretation). Because the morphological axis of this study is linked to TP53 status and to a transcript-level readout of p53 activity, the MDM2 P1/P2 isoforms, it has the advantage of compensating, with molecular axes, for the directional uncertainty that accompanies inferring stemness from morphology alone.**

Comparison with phenotypic analyses by organoid morphology × deep learning (across cancer types). Attempts to classify organoid morphology by deep learning and to link it to biologically and therapeutically meaningful states have also been reported outside colorectal cancer. Zhao et al. (2021) [10] found that organoids derived from claudin-low mammary tumors (mesenchymal triple-negative breast cancer) display a characteristic "spiky" structure in three-dimensional culture and change to a smooth, round shape upon miR-200-induced mesenchymal–epithelial transition (MET); they built a morphological screening method based on a deep neural network and nearest-neighbor classification and identified drugs that reverse EMT (class I HDAC inhibitors and bromodomain inhibitors). This is a representative example, in breast cancer, showing that organoid morphology can serve as a sensitive readout of a molecular state (EMT) and that deep learning can quantify it. This study extends this framework of "linking morphology ↔ molecular state by deep learning" to colorectal cancer, and it is novel relative to Zhao 2021 in that (1) it linked morphology to TP53 status and to a transcript-level readout of p53 activity, MDM2 P1/P2 promoter usage; (2) it captured the morphology ↔ isoform correspondence intrinsic to patient-derived organoids (Section 3.1.2 of the main text) rather than drug-induced morphological change; and (3) it presents the result not merely as a single drug-screening platform but as a classifier whose derived signatures were examined in the large independent cohorts TCGA and GSE39582 (the image-based morphological correspondence was examined outside the discovery cohort only through a morphology gene program in an independent organoid biobank, where it aligned with P2 across samples but not within patients, and the across-sample correspondence was largely explained by TP53 status; main text Section 3.1.5). The two studies support, from different cancer types and different molecular axes, the common view that organoid morphology may serve as a surrogate indicator of molecular subtype and therapeutic target. In this study as well, in the metastatic specimen HCT38-3LM, in which morphology and the MDM2 isoform diverged, we found activation of a cancer-cell-intrinsic partial EMT program (CDH2, VIM, FN1, and others) (Section 3.1.2 of the main text); this provides, at the single-specimen level in colorectal cancer, an observation consistent with Zhao 2021 that EMT can dissociate morphology from the molecular (promoter) state. Quantitatively, the EMT score of HCT38-3LM (the mean z-score of EMT/mesenchymal genes minus the mean z-score of epithelial markers: 4.94) was the highest among all 63 specimens (Supplementary Table S1), and in a patient-paired comparison of EMT scores computed from the expression matrix, metastatic specimens also showed significantly higher EMT scores than primary tumors (metastasis median +0.130 versus primary median −0.590 across all 63 specimens, per-specimen Wilcoxon rank-sum test P = 0.014; patient-level paired Wilcoxon signed-rank test P = 0.004 [17 patients]; by contrast, the difference between P2-dominant and P1-dominant specimens was not significant, P = 0.079; Section 3.1.2 of the main text). This agrees in direction with the observation in Zhao 2021 that miR-200-induced mesenchymal–epithelial transition (MET) produced morphological change, and raises the possibility that a comparable mechanism contributes to morphological change in the intrinsic EMT of colorectal cancer as well. Furthermore, in the DEG data of this study (probe level, 33 comparisons), among the miR-200 family, MIR200C, MIR200B, and MIR141 showed a consistent tendency to be higher in the P2-dominant group (mean linear FC = −1.40/−0.85/−0.77; each in the same direction in 25 of 33 comparisons). Under the convention of this study, a negative value indicates that expression is higher in the P2-dominant group. Because miR-200 is a microRNA that normally targets and represses ZEB1/ZEB2 to maintain the epithelial state (EMT suppression), the relatively high miR-200 in the P2-dominant group appears paradoxical at first sight. However, this is consistent with the interpretation that the mesenchymal tendency of the P2-dominant group does not proceed to complete EMT but remains in a partial EMT state because miR-200 partially represses ZEB1/2. This also agrees in direction with our finding of "partial EMT" in HCT38-3LM in this study and with the observation in Zhao 2021 that morphology changed upon miR-200-induced mesenchymal–epithelial transition (MET). However, this analysis is an exploratory finding at the probe level (MIR200A showed almost no difference), and a causal interpretation would require functional validation at the miRNA level. When we further examined, in this study, the expression dynamics of the EMT transcription factors ZEB1 and ZEB2 themselves, the targets of miR-200, both changed consistently toward higher expression in the P2-dominant group (ZEB1: same direction in 29 of 33 comparisons [88%], mean linear FC −0.90 [median linear FC −1.13]; ZEB2: 28/33 [85%], mean linear FC −0.99 [median linear FC −1.31]; under the convention of this study, a negative value indicates that expression is higher in the P2-dominant group; the values are from Supplementary Table S15, with the mean linear FC, the same measure as in Section 3.1.2 of the main text, shown as the primary value). However, their effect sizes were modest, approximately 1.0-fold by mean linear FC and approximately 1.1–1.3-fold by median linear FC, and did not reach the 2-fold threshold (|log2FC|≥1) of the DEG criteria (Supplementary Table S15). That ZEB1/2 are not completely repressed but are maintained at low levels even while the miR-200 family is elevated corroborates, with the data of this study itself, the above interpretation that partial repression of ZEB1/2 keeps the cells in a partial EMT state without transition to complete EMT. We also examined the direction of expression of other EMT-inducing transcription factors. SNAI2/SLUG, like ZEB1/2, was consistently higher in the P2-dominant group (32/33 [97%], mean linear FC −1.15), with a modest effect size of approximately 1.1-fold that did not reach the 2-fold threshold of the DEG criteria. In contrast, SNAI1 was higher on the tissue-1 side (27/33 [82%] in the tissue-1 direction, mean linear FC +1.88, approximately 1.9-fold), so that the direction of expression diverged among the EMT-inducing transcription factors (SNAI3 showed no between-group difference, 17/33). The higher SNAI1 on the tissue-1 side, where p53 is inactive (TP53 mutation or wild-type p53 inactivity), is not explained by the p53–miR-200 axis: wild-type p53 represses EMT through miR-200 family microRNAs that target ZEB1/2 ([113,114]), and p53 inhibition did not induce SNAI1 or SNAI2 in the study that established this axis ([113]); the cause of the higher SNAI1 remains undetermined. Meanwhile, that SNAI2 and ZEB1/2 are modestly higher in the P2-dominant group (sub-threshold) is consistent with the above interpretation that the P2-dominant group does not proceed to complete EMT but remains in partial EMT (Supplementary Table S15). Furthermore, this principle is not confined to cancer. Lukonin et al. (2020) [11] analyzed the regeneration process of normal mouse intestinal organoids by image-based high-content profiling, extracted multivariate morphological features from hundreds of thousands of organoids to quantify a phenotypic landscape (15 phenotypes), inferred genetic interactions from those phenotypic fingerprints, and showed in vivo that retinoic acid metabolism and RXR antagonism promote regeneration. That is, the principle that quantification of organoid morphology serves as a readout of molecular and genetic states can be generalized from the regeneration of normal tissue to various cancers. This study extends this lineage to colorectal cancer, and its originality lies in linking morphology to TP53 status and to a transcript-level readout of p53 activity (MDM2 P1/P2 promoter usage), and to therapeutic stratification, and in examining the derived signatures in large independent cohorts. A comparison between representative studies using organoid morphology × AI/image analysis and this study is summarized in a separate comparison table (comparison of organoid morphology × AI studies). (Supplementary Table S33)

**Novelty of MDM2 promoter usage as a readout of p53 activity. The dual-promoter structure (P1/P2) of MDM2 has long been known [6,59], but to the best of our knowledge no study has systematically analyzed P1/P2 promoter choice in CRC patient-derived organoids or used MDM2 promoter usage as a locus-level readout of p53 activity in CRC. Bond et al. (2004) [7] showed that SNP309 within P2 accelerates tumor formation, and Wade et al. (2013) [4] comprehensively reviewed the regulation of p53 by MDM2/MDMX, but the former focused on the effects of a specific genetic polymorphism, and neither examined P1/P2 usage as a readout of p53 activity in CRC. In addition, mismatch-repair-deficient (MSI) tumors carry hypermutated genomes [104], and in the CMS classification TP53 mutations are not enriched in the MSI-rich CMS1 subtype [2]. The TP53 analysis of this study (TP53 mutations are concentrated in the P1-dominant group; at the patient level [majority isoform] all three P2 patients were TP53 wild-type, whereas at the specimen level the P2 specimens HCT41-1T and HCT38-3LM carried TP53 mutations yet were p53-active, while HCT67-4LMR carried none) is an observation of this large-scale genome-level finding in organoids, a functional cellular system, in a form consistent with a transcript-level readout, the MDM2 isoforms (TP53 mutations were found in 15 of the 17 P1-dominant patients with determined status, two being wild type and one undetermined [HCT83-4LM assigned to HCT83; Section 3.1.3 of the main text]; p53 was inactive in both specimens of the wild-type P1-dominant patient HCT59 [Section 3.2.1 of the main text], morphology followed TP53 status rather than promoter usage [Section 3.1.4 of the main text], and the resemblance of P2 dominance to the MSI-like program is at the signature level only; P2_index did not differ among the CIN, GS and MSI subtypes and was not associated with MSI status [Sections 3.1.4, 3.2.2 and 4.6 of the main text]).**

**Relation to the stromal subtype. As Calon et al. showed in Nature Genetics (2015) [156], a TGFβ-driven stromal gene expression signature defines the poor-prognosis subtype of CRC (the CMS4 mesenchymal type). The activation of ECM Organization, TGFβ/SMAD3, EMT, and invasion-related pathways observed in tissue 2 (P2-dominant) also overlaps with CMS4-like characteristics, and one feature of this study is that this phenotype is a composite one that does not fit within a single CMS subtype. We propose, as a hypothesis not tested here, that MDM2 P2 induction may accompany partial EMT; the P2–P1 difference in EMT score was not significant (P = 0.079; Section 3.1.2 of the main text).**

**Comparison with the metabolic phenotype. In this study, we found that Oxidative Phosphorylation and the TCA Cycle are simultaneously activated in tissue 1 (which corresponds to CMS2 by the criterion of the differentiation core excluding the proliferation module; the highest overall score is CMS1, but the difference from CMS2 is not significant [Section 3.3.2 of the main text and Supplementary Note Section 5.2]). This can be interpreted in terms of meeting the energy demand that supports a high proliferation rate, and is consistent with an autonomous proliferation program driven by the MYC/FOXM1/E2F axis. Detailed comparison with Okamoto et al. 2022: complementarity of independent analyses using a shared cohort. The comparison with Okamoto et al. (2022) [12] goes beyond a mere comparison of methodology; its significance lies in integrating the results obtained by analyzing organoids derived from the same patients along different analytical axes (morphological typing versus TP53 status and MDM2 isoform analysis). The principal correspondences are summarized below. First, regarding the correspondence of morphological types, Type 1/2 of Okamoto 2022 (the Type A group: budding structures, narrow lumina, and low expression of cell-adhesion genes) is expected to correspond to the Type1-dominant morphology of this study (compact glandular type; the Type1 fraction was higher in TP53-mutant organoids, Section 3.1.4 of the main text), and Type 3/4/5 of Okamoto 2022 (the Type B group: wide lumina, spherical structures, and high expression of cell-adhesion genes) is expected to correspond to the non-Type1-dominant round morphology of this study (Type5 + Type0) (the two typings were made independently and were not matched image by image, and neither typing is equivalent to MDM2 P1/P2 usage). The finding that Okamoto 2022 demonstrated with correlation coefficients, namely that “Type 1 is negatively correlated with Types 3, 4 and 5 (correlation coefficients −0.17, −0.17 and −0.11), so that Types 1/2 and Types 3–5 fall into separate groups,” is consistent with the quantitative correspondence in this study that “P1-dominant and P2-dominant specimens separate as Type1-dominant versus non-Type1-dominant morphology (median Type1 fraction 0.826 versus 0.444; Section 3.1.2 of the main text).” Second, regarding the expression patterns of LGR5, OLFM4, and ribosome biogenesis genes, Okamoto 2022 showed that Type 1 PDOs markedly overexpress ribosome biogenesis-related genes (NES = 2.53) and also highly express intestinal stem cell markers such as LGR5, CD44, and OLFM4 (Figure 5B–D of that paper). In this study as well, OLFM4 was highly expressed in tissue 1/P1 (avg FC +1,252, 73% consistent), and the MYC, FOXM1, and E2F axis was identified in the Upstream analysis. Ribosome biogenesis is a major transcriptional target of MYC, and the finding of Okamoto 2022 is consistent, in an independent analysis of the same-biobank cohort, with a possible link, not tested in this study, “MYC/FOXM1/E2F axis → ribosome biogenesis → maintenance of LGR5-positive stem cells.” Third, regarding CX-5461 resistance, as its most important finding, Okamoto 2022 performed an Upstream Regulator analysis of the RNA Pol I inhibitor CX-5461 and showed that Type 1/2 (group A) is resistant to CX-5461 (Figures 6A and 6E of that paper). Furthermore, in Figure 6D of that paper, MDM2 is explicitly shown as a downstream target molecule of CX-5461. From the viewpoint of this study, one possible interpretation (not tested here; CX-5461 response was not examined in our samples, and Type 1/2 organoids were not matched to our specimens) is that the resistance reflects p53 inactivity (TP53 mutation or wild-type p53 inactivity), which in our specimens accompanied the compact glandular morphology and the MYC/FOXM1/E2F program; under p53 inactivity, MDM2 is transcribed mainly from the basal P1 promoter because p53-dependent P2 induction fails (Sections 3.1.4 and 3.2.1 of the main text). Fourth, regarding the same finding in patient HCT67, Figure 3A of Okamoto 2022 showed that patient HCT67 has organoids of both Type A and Type B. In this study as well, a promoter switch was observed in the same patient (HCT67) between the primary tumor HCT67-1T (P1-dominant, Type1-dominant morphology) and the metastatic specimen HCT67-4LMR (P2-dominant, non-Type1 [cystic–mucinous] morphology, Type1 fraction 0.12; the other metastatic specimen, HCT67-3LM, remained P1-dominant; TP53 R175H was detected in the primary tumor and the liver metastasis but not in the recurrent liver metastasis HCT67-4LMR, Figure 9D of the main text). In the overlapping patient group from the same biobank, the coexistence of Type A and Type B within the same patient described by Okamoto is thus consistent with the within-patient coexistence of morphologies, of P1- and P2-dominant specimens and of different TP53 status observed in this study. In this way, Okamoto 2022 describes morphology, and this study relates morphology to TP53 status and to a promoter-level readout of p53 activity (MDM2 P1/P2 usage; morphology correlates only weakly with the MDM2 axis [ρ = 0.33] and follows TP53 status beyond expression-inferred 20q gain and aneuploidy; Section 3.1.4 of the main text), and the two studies form a mutually complementary body of evidence through cohorts derived from the same biobank.**

### **Supplementary Results 7. Robustness of the differentially expressed gene (DEG) definition (threshold sensitivity analysis and permutation test)**

The DEG definition of this study differs from conventional definitions in that its primary criterion is the consistency of the direction of change across the 33 pairwise comparisons rather than statistical significance (P value or FDR). We quantitatively verified the validity of this definition from two perspectives: a threshold sensitivity analysis and a sign-randomization permutation test (the analysis code is given in Supplementary Methods, and the results in Supplementary Table S5 and Supplementary Figures S1 and S2). The input was the signed linear fold change of all probes (33 comparisons; Supplementary Table S16 [the gene-symbol-restored version]); for genes with multiple probes, the probes were grouped by exact match of the gene symbol and averaged for each comparison to give gene-level values (composite-symbol rows were not allocated to any token; fully reproducible with seed=42).

**Threshold sensitivity analysis.** Centering on the adopted criteria of a consistency rate of 70% and |log2 fold change| ≥ 1 (2-fold), we recomputed the number of DEGs on a grid of consistency rates of 60–80% and fold changes of 1.5-, 2-, and 3-fold. The number of DEGs (total / higher in tissue 1 / higher in tissue 2) changed smoothly with the thresholds, from 1,997 (1,063/934) at 60% and 2-fold to 1,890 (1,013/877) at 70% and 2-fold and 1,658 (928/730) at 80% and 2-fold, and no abrupt discontinuity was observed. The tissue 1:tissue 2 ratio remained within the range 53:47–56:44 (1,063/934 = 53:47 at 60% and 2-fold, 1,013/877 = 54:46 at 70% and 2-fold, and 928/730 = 56:44 at 80% and 2-fold), and the representative markers (OLFM4, SLC26A3, CTSE, REN, MMP7, MDM2, etc.) remained DEGs under every setting. This indicates that the conclusions are robust to the choice of the adopted thresholds.

**Permutation test.** For each gene, the signs of the fold changes across the comparisons were randomized according to a Rademacher distribution while their absolute values were retained, and the DEG criteria (consistency rate 70%, 2-fold) were recomputed for all genes; this operation was repeated 1,000 times. As a result, the null distribution of the number of genes satisfying both criteria by chance had a mean of only 11.3 (SD 3.3, maximum 23), whereas the observed value was 1,890 genes; the empirical FDR was 0.60% (11.3/1,890), with permutation P < 1×10⁻³ and z = 567. That is, of the 26,278 genes, only about 11 satisfy both criteria by chance, which quantitatively supports the conclusion that the observed directional agreement is unlikely to arise by chance even under a consistency-based definition that does not use the P value as its primary criterion. Note that the number of DEGs expected by chance under the consistency criterion alone (without a fold change threshold) is analytically 355.6 (1.35%), confirming that the concomitant use of the fold change criterion suppresses false positives substantially further.

**Restoration of gene symbols.** In the Gene symbol column of Supplementary Table S16 (probe-level linear fold change), the gene symbols of 27 genes and 31 probe sets had been converted into date values by the spreadsheet's automatic date conversion, and we therefore restored them using the intact Description column as the primary source (MARCH1–11, MARC1–2, SEPT1–14, SEP15, and DEC1; these are the symbols current at the time the array was produced, and the approved symbols after the 2020 HGNC renaming correspond to MARCHF1–11, MTARC1–2, SEPTIN1–14, SELENOF, and BHLHE40, respectively). Restoration was performed only for the 31 rows for which all three routes agreed: (i) deriving the symbol from the Description column according to the HGNC Approved name rules, (ii) confirming that the month and day of the date are consistent with the symbol, and (iii) confirming that the symbol actually exists in the R-derived expression matrix that never passed through a spreadsheet (exprs_matrix_symbol.csv); not a single fold change cell was altered. The restoration resolved the state in which MARCH1 and MARC1, and MARCH2 and MARC2, had each been merged into a single symbol, and the gene universe increased from 26,276 to 26,278. None of the 27 restored genes satisfies the DEG criteria (maximum |mean linear fold change| = 1.72), and the 1,890 genes of Supplementary Table S4 and the DEG calls are completely unchanged. The permutation figures given above are the results of re-running the analysis with the restored version as input, and in the threshold sensitivity grid only 5 cells shift by 1 each (3,310→3,311 at a consistency rate of 60% and 1.5-fold). The 31 restored rows can be identified in Supplementary Table S16 by the Gene symbol column (the restored symbols) and the Description column (the primary source of the restoration), as is also noted in that table; the working record of the three-route judgments is available from the corresponding author on reasonable request. In Supplementary Table S16, the Gene symbol column is stored in text format to prevent re-conversion to dates.

### **Supplementary Results 8. Metastatic trajectory analysis (PCA/UMAP)**

From the RMA-normalized, log2-transformed expression matrix of all 63 specimens, we visualized the expression landscape using the 3,000 genes with the highest variance by principal component analysis (prcomp, with each gene scaled) and by UMAP (umap package, n_neighbors=15, random_state=42) (analysis code: Code6, Supplementary Methods). For each patient, the primary tumor → metastasis pair was connected by an arrow, and the trajectories of the patients with both P1- and P2-dominant specimens (HCT38, HCT41, and HCT67) were highlighted with thick lines.

In both Supplementary Figure S3 (PCA: PC1 14.4%, PC2 9%) and Supplementary Figure S4 (UMAP), the primary-to-metastasis trajectories of the three patients with both P1- and P2-dominant specimens (HCT38, HCT41, and HCT67) were long and traversed a large part of the expression space, in contrast to the relatively short trajectories of the other patients. This indicates that the specimens deviating from the typical morphology of their group (Section 3.1.2) also shifted markedly in their expression profile as a whole, and it is consistent with a global expression difference accompanying the within-patient difference in promoter usage (exploratory; based on a small number of patients).

### **Supplementary Results 9. Concordance analysis with mechanotransduction mechanisms**

To test the expression-inferred difference in physical properties between tissue 1 (“dense solid”) and tissue 2 (“highly viscoelastic gel”) in this study (Section 4.5 of the main text and Supplementary Note Section 5.3) against the framework in which the mechanical microenvironment regulates chromatin state and transcription [106], we re-aggregated the probe-level expression of this cohort (Supplementary Table S16) and the IPA Upstream analysis (Supplementary Table S7) for the gene sets that this framework cites: (i) the nuclear lamina–heterochromatin machinery, (ii) mechanosensitive transcription factors (Table 1 of [106]), and (iii) the soft-ECM metabolic axis (Fig 6d of [106]) (Supplementary Table S29, 92 genes in total). The nuclear-lamina tethering and heterochromatin-writing machinery was coordinately activated on the tissue-1 side, and the mechanosensitive transcription factors (TGFβ-SMAD, β-catenin, NF-κB, AP-1, SRF-MRTF, and YAP/TAZ-TEAD) on the tissue-2 side, consistent with the inferred physical states. In contrast, the soft-ECM metabolic axis (NRF2 and SREBP) was not reproduced in tissue 2. This analysis is a re-aggregation of existing data; it shows concordance between the expression-inferred physical state and molecular mechanism, not mechanical causation (physical properties were not measured).

### **Supplementary Results 10. Sensitivity analysis of the IPA adoption criteria**

We evaluated how the choice of the adoption criteria for IPA findings described in Section 2 of the main text—(i) sign consistency of the Activation z-score of 75% or more across the 33 comparisons and (ii) |median z| ≥ 2—affects the conclusions by varying, on a grid, the sign-consistency threshold (70/75/80%), the upper limit of the frequency rank (20/25/30/50/100/unlimited), and the |median z| threshold (not applied / ≥ 2) (Supplementary Table S10). First, when the rank limit was set to the top 25, the number of adopted items remained unchanged at 68 regardless of whether the sign-consistency threshold was set to 70%, 75%, or 80%. That is, once a rank limit is imposed, the sign-consistency threshold does not affect the result, and the two criteria are strongly correlated. Second, when only the rank limit was relaxed, the number of adopted items increased monotonically (rank 25: 68 → rank 30: 80 → rank 50: 135 → rank 100: 251), and the added items included many that satisfied |median z| ≥ 2 and a sign consistency of 85% or more. Specifically, the hypoxia inducers deferoxamine (median z = −4.66, 94%) and dimethyl n-oxalyl-glycine (−4.86, 85%), the EZH2 inhibitor tazemetostat (−3.96, 97%), the LSD1 inhibitor SP2509 (−4.10, 100%), and the XPO1 inhibitor eltanexor (−4.59, 97%) are all added as upstream regulators in the tissue-2 direction, and Spliceosomal Cycle (+3.44, 100%), RNA Polymerase I Transcription (+3.54, 97%), and Nonhomologous End-Joining (+3.53, 97%) are added as Canonical Pathways in the tissue-1 direction. These are directionally consistent with the findings of Section 3.3.1 (hypoxia adaptation), Section 4.3 (chromatin repression machinery), and Section 3.3.1 (RNA processing/ribosome biogenesis) of the main text and are consistent with them (they come from the same IPA output and are not independent evidence). On this basis, we judged that the frequency rank, as an index of how frequently an item appears in the IPA output, introduces information of a different dimension from effect size and directional reproducibility, and we excluded it from the final adoption criteria (ranks are listed alongside, as descriptive information, in Supplementary Tables S6–S9). Note that the aggregation rule for multi-probe genes differs among analyses. In aggregating from Supplementary Table S16 to the gene level, the 1,890-gene DEG set of the main text was calculated by the rule of “averaging the probes within each comparison and then averaging across the 33 comparisons,” whereas Supplementary Table S27 was calculated, as stated in its note, by the “representative probe (maximum |linear FC|)” rule. For single-probe genes the two coincide, and the values diverge only for multi-probe genes (NKD1: −30.69/−26.17; TCF7L2: −3.04/−2.91; CD44: +1.69/+1.50; EPHB2: +2.40 [85%]/+1.43 [61%]). The description of the WNT pathway in Supplementary Note Section 5.1 follows the rule of Supplementary Table S27, and EPHB2 is not included in the 1,890-gene DEG set because it meets the DEG criteria under the representative-probe rule but not under gene-level aggregation. When criteria (i) and (ii) are adopted, 21 of the major upstream regulators discussed in the main text (the 25 regulators listed in the Main_text_regulators sheet of Supplementary Table S10) meet the criteria. Of the remaining 4, EGR1 (median z = −1.95, sign consistency 91%), CDX1 (+1.61, 93%), and E2F2 (+1.74, 90%) show high directional reproducibility but an effect size below the threshold and are explicitly labeled sub-threshold where they appear. HNF4A (−0.14, 52%) falls far below the sign-consistency criterion and its direction of activation cannot be determined; we therefore make no assignment to a tissue class on the basis of the IPA Upstream analysis (the mRNA expression of HNF4A itself, however, is robust—a mean of approximately +2.0-fold on the tissue-1 side with a sign consistency of 94%—and the description in the Supplementary Report (Additional file 1), Section R-20, is based on this expression-level finding).

### **Supplementary Results 11. Aggregation rule in the TCGA-COAD/READ analyses (from aliquot rows to the specimen level)**

*Corresponds to Section 3.3.2 of the main text.*

All TCGA-COAD/READ analyses in the main text were performed at the specimen level (sample level). The columns of the expression matrix obtained from GDC are at the aliquot level, and in some cases multiple aliquots from the same specimen are registered. If aliquot rows are used directly as the unit of analysis, the same specimen is counted more than once, and the degrees of freedom of the tests are overestimated because of pseudo-replication. We therefore used the first 15 characters of the sample barcode (TCGA-XX-XXXX-01) as the specimen key and aggregated the rows to one record per specimen, taking the arithmetic mean for continuous values and, for categorical values, adopting a single value with priority given to valid values (a value was treated as missing only when all aliquots of the same specimen were indeterminate). This aggregation rule was adopted for this study and for the companion study of the 15q11-q13 locus (T. Tsukui, R. Yao, and K. Tsuda, unpublished observations), which analyzes the same TCGA specimens. The correspondence before and after aggregation in this study is as follows.

(i) Primary tumors: 647 aliquot rows → 624 specimens, after excluding 51 normal tissues, 2 recurrent specimens, and 1 metastatic specimen from the 701 entries obtained from GDC. (ii) CMS classification (CMScaller, Entrez ID, RNAseq=TRUE, FDR<0.05): 574 of the 647 rows could be classified → 563 specimens (CMS1 96, CMS2 167, CMS3 96, CMS4 204; indeterminate 61). Among the 13 specimens with duplicate aliquots, there was no discrepancy in the CMS calls. (iii) TP53 mutation status (Masked Somatic Mutation): 578 specimens for which somatic mutation data were available (wild type 234, mutant 344). Cases for which somatic mutation data were not available were excluded from the analysis rather than being treated as wild type. (iv) Survival analysis: 591 cases with overall survival information, with duplicates removed at the patient level (533 cases for the Kaplan–Meier analysis by CMS). (v) MDM2 P2 promoter usage (P2_index, UCSC Xena TOIL): P2_index is non-missing in 380 specimens; of these, 374 specimens can be matched with TP53, 342 with CMS, and 368 with overall survival, and there are 328 complete cases for the multivariable linear model (TP53 status, CMS, tumor purity, and MDM2 copy number) (Section 3.2.2 of the main text). Because this model includes CMS as an explanatory variable, the upper bound is 342 specimens; excluding from these the 5 specimens for which somatic mutation data were not available (TCGA-A6-6140, TCGA-A6-6141, TCGA-D5-6923, TCGA-DC-6156, and TCGA-EI-6511) and the 9 specimens for which MDM2 copy number was not available (TCGA-AH-6547, TCGA-AZ-4684, TCGA-BM-6198, TCGA-CK-4952, TCGA-CM-4744, TCGA-CM-5868, TCGA-CM-6679, TCGA-DM-A285, and TCGA-F4-6855) leaves 328 cases.

(vi) Notation of P values: for the dMMR/pMMR comparison of the P2 signature in GSE39582 (dMMR 75, pMMR 444), because both groups are of size 50 or more, the normal approximation of wilcox.test was used, yielding P = 1.14×10⁻¹⁷. The minimum P value attainable by the Wilcoxon rank-sum test at this sample size is 1.1×10⁻⁴³ under the normal approximation with continuity correction, so double-precision underflow (below 5×10⁻³²⁴) cannot be reached. Section 3.3.2 of the main text reports this measured value as the full-score result; the primary score in the main text and Supplementary Figure S6C is the immune-excluded P2 score (P = 6.59×10⁻¹⁰). The reported value was computed by R49. The P-value formatting function of Code8 v11h is designed to print "< 5e-324 (underflow)" only in the case of underflow (P=0), and the measured value of this comparison does not fall into this category. For the quantities shared by the two papers, agreement was confirmed by cross-checking through the primary data, not by comparing the main texts against each other.

All values that changed as a result of the change in the unit of aggregation (Kruskal–Wallis P by functional module, medians by CMS, n for the TP53 orthogonal axis, and the number of modules concordant with ssGSEA) are reported at the specimen level throughout the main text, the figure legends, and the supplementary tables. Supplementary Table S34 is calculated at the specimen level, and the agreement between the values in the body of the table and the summary in its notes has been verified by machine. Note that the GSVA calculation of the signature scores itself was performed at the aliquot level, and the scores were then averaged to the specimen level (the expression matrix was not averaged before scoring).

### **Supplementary Results 12. Index of the Supplementary Methods**

*Corresponds to Sections 2 and 3 of the main text.*

The analysis code of this study is deposited in the Supplementary Methods as Code1–Code21 (Code10 is a vacant number) and, for the scripts whose numbering is pending, as R14, R16, R38, R39, and R42–R49 (including R47b, R47c and R47d). The analyses added for Sections 3.1.2, 3.1.4, 3.1.5 and 3.2 are deposited as scripts that keep their internal analysis numbers (R68–R177) and are summarized after the list below. Code12–Code15 correspond to the four external-validation analyses (D1–D4), numbered in the order in which they are described in Sections 3.2.2, 3.2.4 and 3.3.3 of the main text. The code numbers are identifiers and do not coincide with the order of first appearance in the current main text. The full index—inputs, outputs, dependencies, run order, and the bundled execution records of every script—is README_SupplementaryMethods in Additional file 2; the correspondence between each code and the main text is summarized as follows.

Code1: the full set of training and inference scripts for deep-learning morphological classification of organoids (VGG16 transfer learning) → Section 2 (image classification by deep learning), Section 3.1.1, Table 1 (confusion matrix), and Figure 1. Code1b (S178f_reinfer_792.py): re-classification of the same 792 images with training-matched normalization → Sections 2, 3.1.1 and 3.1.2 and Supplementary Tables S2 and S46. Code2: the script that derives the RMA-normalized expression matrix from the CEL files (oligo::read.celfiles() → oligo::rma()) → Section 2 (whole-transcriptome analysis by microarray). Code3: sample summary of the 63 specimens and calculation of the EMT score → Section 3.1.2 and Supplementary Table S1. Code4: annotation of the HTA2.0 array (the SQLite database of pd.hta.2.0 is queried directly with DBI::dbGetQuery to convert probe set IDs into gene symbols) → generation of exprs_matrix_symbol.csv → Section 2. It does not itself perform RMA normalization (RMA is performed by Code2). Code5: robustness of the DEG definition (threshold sensitivity analysis and permutation test) → Section 2, Supplementary Table S5, and Supplementary Figure S1 (Supplementary Results 7). Code6: metastatic trajectory analysis (PCA/UMAP) → Section 3.1.2, Supplementary Figure S3, and Supplementary Figure S4 (Supplementary Results 8). Code7: hierarchical clustering with the 39 genes from IPA Biomarker Detection → Section 3.3.3 and Supplementary Figure S5. Code8: external validation of the downstream signatures (GSVA/ssGSEA; CMS, MSI, TP53, and purity; division into functional submodules) → Section 3.3.2, Supplementary Figure S6A–B, and Supplementary Figure S9 (the GSE39582 values reported in Sections 2 and 3.3.2 and in Supplementary Figure S6C come from R49; see below). Code9: direct external validation of MDM2 P1/P2 (P2_index and its associations with TP53 status, CMS and the signatures) → Section 3.2.2 and Figure 5. Code10: vacant (the statistical analysis code for Section 3.2.4 is Code12; the plotting script for Supplementary Figure S10, make_FigureS10_v6.py, is included; see below). Code11: generation of the Kaplan–Meier curves for the P1/P2 scores, the KM curves by CMS, and the multivariable Cox forest plot → Section 3.3.2, Supplementary Figure S7A, Supplementary Figure S7B, Supplementary Figure S7C, and Supplementary Figure S7D. Code12: validation of sensitivity to the 4 MDM2–p53 pathway drugs in GDSC1/GDSC2 and of MDM2 dependency by DepMap CRISPR (by TP53 status and by MSI status; integrated sensitivity score) → Section 3.2.4 and Supplementary Table S38. Code13: immune deconvolution (quanTIseq and MCP-counter) and prediction of ICI response by TIDE, together with the MSI-stratified sensitivity analysis → Section 3.3.3, Supplementary Table S35, and Supplementary Table S36. Code14: comparison of P2_index and p53 target module output by functional class (WT/LOF/GOF; definition A/definition B) using the mutation annotation of the TP53 Database → Section 3.2.2 and Supplementary Table S37. Code15: projection of sensitivity to the 198 drugs in GDSC2 by oncoPredict, and its correlation with the P2/P1 scores, degree of separation, and comparison by CMS → Section 3.2.4 and Supplementary Table S39 (sheet S39). Code16: generation of the TCGA expression matrix (tcga_coadread_log2tpm.rds) and the covariate table (tcga_coadread_covariates.csv) that serve as input to Code9 → input to Section 3.2.2 of the main text. TP53 status is assigned by a three-way rule—specimens in which a somatic mutation was detected are mutant, specimens for which somatic mutation data exist but no mutation was detected are wild-type, and specimens for which somatic mutation data themselves were not available are missing—which is consistent with the declaration in Section 3.2.2 of the main text (non-detection is not treated as wild-type). Code17: calculation of tumor purity (ESTIMATE) → Section 2 and the sensitivity analysis of Section 3.3.2. Tumor purity is a variable shared with the companion study of the 15q11-q13 locus (T. Tsukui, R. Yao, and K. Tsuda, unpublished observations), and this single script was written for both reports. Code18: export of the per-specimen GSVA scores and CMS assignments → the paired comparison in Section 3.3.2 of the main text. Code19: retrieval of MDM2 gene-level copy number (ASCAT3) and decomposition of composite barcodes → Sections 2 and 3.2.2 of the main text. In addition, Code20 (R15_S363_S365_sample_level_numbers_v1.R; the partial correlations of Section 3.2.2), Code21 (analyze_s365_v1.py; the multivariable linear model and the stratification by high-level amplification of Section 3.2.2), and the scripts already deposited in the Codes/ directory with their code numbering pending—R38 and R39 (the analysis of the 65 organoid lines in Sections 3.2.4 and 4.6 of the main text), R42–R44 (the colibactin-related analyses of Section 2 of the main text, Supplementary Note Section 5.2, and Section 4.6), R45 (the BBC3 sensitivity analysis of Section 2 of the main text [GSE39582; 211692_s_at]), R46 (the sensitivity analysis of the CMS input scale of Section 2 of the main text [the double log transformation in CMScaller]), R47 and R47b (measured GDSC2 drug sensitivity and DepMap CRISPR dependency versus the P1/P2 scores, unadjusted and adjusted for cancer type and TP53 status; Section 3.2.4 and Supplementary Table S39, sheets S39b and S39c), R47c (the same analysis additionally adjusted for RAS and BRAF mutation status and restricted to RAS/RAF wild-type lines; Section 3.2.4 and sheet S39d), R47d (the drug-class comparison of the per-drug partial correlations written by R47b [two-sided Wilcoxon rank-sum test]; Section 3.2.4 and sheet S39b, notes 2 and 3), R48 (the patient-level sensitivity analysis of the morphology–isoform correspondence; Sections 2 and 3.1.2 of the main text), and R49 (the tumor-only re-analysis of GSE39582 [the 19 non-tumoral mucosa samples removed before scoring], including the BBC3 sensitivity analysis, which is the source of the GSE39582 values reported in Sections 2 and 3.3.2 of the main text and in Supplementary Figure S6C; Code8 and R45 are kept as the records of the analysis that retained these samples)—together with the records of the execution environment, are listed in the index README_SupplementaryMethods.

Scripts added for Sections 3.1.2, 3.1.4, 3.1.5 and 3.2 (internal analysis numbers R68–R177; these numbers identify runs and do not follow the order of the main text). Patient assignment and replication arrays: R71 (joint RMA of the 63 discovery arrays and the additional HTA 2.0 arrays of the same biobank) with r71_prep_v1.py and r71_analysis_v1.py (per-specimen MDM2 P2 index), r84_identity_v1.py (patient identity of the arrays registered as HCT84 → Section 2), r85_replication_v1.py and r89_public_only.py (replication of the P1/P2 separation in the arrays not used for discovery, the main values restricted to the 10 arrays identical to GSE128213 files → Sections 3.1.2 and 3.2.1 and Supplementary Table S24, sheet 4), R86 (R48 and Code6 rerun with HCT83-4LM (array file HCT84-4LM) assigned to patient HCT83 → Section 3.1.2 and Supplementary Figures S3 and S4) and R87 (check of the MSI coding of Code8 → Section 3.3.2). Section 3.2: R68 with r68_load.py, r68_p53activity_v1.py, s222_prep_v1.py and p53_organoid_checks_S229.py (organoids → Section 3.2.1 and Figure 3) and r68_screen_v1.py with its output R68_screen_iso_hits.csv (genome-wide exon-usage screen, 40 probe sets; the output is read by r129 → Section 2 [genome-wide survey; p53 activity score] and Section 3.2.5); R82 (nutlin RNA-seq, GSE86221 → Section 3.2.1 and Figure 4); R73 with r73_expr_v1.py, r73_analyze_v1.py, r73_analyze_part2.py and r73_analyze_part3.py, R90, R97 and r100.py–r100e.py (TCGA tumors → Section 3.2.3, Figure 6 and Supplementary Table S48, sheet A); R70 with r70_cellline_p53activity_v1.py and r70_sonkin_check_v1.py, R74 and R74b with r74_ccle_p2_v1.py, R76 with r76_analysis.py, and r93_ctrp.py (cell lines → Section 3.2.4, Figure 7 and Supplementary Table S48, sheets B1 and B2); r109_S240c_organoid_mdm2total_v1.py, r109b_S240c_celllines_v1.py, r109c_S240c_gdsc_v1.py and r110_S241_S48_sheetC_v1.py (whether P2 usage carries information beyond total MDM2 in tumors, organoids and cell lines → Sections 3.2.1, 3.2.3 and 3.2.4 and Supplementary Table S48, sheet C). Section 3.2.2 (Supplementary Report, Section R-12): R117_S242_recount3_MDM2_junctions_v1.R and r117d_S242_crc_junction_v1.py (MDM2 promoter usage counted from splice-junction reads in TCGA-COAD/READ → Supplementary Table S48, sheet D); r120_S243_paper1_rppa_v1.py (protein levels from RPPA → Section 3.2.3 and Supplementary Table S48, sheet E); r137_sgnex_mdm2_v1.py with bamremote.py (MDM2 junction reads counted directly in the public SG-NEx short- and long-read alignments, reading only the MDM2 region) and r142_sgnex_p2j_vs_ccle_v1.py (comparison with the DepMap P2 index → Section 3.2.4 and Supplementary Table S48, sheet G). Section 3.1.4: r77_load.py, r77_screen.py, r77_layer2_adj.py, r77_detail.py and r77_arm_enrichment_patient21_S229.py (genome-wide screen and chromosome-arm enrichment → Section 2 and Section 3.1.4, Figure 2A), r88_cyto_go_v3.py (cytoskeleton and adhesion GO terms) and r101.py–r101c.py (morphology gene score in TCGA tumors → Section 3.1.4 and Figure 2). Sections 3.1.5 and 3.2.5: R129_S247c_GSE294511_HTA2_export_v2.R with R129_S247c_run.command (download of GSE294511 and RMA of its 119 organoid and 50 tissue arrays), r129_S247c_GSE294511_replication_v1.py (replication in the independent organoid biobank → Sections 3.1.5 and 3.2.5) and r129b_S248_TableS48_sheetF_v1.py (Supplementary Table S48, sheet F) and r130_S248_morphology_only_sensitivity_v1.py (sensitivity analysis with the 124 morphology-only clusters) and r130b_S248_cluster_robust_v1.py (drug-sensitivity models with standard errors clustered by patient; patient-mean partial correlations). Figures 3, 4, 6 and 7 and Figure 2 are drawn, with every plotted value recomputed, by make_Figures10_13_v1.py and make_Figure14_v1.py, respectively (the script names do not follow the figure numbers). r93_pancan_p2.py (29 TCGA cancer types) is kept as a run record only; its results are not reported in this article. TP53 status versus promoter usage (Sections 2, 3.1.2, 3.1.3, 3.1.4, 3.1.5 and 3.2.1, Figures 1D, 2E–G, 3 and 9, and Supplementary Tables S11, S23 and S48, sheets C and H): r148_tp53_recall_v1.py with its input tables R148_TP53_calls_discovery.csv and R148_type1_fraction.csv (every TP53-genotype-dependent result computed with the final per-specimen calls), r148b_sheetC2_v1.py (sheet C with the final calls), r148c_gse294511_morph_tp53_v1.py (morphology score versus TP53 status in GSE294511), r144_organoid_arm_p53_v1.py (chromosome-arm expression scores used as 20q and aneuploidy proxies), r149_morph_tp53_cin_v1.py (morphology by TP53 status beyond 20q and aneuploidy in the discovery organoids, GSE294511 and TCGA-COAD/READ), r150_programs_tp53_vs_p2_v1.py (which expression programs follow TP53 status and which follow P2 beyond TP53 status), r151_tp53_class_morph_v1.py (TP53 mutation class, MSI and BRAF in TCGA-COAD/READ), r152_S257b_tp53_morph_cimp_v1.py with R152_TCGA_CIMP_calls.csv (adjustment for CIMP), r153_S257b_discovery_genomewide_tp53_vs_p2_v1.py (genome-wide comparison of TP53 status and promoter usage in the discovery organoids), and the table builders build_S11_S23_v2.py, build_S48_sheetH_v1.py, build_S48_sheetH_R152_v1.py and build_S48_sheetH4_R153_v1.py. Normal-tissue baseline and published classifiers: r160_S264_normal_baseline_P2_index_v1.py (P2_index of TCGA adjacent normal tissue, tumors and patient-matched pairs from the Toil recompute; Table S48 sheet I), r159_S264_published_p53_classifiers_v1.py and r159b_S264_li2024_v1.py (comparison of the p53 activity score and P2_index with the TP53-loss phenocopy score of Fito-Lopez et al. [20] and the classifier of Li et al. [21]; Table S48 sheet J), and the table builder build_S48_sheetsIJ_S265_v1.py. ENCODE PRO-cap initiation at MDM2 P1 and P2: R161_S267_public_chromatin_and_queries_v1.py with R161_S267_run.command (remote extraction of the ENCODE GRCh38 signal tracks around MDM2, with the experiment and file tables A_encode_experiments.tsv and A_encode_bigwig_files.tsv), r161_enc_lib.py and r161_features_v1.py (per-file region means; output R161_features_by_file.tsv), r161_mdm2_summary_v1.py and r161_mdm2_final_v1.py (fraction of transcription initiation at P2 per experiment, reprocessed copies of the same replicate counted once; output R161_mdm2_final_output.txt → Section 2 [cell lines] and Section 4.2), and the table builder build_S48_sheetK_and_P2_ENCODE_sheet_S267e_v1.py (Supplementary Table S48, sheet K; the same script also writes an ENCODE-chromatin sheet of a supplementary table of the companion study of the 15q11-q13 locus and stops at that step in this archive because the inputs of that sheet are not bundled). Further scripts behind Supplementary Table S48: r163_S270_emt_axis_stratified_v1.py with independent_verify_R163_S270.py (EMT score by sampling site, MDM2 isoform and TP53 status → Sections 3.1.2 and 4.2.1, Figure 9 and Supplementary Table S48, sheet L), r165_S272_definition_sensitivity_v1.py with independent_verify_R165.py and build_S48_sheetN_S272.py (sensitivity of the Section 3.2.3 analysis to the TP53-call, hypermutation and expression-source definitions → sheet N), build_S48_sheetO_S272f.py (the list of organoid morphology genes → Section 3.1.4 and sheet O), and r168_S273_colibactin_p53_v1.py with build_S48_sheetP_S273.py (colibactin mutational-signature activities in TCGA-COAD/READ by TP53 status, hypermutation and p53 activity, and in the Nunes cohort, where TP53 status is not available, against the p53-target score in all and in MSS specimens → Sections 3.3.3 and 4.6 and sheet P), and r177_S285_hypermut_p53_v1.py (p53 activity score of hypermutated versus non-hypermutated TP53 wild-type tumors, the reason for the restriction in Section 3.2.3 → sheet N, block N2).

The plotting and table-formatting scripts that reproduce the published values of Figures 1B, 1D and 8 of the main text and Supplementary Figure S5 (make_Figure1D_v9.py with compose_Figure1_v9.py for the composite of Figure 1 [make_Figure1D_morphology_isoform.py draws panel D alone], make_Figure1B_v10.py with compose_Figure1_v10.py for panel B of all 63 comparison sets, make_Figure2_p53_immune_v7.py and make_Figure3_Biomarker39_clustering_v3.py), the panels of Figures 2, 3 and 9 redrawn with TP53 status (make_Figure8_top_v2.py; make_Figure12EFG_v1.py with compose_Figure12_v2.py; make_Figure13_v2.py with render_svg.py; the script names do not follow the figure numbers), Figure 10 (make_Figure13_overview_v1_S270c.py with its drawn-value file Figure13_values.csv and the cell-by-cell verifier verify_Figure13_overview_S270c.py, the per-specimen table of its discovery block [r164_S270c_discovery_p53_overview_v1.py, independently re-implemented by independent_verify_R164_S270c.py] and the builder and verifier of Supplementary Table S48, sheet M [build_S48_sheetM_S270c.py, verify_S48_sheetM_S270c.py]), Supplementary Figure S10 (make_FigureS10_v6.py; the per-cell-line re-export of the primary table is R29_export_D2_percell_v2.R), Supplementary Figure S12 (make_FigureS13_WSe_TCGA_v1.py; the script name does not follow the figure number) and Supplementary Table S34 (make_TableS34_v2.py), together with S239_FigureS3S4_label_and_key_v1.py, which relabels one specimen with its GSE128213 title and fills the legend keys of Supplementary Figures S3 and S4 drawn by Code6, are included in the Supplementary Methods (Additional file 2). The Supplementary Methods also contain make_Figure4_P2_subgroups_v2.py (patient-level subgroups among the P2 patients), which does not draw any figure of this article. The scripts are provided as executed; the published figures and tables differ from their output only in the English wording and the P1/P2 labels, not in any value. Supplementary Figure S9 is drawn from the values of Supplementary Table S34 (make_FigureS9_v3.py, available from the corresponding author on request), and Supplementary Tables S23 and S33 were compiled from Supplementary Tables S11 and S2 and from the cited literature, respectively.

Code1 and Code2 are existing scripts held by the authors (image classification and RMA normalization); Code1 (the full set of training and inference scripts for image classification) and Code2 (the execution script for RMA normalization—a version restored from the execution record of 2024-12-31 that includes a stage for checking against the normalized output) have both been deposited in the Codes/ directory (indexed in README_SupplementaryMethods). The plotting scripts that are not included with this manuscript (Figures 1A–1C and 1E and the base drawing of Figure 9B–D; Supplementary Figures S2, S9, S11, and S14) are also explicitly identified in the same index.

Additional file numbers follow the order of first citation in the main text; the Supplementary Table numbers (S1–S48) are identifiers and do not follow the order of first citation. The numbers Supplementary Tables S17 and S25 and Supplementary Figure S13 are not assigned in this article. The main text cites all of the remaining 46 tables, this Supplementary Information cites S1, S2, S4–S11, S15, S16, S18, S20, S21, S23, S24, S26, S27, S29, and S31–S48, and the Supplementary Note (Additional file 14) cites S1, S6–S10, S13, S15, S18–S21, S27–S29, S31, S32, S34, S42, S43, and S45; every table in the current set is cited. The contents of Supplementary Tables S35–S39 are as follows. Supplementary Table S35: all results of immune deconvolution (quanTIseq and MCP-counter) and TIDE (Section 3.3.3 of the main text). Supplementary Table S36: MSI-stratified sensitivity analysis of the immune findings (Section 3.3.3 of the main text). Supplementary Table S37: P2_index and p53 target module output by TP53 functional class (definition A and definition B; Section 3.2.2 of the main text). Supplementary Table S38: list of all comparisons by GDSC1/GDSC2 and DepMap (Section 3.2.4 of the main text). Supplementary Table S39 (four sheets; Section 3.2.4 of the main text): sheet S39, P2/P1 correlations, degree of separation, and comparison by CMS for the 198 drugs by oncoPredict; sheet S39b, measured GDSC2 LN_IC50 of cancer cell lines versus the P1/P2 scores for 295 drug entries (DRUG_IDs; they correspond to the 286 GDSC2 drug names of Supplementary Results 13, some drugs being represented by two entries), unadjusted and as partial correlations adjusted for cancer type and TP53 status; sheet S39c, DepMap CRISPR dependency of 17 target genes by TP53 status and versus the scores; sheet S39d, the analysis of sheet S39b additionally adjusted for RAS and BRAF mutation status and restricted to RAS/RAF wild-type lines. The contents of Supplementary Tables S40 and S41 are as follows. Supplementary Table S40: per-line list of nutlin-3 sensitivity (log(IC50)), somatic mutations (TP53_mut, TP53_LOF, KRAS, BRAF, MDM2 gain of function, and MSI), ploidy, and the GSVA scores of the P1/P2 signatures for 65 patient-derived colorectal cancer organoid lines (TP53-mutant 48/wild-type 17) (Section 3.2.4 of the main text). Supplementary Table S41: in the same 65 lines, nutlin-3 sensitivity by TP53 status (exact Wilcoxon rank-sum test and Cliff's delta), linear models with added covariates, Spearman correlations between signature scores and sensitivity (BH correction with a family of 13), partial correlations adjusted for TP53 status, the stratified analysis restricted to the 48 TP53-mutant lines and its sensitivity analysis, and the coverage of the gene sets (Sections 3.2.4 and 4.6 of the main text). The two tables are provided as two separate workbooks (Additional files 32 and 33). Note that although the nominal family size is 13, the coverage of Type5_lncRNA is 0, so that Type5_noimmune is the same set as Type5_core and the number of independent tests is 12; BH correction with a family of 12 does not change any decision with respect to the 0.05 threshold (Supplementary Table S41, note 4). The contents of Supplementary Table S42 are as follows. Supplementary Table S42: list of all tests of the association between the colibactin-related signatures (SBS88 and ID18) and the P1−P2 GSVA axis in the Nunes cohort (U-CAN; WGS + RNA-seq, 1,051 specimens) (block 1: 6 primary tests; block 2: 6 sensitivity/adjusted tests; BH correction with each block as a family; including 9 notes on the test implementation, zero inflation, and the provenance of the numbers of positives per tertile; Section 2 of the main text, Supplementary Note Section 5.2, and Section 4.6). The contents of Supplementary Tables S43–S48 are as follows. Supplementary Table S43: key items of the biomarker and GO analyses (representative genes and top Gene Ontology terms for eight biomarker categories). Supplementary Table S44: comparison of the upstream and downstream aspects of MDM2 promoter regulation (tissue 1, P1-dominant, versus tissue 2, P2-dominant). Supplementary Table S45: comparison of expression-inferred physical properties and Tox functions (tissue 1, autonomous-proliferation type, versus tissue 2, environment-adaptive type). Supplementary Table S46: per-sample morphology-class fractions under both preprocessing conditions (one sheet, 11 columns; documents, per sample, the robustness of the two-group separation to image preprocessing; Section 3.1.2 of the main text). Supplementary Table S47: per-tumor P2_index, TP53 status (with functional class under definitions A and B) and CMS class for the 380 TCGA-COAD/READ primary tumors with P2_index (one sheet, 6 columns; provides the underlying values of Section 3.2.2 and Figure 5 of the main text, cited in the Availability of data and materials). Supplementary Table S48 (Additional file 7): P2 usage and p53 activity in TCGA tumors, organoids and cell lines, and replication in an independent organoid biobank (eighteen sheets, A to P, with a README; the content of each sheet is listed in the description of Additional file 7 at the end of the main text; Sections 2, 3.1.2–3.1.5, 3.2 and 3.3.3 of the main text, also cited in the Availability of data and materials). The number of specimens in the title of Supplementary Table S37 is 374, consistent with Section 3.2.2 of the main text. The index document of the Supplementary Methods is README_SupplementaryMethods.

### **Supplementary Results 13. External data and releases used in the four external validation analyses (D1–D4)**

*Corresponds to Sections 3.2.2, 3.2.4 and 3.3.3 of the main text.*

*The external validations in Sections 3.2.2, 3.2.4 and 3.3.3 of the main text all rely on specific releases of public databases. To guarantee that the values were computed on the same set of cell lines and with the same mutation annotation, we specify here the releases used and the conditions under which they were obtained.*

***DepMap.*** *The genotypes and CRISPR dependencies of the cell lines were all unified to the DepMap 24Q4 Public release (Model.csv, OmicsSomaticMutations.csv, and CRISPRGeneEffect.csv). All 18 contrasts were computed with this release. TP53 status was determined from the VariantInfo column of OmicsSomaticMutations.csv, with cell lines carrying a non-synonymous variant classified as mutant. In 24Q4, among the 1,316 TP53 mutation rows (1,174 cell lines), 0 cell lines carried only synonymous variants, so the rule "cell lines with only synonymous variants are wild-type" and the rule "any variant, regardless of type, is mutant" coincide completely. The 176 cell lines for which no mutation profile was available were excluded from the analysis rather than being defaulted to wild-type. CRISPRGeneEffect.csv comprises 1,178 cell lines × 17,916 genes, and TP53 status could be assigned to all 1,178 cell lines.*

***GDSC.*** *For the dose–response data we used GDSC1 (fitted_dose_response 24Jul22; 333,161 rows / 378 drugs / 970 cell lines) and GDSC2 (same release, 24Jul22; 242,036 rows / 286 drugs / 969 cell lines). Mechanical name matching of the 542 drug names identified four agents targeting the MDM2–p53 pathway: Nutlin-3a, Serdemetan, PRIMA-1MET, and MIRA-1. RG7112, Idasanutlin, AMG-232, Milademetan, HDM201, SAR405838, and APG-115, which are in clinical development, are included in neither GDSC1 nor GDSC2. MSI status follows the annotation in Cell_Lines_Details.xlsx.*

***TP53 Database.*** *For the classification of functional classes we used the MutationView and Function downloads of the TP53 Database release 21 (formerly the IARC TP53 Database; currently maintained by the U.S. National Cancer Institute). Definition A is the conservative definition, which counts as GOF only missense mutations at the 6 hotspot residues (R175H, G245S, R248Q/W, R249S, R273H/C, R282W); definition B is the broader definition, which assigns to the GOF side all missense mutations whose transactivation class is non-functional. The assignment of WT and LOF is identical under either definition, and the two definitions differ only at the boundary of GOF.*

*oncoPredict. For the sensitivity projection in Section 3.2.4 we used oncoPredict 1.3.1; the training set was the GDSC2 expression matrix (17,419 genes × 805 cell lines) and the response matrix of 198 drugs. The TCGA-COAD/READ expression matrix was aggregated per specimen under the same rule as in Supplementary Results 11 (647 aliquot rows → 624 specimens, 36,650 genes), and the 198 drugs were predicted in 10 batches (nine batches of 20 drugs and one of 18). We confirmed that the batching does not affect the results: the maximum absolute difference from the values computed for a single drug alone was 0. The settings of calcPhenotype were batchCorrect = eb (ComBat), removeLowVaryingGenes = 0.2 (removal of the bottom 20% on the raw data), minNumSamples = 10, and powerTransformPhenotype = TRUE (a Box-Cox transformation applied to the drug response on the training side). Note that, because calcPhenotype in oncoPredict 1.3.1 does not write files but returns the matrix as its return value, the way in which the output is captured is version-dependent.*

***TIDE. For the prediction of response to immune checkpoint inhibitors in Section 3.3.3 we used TIDEpy 1.3.8. The TIDE web server (tide.dfci.harvard.edu) has been discontinued, and the successor site, the Cancer Immunology Data Engine (cide.ccr.cancer.gov/tide/), displays the notice "TIDE Server Retired" (an explicit date of discontinuation could not be confirmed). TIDEpy is not a reimplementation of the method but the Python implementation by the original authors' own group (GPLv3, developer Jingxin Fu, liulab-dfci/TIDEpy, PyPI: tidepy), and it has no bibliographic reference of its own. Both TIDEpy itself and the TIDE web platform request citation of the following two reports. (1) Jiang P, Gu S, Pan D, Fu J, Sahu A, Hu X, Li Z, Traugh N, Bu X, Li B, Liu J, Freeman GJ, Brown MA, Wucherpfennig KW, Liu XS. Signatures of T cell dysfunction and exclusion predict cancer immunotherapy response. Nat Med. 2018;24(10):1550-1558. doi:10.1038/s41591-018-0136-1. PMID: 30127393. (2) Fu J, Li K, Zhang W, Wan C, Zhang J, Jiang P, Liu XS. Large-scale public data reuse to model immunotherapy response and resistance. Genome Med. 2020;12(1):21. doi:10.1186/s13073-020-0721-z. PMID: 32102694. Section 3.3.3 of the main text cites these two reports (references [64] and [65]). Note that the only input to TIDE is the pre-treatment tumor expression profile; MSI status, tumor mutational burden, and PD-L1 expression are not among the inputs (Jiang et al. 2018). Because TIDE measures axes independent of MSI (T cell dysfunction and T cell exclusion), its prediction is not a substitute for MSI-High/dMMR status in determining eligibility; Section 3.3.3 of the main text therefore treats the TIDE result as exploratory (the P2-high versus P2-low difference was not reproduced in the specimens with MSI annotation). Deconvolution was performed with immunedeconv 2.1.4 (quanTIseq and MCP-counter). xCell and EPIC were not used because their execution failed.***

***Location of the primary outputs.*** *The primary outputs of D1–D4 are stored under results/WSd/ as D1_*.csv (7 files), D2_*.csv (5 files), D3_*.csv (4 files), and D4_*.csv (4 files), and all values in the main text and in Supplementary Tables S35–S39 are mechanical transcriptions from these primary tables. No estimation or imputation was performed. However, the CMS-specific values in Section 3.2.4 (Kruskal–Wallis P and the medians by CMS) are based on the output (R30_drug_by_CMS_563.csv) of R30_verify_D1_CMS_v1.R, which reconstructed the CMS assignment on a per-specimen basis for 563 specimens.*

***Execution environment.*** *The analyses in Section 3.3.2 (Code8) were run in R 4.6.0 (released 2026-04-24; x86_64-apple-darwin20; macOS Ventura 13.7; locale en_US.UTF-8; random number generator Mersenne-Twister / Inversion / Rounding). The versions of the main packages were GSVA 2.6.3, CMScaller 2.0.1, TCGAbiolinks 2.40.0, SummarizedExperiment 1.42.0, org.Hs.eg.db 3.23.1, AnnotationDbi 1.74.0, GEOquery 2.80.0, survival 3.8-9, survminer 0.5.2, limma 3.68.4, matrixStats 1.5.0, data.table 1.18.4, ggplot2 4.0.3, and dplyr 1.2.1. Code8 is designed to write out sessionInfo() to session_info.txt at the end of the analysis, and that output is included as Codes/Code8_sessionInfo.txt (27 attached packages and 132 namespaces). This record covers Code8 and the scripts run in the same environment (Code11, Code12, Code14, Code16, Code18, Code19, and R30); the tumor-only GSE39582 re-analysis (R49), whose values are reported, has its own record, R49_sessionInfo.txt (R 4.6.0, the same computer; GSVA 2.6.3 / GEOquery 2.80.0). For Section 3.2.2 (Code9), a sessionInfo() output was obtained separately and is included as Code9_v3b_sessionInfo.txt (R 4.6.0, the same computer; UCSCXenaTools 1.7.0 / GSVA 2.6.3 / CMScaller 2.0.1 / TCGAbiolinks 2.40.0 / survival 3.8-9 / survminer 0.5.2 / data.table 1.18.4 / matrixStats 1.5.0 / ggplot2 4.0.3 / dplyr 1.2.1 / org.Hs.eg.db 3.23.1 / AnnotationDbi 1.74.0 / GEOquery 2.80.0). The execution environment for the R analyses in this study was common to all code, including Code13 (immunedeconv), Code15 (oncoPredict), and Code3–Code7 on the discovery-cohort side (the same computer, R 4.6.0). In addition to the Code8, Code9, and R49 records, the Supplementary Methods bundle the sessionInfo() records of R38 and R39 (A13b_sessionInfo.txt and A14c_sessionInfo.txt), R47, R47b, R47c, and R48. In addition, R21, R23, R24, R38, R39, R42, R43, R45, R46, R47, R47b, R47c, R48, and R49 write their own sessionInfo() when run (Section 5 of README_SupplementaryMethods). Because TIDEpy is a Python implementation, it does not appear in the R sessionInfo(). Note that estimate 1.0.13 appears in that record because compute_purity() in Code8 references that package via requireNamespace("estimate"); it is not a record of tidyestimate 1.1.1, with which Code17 computed the tumor purity used in this paper and shared with the companion study of the 15q11-q13 locus (T. Tsukui, R. Yao, and K. Tsuda, unpublished observations) (the two should not be confused).*

### **Supplementary Figure Legends**

*This section collects the legends for Supplementary Figures S1–S12 and S14 (Additional files 10, 11, 22, 23, 52, 43, 44, 27, 45, 30, 61, 62 and 49, respectively; the number S13 is not used). Each figure is referenced from the corresponding section of the main text. Note that the tables of the main text are Tables 1–5, and Table 1 (the confusion matrix of the deep-learning classifier) corresponds to Section 3.1.1 of the main text.*

**Supplementary Figure S1.** Robustness of the differentially expressed gene (DEG) definition (threshold sensitivity analysis and permutation test). With a consistency rate of 70% and |log2 fold change| ≥ 1 (twofold) as the center, the number of DEGs was recalculated over a grid of consistency rate 60–80% × fold change 1.5/2/3-fold. The total number of DEGs changed smoothly with the thresholds—1,997 at 60%/twofold, 1,890 at 70%/twofold, and 1,658 at 80%/twofold (no discontinuous cliff)—and the tissue 1 : tissue 2 ratio remained within the range 53 : 47 to 56 : 44 (928/730 = 56 : 44 at the 80%/twofold grid point). In addition, against the null distribution of a permutation test in which the signs of the fold changes across comparisons were randomized 1,000 times with a Rademacher distribution (number of genes satisfying both criteria by chance: mean 11.3, SD 3.3), the observed value was 1,890 genes (empirical FDR 0.60%, P < 1×10⁻³, z = 567). These results demonstrate the robustness of the threshold choice and of the consistency-based definition. The values were obtained by re-running the analysis with the gene-symbol-restored Supplementary Table S16 as input (generated by Code5 [Supplementary Methods]).

**Supplementary Figure S2.** Relationship between the consistency rate of the direction of change and expression level (log2 fold change). For all genes, the consistency rate of the direction of change across the 33 two-group comparisons (vertical axis) is plotted as a scatter plot against the mean log2 fold change (horizontal axis). The region corresponding to the adopted DEG criteria (consistency rate ≥ 70% and |log2 fold change| ≥ 1) is delineated, and representative marker genes are annotated. The figure visualizes the DEG definition of this study, which uses directional consistency rather than statistical significance (P value, FDR) as the primary criterion. The per-gene values follow the gene-level aggregation of Code5 [Supplementary Methods] applied to Supplementary Table S16; the plotting script is not included in the Supplementary Methods and is available from the corresponding author on request.

**Supplementary Figure S3.** Metastatic trajectories in meta-expression space (principal component analysis, PCA). Principal component analysis of the RMA-normalized expression matrix (log2-transformed) of all 63 specimens using the top 3,000 genes by variance (prcomp, with gene scaling; PC1 = 14.4%, PC2 = 9%). Primary tumor → metastasis pairs from the same patient are joined by arrows, and the trajectories of the three patients with both P1- and P2-dominant specimens (HCT38, HCT41, and HCT67) are emphasized with thick lines. These patients describe longer trajectories that traverse the expression space more widely than the other patients, consistent with a global expression difference accompanying the within-patient difference in promoter usage (exploratory; based on a small number of patients). Generated by Code6 [Supplementary Methods].

**Supplementary Figure S4.** Metastatic trajectories in meta-expression space (UMAP). The same input as in Supplementary Figure S3 (all 63 specimens, top 3,000 genes by variance) was visualized with UMAP (umap package, n_neighbors = 15, random_state = 42). The arrows and thick lines have the same meaning as in Supplementary Figure S3. The long primary → metastasis trajectories of the three patients with both P1- and P2-dominant specimens (HCT38, HCT41, and HCT67) agree with the PCA (Supplementary Figure S3). Generated by Code6 [Supplementary Methods].

**Supplementary Figure S5.** Hierarchical clustering heatmap based on the 39-gene panel identified by IPA Biomarker Detection. Using the 39 genes as input, hierarchical clustering (Manhattan distance, average linkage; the framework follows Eisen et al. [74], without row or column scaling) was performed with an R script written by the authors. Horizontal axis: 63 organoid samples from 21 patients (patient ID–tissue number); the bar above the heatmap shows the MDM2 isoform of each sample (P1, exon 1; P2, exon 2; Additional file 3: Supplementary Table S2). Vertical axis: the 39 biomarker genes (9 wild-type TP53 target genes, 2 gastric metaplasia markers [CTSE, REN], 5 invasion/ECM genes [MSLN, MMP7, SERPINB5, COL17A1, DCBLD2], the DNA damage response gene PRKDC, 9 immune/signaling genes, 2 genes of unresolved annotation and 11 genes of other functions). Color scale: green (low) → black → red (high) log2 RMA intensity over the data range (3.1–12.3). All 63 samples divided into two clusters (51 and 12 specimens; final merge height 57.93 versus 48.66; cophenetic correlation 0.833), agreeing with MDM2 promoter usage in 62 of 63 specimens (98.4%), the sole exception being HCT41-1T; because the 39 genes were selected from the same P1-versus-P2 comparisons, this 62/63 agreement is a within-sample (non-independent) result. The heatmap was redrawn in Python from the same probe-set-level values under the same clustering conditions (script in Additional file 2), reproducing the split, merge heights, cophenetic correlation and 62/63 agreement.

**Supplementary Figure S6.** GSVA signature scores by CMS subtype in TCGA-COAD/READ and by mismatch-repair status in GSE39582. (A) P1 signature score across CMS subtypes (n = 563; Kruskal–Wallis P = 7.2×10⁻¹⁵; CMS1 had the highest median but did not differ significantly from CMS2, P = 0.349, whereas the differentiation core excluding the proliferation module, 13 genes, was highest in CMS2, P = 0.038). (B) P2 signature score without the six immune checkpoint genes (the primary bulk-tumor score, 23 genes; Kruskal–Wallis P = 2.54×10⁻³⁰; highest in CMS3, median 0.397 versus 0.202 in CMS1, P = 1.95×10⁻⁴; full 29-gene score P = 3.3×10⁻²⁸, CMS1 and CMS3 comparable, P = 0.929). (C) The immune-excluded P2 score in GSE39582 tumors with MMR annotation (n = 519; dMMR 75, pMMR 444), higher in dMMR (Wilcoxon rank-sum, normal approximation, P = 6.59×10⁻¹⁰; full 29-gene score P = 1.14×10⁻¹⁷); the 19 non-tumoral mucosa samples were removed before scoring. Pairwise P values are BH-adjusted; the breakdown by functional module is in Additional file 42: Supplementary Table S34.

**Supplementary Figure S7.** Overall survival in TCGA-COAD/READ (n = 591). (A) Overall survival of the P1-high (295) versus P1-low (296) groups by median split of the P1 signature score (log-rank P = 0.114). (B) The same comparison by the P2 signature score (P2-high 295 versus P2-low 296; log-rank P = 0.104). (C) Overall survival by CMS subtype (log-rank P = 0.0072, n = 533; CMS1 91, CMS2 158, CMS3 92, CMS4 192). (D) Forest plot of the multivariable Cox proportional hazards analysis with mutual adjustment of the P1 and P2 scores (clinical stage not adjusted): P1 score HR per 1 SD = 0.852 (95% CI 0.713–1.019), P = 0.080; P2 score HR = 0.893 (95% CI 0.747–1.066), P = 0.210; both confidence intervals cross 1, and after adjustment for age, stage and MSI no independent prognostic contribution of the P1 score was confirmed (Section 3.3.2).

**Supplementary Figure S8.** Per-sample linear fold changes of the key marker genes (table-format figure). The linear fold changes of the key genes defining the tissue-1/tissue-2 contrast (SNORD116, EGR1, MDM2, MMP7 and others; the SNORD116 rows are shown as data only, and their interpretation is given in the companion study on the 15q11-q13 locus [T. Tsukui, R. Yao, and K. Tsuda, unpublished observations]) are shown numerically for each TAC comparison, together with the patient of origin of the tissue-2 (P2)-side samples of each comparison (all values are identical to Supplementary Table S26). MDM2 and MMP7 are consistently more highly expressed on the tissue-2 side in all patients, whereas EGR1 varies between patients (Section 3.1.3 of the main text). This is compatible with upstream inputs to P2 activation (inducing factors) differing between patients (an observation in three patients; not tested), while the core of the P2-dominant program (MDM2 and the metaplastic markers) is commonly retained, in line with the mechanistic hypothesis of this study (Supplementary Note Section 5.2). The figure was generated by make_figS8_v3c.py [Supplementary Methods], whose inputs are Supplementary Tables S26 and S18.

**Supplementary Figure S9.** Specificity of the P1/P2 signatures with respect to the CMS classification (sensitivity analysis). (A) The number of genes shared between each signature (P1, P2) and the CMS calling templates of CMScaller, shown by CMS class. The overlap is at most 3 genes per CMS class on each signature side (CMS1: P1 2/P2 1; CMS2: 2/0; CMS3: 1/3; CMS4: 0/0. Even when summed per class, the maximum is the 4 genes of CMS3). The overlapping genes are CALB1 and DPP4 on the tissue-1 side and CTSE on the tissue-2 side against the CMS1 template (126 genes); OLFM4 and SLC9A3 on the tissue-1 side and none on the tissue-2 side against the CMS2 template (82 genes); OLFM4 on the tissue-1 side and CTSE, TFF1, and TFF3 on the tissue-2 side against the CMS3 template (84 genes); and none on either side against the CMS4 template (237 genes) (the templates comprise 529 genes in total; in this analysis we restricted them to the genes present on the array and used 123 genes for CMS1, 77 for CMS2, 82 for CMS3, and 224 for CMS4). The overlaps are confined to intestinal absorptive epithelium markers (CALB1, DPP4, OLFM4, SLC9A3) and gastric metaplasia markers (CTSE, TFF1, TFF3); the 8 genes of the P1 proliferation module and the 10 p53 target genes, 6 immune checkpoint genes, and 2 cancer-associated lncRNA genes of P2 do not share a single gene with any CMS template. This indicates that the association between the signatures and CMS does not derive from an artifact of gene overlap. (B) The strength of the association between the scored functional submodules (P1: identity/proliferation; P2: metaplasia/p53 targets/immune/lncRNA) and CMS, shown as −log10(Kruskal–Wallis P). Each module is significantly associated with CMS on its own, indicating that the association is not driven by a single module. Note that the P1 identity-TF module consists of HNF4A alone; because GSVA requires gene sets of two or more genes by design, no module score can be computed for it, and it was therefore excluded from this panel (HNF4A itself is included in the P1 signature as a whole). All tests in this panel were performed at the per-specimen (sample) level on 563 specimens (obtained by averaging 574 aliquot rows; the aggregation rule is given in Supplementary Results 11), and the number of genes, composition, per-class medians, Kruskal–Wallis P, and BH-adjusted pairwise comparisons of all 11 modules are listed in Supplementary Table S34. The values correspond to Section 3.3.2 of the main text; the canonical analysis is Code8 (Table_Module_CMS_KW.csv; aliquot-level scores averaged to the 563 specimens), and the figure was drawn from the 563 per-specimen values with make_FigureS9_v3.py, which is not included in the Supplementary Methods (available from the corresponding author on request).

**Supplementary Figure S10.** Comparison of MDM2 inhibitor sensitivity by TP53 and MSI status in the GDSC2 cell line panel. Using the cancer cell line drug sensitivity database GDSC2, we compared the LN_IC50 of drugs targeting the MDM2–p53 pathway stratified by TP53 status (wild-type/mutant) and by MSI status (colorectal cancer cell lines and pan-cancer). Here the MDM2 inhibitors are Nutlin-3a and Serdemetan, whereas PRIMA-1MET and MIRA-1 are mutant p53 reactivators with a different mechanism of action (Section 3.2.4 of the main text). Cell lines with wild-type TP53 (and MSI-High) showed higher sensitivity to the MDM2 inhibitors (lower LN_IC50), supporting, at the cell line level and with publicly available cell line pharmacogenomic data, the conclusion that the molecular background corresponding to the P2-dominant group (wild-type TP53, retained p53 function) matches the pharmacological requirements of MDM2 inhibitors (this is not a validation of drug response in patient specimens; Section 4.6 of the main text). The comparison by MSI status is a background observation; MSI-High is not a proxy for the P2-dominant group (the resemblance of P2 dominance to the MSI-like program is at the signature level only, and in TCGA-COAD/READ P2_index did not differ among the CIN, GS and MSI subtypes and was not associated with MSI status; Sections 3.1.4, 3.2.2 and 4.6 of the main text). The panel layout is: A, pan-cancer, Nutlin-3a; B, colorectal, Nutlin-3a; C, pan-cancer, integrated sensitivity score; D, colorectal, integrated sensitivity score (by MSI status); E, pan-cancer, Serdemetan; F, colorectal, integrated sensitivity score (by TP53 status); G, colorectal, comparison of the four drugs by MSI status. The integrated sensitivity score is the mean of the GDSC1/GDSC2 z-values of Nutlin-3a and Serdemetan; D and F cover the 47 colorectal cell lines, and G covers 46–47 lines depending on the drug (46 for Serdemetan and MIRA-1); they differ only in the stratification axis (D and G by MSI status, F by TP53 status). All of these 47 lines have known TP53 status and MSI status, and they do not include cell lines for which no mutation profile was available. TP53 status and lineage were determined with DepMap 24Q4 Public, and cell lines for which no mutation profile was available were not defaulted to wild-type but were excluded from the analysis. No multiplicity correction was applied across panels. The FDR values, listed alongside the drawn values in the CSV cited below, were Benjamini–Hochberg-adjusted within each block of Supplementary Table S38 for panels A–F and within the colorectal/MSI block of the same table for panel G. Statistical details and all contrasts are given in Supplementary Table S38. All statistics in the figure are mechanically transcribed from the primary outputs of Code12 (results/WSd/), and the drawn values are disclosed in FigureS10_v6_drawn_values_20260803.csv [Supplementary Methods]. Generated with make_FigureS10_v6.py [Supplementary Methods] (the per-cell-line re-export of the primary tables is R29_export_D2_percell_v2.R [Supplementary Methods]).

**Supplementary Figure S11.** Association between colibactin mutational signature burden (SBS88, ID18) and the P1−P2 expression axis (Nunes 2024 cohort). From the Swedish U-CAN colorectal cancer cohort (Nunes et al. Nature 2024 [82]; WGS + RNA-seq), we analyzed the 1,051 specimens (MSS 828, MSI-H 223) for which both expression (ArrayExpress E-MTAB-12862) and COSMIC-decomposed signature activities (Supplementary Table 13 of that paper) were available. The axis is the difference between the GSVA scores of the full P1 and P2 signatures of this study (named Type1_full/Type5_full in the code; gene sets and calculation method identical to those in Section 2 of the main text); larger values indicate P1 dominance. (A, B) Scatter plots of SBS88 (A) and ID18 (B) activity (log₁₀(x+1)) against the axis in all specimens. Spearman ρ = −0.085 (P = 0.0059) / −0.104 (P = 6.9×10⁻⁴). The negative correlations indicate that the colibactin exposure footprint is only slightly shifted toward the P2-dominant side (the correlations are weak). (C) Signature positivity rates (activity > 0) in the P1-high and P2-high groups defined by the tertiles of the axis (351 specimens each). SBS88 4 versus 16 (Fisher's exact P = 0.011); ID18 20 versus 38 (P = 0.019). (D, E) When restricted to MSS (n = 828), the negative correlations are retained and, if anything, strengthened (SBS88 ρ = −0.121, P = 5.0×10⁻⁴; ID18 ρ = −0.156, P = 6.4×10⁻⁶). (F) Summary of effect sizes. In partial Spearman correlation (covariates: MSI, stage, site, age), ID18 is significant (ρ = −0.084, P = 0.0067) and SBS88 is borderline (ρ = −0.060, P = 0.053). The tests were BH-corrected with each block of 6 tests as a family (Supplementary Table S42). The Spearman P values are based on the t approximation because of ties, and the Wilcoxon tests (Supplementary Table S42) used the normal approximation because both groups were ≥ 50. The activities are zero-inflated (positives: SBS88 31, ID18 82; all positives are MSS), and the effects should be read from the rank correlations and the positivity rates. The values were computed by R42_WSe_colibactin_Nunes_v2.R [Supplementary Methods]; the plotting script is not included in the Supplementary Methods and is available from the corresponding author on request.

**Supplementary Figure S12. Reproduction analysis in TCGA-COAD/READ of the association between colibactin mutational signature burden and the P1/P2 axis. We analyzed the 374 specimens for which all of the following were available: the SBS88 and ID18 activities fitted with sigminer (sig_fit) to the deposited somatic mutation data (maf_coadread.rds; predominantly WES, 616 specimens, GRCh38; COSMIC v3.1-series reference), P2_index (isoform quantification), and the P1−P2 GSVA axis. The forest plot shows Spearman ρ with the signs aligned to the direction of reproduction of the Nunes cohort (denser on the P2 side = positive; because P2_index is negatively correlated with the P1−P2 axis [ρ = −0.156], the P2_index rows keep their sign and the expression-axis rows are sign-reversed). Only P2_index × ID18 was significant in the direction of reproduction (ρ = +0.143, P = 0.0055, FDR = 0.044; tertile positivity rates 20/125 versus 7/125, Fisher's exact P = 0.013, OR = 3.20); the expression GSVA axis (P1−P2) showed no significant association in the same direction, and SBS88 was not significant on either axis (BH correction with each block of 12 tests as a family; Spearman by t approximation because of ties, Wilcoxon by normal approximation, positivity rates by Fisher's exact test). The gray squares are the observed values in the Nunes cohort (for reference). Because TCGA is predominantly WES, the detection sensitivity for ID18 and SBS88 is lower than that of WGS (activity-positive 11.0% and 13.6%). The MSS-restricted sensitivity analysis is uninformative because, even after supplementation from the public annotation, per-patient MSI annotation was available for only 32 of the 374 specimens (MSS 24) (FDR = 1 in all tests with n = 24). This analysis is exploratory, and the reproduction depends on the choice of axis (Section 4.6 of the main text). Stratification of these 374 specimens by TP53 status and hypermutation (analysis R168; Supplementary Table S48, sheet P) shows that the reproduced association does not run through p53 activity: the activities are unrelated to the p53 activity score in TP53 wild-type tumors, the positivity rate is the same in wild-type tumors with and without evidence of p53 activity, and the correlation with P2_index is present in TP53-mutant tumors (rho = +0.209) and absent in wild-type tumors (rho = +0.026).**

**Supplementary Figure S14.** Integrated hub-molecule network (Cytoscape synthesis of all 33 IPA analyses by the authors). The factors that appeared among the top ranks in each of the modules — Canonical Pathway, Upstream Analysis, Disease and Bio Functions, Tox Function, Regulator Effects analysis and Networks analysis — across all 33 two-group comparisons were aggregated, and the central hub molecules that recurred in common across multiple analyses were integrated and visualized by the authors as a single network diagram using Cytoscape version 3.10.4 [157] (Supplementary Tables S20 and S21; Additional files 50 and 51). Red = nodes in the P1/tissue-1 direction; blue = nodes in the P2/tissue-2 direction. The figure shows a structure in which a proliferation hub centered on the FOXM1–MYC–E2F axis (P1/tissue 1) is juxtaposed with an apoptosis/environment-adaptation hub that includes TP53, CDKN1A, RBL1/2 and immune-related molecules (P2/tissue 2). This is not the IPA-native Graphical Summary but an original integrated diagram produced by the authors.

### **References not cited in the main text**

*Numbering continues from the reference list of the main text; only the references that are cited in this Additional file and not in the main text are listed.*
