## Additional files for "TP53 status aligns with morphology and metaplastic programs in colorectal cancer organoids, and MDM2 P2 promoter usage reflects p53 activity": Additional_file_14_Supplementary_Note.docx

### Supplementary Note: Molecular landscape of the two organoid groups

This Supplementary Note contains the details (methods, results, and discussion) of the IPA (Ingenuity Pathway Analysis) and EnrichR GO analyses that are summarized in the main text (Sections 2, 3.3.1, 4.2.1 and 4.5). The figures, tables, supplementary tables, supplementary figures, and Supplementary Results referred to in this Note, as well as the bracketed reference numbers [N], are those of the main text and of the Supplementary Information (Additional file 12); references cited only in this Note are listed at its end, with numbering continuing from the main text. Section numbers refer to sections of the main text unless explicitly written as “Section N of this Note.” As in the main text, “tissue 1” and “tissue 2” denote the MDM2 P1-dominant and P2-dominant specimen groups, which largely coincide with TP53-mutant and TP53 wild-type specimens, respectively; the descriptive labels “autonomous-proliferation type” and “environment-adaptive type” summarize their expression programs and do not imply that promoter usage determines them.

### 1 Supplementary Methods: details of the IPA and EnrichR GO analyses

#### 1.1 Ingenuity Pathway Analysis (IPA)

Using Ingenuity Pathway Analysis (IPA; Qiagen) [72], we performed the following analyses. Data were analyzed through the use of IPA (QIAGEN Inc., https://www.qiagenbioinformatics.com/products/ingenuity-pathway-analysis)[72]. Each IPA analysis was performed independently on the comparison data for each of the 33 pairs contrasting tissue 1 with tissue 2, and the results were aggregated on the basis of frequency of appearance using the comparison function of IPA. The aggregation was carried out in two groups, the first half (the first 17 of the 33 analyses in order of analysis ID, analyses 49–92) and the second half (the remaining 16, analyses 93–109); “first-half Rank N” and “second-half Rank N” in the main text denote the rank of frequency of appearance within each group, and “N/M consistent” indicates that the same result was reproduced in N of the M analyses examined. An M smaller than 33 indicates missing data. The assignment of each analyzed item to a tissue class (direction of activation) was determined by a majority vote of the signs of the Activation z-score of that item across the 33 comparisons; that is, items for which positive signs predominated were assigned to tissue 1 (P1-dominant, autonomously proliferative), and items for which negative signs predominated were assigned to tissue 2 (P2-dominant, environment-adaptive).

Furthermore, as criteria for adopting only highly reproducible, robust findings in the main text, we required that both of the following be satisfied: (i) the sign consistency of the Activation z-score across the 33 analyses be at least 75%, and (ii) the absolute value of the median Activation z-score across the 33 analyses be at least 2 (|median z| ≥ 2). Criterion (i) is a requirement specific to this study: because a single IPA run may be unstable depending on the choice of comparison, it requires reproducibility of the direction of activation across the 33 comparisons (which reuse the same P2 specimens and are therefore not independent replicates). Criterion (ii) corresponds to the threshold conventionally used in IPA to judge significant activation or inhibition (|z| ≥ 2). Under these criteria, items whose sign distribution was nearly even and whose direction of activation could not be determined (e.g., HNF4A; sign consistency 52%, median z = −0.14) were excluded from the claims of assignment to a tissue class in the main text, and for items whose direction was consistent but whose effect size did not reach the threshold (e.g., CDX1; sign consistency 93%, median z = +1.61), we explicitly stated at the relevant places that they are sub-threshold. The frequency ranks (first-half group: analyses 49–92; second-half group: analyses 93–109) are given alongside each item as descriptive information in Supplementary Tables S6–S9 but were not used as a criterion for adoption, because the rank is an index reflecting how frequently the item appears in the IPA output and would introduce information of a different kind from the effect size and the reproducibility of the direction of activation. The robustness of the conclusions to the choice of adoption criteria was verified by a sensitivity analysis in which the sign consistency (70/75/80%), the rank cutoff (20/25/30/50/100/unlimited), and the |median z| threshold (not applied/≥ 2) were varied over a grid (Supplementary Table S10; Supplementary Results 10).

The two criteria above were applied to Upstream Analysis, for which assignment to a tissue class is claimed on the basis of the effect size of activation. All of the upstream regulators for which the main text claims assignment to a tissue class on the basis of the Upstream Analysis results (listed in Sections 2.2, 3.2, 4.3, and 5.3 of this Note) satisfy a sign consistency of at least 75% and |median z| ≥ 2 (among the listed factors, the smallest |median z| is 2.04, for CTNNB1). In contrast, the listings in the results sections for Canonical Pathway and for Disease and Bio Functions (Sections 2.1, 2.3, 3.1, and 3.3 of this Note) are descriptions based on the rank of frequency of appearance across the 33 analyses and on the reproducibility of the direction of activation, and no selection by effect size was performed. Because these modules include items with |median z| < 2, such items are explicitly indicated as sub-threshold in each section and are treated not as independent claims of activation but as descriptions based on frequency of appearance and reproducibility of direction. Tox Function, in turn, includes items with |median z| < 2 because this module by its nature contains many items with small absolute z-scores; the findings of this module were therefore used only as corroborating evidence supporting the findings of the other modules, not as independent claims. However, for the Regulator Effects analysis (5) described below—and for that analysis alone—the integration of all 33 analyses could not be completed because of the large computational load; the above framework of sign aggregation and first-/second-half ranking across the 33 analyses was therefore not applied, and the results of the first 8 of the 33 analyses were integrated (see (5) for details).

**(1) Canonical Pathway analysis:** significant enrichment in known biological signaling and metabolic pathways was evaluated.

**(2) Upstream Analysis:** transcription factors, cytokines, kinases, and other molecules that function as upstream regulators were identified, and their activation/inhibition states were inferred.

**(3) Disease and Bio Functions:** disease phenotypes and biological functions associated with the analyzed gene sets were evaluated.

**(4) Tox Function:** associations with toxicity-related signals and with clinical-chemistry changes were evaluated.

**(5) Regulator Effects analysis:** the causal chain from upstream regulator through regulated target molecules to downstream disease phenotype was quantitatively evaluated with the Consistency Score. This analysis is a module unique to IPA that answers the causal-inference question of which molecules cause which disease phenotypes through which targets. Whereas the other IPA modules integrated all 33 analyses, for this Regulator Effects analysis alone the integration of all 33 analyses could not be completed because of the very large computational load, and the results of the first 8 of the 33 analyses (analyses 49, 50, 51, 52, 53, 54, 59, and 63) were integrated. Accordingly, the ranks and Consistency Scores presented for this module are based on the integration of these 8 analyses.

**(6) Networks analysis:** molecular interaction networks within the analyzed gene sets were constructed on the basis of the IPA database of known interactions, functional clustering and scoring were performed, and the central interaction modules were identified.

**(7) Graphical Summary: the top-ranked factors from each module — Canonical Pathway analysis (1), Upstream Analysis (2), Disease and Bio Functions (3), Tox Function (4), Regulator Effects analysis (5), and Networks analysis (6) — were aggregated across all 33 analyses, and the central hub molecules that recurred across multiple analyses were integrated and visualized by the authors as a single network diagram in Cytoscape (version 3.10.4) [157]. By identifying the network hubs on which the top factors of each analysis converge, we summarized the principal molecular differences between the two groups.**

#### 1.2 EnrichR GO analysis

The lists of differentially expressed genes obtained from the two-group comparison analysis (TAC) were subjected to Gene Ontology (GO) enrichment analysis using the Enrichr R package [73]. The databases referenced were the three categories GO Molecular Function (GO MF), GO Cellular Component (GO CC), and GO Biological Process (GO BP) in their 2026 versions, and statistical significance was judged by the FDR-corrected P value obtained with the Benjamini–Hochberg (BH) procedure (adjusted P-value; adj. P < 0.05) [73]. For each list, the top 400 differentially expressed genes were entered. The GO terms characteristic of each tissue (tissue 1 and tissue 2) are described in this Note. GO MF describes the molecular functions of proteins (enzymatic activity, binding activity, etc.), GO CC describes subcellular localization and cellular structures (organelles, complexes, etc.), and GO BP describes the biological processes carried out by cells (metabolism, signal transduction, morphogenesis, etc.). This allows functional features to be captured in an unbiased manner, independent of the IPA database of known pathways.

### 2 Molecular pathway landscape of tissue 1 (P1-dominant, mostly TP53-mutant specimens)

##### 2.1 IPA Canonical Pathway analysis: simultaneous maximization of RNA processing, ribosome biogenesis, and DNA replication

In the IPA Canonical Pathway analysis, the pathway that ranked first overall (present in all 33 analyses, 33/33) was “Processing of Capped Intron-Containing Pre-mRNA,” followed in second place by Cell Cycle Checkpoints (32/33 consistent) (Table 3). The position of the pre-mRNA processing pathway at the very top reflects the large volume of transcriptional activity that sustains the vigorous proliferation of tissue 1 and also indicates high activity of the splicing machinery. “Major pathway of rRNA processing in the nucleolus and cytosol” (29/33 consistent) was ranked 4th and “rRNA modification in the nucleus and cytosol” (32/33 consistent) 10th, such that pathways related to ribosome biogenesis accumulated among the top ranks. This is mechanistically consistent with the high expression of ribosome biogenesis genes (NES = 2.53) reported by Okamoto et al. (2022) [12] in the Type 1 PDOs of the same-biobank cohort. Pathways related to cell cycle control and DNA repair, such as Cell Cycle Checkpoints (2nd), Mitotic Metaphase/Prometaphase (3rd and 5th), Synthesis of DNA (6th), DNA Replication Pre-Initiation (7th), and HDR/NHEJ (9th), also accumulated among the top ranks. In addition, Telomere Maintenance (first-half Rank12, second-half Rank20), which is involved in maintaining replicative lifespan, was found among the top ranks. The SUMOylation pathways of chromosome organization, DNA replication, and DNA damage response proteins were also enriched at high ranks (SUMOylation of chromatin organization proteins: first-half Rank14, second-half Rank13; SUMOylation of DNA replication proteins: first-half Rank17, second-half Rank11; SUMOylation of DNA damage response and repair proteins: first-half Rank35, second-half Rank16), indicating activation of a mechanism that supports vigorous DNA replication and repair and chromosome maintenance at the level of post-translational modification (SUMOylation). It is also noteworthy that Cilium Assembly (ciliogenesis; first-half Rank38, second-half Rank23) was ranked among the top pathways. The RB-family pocket proteins RBL1/RBL2 are known to regulate ciliogenesis, which is consistent with the inactivation of RBL1/RBL2 in tissue 1 (Section 4.3 of this Note); together with Activation of anterior HOX genes (first-half Rank57, second-half Rank50), this may constitute a molecular point of contact linking the cell cycle, cilia, and morphogenesis. Furthermore, the RHO GTPases Activate Formins pathway (first-half Rank24, second-half Rank12, 94% sign consistency), which is responsible for nucleating polymerization of the actin cytoskeleton, was also robustly activated in tissue 1 and, together with Cilium Assembly, is consistent with a mechanism that supports the dense, tubular, well-organized epithelial morphology of tissue 1 at the level of the cytoskeleton. This pathway profile is consistent with that of CRC arising through the chromosomal instability (CIN) pathway (at the cohort level, the differentiation core of the P1 signature is highest in CMS2, whereas the whole P1 signature, including the proliferation module, is highest in CMS1; Section 3.3.2 of the main text).

In addition, the accumulation of Gene Silencing by RNA and the MicroRNA Biogenesis Signaling Pathway indicates that gene expression is finely controlled through post-transcriptional regulation.

With respect to energy metabolism, a group of aerobic metabolic pathways, including oxidative phosphorylation, the TCA cycle, and the pentose phosphate pathway, accumulated (Table 3). This indicates efficient energy metabolism with maximized mitochondrial function, going beyond simple dependence on anaerobic glycolysis (the Warburg effect), and the accumulation of Complex I/III/IV assembly and Cristae formation corroborated that the structural integrity of the mitochondrial inner membrane is maintained to a high degree.

Among DNA repair pathways, the major repair pathways, from HDR and NHEJ to BER, accumulated comprehensively (Table 3). Although DNA damage occurs at high frequency in the CIN pathway, the expression data suggest that tissue 1 maintains DNA repair and a high proliferation rate simultaneously, with these multiple repair mechanisms upregulated together, which would help preserve the functional integrity of the genome while sustaining that rate [158].

With respect to signal transduction, Hedgehog (both the ‘on’ and ‘off’ states) and the bidirectional regulatory pathways of WNT/β-catenin accumulated simultaneously (Table 3). This seemingly contradictory finding suggests that WNT signaling oscillates dynamically and that a fine balance is being struck between the maintenance of stemness and differentiation. The accumulation of Notch4 signaling, PPAR and PPARα/RXRα activation, FXR/RXR activation, and estrogen-related signaling was also characteristic.

##### 2.2 IPA Upstream Analysis: coordination of the MYC/E2F axis and intestinal identity factors

In the Upstream Analysis, the activated factors in the tissue-1 direction that met the adoption criteria (sign consistency of at least 75% and |median z| ≥ 2) were headed by MYC (median z = +5.76, 32/33 consistent, first-half Rank3), Eldr (+6.29, 31/32, first-half Rank4), CEBPB (+5.46, 31/33, first-half Rank8), RABL6 (+5.17, 30/31, first-half Rank10), and TFEB (+4.79, 32/33, first-half Rank12); a total of 121 factors (101 molecules and 20 chemicals), further including TBX2, E2F, FOXM1, MDM4, and AURKB, met the same criteria (the full list is given in Supplementary Table S7; E2F1 and MYBL2, described in Section 3.3.1, also meet the same criteria). On the other hand, ERBB2 (median z = +1.96, sign consistency 88%), PPARGC1A (+1.88, 88%), CDX1 (+1.61, 93%), BCL6 (+1.54, 94%), and TFAP2C (+0.94, 84%) showed high sign consistency in the tissue-1 direction, but their effect sizes did not reach the adoption threshold (sub-threshold), and NRF1 (+1.29, 74%) and HNF4A (−0.14, 52%) did not meet the sign consistency criterion; these were therefore excluded from the interpretation below (Supplementary Table S7).

With respect to intestinal identity, the intestinal-type homeobox transcription factor CDX1 showed high sign consistency (93%) in the tissue-1 direction, but its effect size, median z = +1.61, remained at a sub-threshold level that did not reach the adoption threshold (|median z| ≥ 2). As Verzi et al. showed in Proceedings of the National Academy of Sciences (2010) [159], CDX2 and TCF4 bind cooperatively to intestine-specific cis-regulatory regions, thereby maintaining the intestinal-type gene expression program and defining the intestinal phenotype of differentiated colorectal cancer. By contrast, for HNF4A, the transcriptional master regulator of intestine-specific genes, the sign consistency across the 33 comparisons was 52% (median z = −0.14), so the direction of activation was undetermined and HNF4A could not be assigned to either tissue class (Supplementary Table S7). Therefore, the interpretation that tissue 1 retains its identity as “colon” is suggested by the IPA Upstream Analysis only as a sub-threshold tendency toward activation of CDX1 (predicted activity; CDX1 mRNA itself is slightly higher on the tissue-2 side, Supplementary Report Section R-20); its confirmatory basis must be sought in the expression-level findings for the intestinal markers (OLFM4, LGR5, SLC26A3) (Section 3.3.1). In contrast, activation of MYC (median z = +5.76), E2F (+3.87; E2F1 +2.15, E2F3 +3.54), FOXM1 (+2.61), MYBL2 (+2.44), MYCN (+2.67), MYCL (+2.77), AURK (+3.29), CDK19 (+3.92), and LIN9 (+3.28) met the adoption criteria in every case, indicating that a transcriptional program that comprehensively accelerates the G1/S transition of the cell cycle (MYC/E2F), the execution of M phase (FOXM1), and mitosis (AURK) has been established (E2F2, at +1.74, is sub-threshold).

In addition, pifithrin alpha (a p53 inhibitor; median z = +1.999, sign consistency 100%, 31/31 consistent [z-scores were calculated in 31 of the 33 analyses]) was identified as an upstream regulator. Its effect size is at a sub-threshold level, falling just short of the adoption threshold (|median z| ≥ 2), but its sign consistency of 100% is the highest of all. The activation of MDM4 (+2.73, 87%) meets the adoption criteria. From the IPA side, these findings are consistent with p53 being inactive on the tissue-1 side—through TP53 mutation or p53 inactivity in the wild type (Section 3.2.1: two of the three TP53 wild-type P1-dominant specimens, both from HCT59, had p53 activity scores at or below the 90th percentile of mutants, and the p53 suppressors MDM4, PPM1D and USP7 did not differ significantly between groups)—and provide corroborating support for a lack of P2 promoter activation and a P1-led pattern of MDM2 expression (see Section 3.1.3 and Additional file 1, Section R-18). The tendency toward activation of ERBB2 (median z = +1.96, sign consistency 88%; sub-threshold) was directionally consistent with the features of HER2-amplified CRC.

##### 2.3 IPA Disease & Bio Functions: convergence on proliferation, DNA repair, and gland formation

In the IPA Disease & Bio Functions analysis, the following biological processes were identified as functional consequences of tissue 1 (Supplementary Table S8: details of IPA Disease & Bio Functions). Repair of DNA, Cell viability of tumor cell lines, Checkpoint control, Double-stranded DNA break repair, Cell proliferation of tumor cell lines, Cell viability, Survival of stem cell lines, DNA replication, Colony formation, Abnormal morphology of gland, and Colorectal adenoma occupied the top ranks. Of these, DNA replication (median z = +1.38, sign consistency 97%), Abnormal morphology of gland (+1.50, 75%), and Colorectal adenoma (+0.86, 85%) are sub-threshold in effect size. In particular, the accumulation of “Abnormal morphology of gland” and “Colorectal adenoma” is directionally consistent with the morphological features of colorectal adenocarcinoma, which histologically forms glandular structures.

##### 2.4 IPA Tox Function: organ-spanning activation topped by nephritis signals

In the Tox Function analysis of tissue 1, a renal and cardiac proliferative toxicity signature headed by Nephritis (median z = +2.47, first-half Rank1, 30/33 consistent) accumulated. The signals meeting the adoption criteria (sign consistency of 75% or higher and |median z| ≥ 2) were Nephritis and Hyperplasia of kidney cells (+2.00, first-half Rank6, 25/27); Glomerulonephritis (+1.47, first-half Rank5, 32/33) and Hypertrophy of cardiomyocytes (+0.81, 85%) were consistent in direction but sub-threshold in effect size, and Congenital heart disease (+0.14, 55%) did not meet the sign consistency criterion either (Supplementary Table S9). These signals reflect the vigorous proliferative program of tissue 1 and contrast with the metabolic ALP signature of tissue 2 (Section 3.4 of this Note; all signals and the details of their interpretation are given in Supplementary Results 1 of the Supplementary Information).

##### 2.5 EnrichR GO analysis: accumulation of amino acid transport, DNA replication, and apico-basal polarity of the intestinal epithelium

The EnrichR GO analysis confirmed enrichment of the apico-basal polarity characteristic of the intestinal absorptive epithelium and of proliferative metabolism, represented by L-leucine/branched-chain amino acid transport (BP), DNA replication and DNA metabolic process (BP), and Basolateral/Apical Plasma Membrane (CC) (the list of all enriched terms [BP/CC/MF] is given in Supplementary Results 2 of the Supplementary Information).

### 3 Molecular pathway landscape of tissue 2 (P2-dominant, mostly TP53 wild-type specimens)

##### 3.1 IPA Canonical Pathway analysis: the triangle of inflammation, invasion, and hypoxic adaptation

In the IPA Canonical Pathway analysis, the top ranks were occupied by Pathogen Induced Cytokine Storm Signaling Pathway, Extracellular Matrix Organization, Role of Osteoclasts/Chondrocytes in Rheumatoid Arthritis, Cell Junction Organization, RAF/MAP Kinase Cascade, Degradation of the Extracellular Matrix, Assembly of Collagen Fibrils, IL-17 Signaling, Mitochondrial Dysfunction, NF-κB Signaling, Neuroinflammation Signaling, Death Receptor Signaling, Colorectal Cancer Metastasis Signaling, VEGF Signaling, Interleukin-4/13 Signaling, Regulation of EMT by Growth Factors, HIF1α Signaling, Tumor Microenvironment Pathway, Ferroptosis Signaling Pathway, and CGAS-STING Signaling Pathway (Table 3; Supplementary Table S6). Of these, nine pathways—IL-17 Signaling (median z = −1.77, sign consistency 91%), Neuroinflammation Signaling (−1.09, 85%), Death Receptor Signaling (−1.83, 100%), Colorectal Cancer Metastasis Signaling (−1.86, 85%), Interleukin-4/13 Signaling (−1.52, 91%), Regulation of EMT by Growth Factors (−1.46, 82%), HIF1α Signaling (−1.52, 94%), Ferroptosis Signaling Pathway (−1.52, 91%), and CGAS-STING Signaling Pathway (−1.72, 94%)—showed high reproducibility of the direction of activation but effect sizes below the threshold (sub-threshold), and their description here is based on frequency of appearance and reproducibility of direction. Note that IL-17A Signaling in Fibroblasts (sign consistency 73%) and Necroptosis Signaling Pathway (67%) did not meet the primary criterion of 75% sign consistency and were therefore excluded from the list. In addition, “Neutrophil degranulation” (second-half Rank6, 30/33 consistent) was enriched on the tissue-2 side. Because organoids lack stroma and immune cells (Section 4.6), this does not indicate neutrophil involvement; it is interpreted as an overlap between secretory and granule-associated genes expressed by the epithelial cells themselves (TCN1, CTSH, B2M, and others) and the neutrophil-granule annotation. The same annotation overlap underlies the significant enrichment of Tertiary Granule Lumen (adj. P = 0.0007; annotated as neutrophil tertiary granules) and Specific Granule Lumen (adj. P = 0.023) in the EnrichR GO CC analysis (Section 3.5 of this Note; Supplementary Results 4 of the Supplementary Information).

The fact that “Pathogen Induced Cytokine Storm” is among the top-ranked tissue-2 pathways (sign consistency 100%, median z = −3.32) is a decisive difference from tissue 1. This indicates that the epithelial cells of tissue 2 themselves strongly express an inflammatory and stress-response program (for the molecular link between inflammation and cancer, see reference [160]) and is consistent with the simultaneous activation of multiple pro-inflammatory cytokines (TNF, IL-1B, IL-6, IL-17, IL-33, CXCL8, CXCL14, CCL20). Because organoids lack stroma and immune cells, this finding does not itself indicate inflammation of the tumor microenvironment.

The accumulation of “Colorectal Cancer Metastasis Signaling” (effect size below the threshold) is directionally consistent with expression of a molecular program associated with metastasis. At the same time, the accumulation of ECM Organization, Degradation of ECM, Regulation of EMT, and Integrin Signaling is consistent with an ECM-degradation expression program centered on MMP7 (100% consistent, avg FC −111) (invasive capacity itself was not measured). MMP1 has an avg FC of −279, but its median is +1.2 and its consistency is only 42.4% (14/33), indicating a patient-specific expression change.

The accumulation of HIF1α Signaling (median z = −1.52, sign consistency 94%; effect size below the threshold) suggests that tissue 2 may be in a hypoxic environment [161]. Note that HIF-1α has been reported to interact directly with MDM2 and thereby modulate p53 function [153], and it may thus constitute a point of contact between the hypoxic environment and the MDM2–p53 axis. The elevation of AST/ALT/ALP/LDH predicted in the IPA Tox Function analysis (of which only ALP meets the adoption criteria; Section 3.4 of this Note) may reflect cell necrosis, and a vicious cycle may be formed in which the “hypoxic niche” created by necrosis further activates the HIF1α pathway. The high expression of CA9 (carbonic anhydrase 9) within the gene list also supports survival adaptation under hypoxia [162]. B2M (β2-microglobulin; avg FC −115) is a component of the MHC class I complex and was higher on the tissue-2 side in 73% of the 33 comparisons (median −0.5), consistent with a signature-level resemblance to the CMS1 immune module (not an indication of MSI status; Section 4.2.1; the B2M cluster proper on chromosome 15 shows even higher consistency, 79–88% each, and the mean consistency is diluted when the B2M-like sequences on chromosome 16 are included). The accumulation of CGAS-STING Signaling Pathway (median z = −1.72, sign consistency 94%; effect size below the threshold) deserves particular attention. This pathway senses DNA leaked into the cytoplasm and activates an immune response, and, as Bakhoum et al. showed in Nature (2018) [163], chromosomal instability (CIN) induces cytoplasmic DNA leakage and activates the STING pathway. However, the Aneuploidy and DNA damage predictions for tissue 2 are likely inferred from the lower expression in tissue 2 of mitotic and checkpoint genes that are more highly expressed in tissue 1 (Section 3.3 of this Note) and do not indicate chromosomal instability in tissue 2 (Section 3.1.4). The accumulation of CGAS-STING (sub-threshold) therefore cannot be attributed to CIN-derived cytoplasmic DNA, and its origin remains unresolved.

The accumulation of Ferroptosis Signaling Pathway (median z = −1.52, sign consistency 91%; effect size below the threshold) is directionally consistent with the Necrosis and cell-death signals observed in the IPA Tox Function analysis and suggests that modes of cell death other than classical apoptosis may be involved in tissue 2. Note that Necroptosis Signaling Pathway also appeared on the tissue-2 side, but because it has a sign consistency of 67% and a median z of −0.73 and thus meets neither of the adoption criteria of this study, we do not interpret it as activated.

##### 3.2 IPA Upstream Analysis: dominance of wild-type p53 and stress-response factors

The factors identified as activated upstream regulators in the Upstream Analysis, headed by NUPR1 (median z = −8.59, sign consistency 94%) and TP53 (−7.14, 100%) and including IGF2BP1, CDKN2A, CDKN1A, TGFB1, SMARCA4, SMAD3, IL1B, TGFB3, TNF, EGF, FGF2, HIF1A, IL6, and CTNNB1, all met the adoption criteria (sign consistency of 75% or higher and |median z| ≥ 2) and showed a profile diametrically opposite to that of tissue 1 (PRKCD, STAT4, TP73, RBL2, MAPK3, MAPK8, FOXO3, HNRNPK, and others also met the criteria and were identified in the same direction. RELA, SRC, E2F6, IFNG, and AKT1 agreed in direction but had effect sizes below the threshold. The median z and sign consistency of each factor are given in Supplementary Table S7). Note that EGR1, although its sign consistency in the tissue-2 direction was high (91%), was sub-threshold, with its effect size (median z = −1.95) falling slightly short of the adoption threshold (see the discussion later in Section 3.2 of this Note).

The most important finding is that TP53 is listed as an activated upstream regulator. This agrees with p53 being actually active in the P2-dominant specimens (the p53 activity score separated wild-type P2-dominant specimens from TP53-mutant specimens [AUC 0.99]; because the P2 signature shares genes such as CDKN1A, BAX, and BBC3 with the p53 activity score, and the P2-dominant group was defined in part through differences in p53 target expression, this separation may partly reflect circularity; Section 3.2.1). The activation of wild-type TP53 (with EGR1 in the same direction but below the adoption threshold) is consistent with transcriptional induction of the MDM2 P2 promoter (see Section 3.1.2 and Figure 1) and indicates activation of the p53–MDM2 autoregulatory loop. Increased production of MDM2 protein may promote the ubiquitination and degradation of p53 and contribute to the maintenance of cancer cell survival [3] (for the detailed mechanism, see Section 5.2 of this Note).

The high-ranking activation of NUPR1 (Nuclear Protein 1, also known as p8 or COM1) is another noteworthy finding. NUPR1 is a stress-responsive transcriptional cofactor and, as Goruppi and Iovanna showed [164], is involved in the stress tolerance of cancer cells and in the regulation of p53 function. The activation of NUPR1 in tissue 2 suggests the existence of a mechanism regulating p53 activity through stress-response pathways, in addition to the MDM2 P2-dependent regulation of p53.

The activation of TGFB1, TGFB2, TGFB3, and SMAD2/3/4 is a major driver of EMT (epithelial–mesenchymal transition). As Kalluri and Weinberg showed in the Journal of Clinical Investigation (2009) [111], EMT, for which TGFβ/SMAD signaling is a major inducer, confers mesenchymal properties (enhanced migratory capacity, invasiveness, and resistance to apoptosis) on epithelial cells. Thiery et al. (2009) [110] comprehensively described the roles of EMT in development, tissue repair, and cancer progression and established that EMT confers migratory capacity, invasive capacity, resistance to apoptosis, and stem cell-like properties on cells. Furthermore, the consensus statement by Yang et al. (2020) [112] defines EMT not as a single binary switch but as a continuum of intermediate states (partial EMT) between the epithelial and mesenchymal types, and the complex phenotype observed in tissue 2, in which gastric-type and intestinal-type features coexist, is consistent with this concept of “partial EMT.” In the Upstream Analysis of this study, CTNNB1 (β-catenin), EGF, and FGF2 were identified as activated upstream regulators in addition to SMAD3 and TGFB1, and the combined activation of TGFβ, Wnt, and FGF signaling may be directionally consistent with partial EMT. However, the difference in EMT score between P2-dominant and P1-dominant specimens was not significant (P = 0.079); the EMT score was higher mainly in metastatic specimens (patient-level paired Wilcoxon P = 0.004, 17 patients), and confounding by sampling site is possible (Sections 3.1.2 and 4.6). EGR1 is a transcription factor that may regulate MDM2 expression in a context-dependent manner through the EGR1-binding sequence of the MDM2 promoter, and the P2 promoter itself has been reported to be activated by multiple transcription factor response elements [9]. Indeed, EGR1 has also been reported to repress MDM2 transcription in head and neck squamous cell carcinoma (demonstrated by ChIP and promoter assays) [100], and the action of EGR1 on MDM2 can therefore be bidirectional, either activating or repressive. However, whether EGR1 drives P2 directly or acts indirectly and cooperatively through MAPK signaling (MAPK3/MAPK8) or similar pathways remains unresolved, and it should be noted that the involvement of EGR1 in this study is based on an inference of activity from downstream targets by the IPA Upstream analysis.

Note that EGR1 mRNA expression varied widely among patients (Supplementary Figure S8 and Supplementary Table S18). Among the P2 patients, expression in HCT27 was higher on the tissue-2 side in all comparisons (mean linear fold change −1.7), whereas in HCT64 expression leaned toward tissue 2 but fluctuated widely depending on the comparison partner, and in HCT33, conversely, expression was higher on the tissue-1 side. Because EGR1 is an immediate-early gene and its absolute mRNA level is inherently unstable, in this study we evaluated the involvement of EGR1 not by its expression level itself but by inferring its activation state from downstream targets with the IPA Upstream analysis. Whether EGR1 contributes to P2 induction was not examined; the differences between patients are compatible with other P2-inducing inputs, such as wild-type p53 itself or the MAPK stress pathway, being relatively predominant in HCT33 and other patients. This is consistent with the mechanistic hypothesis of this study (Section 5.2 of this Note) that P2 promoter activation is controlled not by a single factor but by the integration of multiple stress-response inputs. Similar patient-specific profiles were also observed for other key genes (Supplementary Figure S8); this is an observation in three patients (Section 3.1.3).

##### 3.3 IPA Disease & Bio Functions: simultaneous accumulation of predicted chromosome misalignment and damage, apoptosis, and cell death

In the IPA Disease & Bio Functions analysis, the following biological processes were identified as functional consequences of tissue 2 (Supplementary Table S8: details of IPA Disease & Bio Functions). Misalignment of chromosomes, Abnormal morphology of Nucleus, Cell death of tumor cell lines, Apoptosis, Necrosis, DNA damage, Senescence of tumor cell lines [165], Formation of γH2AX nuclear focus (written as "Formation of gamma H2AX nuclear focus" in Supplementary Table S8, following the official IPA term name), Damage of chromosomes, Autophagy, Aneuploidy, Chromosomal instability, Angiogenesis, Invasion of tumor, Invasion of tumor cells, and Vasculogenesis occupied the top ranks. Of these, Autophagy (median z = −1.05, sign consistency 78%), Aneuploidy (−1.87, 100%), Angiogenesis (−1.03, 79%), Invasion of tumor (−1.66, 93%), Invasion of tumor cells (−1.57, 88%), and Vasculogenesis (−0.76, 75%) had effect sizes below the threshold (sub-threshold). "Misalignment of chromosomes" ranked first in tissue 2, but the predictions of Misalignment of chromosomes, Damage of chromosomes, Aneuploidy, Chromosomal instability, and related terms are likely inferred from the lower expression in tissue 2 of mitotic and checkpoint genes that are more highly expressed in tissue 1 (AURKB, PLK1, and others) and do not indicate chromosomal instability in tissue 2. The expression features of chromosomally unstable tumors instead align with the tissue-1 (P1) side (Section 3.1.4).

The accumulation of Formation of γH2AX nuclear focus is likewise an inference from expression and does not measure the frequency of DNA double-strand breaks (DSBs). The simultaneous accumulation of the Senescence, Apoptosis, and Necrosis signals indicates that both the cellular senescence and apoptosis pathways are activated. Part of this apoptotic signaling may be offset by MDM2 P2-dependent suppression of p53 and may coexist with the increased expression of EMT-, invasion-, and metastasis-related genes (Section 3.3.1) (see Section 5.2 of this Note for details). Note that "Epithelial-mesenchymal transition" in the IPA Disease & Bio Functions has a median z of −0.32 and a sign consistency of 61%, and thus satisfies neither of the adoption criteria of this study (|median z| ≥ 2 and sign consistency of 75% or higher). The basis on which this study discusses EMT is not the IPA activation prediction but the EMT score and the direction of expression of EMT-related genes (Section 3.1.2, Supplementary Tables S1, S13, and S15). "Invasion of tumor" also has an effect size below the threshold (median z = −1.66) and is treated as a tendency based on its sign consistency (92.6%).

##### 3.4 IPA Tox Function: clinical-chemistry indicators of necrotic cell breakdown through elevated ALP (AST, ALT, and LDH in the same direction below threshold)

In the Tox Function analysis of tissue 2, Increased Levels of Alkaline Phosphatase (ALP; median z = −2.12, sign consistency 94%, first-half Rank3 and second-half Rank2) was detected as the top-ranked term in the tissue-2 direction and met the adoption criteria (sign consistency of 75% or higher and |median z| ≥ 2). AST (−1.44, 97%), ALT (−1.45, 94%), and LDH (−1.50, 87%) were also consistent in the same direction, but their effect sizes were all below the threshold; we describe these together as a clinical-chemistry profile of necrotic cell breakdown (Section 5.2 of this Note and Supplementary Table S9; details in Supplementary Results 3 of the Supplementary Information).

##### 3.5 EnrichR GO analysis: accumulation of extracellular vesicles, secretory granule structures, and endopeptidase inhibitors

The EnrichR GO analysis confirmed enrichment of secretory and granular cell structures and of protease inhibitor activity, represented by extracellular vesicle, exosome, and secretory granule lumen (CC; top-ranked, adj. P ≈ 0) and by Endopeptidase Inhibitor Activity (MF; adj. P ≈ 0) (the full list of enriched terms is given in Supplementary Results 4 of the Supplementary Information).

### 4 Common to tissues 1 and 2: IPA integrated analyses

##### 4.1 IPA Regulator Effects analysis

In the IPA Regulator Effects analysis, the highest-scoring causal cascade (Rank1, Consistency Score 25.93) ran from upstream regulators such as EGR1, BMP4, and HIF1A, through targets such as TP53 and VEGFA, to elevation of ALP, suggesting that P2 usage, a readout of p53 activity, and the elevation of ALP co-vary in these specimens, possibly under shared upstream regulators that include EGR1 (a transcription factor that can regulate MDM2 expression in a context-dependent manner [100]); this is a computational cascade, and a causal role of EGR1 was not tested. The Rank2 cascade (Score 25.02) showed that mitotic regulators such as ANLN and the AURK family bidirectionally regulate MDM2, TP53, CDKN1A, BAX, and others. Across all 2,442 entries in Supplementary Table S32, we identified a group of recurrent upstream regulators—11 factors that appeared repeatedly, including ANLN, AURK, CKAP2L, Eldr, LIN9, NEDD8, and PPM1A (details of all cascades [Rank1/2/10 and others] are given in Supplementary Results 5 of the Supplementary Information; discussion in Section 4.2 of the main text). Note that, owing to constraints on computational load, this Regulator Effects analysis is based on the integrated results of the first 8 of the 33 analyses (analyses 49–54, 59, and 63) (see Section 1.1 of this Note).

##### 4.2 IPA Networks analysis

In the IPA Networks analysis (analysis 49, Network 3, Score 37; functions: Cancer/Cell Cycle/RNA Post-Transcriptional Modification), MDM2 was confirmed to coexist in the same protein–protein interaction module with FBXW7 (the E3 ligase responsible for MDM2 degradation), CSNK2A1/2 (CK2; phosphorylation control of the MDM2–p53 interaction), XPO1/CRM1 (responsible for nuclear export of MDM2), LBR (a nuclear-envelope anchoring protein), MCM2 (a DNA replication helicase), and NOLC1 (a nucleolar phosphoprotein) (Supplementary Table S19: IPA Networks; see also Supplementary Figure S14). This network structure indicates that the production, phosphorylation, nuclear export, and degradation of MDM2 are regulated in an integrated manner within a single functional module. The repeated appearance of RNA Post-Transcriptional Modification across all analyses as one of the most frequent network function categories suggests that the differences between the two groups include broad changes in RNA regulation beyond MDM2 splicing. Cell Death and Survival likewise appeared across all analyses as the most frequent network function category, and all network analyses indicated that the regulation of cell death and survival is one of the principal biological differences between the two groups. Viewed quantitatively, among the top-scoring networks the highest score (Score 42) was obtained in analysis 50 (core function: Cell Death and Survival) and analysis 52 (Developmental Disorder/Cancer), and the next highest (Score 40) in analysis 50 and analysis 54 (Cancer/Cellular Compromise, Metabolic Disease). That the top-scoring network has Cell Death and Survival as its core function corroborates the frequency-based tendency described above from the quantitative side as well and suggests that the antagonism between p53-dependent cell death and proliferation/survival signaling is a prominent feature of the network structure.

##### 4.3 IPA Graphical Summary (tissue 1: hub-molecule structure of the autonomous-proliferation type)

In this study, we integrated the top-ranked factors from each module—Canonical Pathway analysis, Upstream Analysis, Disease and Bio Functions, Tox Function, IPA Regulator Effects analysis, and IPA Networks analysis—and extracted the hub molecules that appeared in common across multiple analyses. In tissue 1 (P1-dominant, autonomously proliferative), the following central hub molecules were identified. The overall picture, drawn with Cytoscape by integrating the data from the 33 analyses, is shown in Supplementary Figure S14 (the full lists of hub molecules and relationships are given in Supplementary Tables S20 and S21).

In the IPA Graphical Summary, 32 factors including FOXM1, MYC, MYBL2, E2F1/E2F2/E2F3, AURKB, PLK1, MDM4, and VHL were located at the central hubs (the full list is given in Supplementary Table S20). This is consistent with a positive feedback loop centered on the FOXM1–MYC–E2F axis and with cell-cycle drive that depends little on external proliferative stimuli (an inference from IPA predictions, not a functional test) (for the role of FoxM1 in executing the mitotic program, see reference [166]). Activation of VHL suggests regulation of the HIF pathway and possibly a capacity to adapt to hypoxic environments. Furthermore, the RB-family pocket proteins RBL1 (p107) and RBL2 (p130) were identified as inactivated factors in tissue 1, consistent with autonomous driving of the cell cycle through derepression of the E2F transcription factors (E2F1/E2F2/E2F3). This stands in clear contrast to tissue 2, in which RBL1/RBL2 were identified as activated factors (Section 4.4 of this Note; constituting the TP53–CDKN1A–RBL1/RBL2 cell-cycle arrest hub) consistent with suppressive control of the cell cycle, suggesting that the RB–E2F axis is regulated in opposite directions in the two types (IPA-predicted activation states) (tissue 1: RBL1/RBL2 inactivation → E2F derepression → proliferation; tissue 2: RBL1/RBL2 activation → E2F repression → cell-cycle arrest).

##### 4.4 IPA Graphical Summary (tissue 2: antagonistic hub structure of p53-induced apoptotic signaling and MDM2-dependent survival signaling)

The IPA Graphical Summary of tissue 2 (P2-dominant, environment-adaptive) showed the following hub molecules in the same integrated hub network (Supplementary Figure S14). In the IPA Graphical Summary, 33 factors including TP53, CDKN1A, RBL1, RBL2, IL1B, TGFB1, EGF, TNF, NUPR1, and AKT1 were located at hubs, with FGF2 and AGT also present as lower-degree hubs (degrees 5 and 2, respectively; the full list with degrees is given in Supplementary Table S20; AURKA and HELLS appear as hubs on the tissue-2 network even though their direction of expression is on the tissue-1 side). The configuration in which the cell-cycle arrest factors TP53–CDKN1A–RBL1/RBL2 occupy the core of the hub while the survival and proliferation signals AKT1, EGF, and FGF2 also coexist at the hub indicates a state of dynamic equilibrium in which p53-dependent apoptosis-inducing signaling and AKT/EGF-mediated survival signaling antagonize each other. The presence of AGT (angiotensinogen) at a hub, together with REN, which ranked second in the gene list, is consistent with expression of components of the renin–angiotensin system in tissue 2 (the function of the system was not tested).

### 5 Discussion

#### 5.1 Biological interpretation of tissue 1 (P1-dominant)

In tissue-1 cases in which p53 is inactive, through TP53 mutation or p53 inactivity in the wild type (TP53 mutation is found in most of the P1 group but is not obligatory: 15 of the 17 P1-dominant patients with determined status were mutant [HCT83-4LM assigned to HCT83], and p53 was inactive in both specimens of the wild-type patient HCT59; Sections 3.1.3 and 3.2.1), the P2 promoter would not be induced, and MDM2 would therefore be stably expressed at a basal level from the constitutive P1 promoter. MDM2 and MDM4 (MDMX) have been reported to participate in the control of the cell cycle and proliferation not only through negative regulation of p53 but also in a p53-independent manner [4]. In light of this finding, the steady expression of MDM2 from P1 in tissue 1 can be understood as a state that acts in concert with the autonomous proliferation program driven by the MYC/FOXM1/E2F axis (Sections 2.1 and 2.2 of this Note). However, the data from this study show only that MDM2 expression co-varies with proliferation-related gene sets on the tissue-1 side; they do not demonstrate a functional causal relationship in which MDM2 regulates proliferative signaling.

P1 organoids display the features of a tumor that proliferates autonomously while retaining intestinal identity. The top-ranked Canonical Pathways (pre-mRNA and rRNA processing, cell-cycle checkpoints, and DNA synthesis and repair) and the top-ranked Upstream factors (MYC, Eldr, CEBPB, RABL6, FOXM1, E2F1–3, AURKB, and PLK1) are together consistent with a largely autonomous cell-cycle-driving circuit that depends little on external proliferative stimuli (ranks and effect sizes are given in Sections 2.1 and 2.2 of this Note, Table 3, and Supplementary Table S7). This is consistent with the features of CMS2 (canonical/WNT type) colorectal cancer [2,167]. In the expression data of this cohort, the WNT/β-catenin pathway [151] presents a divided picture (the mean linear FC and sign consistency rate of each gene are given in Supplementary Table S27). The WNT targets that constitute the stem cell program at the base of the intestinal crypt (LGR5, OLFM4, EPHB2, CD44, and ASCL2) were all higher on the tissue-1 side [91,92], consistent with P1/CMS2 retaining intestinal stem cell identity (when this study states that "P1 corresponds to CMS2," the basis is the score of the differentiation core, which excludes the proliferation module, together with gene-level enrichment; the highest overall score is for CMS1, but its difference from CMS2 is not significant; Section 5.2 of this Note). In contrast, the activation and negative-feedback targets of canonical WNT (AXIN2, NKD1, APCDD1, TNFRSF19/TROY, and BMP4), as well as β-catenin (CTNNB1), TCF7L2, and APC itself, were all higher on the tissue-2 side (sign consistency 79–97%). That AXIN2 and NKD1, the standard readouts of canonical WNT activity, are higher on the tissue-2 side indicates that the transcriptional output of the WNT pathway is, if anything, stronger in the P2-dominant group, and is mechanistically consistent with ligand-dependent WNT activation, as seen with RSPO2/3 fusions, which occur in APC-wild-type colorectal cancers [168], and with RNF43-inactivating mutations, which are frequent in MSI colorectal cancers [169]. The two tissue types are therefore interpreted as exhibiting distinct WNT modes—an intestinal stem cell-type WNT target program on a background of APC loss of function in P1/CMS2, and a strong, ligand-dependent transcriptional WNT output in P2/serrated (MSI-like at the signature level). Of note, MYC, a direct target of canonical WNT [170] that drives the proliferation of P1, is of indeterminate direction at the mRNA level (Supplementary Table S27), whereas it shows strong activation in the IPA Upstream analysis (median z = +5.76), suggesting regulation at the level of activity rather than transcript abundance. The high expression of OLFM4 suggests maintenance of the intestinal stem cell program [91], and the increased expression of HNF4A (Additional file 1, Section R-20) indicates maintenance of the intestinal transcription factor network. Note that this interpretation is based on expression-level findings; in the IPA Upstream analysis, the direction of HNF4A activation was undetermined (sign consistency 52%), and the activation of CDX1 was also sub-threshold (Section 2.2 of this Note). The enrichment of terms related to apico-basal polarity specific to the intestinal absorptive epithelium, amino acid transport, and DNA replication in the EnrichR GO analysis (Section 2.5 of this Note and Supplementary Table S31) reveals a dual nature in which proliferation and intestinal differentiation identity coexist. Moreover, the Rank1 Canonical Pathway, "Processing of Capped Intron-Containing Pre-mRNA," is consistent with a high volume of transcriptional activity and with high activity of the splicing machinery, and the accumulation of ribosome biogenesis terms (Ranks 4 and 10) corresponds mechanistically to the high expression of ribosome biogenesis in Type1 PDOs reported by Okamoto et al. [12] (Section 2.1 of this Note) and to the high expression of TERT. The coordinated activation of G2/M genes by the MuvB–FOXM1 axis, inferred by the IPA Regulator Effects analysis (Section 4.1 of this Note and Supplementary Results 5 of the Supplementary Information), is a candidate molecular mechanism underlying the highly efficient proliferation of the P1-dominant group.

#### 5.2 Biological interpretation of tissue 2 (P2-dominant): a dynamic equilibrium between wild-type p53-induced apoptotic signaling and MDM2-dependent survival signaling

The significance of the predominant activity of the MDM2 P2 promoter in tissue 2 extends beyond mere suppression of p53. As Phelps et al. showed in Cancer Research (2003) [9], the P2 promoter can be induced not only by p53 but also through multiple transcription factor response elements. In the Upstream Analysis, MAPK3 was detected as an activating factor in the tissue-2 direction that met the adoption criteria (median z = −3.51, sign consistency 90%), and EGR1 (−1.95, 91%) and SRC (−1.85, 93%) were also consistent in direction but had effect sizes below the adoption threshold (Section 3.2 of this Note and Supplementary Table S7). All of these are consistent with activation of the RAS/MAPK pathway and could act in concert to activate the P2 promoter persistently. However, in nutlin-treated cells P2 was induced in a p53-dependent manner and was essentially not induced in cells lacking p53 (Section 3.2.1), so these inputs are considered to act in the presence of active p53. The production of P2-derived MDM2 would establish a dynamic suppression cycle in which wild-type p53 is repeatedly activated and degraded, and this apoptosis evasion could support the survival of cancer cells during a complex process of lineage conversion in which TGFβ/SMAD3-mediated partial EMT and acquisition of gastric metaplasia coexist (the causal chain linking them was not verified in this study; Section 4.3 of the main text).

P2 organoids are interpreted as being in a dynamic equilibrium in which, in response to sustained activation of wild-type TP53 (the top-ranked factor in the Upstream analysis), the MDM2 P2 promoter is induced by p53, which would suppress p53-dependent apoptosis. The correspondence between this picture and the environment of immune activation and chronic inflammation that Guinney et al. described as characteristic of CMS1 (the MSI immune type) [2] is limited to a signature-level resemblance in the immune module. P2 dominance is not equivalent to microsatellite instability: in TCGA-COAD/READ, P2_index did not differ among the CIN, GS and MSI subtypes and was not associated with MSI status (Sections 3.1.4, 3.2.2 and 4.6). The seemingly contradictory configuration in which the p53 target genes (CDKN1A, DDB2, FAS, and DR5), MDM2 itself, and additionally AKT1 and EGF signaling simultaneously occupy hubs of the Graphical Summary is interpreted as antagonism between p53-induced apoptotic signaling and MDM2/AKT/EGF-dependent survival signaling (an inference from bulk expression and IPA prediction; co-occurrence within the same cells was not tested). As Purvis et al. showed [171], the amplitude and period of these p53 dynamics determine whether the cell lives or dies.

Gastric-type and Paneth-cell-type metaplasia (CTSE, REN, TFF1/3, DEFA5/DEFA6, and MUC5B/5AC) indicate a composite metaplasia-like phenotype comprising loss of colorectal identity and simultaneous acquisition of multiple ectopic differentiation traits (the higher expression of transcripts of the imprinted 15q11-q13 locus, including the SNORD116 cluster, is analyzed in the companion study on the 15q11-q13 locus [T. Tsukui, R. Yao, and K. Tsuda, unpublished observations]). The ALP-elevation signal detected at the top of the Tox Function analysis (Section 3.4 of this Note) is hypothesized to reflect necrotic cell breakdown accompanying the antagonism between p53-induced cell death and MDM2-dependent survival signaling within P2 tumors (enzyme levels were not measured in this study). AST, ALT, and LDH are in the same direction, but their effect sizes are all below the adoption threshold (Section 3.4 of this Note and Supplementary Table S9). IPA Regulator Effects Rank1 (EGR1/HIF1A/NF-κB → TP53/VEGFA → ALP elevation; Consistency Score 25.93) provides a computational hypothesis for this ALP signal. In the EnrichR GO CC analysis, Extracellular Vesicle and Exosome were enriched as the top two terms, with by far the highest significance, suggesting increased production of tumor-derived extracellular vesicles [154] (vesicle production was not measured). The enrichment of Golgi Lumen, Tertiary Granule Lumen, and Secretory Granule Lumen suggests the emergence of secretory features characteristic of gastric chief cells and pancreatic acinar cells. In the GO MF analysis, Endopeptidase Inhibitor Activity (15 factors including SERPINA1) and Death Receptor Activity (FAS, TNFRSF10B, and others) occupied the top ranks, suggesting high expression of serine protease inhibitors and death receptors (the adj. P of each term is given in Supplementary Results 4 of the Supplementary Information; summary in Section 3.5 of this Note and Supplementary Table S43). We also explored the relationship with the mutational footprint of exposure to an exogenous mutagen (Supplementary Figures S11 and S12, Supplementary Table S42; Section 2, Methods). In the Nunes cohort (WGS plus RNA-seq, n = 1,051) [82], the burden of the colibactin-associated signatures (SBS88 and ID18) was only slightly shifted toward the P2-dominant side (correlations were weak; ID18: overall ρ = −0.104, FDR = 0.0041; MSS only, ρ = −0.156, FDR = 3.8×10⁻⁵; SBS88: overall ρ = −0.085, FDR = 0.013; MSS only, ρ = −0.121, FDR = 7.6×10⁻⁴), and ID18 remained significant after covariate adjustment (partial Spearman ρ = −0.084, P = 0.0067; SBS88 was borderline, P = 0.053). Replication in TCGA-COAD/READ (mainly WES, n = 374) was partial: a significant association in the same direction was found only between P2_index and ID18 (ρ = +0.143, FDR = 0.044; high P2_index = the P2 side), whereas no significant association was found on the expression GSVA axis, and SBS88 was not significant on either axis (Supplementary Figure S12). This analysis therefore remains a cross-sectional, exploratory association, and it should be noted that replication in TCGA depends on how the axis is defined (limitations are given in Section 4.6).

To summarize the positioning as molecular subtypes, the P1-dominant group corresponds to CMS2 (Canonical/CIN type)—the GSVA score of the differentiation core excluding the proliferation module was highest in CMS2 and significantly higher than in CMS1, whereas the whole-signature score was highest in CMS1 but did not differ significantly from CMS2 (Section 3.3.2, Supplementary Table S34); at the gene level, with the TCGA-derived class markers (reference set B) the tissue-1 side DEGs were specifically enriched in CMS2 markers (OR = 3.92), with 0 genes overlapping CMS4 markers, whereas with the CMScaller templates (reference set A) they overlapped the CMS1–CMS3 templates to a similar degree (OR = 3.09–3.77) (Section 3.3.3, Supplementary Table S28)—whereas the class to which the P2-dominant group is assigned differs depending on the layer evaluated. At the score (pathway) level, CMS1 and CMS3 were equivalent, with no significant difference between them (Section 3.3.2). The composition of this equivalence becomes clear when it is decomposed into functional modules. The immune checkpoint module (6 genes) stands out in CMS1 (Section 3.3.2, Supplementary Table S34), and this corresponds to the core features of CMS1—wild-type TP53, MSI, and immune activation—and to the dMMR/pMMR difference in the independent cohort GSE39582 (full P2 score P = 1.14×10⁻¹⁷; the difference remains with the immune checkpoint genes excluded, P = 6.59×10⁻¹⁰, the primary score in the main text). In contrast, for the core excluding the immune and lncRNA modules (metaplasia plus p53 targets, 21 genes), CMS3 scored highest and significantly higher than CMS1, and for the gastric/Paneth metaplasia module alone the difference was even larger (all in Section 3.3.2, Supplementary Table S34). That is, the P2 score appears CMS1-like because of the immune axis, whereas the lineage and differentiation traits correspond to CMS3.

However, in the gene-level overlap analysis, the tissue-2 side DEGs were significantly enriched in CMS1 markers (OR = 2.63), whereas the enrichment in CMS3 (metabolic) markers was stronger (OR = 5.91; Section 3.3.3, Supplementary Table S28). The 28 genes overlapping on the CMS3 side are dominated by secretory, mucinous, and metabolic systems such as TFF1, TFF3, and CTSE (all 28 genes are listed in Supplementary Table S28, reference set B, top 200) and correspond to the phenotype of gastric metaplasia and lineage plasticity shown in this study. This tendency was consistent whichever of the two independent families of reference sets was used for evaluation (the CMS templates of CMScaller, and class-specific markers derived empirically from TCGA-COAD/READ), and enrichment in CMS3 was the strongest of the four classes (OR = 5.91–18.21; the strongest under any of the top-100, top-200, and top-500 reference-set definitions; Supplementary Table S28). To confirm that this enrichment is not a circularity arising from the signature definitions of this study itself, we recalculated after removing from the query side the 29 genes of the P2 signature and a 26-gene panel of gastric-metaplasia, Paneth-cell-metaplasia and intestinal-identity markers (gastric metaplasia: MUC5AC, MUC6, TFF1, TFF2, GKN1, GKN2, ANXA10, AQP5, CTSE, LGALS4, VSIG1, CLDN18, MUC5B and PGC; Paneth-cell metaplasia: DEFA5, DEFA6, MMP7, REG4 and PLA2G2A; intestinal identity: CDX2, VIL1, MS4A12, SLC26A3, MUC2, HNF4A and SATB2; the same panel is used as a sensitivity panel in the companion study of the 15q11-q13 locus, T. Tsukui, R. Yao, and K. Tsuda, unpublished observations), 47 genes in all after removing duplicates; the enrichment in CMS3 was hardly attenuated (OR = 5.91 → 5.30, BH-FDR = 2.0×10⁻¹¹ → 2.0×10⁻⁹), and the same held for CMS1 (OR = 2.63 → 2.31, BH-FDR = 3.2×10⁻³ → 1.8×10⁻²). The exclusion was repeated with a narrower 17-gene panel of the same marker classes (AGR2, AQP5, CLDN18, GIF, GKN2 and TFF2; ANG, MMP7, PLA2G2A, REG4 and RNASE4; CA4, CDX2, FABP2, MS4A12, SATB2 and VIL1); interchanging the two does not change the conclusion. Of the 877 tissue-2 side DEGs, 11 genes are contained in the 26-gene panel, whereas only 2 genes, AQP5 and MMP7, are contained in the 17-gene panel; when the latter is substituted, the enrichment is OR = 5.29 for CMS3 and OR = 2.31 for CMS1, essentially identical to the values above (5.30 and 2.31). That is, exclusion by the 26-gene panel removes a larger number of genes and is the more conservative test. Moreover, of the 6 genes of the 17-gene panel that are not contained in the 26-gene panel (AGR2, GIF, ANG, RNASE4, FABP2, CA4), only FABP2 satisfies the DEG criteria of this study, and because FABP2 belongs to the tissue-1 side DEGs (1,013 genes), it cannot enter the present sensitivity analysis, which targets the tissue-2 side DEGs. Therefore, the circularity check in this section does not depend on which of the two panels is used. Of the 28 genes overlapping on the CMS3 side, 25 genes (89%) are not used in any signature of this study, and the CMS markers contained in the P2 signature are nearly balanced, with 2 genes for CMS1 and 3 genes for CMS3 (the 6 immune checkpoint molecules are not among the genes overlapping on the CMS1 side). That is, neither the score-level assignment to CMS1 nor the gene-level assignment to CMS3 is an artifact arising from the definition of the signatures.

On the other hand, the P2-dominant group simultaneously shows TGFβ1/SMAD3-associated EMT, ECM remodeling, and invasion-related pathways (the difference in EMT score between P2-dominant and P1-dominant specimens was not significant, P = 0.079; the score follows the sampling site rather than the two poles and is treated as a separate axis of progression, Section 3.1.2 of the main text) (EMT and Tumor Microenvironment in the IPA Canonical Pathway analysis; the extracellular vesicle and protease groups in the GO analysis), and these also overlap with features of the mesenchymal CMS4 (consistent in direction with the finding of Calon et al. [156] that stroma-derived expression signatures define poor-prognosis subtypes). Therefore, rather than converging on a single CMS as the P1-dominant group does, the P2-dominant group is positioned as a composite subtype in which two axes map onto separate CMS classes: the lineage and differentiation traits (gastric metaplasia, secretion, and metabolism) correspond to CMS3 (metabolic), and the immune-evasion traits correspond to CMS1 (MSI immune). Because two readouts—module-specific scores and gene-level enrichment—both support CMS3, this assignment does not depend on a single analysis. The CMS4-like mesenchymal and stromal traits appear only in the pathway-level findings, and at the gene level no enrichment in CMS4 markers is observed (OR = 0.32 with the TCGA-derived markers, 2 overlapping genes). This dual assignment is a substantive one arising from the difference in the layer measured, and there is no need to treat either one as primary. This non-singularity is consistent with the biological picture of lineage plasticity proposed in this study (acquisition of mesenchymal traits through partial EMT in addition to gastric-type and Paneth-cell-type metaplasia), and suggests the existence of dynamic, transitional tumor states that cannot be fully captured within a fixed CMS framework.

#### 5.3 Physical properties and drug accessibility

The contrast in physical properties between the “dense solid” of tissue 1 and the “highly viscoelastic gel” of tissue 2 (both inferred from gene expression rather than measured; see the note at the end of this section; for the jamming transition of tumors, see reference [150]) is a difference that bears directly on therapeutic efficacy through the relationship between the physical characteristics of tumors and drug accessibility that Jain et al. showed in Annual Review of Biomedical Engineering (2014) [127]. This difference in physical properties is supported by the contrast in cellular architecture revealed by the EnrichR GO CC analysis. Basolateral/External Side of Apical Plasma Membrane (adj. P = 0.013), a polarized structure specific to the intestinal absorptive epithelium, and the CMG complex (the Cdc45–MCM2-7–GINS replicative helicase [152]), both enriched in tissue 1, reflect the maintenance of a densely arrayed epithelial sheet structure equipped with epithelial apico-basal polarity, tight junctions, and a brush border. Such a polarized, densely arrayed epithelium may be directionally consistent with a “dense solid” state and with reduced drug accessibility due to solid stress (the epithelial marker CDH1 showed no clear difference between the groups [Supplementary Table S15], and the adhesion and cytoskeleton GO terms were instead enriched on the cystic–mucinous side; Section 3.1.4). In contrast, the terms most highly enriched in tissue 2—Extracellular Vesicle/Exosome (adj. P ≈ 0), Golgi Lumen (adj. P = 0.0005), and Tertiary/Secretory Granule Lumen (adj. P = 0.0007–0.005)—indicate a hypersecretory cellular architecture characterized by massive vesicle and granule trafficking and extracellular release via the Golgi apparatus, secretory granules, and extracellular vesicles. The vigorous secretion of mucins (MUC5B/MUC5AC/MUC17) and extracellular vesicles may form a hydrated, viscoelastic gel-like extracellular environment and high interstitial fluid pressure (IFP), and could constitute the cellular-architectural basis of the inferred “highly viscoelastic gel” state and of a drug-delivery barrier that impedes convection and diffusion. That is, the differences in cellular architecture and vesicle biogenesis captured by GO Cellular Component (Supplementary Table S43) are directionally consistent with the contrasting physical properties inferred for the two tissue types (solid versus gel) and provide a working hypothesis linking the molecular-level findings to the physical properties of the tumor and to drug accessibility. These physical properties are inferred from P1/P2 expression differences, whereas organoid morphology itself follows TP53 status beyond 20q and aneuploidy and correlates only weakly with the MDM2 axis (ρ = 0.33; Section 3.1.4) (the contrast in physical properties is given in Supplementary Table S45, and the details of GO Cellular Component in Supplementary Table S43).

The high IFP of tissue 2 is, as Nia et al. showed in Nature Biomedical Engineering (2016) [128], the most serious barrier that physically impedes the convection and diffusion of drugs. The mucus barrier formed by MUC5B, MUC5AC, and MUC17 could act as a hydrophobic wall and impede the penetration not only of small-molecule compounds but also of large molecules such as antibody–drug conjugates (ADCs) (the mucus barrier was not measured). The possibility that vascular normalization by anti-VEGF therapy (Jain, 2001) [129] lowers IFP and improves the intratumoral penetration of subsequent chemotherapy and immunotherapy is an important perspective in the design of combination strategies against tissue 2. Furthermore, because the hypoxia and acidification in a high-IFP environment continuously activate p53-mediated stress responses, the physical environment of tissue 2 may form a positive feedback mechanism that sustains the feedback loop of P2 promoter activation (a hypothesis; IFP and hypoxia were not measured).

It should be noted that two distinct axes are involved in the physical state of a tumor. The first is the jamming/unjamming transition of cell collectives: as Oswald, Käs, and colleagues discussed [150], cancer cell collectives undergo a phase transition between a solid-like state (jamming; the cells are aggregated, unable to move, and non-invasive) and a fluid-like state (unjamming; the cells flow, migrate collectively, and are invasive), and this transition bears directly on collective invasion and metastatic capacity. The second is the mechanics of the bulk tissue and extracellular matrix (solid stress versus mucinous gel and interstitial fluid pressure), which governs drug accessibility [127,128]. The two groups in this study contrast on both of these axes. Tissue 2 is accompanied by partial EMT via TGFβ/SMAD3 (Sections 3.2 and 5.2 of this Note) and by an invasion program centered on MMP7 (Section 3.1 of this Note), and at the level of the cell collective this may be directionally consistent with a shift toward the fluid-like (unjammed) state [105,150]. However, the difference in EMT score between P2-dominant and P1-dominant specimens was not significant (P = 0.079); the highest EMT score was in the metastatic specimen HCT38-3LM (4.94), and HCT67-4LMR, the P2-dominant recurrent liver metastasis of HCT67, also scored high (+4.50), so confounding by sampling site is possible (Sections 3.1.2 and 4.6). Tissue 1, in contrast, retains a polarized, densely arrayed epithelium and as a cell collective is directionally consistent with a state close to the solid-like (jammed) state. Separately from this, on the axis of bulk mechanics, tissue 1 presents solid stress and tissue 2 is inferred to present a highly viscoelastic mucus gel and high IFP, both of which restrict drug accessibility, although in different modes. That is, “gel” (the mechanics of the extracellular environment) and “fluid-like” (the fluidity of the cell collective) are concepts at different hierarchical levels, and once the two are distinguished, tissue 2 can be described without contradiction as having a dual physical character: it may be close to fluid-like as a cell collective, while its extracellular environment is a highly viscoelastic gel. Taken together, as a working hypothesis, the P1/P2 expression programs may be directionally consistent not only with proliferation and differentiation but also with the physical state of the cell collective (jamming ↔ unjamming). However, observed organoid morphology follows TP53 status beyond expression-inferred 20q gain and aneuploidy and correlates only weakly with the MDM2 (P1/P2) axis (ρ = 0.33; Section 3.1.4); we therefore do not regard differences in physical properties or morphology as consequences of differences in P1/P2 promoter usage. Note that this finding is an interpretation based on expression programs, and direct rheological measurement of the fluidity of the cell collective itself remains a subject for future work.

When viewed in light of the recent framework in which the mechanical microenvironment regulates chromatin state and transcription [106], the above difference in physical properties (tissue 1 = dense solid, tissue 2 = highly viscoelastic gel) also appears as a consistent pattern at the level of transcription factor activity. The transcription factors that showed activation on the tissue-2 side in this study included many known mechanosensitive transcription factors. Specifically, TGFβ–SMAD (TGFB1/2/3, SMAD2/3/4), which is activated by release from the ECM through traction force, Wnt/β-catenin (CTNNB1), which responds to tissue compression, and AP-1 (FOS: median z = −2.33, sign consistency 88%; JUNB: −2.36, 96%) were identified in the IPA Upstream analysis as activating factors on the tissue-2 side that met the adoption criteria. NF-κB (NFkB complex: −1.51, 88%; RELA: −1.63, 88%) and EGR1 (−1.95, 91%), which respond to cell stretching, were also consistent in the same direction, but their effect sizes were below the adoption threshold (Section 3.3.1; Sections 3.2 and 5.2 of this Note). Furthermore, the two major mechanosensitive transcriptional modules, SRF–MRTF (MRTFB) and YAP/TAZ–TEAD (TEAD2/TEAD3, together with YAP/TAZ targets such as ANKRD1), were also activated on the tissue-2 side (Supplementary Table S29). This convergence suggests that the partial EMT and metaplasia program of tissue 2 is supported by a mechanosensitive transcriptional network consistent with a soft, gel-like mechanical microenvironment, and is consistent with (but does not independently test) the physical-property interpretation of this study.

By contrast, the metabolic axis [106,172]—in which reduced actomyosin tension in the soft ECM of the metastatic site activates NRF2 via mitochondrial fission and SREBP via the Golgi, conferring chemotherapy resistance through reprogramming of antioxidant and lipid metabolism—was not supported in tissue 2 of the present cohort. NRF2 (NFE2L2) was instead an activated upstream regulator on the tissue-1 side (IPA activation z=+2.49), and the lipogenic enzymes (FASN, ACACA) were also more highly expressed on the tissue-1 side (Supplementary Table S29). This negative finding is consistent with the interpretation of this study that any chemotherapy resistance of tissue-2-type tumors would arise mainly not from that metabolic axis but from the physical drug-accessibility barrier of the mucus gel [138] (inferred from gel-forming mucin expression) and from the dynamic equilibrium between wild-type p53-induced apoptosis and MDM2 P2-dependent survival signaling (Section 5.2 of this Note; Additional file 1, Section R-18); neither mechanism was tested directly.

Note that, because tissue elasticity and tension were not measured directly in the organoid system of this study, the above correspondence between the transcriptional and chromatin mechanisms and the physical-property states does not establish mechanical causation; it is convergent corroborating evidence showing that the inferred physical-property states and the molecular mechanisms are mutually consistent.

### References not cited in the main text

Numbering continues from the reference list of the main text; only the references that are cited in this Additional file and not in the main text are listed. Because these numbers are shared with Additional files 1 and 12, a number not cited in this Note (155) is skipped.
