## Additional files for "TP53 status aligns with morphology and metaplastic programs in colorectal cancer organoids, and MDM2 P2 promoter usage reflects p53 activity": Additional_file_27_Figure_S8.pdf

### Sample-specific behavior of key genes: linear fold change per TAC comparison (table format)

Negative = higher in the tissue-2-side samples; positive = higher in the tissue-1-side samples.
All values are identical to Table S26 (Additional file 26); color is auxiliary (direction only).

| Analysis no. | Tissue-2-side samples (patient of origin) | OLFM4† | EGR1 | MDM2 | CTSE | REN | SNORD116 | TFF1 | DEFA5 | MMP7 |
|---|---|---|---|---|---|---|---|---|---|---|
| 49 | HCT27/HCT64 | -27.16 | -1.5 | -5.99 | -192.72 | -106 | 2.25 | -25.01 | -34.6 | -9.16 |
| 50 | HCT27/HCT64 | -21.34 | -1.72 | -7.88 | -46.62 | -58.95 | 43.49 | -14.75 | -28.93 | -8.85 |
| 51 | HCT27/HCT64 | -44.98 | -2.94 | -9.38 | -63.8 | -33.77 | 22.74 | -6.86 | -1 | -75.44 |
| 52 | HCT27/HCT31/HCT64/HCT71 | 4.02 | -1.35 | -3.04 | -58.5 | -12.56 | 5.66 | -3.57 | -1.02 | -3.1 |
| 53 | HCT64 | 428.65 | -1.5 | -6.03 | -3309.31 | -1129.18 | -48.18 | -449.35 | -783.56 | -10.73 |
| 54 | HCT64 | 119.02 | -3.79 | -4.51 | -2281.99 | -362.88 | -19.63 | -82.28 | -483.53 | -9.13 |
| 59 | HCT27 | 7.56 | -1.01 | -35.72 | -26.77 | -32.91 | -0.87 | 1.2 | 48.41 | -79.38 |
| 63 | HCT67 | 3.17 | 3.17 | -2.59 | -248.74 | 737.01 | -519.74 | -3.95 | 10.28 | -453.54 |
| 67 | HCT33 | -1.2 | 5.13 | -6.71 | -16.4 | -2.31 | -223.55 | 32.98 | -7.2 | -123.45 |
| 68 | HCT33 | -1.12 | 1.96 | -5.64 | -25.86 | -6.84 | -158.54 | 5.34 | -7.43 | -13.91 |
| 69 | HCT33 | 3.13 | 8.75 | -5.5 | -15.72 | -2.63 | -513.78 | 2.06 | 63.98 | -9.09 |
| 70 | HCT33 | 1.53 | 1.7 | -9.42 | -81.89 | -3.14 | -302.9 | 6.26 | 55.85 | -25.3 |
| 76 | HCT38 (deviating) | 2.98 | -8.48 | -5.54 | -3053.54 | -1.42 | -1017.74 | -9.39 | 51.47 | -2689.07 |
| 85 | HCT64 | 270.72 | 1.36 | -7.62 | -508.48 | -43.4 | -43.01 | -1303.22 | -10.47 | -8.74 |
| 90 | HCT64 | 65.06 | -7.08 | -10.28 | -3374.2 | -942.08 | -39.25 | -792.72 | -282.95 | -10.73 |
| 91 | HCT64 | -1.55 | -19.46 | -9.84 | -2814.65 | -2258.41 | -78.06 | -1797.39 | -883.27 | -11.18 |
| 92 | HCT64 | -1.78 | -9.58 | -8.55 | -3219.94 | -2152.69 | -61.69 | -938.38 | -1169.29 | -12.11 |
| 93 | HCT64 | -1.75 | -7.53 | -6.45 | -4447.66 | -1386.27 | -69.77 | -1317.81 | -751.64 | -10.57 |
| 94 | HCT64 | 10.08 | -1.17 | -4.89 | -579.84 | -2671.6 | 1.89 | -205.35 | -794.06 | -6.9 |
| 96 | HCT64 | 2.32 | 16.18 | -10.27 | -846.22 | -376.56 | 10.25 | -523.46 | -4.97 | -6.16 |
| 97 | HCT64 | 3.73 | -1.07 | -6.34 | -2980.5 | -168.84 | 6.56 | -660.69 | -748.55 | -5.18 |
| 98 | HCT64 | 11.56 | 1.76 | -4.75 | -4357.15 | -1882.09 | 7.32 | -789.32 | -790.14 | -5.52 |
| 99 | HCT64 | 77.3 | -1.14 | -3.9 | -1034.13 | -1491.61 | 1.88 | -292.26 | -823.07 | -4.7 |
| 100 | HCT64 | -1.36 | -9.94 | -3.47 | -4728.19 | -3286.66 | -25.19 | -617.11 | -1181.1 | -7.27 |
| 101 | HCT64 | 3753.08 | 4.82 | -1.89 | -4.35 | -2.33 | -46.42 | 1.05 | -220.34 | -5.38 |
| 102 | HCT64 | 1.94 | 40.33 | -3.7 | -120.47 | -4.89 | -36.98 | 1.14 | -68.46 | -8.33 |
| 103 | HCT64 | 8197.67 | -6.17 | -11.4 | -5385.52 | -3831.45 | -64.92 | -56 | -117.01 | -1.98 |
| 104 | HCT64 | 5474.57 | -4.5 | -4.8 | -2922.91 | -3813.6 | -50.71 | -826.38 | -18.65 | -9.93 |
| 105 | HCT64 | 717.72 | 1.69 | -3.81 | -2424.34 | -2795.07 | -50.97 | -1864.75 | -759.03 | -9.25 |
| 106 | HCT64 | 7990.33 | 1.31 | -5.2 | -253.3 | -1971.01 | -50.57 | -80.16 | -1025.81 | -5.42 |
| 107 | HCT64 | 7562.09 | -14.85 | -6.3 | -4875.18 | -4427.14 | -70.68 | -520.09 | -54.1 | -12.99 |
| 108 | HCT64 | 3506.7 | -4.21 | -3.24 | -1947.19 | -3699.45 | -2.89 | -334.74 | -987.58 | -5.57 |
| 109 | HCT64 | 3205.83 | -1.77 | -7.69 | -6473.68 | -4055.61 | -43.07 | -5.72 | -766.65 | -4.77 |

#### Mean linear FC per patient of origin of the tissue-2-side samples (Table S26, sheet "Per-patient summary")

| Gene | HCT27 (n=5) | HCT31 (n=1) | HCT33 (n=4) | HCT38 (n=1) | HCT64 (n=26) | HCT67 (n=1) | HCT71 (n=1) | All 33: mean; % negative |
|---|---|---|---|---|---|---|---|---|
| **OLFM4†** | -16.38 | 4.02 | 0.58 | 2.98 | 1588.56 | 3.17 | 4.02 | +1252.1; 27% |
| **EGR1** | -1.7 | -1.35 | 4.38 | -8.48 | -1.3 | 3.17 | -1.35 | -0.7; 64% |
| **MDM2** | -12.4 | -3.04 | -6.82 | -5.54 | -6.2 | -2.59 | -3.04 | -7.0; 100% |
| **CTSE** | -77.68 | -58.5 | -34.97 | -3053.54 | -2278.88 | -248.74 | -58.5 | -1900.6; 100% |
| **REN** | -48.84 | -12.56 | -3.73 | -1.42 | -1652.47 | 737.02 | -12.56 | -1281.1; 97% |
| **SNORD116** | 14.66 | 5.66 | -299.69 | -1017.74 | -26.92 | -519.74 | 5.66 | -104.2; 73% |
| **TFF1** | -9.8 | -3.57 | 11.66 | -9.39 | -519.43 | -3.95 | -3.57 | -408.2; 79% |
| **DEFA5** | -3.43 | -1.02 | 26.3 | 51.47 | -491.91 | 10.28 | -1.02 | -381.0; 85% |
| **MMP7** | -35.19 | -3.1 | -42.94 | -2689.07 | -10.35 | -453.54 | -3.1 | -111.0; 100% |

Each row of the upper table is one TAC pairwise comparison between organoid samples; FC values are shown per comparison, and the patient of origin identifies the samples on the tissue-2 side of that comparison. †OLFM4 is a tissue-1 marker; the sign of its FC depends on the tissue-1-side samples of each comparison (shown for reference). "All 33" gives the mean FC across the 33 comparisons and the percentage of comparisons with negative FC (higher in the tissue-2-side samples). FC, linear fold change.
