## Additional files for "TP53 status aligns with morphology and metaplastic programs in colorectal cancer organoids, and MDM2 P2 promoter usage reflects p53 activity": Additional_file_30_Figure_S10.pdf

Supplementary Figure S10. GDSC2: sensitivity to MDM2 inhibitors and p53-pathway drugs stratified by TP53 status and MSI status

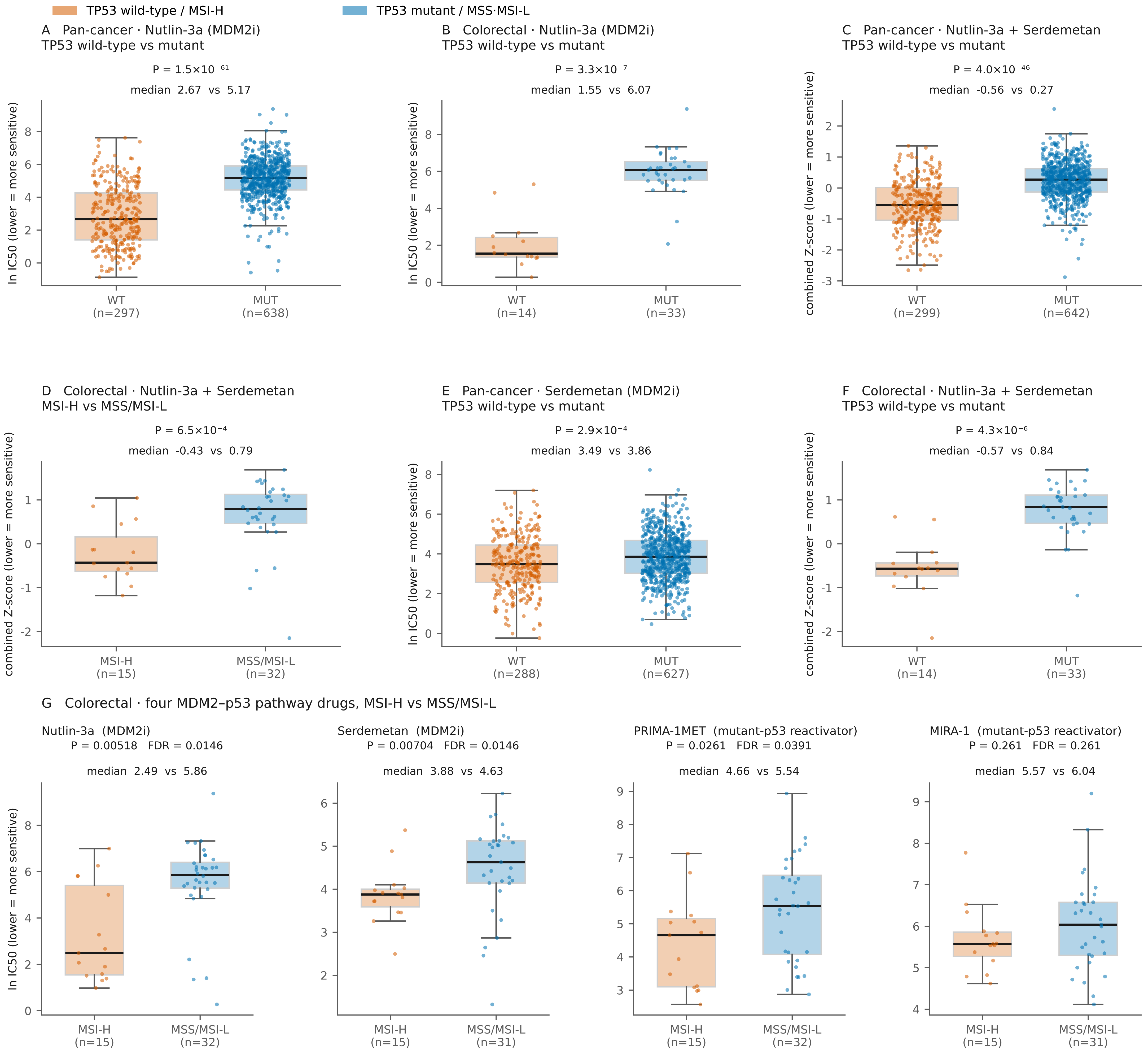


Dose–response: GDSC2 (fitted_dose_response, 24 Jul 2022). TP53 status and lineage: DepMap 24Q4 Public (Model.csv / OmicsSomaticMutations.csv); cell lines without a mutation profile are excluded rather than called wild-type. MSI status: GDSC Cell_Lines_Details. Values are per-cell-line medians over duplicate curves.

Two-sided Mann-Whitney (normal approximation, continuity- and tie-corrected). Panels A-F carry no multiplicity correction across panels; panel G shows the Benjamini-Hochberg FDR computed within the colorectal MSI block of Supplementary Table S38.

Combined Z-score = mean of the GDSC1/GDSC2 Z-scores of Nutlin-3a and Serdemetan (4 measurements per cell line where available). Panels D, F use colorectal lines with a known TP53 status.

PRIMA-1MET and MIRA-1 are mutant-p53 reactivators; greater sensitivity on the MSI-H side (which is enriched for TP53 wild-type) is not the direction their putative mechanism predicts and is reported as an observation only.

All statistics are transcribed from the primary outputs of the deposited code (D2_mdm2i_by_*.csv, S56_D2_integrated.csv); the drawn values are in FigureS10_v6_drawn_values_20260803.csv (Additional file 2).
