## Additional files for "TP53 status aligns with morphology and metaplastic programs in colorectal cancer organoids, and MDM2 P2 promoter usage reflects p53 activity": Additional_file_45_Figure_S9.pdf

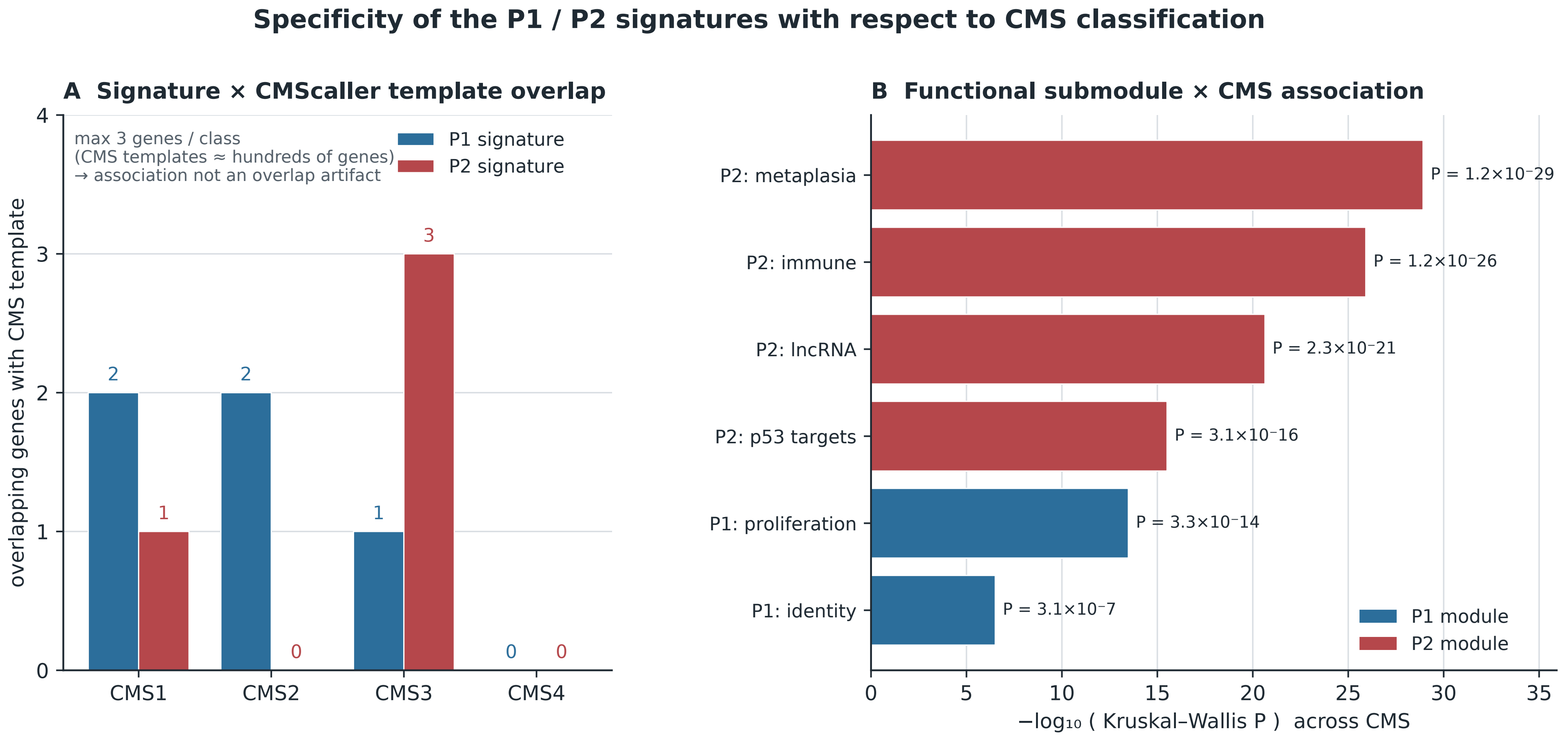


Overlap counts and module association P-values from main-text Section 3.3.2 (Code 8, Additional file 2 → Table_Module_CMS_KW.csv); Kruskal–Wallis P-values are computed at the sample level (n = 563 samples, aliquots averaged within a sample). Each scored functional submodule is independently associated with CMS, and the signature shares ≤3 genes with any CMS template. The P1 identity-TF module (HNF4A only) is not scored: GSVA requires ≥2 genes per set.
