## Additional files for "TP53 status aligns with morphology and metaplastic programs in colorectal cancer organoids, and MDM2 P2 promoter usage reflects p53 activity": Additional_file_61_Figure_S11.pdf

Colibactin mutational-signature burden (SBS88 / ID18) versus the P1−P2 axis (Nunes et al. 2024 cohort, WGS + RNA-seq, n = 1,051)

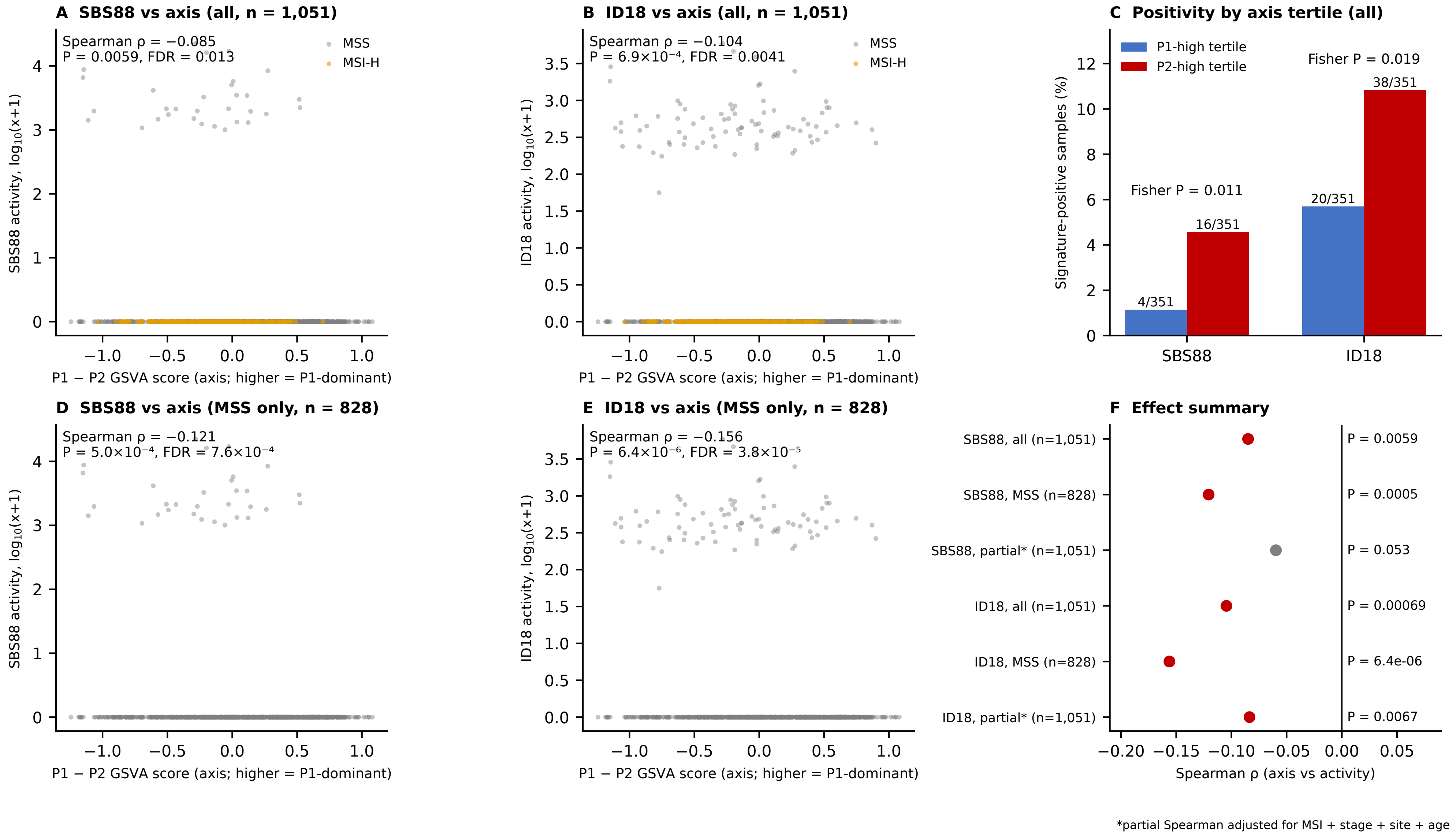
