## Additional files for "TP53 status aligns with morphology and metaplastic programs in colorectal cancer organoids, and MDM2 P2 promoter usage reflects p53 activity": Additional_file_01_Supplementary_Report.docx

Note that ‘tumor purity’ in this manuscript denotes a single measure except in one analysis: in the external cohort validation (Code8) we used the value obtained by converting the ESTIMATE score into an approximate purity with the transformation of Yoshihara 2013 [80] (purity = cos(0.6049872018 + 0.0001467884 × ESTIMATE score)), and the multivariable model of the direct validation of MDM2 P1/P2 (Code21, with the covariate table of R16) used the same approximate purity (median 0.826, range 0.283–0.984; identical for all 624 specimens); only the purity-adjusted analyses of Section 3.2.3 of the main text used ABSOLUTE purity (inferred from copy-number data; R93). This transformation was calibrated on Affymetrix arrays, and its application to RNA-seq is an approximation. Higher values of this approximate purity indicate higher tumor purity. The purity-adjusted Cox regression in Section 3.3.2 also uses this approximate purity as a covariate, and wherever this manuscript writes ‘tumor purity (ESTIMATE)’ it refers to this approximate purity. The companion study of the 15q11-q13 locus (T. Tsukui, R. Yao, and K. Tsuda, unpublished observations) likewise uses this approximate purity as its covariate and records the ESTIMATE score alongside it; because the ESTIMATE score reflects the amount of stromal and immune components, higher values of that score indicate lower tumor purity. The Spearman rank correlation between the two measures is −1.000 per aliquot (−0.99992 after averaging and aggregating per specimen; companion study), and rank-based partial correlations give the same results with either measure, but caution is required in that scale-dependent estimates such as Cox regression reverse direction.

A note on the definition of stratification. The stratification used in this section (tertiles of the P2 signature score) is not identical to the stratification used in the companion study of the 15q11-q13 locus (T. Tsukui, R. Yao, and K. Tsuda, unpublished observations; tertiles of the GSVA difference P2 − P1). Although the rank correlation between the two axes is strong (Spearman ρ = +0.718), agreement of the tertile assignment is limited to 338 of 624 specimens (54.2%), and of the 208 specimens designated P2-high in this section, 132 are also P2-high in that study. While reversal between the two extremes is rare (1 specimen is high here and low in that study, and 9 the converse), the membership of the middle tertile differs considerably. Within each paper the stratification is internally consistent and this is not a numerical error, but when the two reports are read side by side, P2-high cannot be read as referring to the same set of specimens.
