## Supplementary figures and images for "TP53 status aligns with morphology and metaplastic programs in colorectal cancer organoids, and MDM2 P2 promoter usage reflects p53 activity"

### Additional_file_10_Figure_S1.pdf

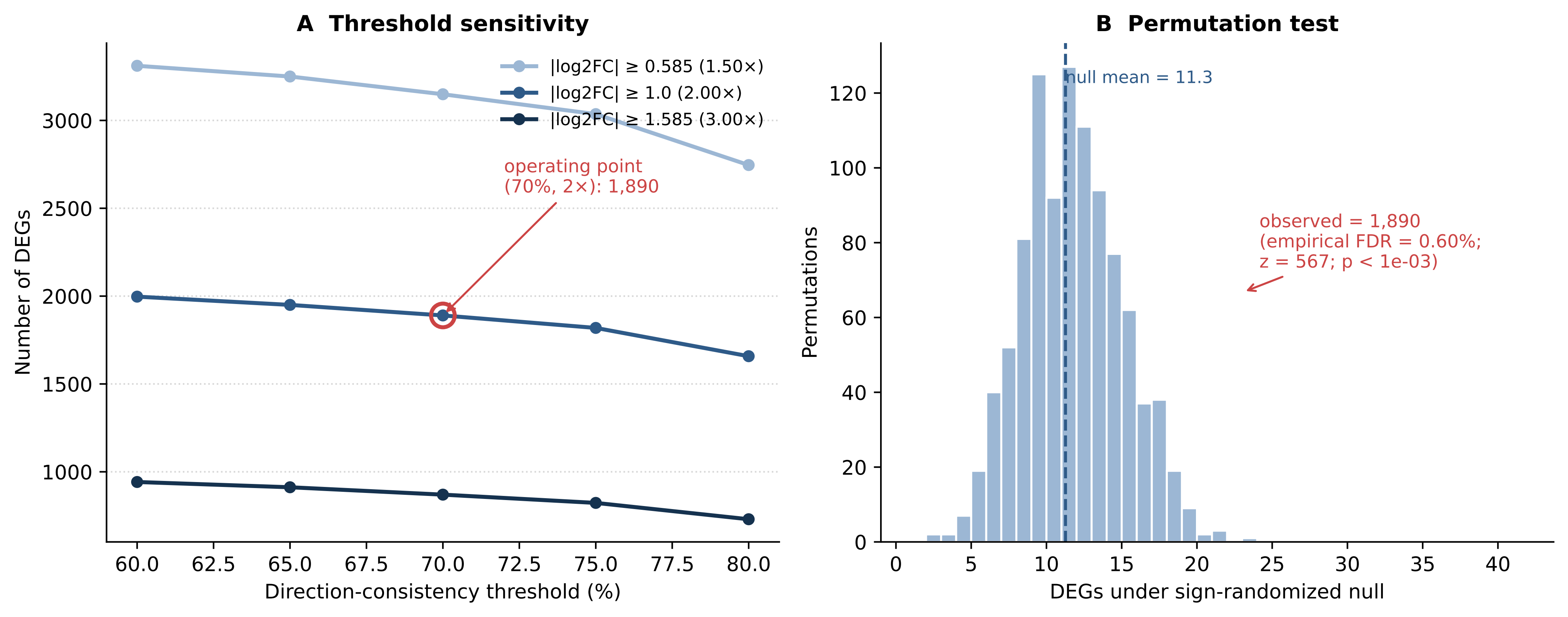

### Additional_file_11_Figure_S2.pdf

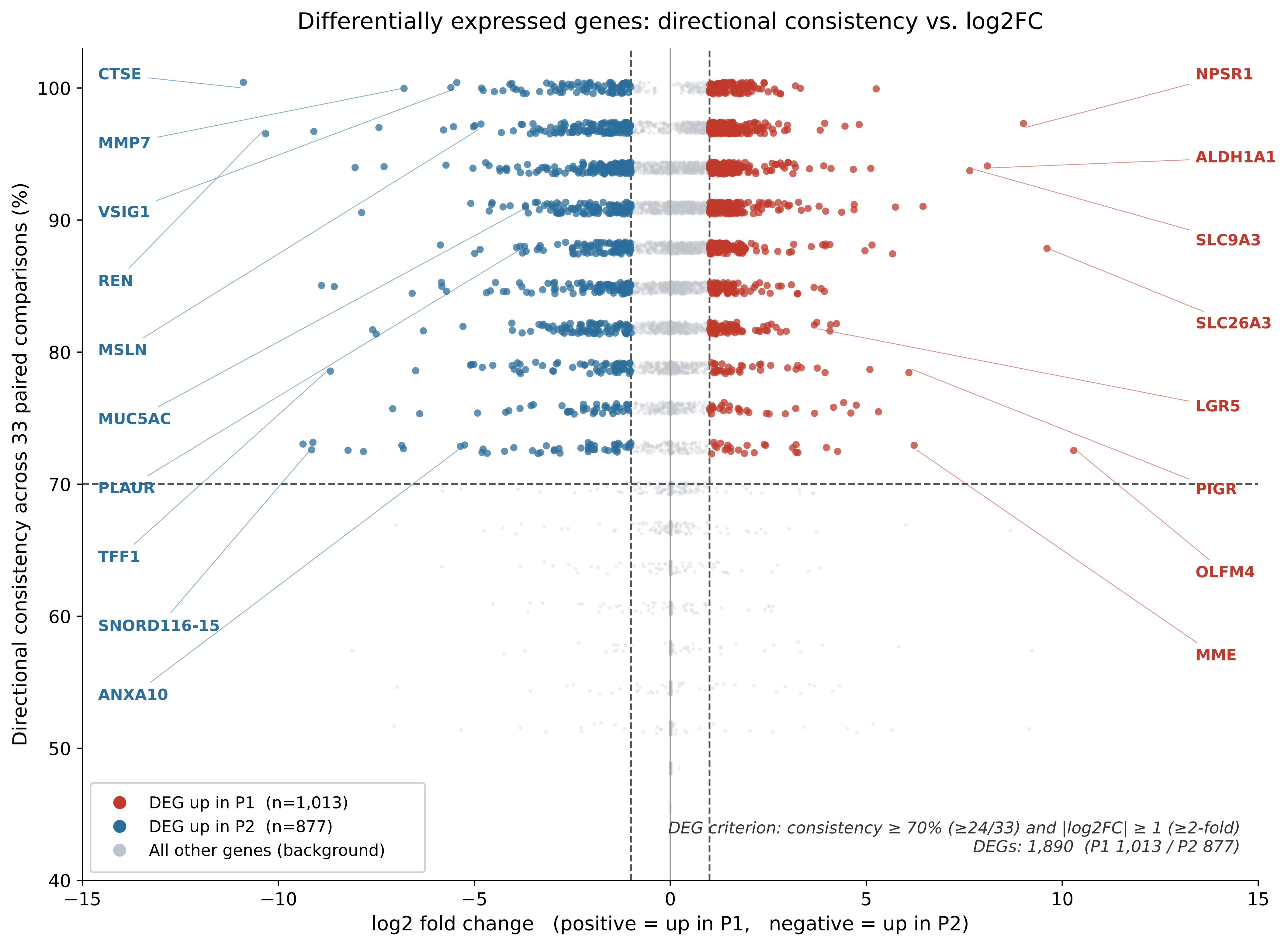

### Additional_file_22_Figure_S3.pdf

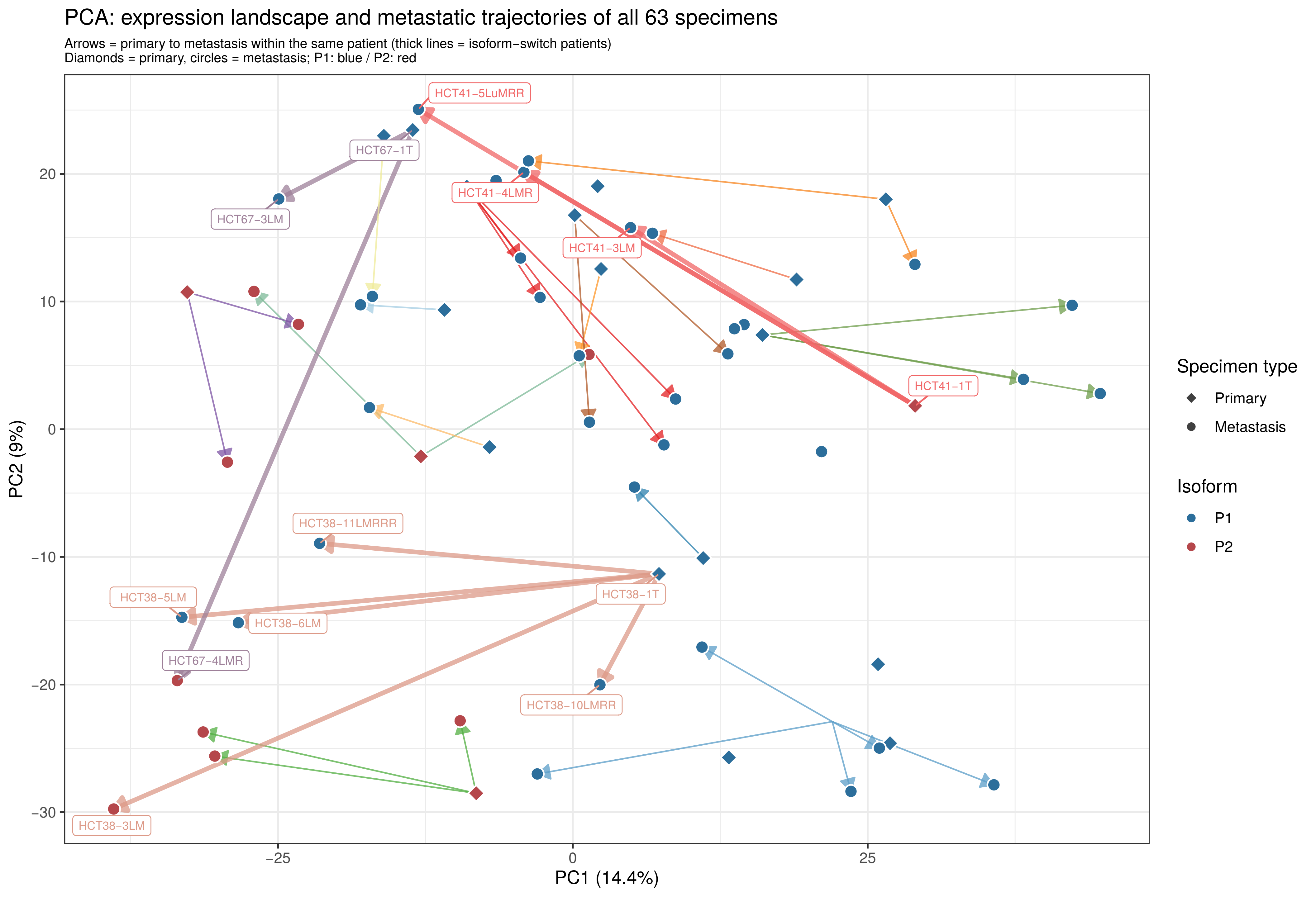

### Additional_file_23_Figure_S4.pdf

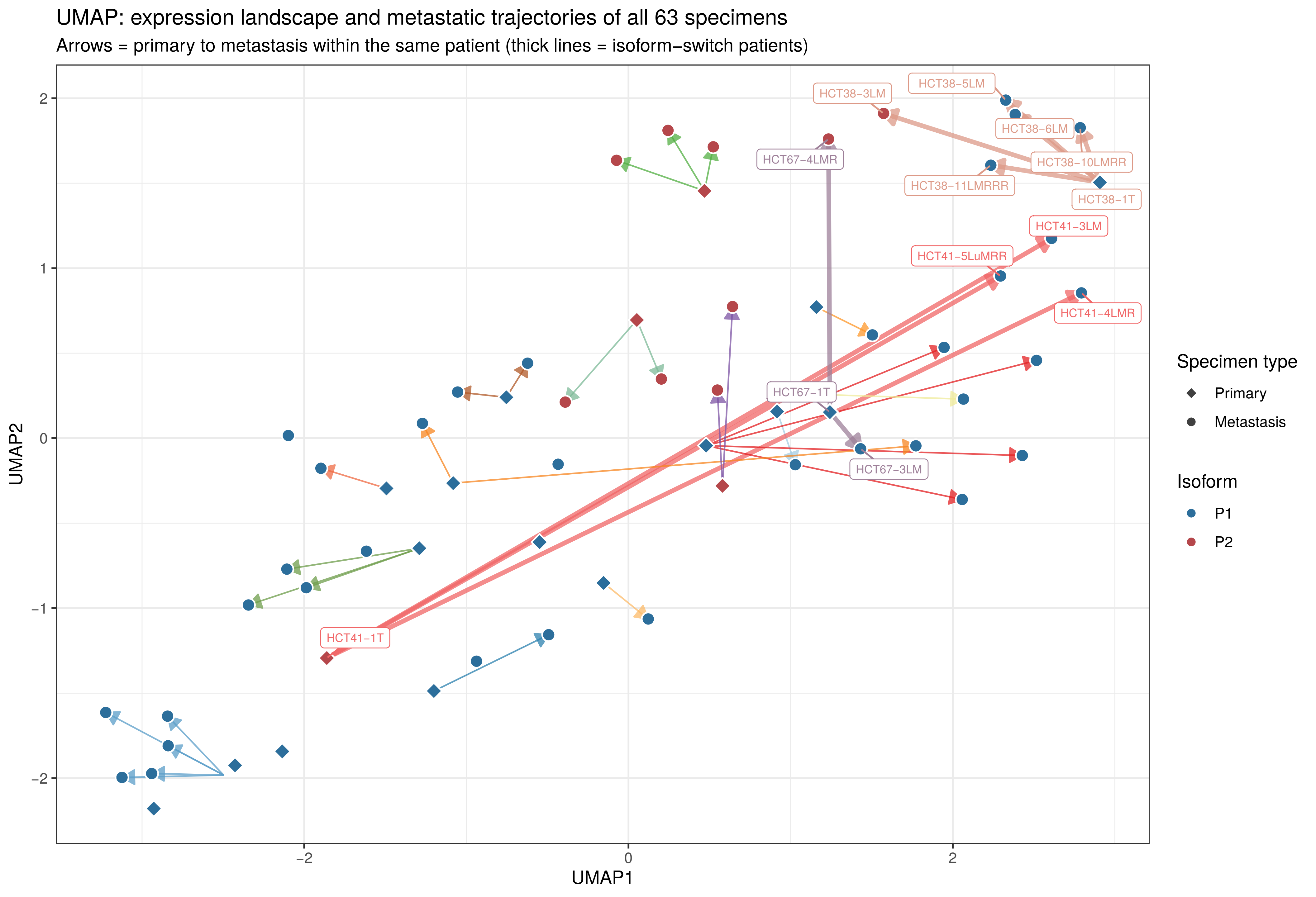

### Additional_file_43_Figure_S6.pdf

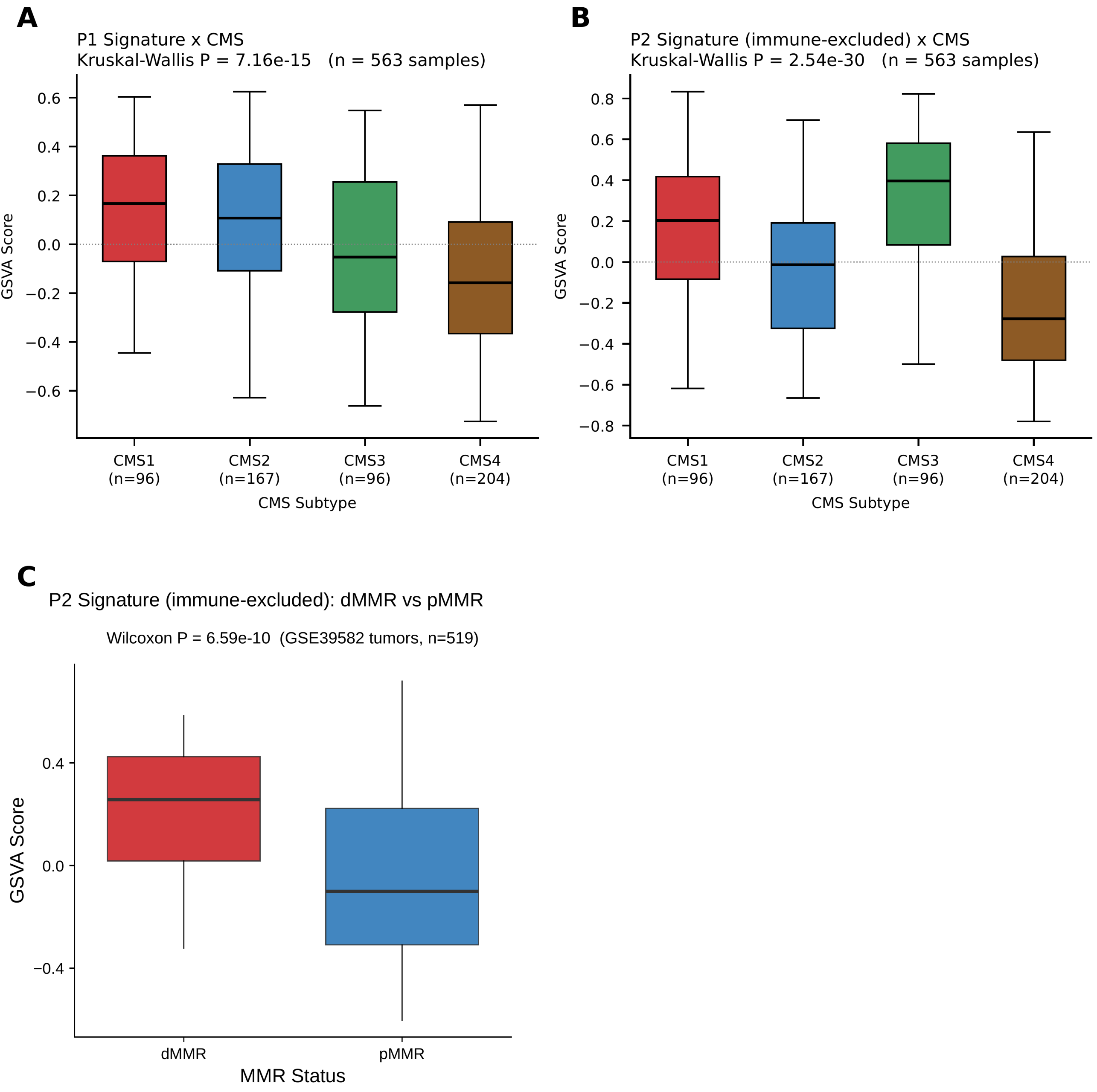

### Additional_file_44_Figure_S7.pdf

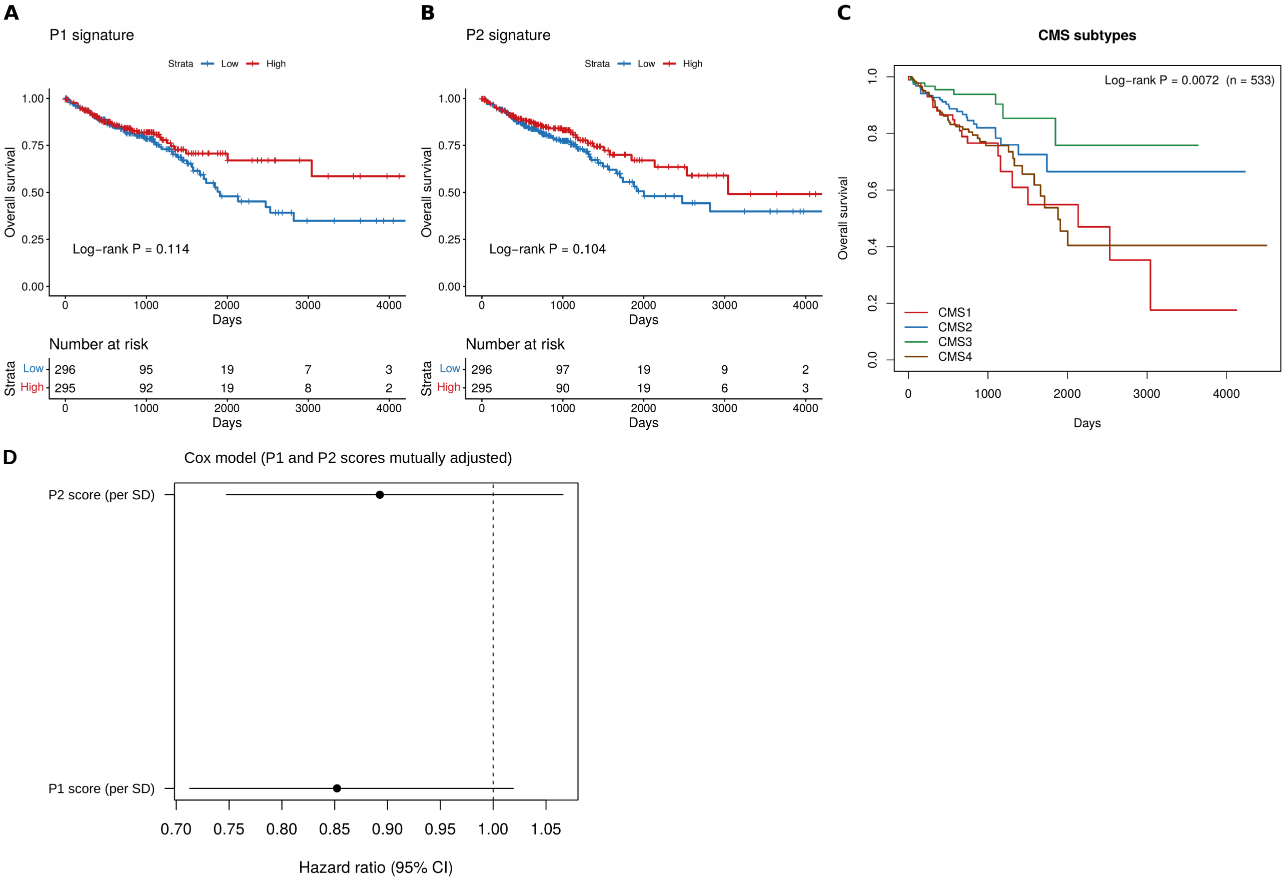

### Additional_file_49_Figure_S14.jpg

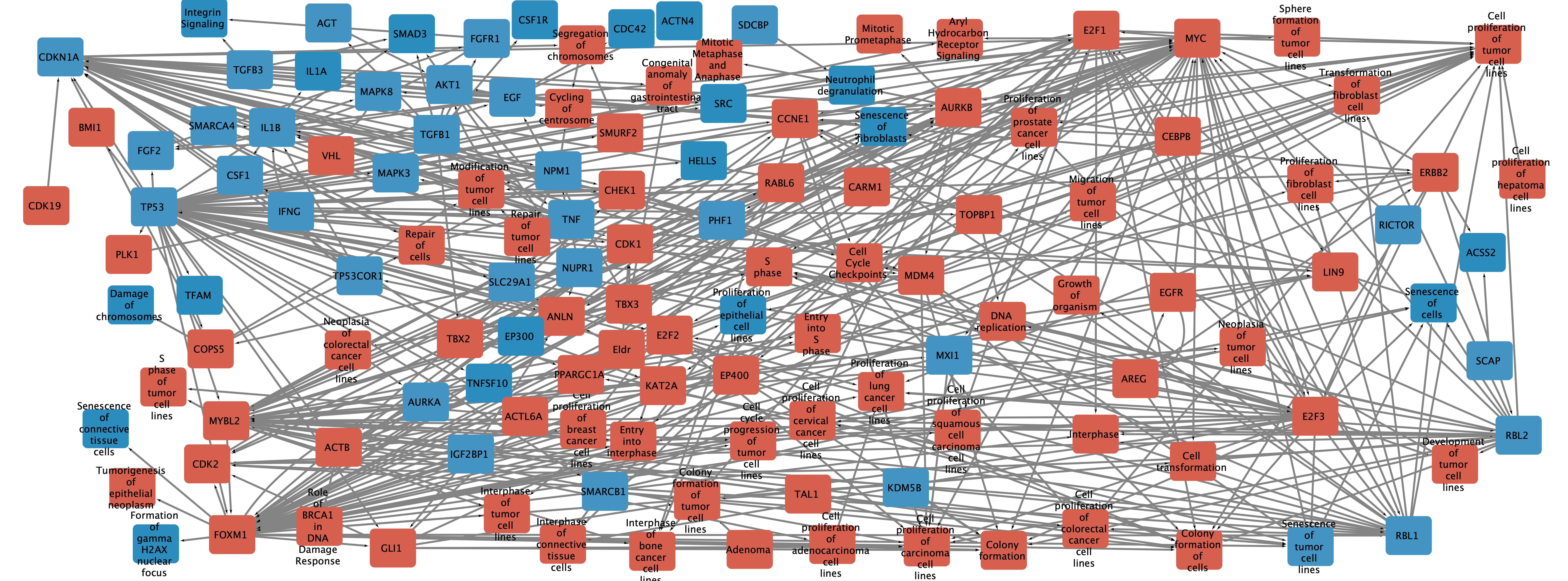

### Additional_file_52_Figure_S5.pdf

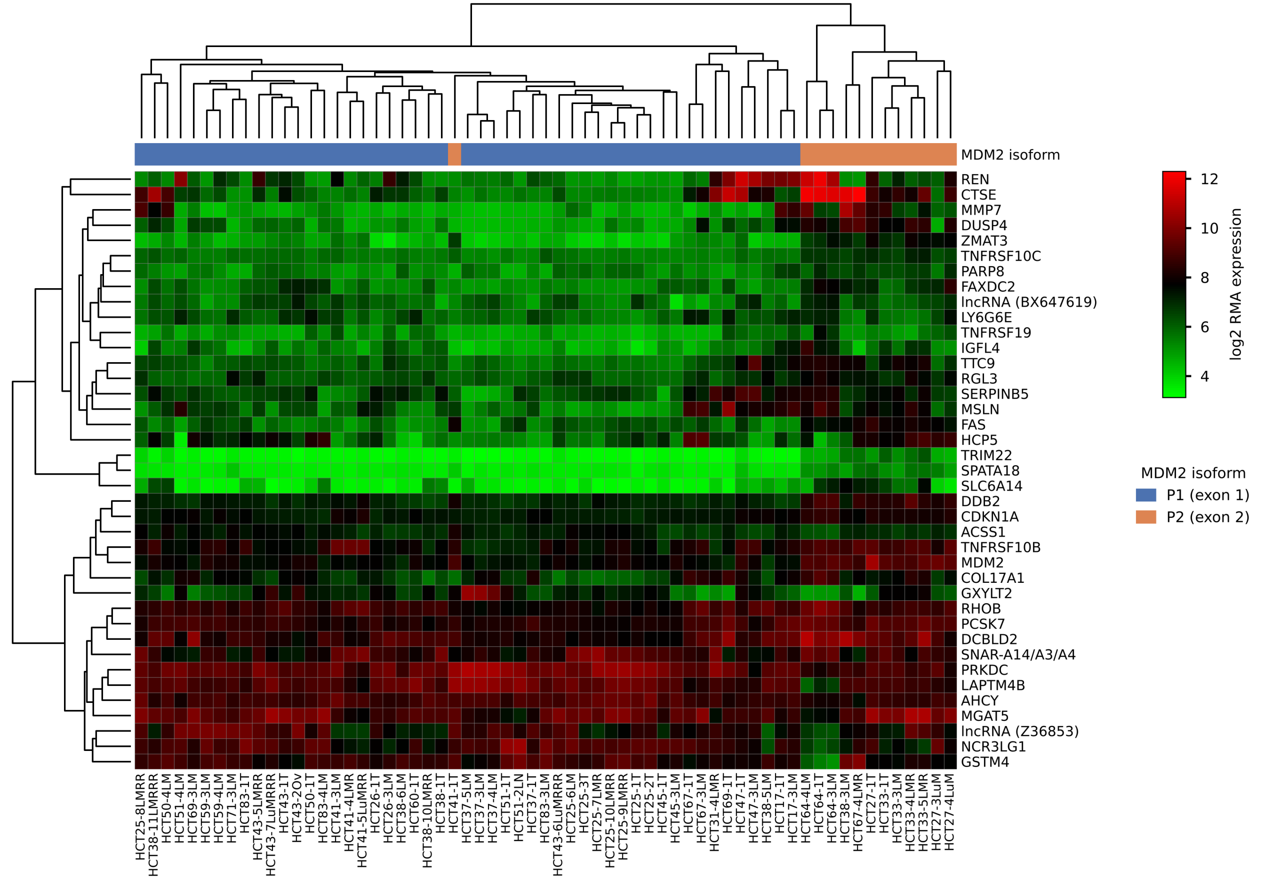

### Additional_file_62_Figure_S12.pdf

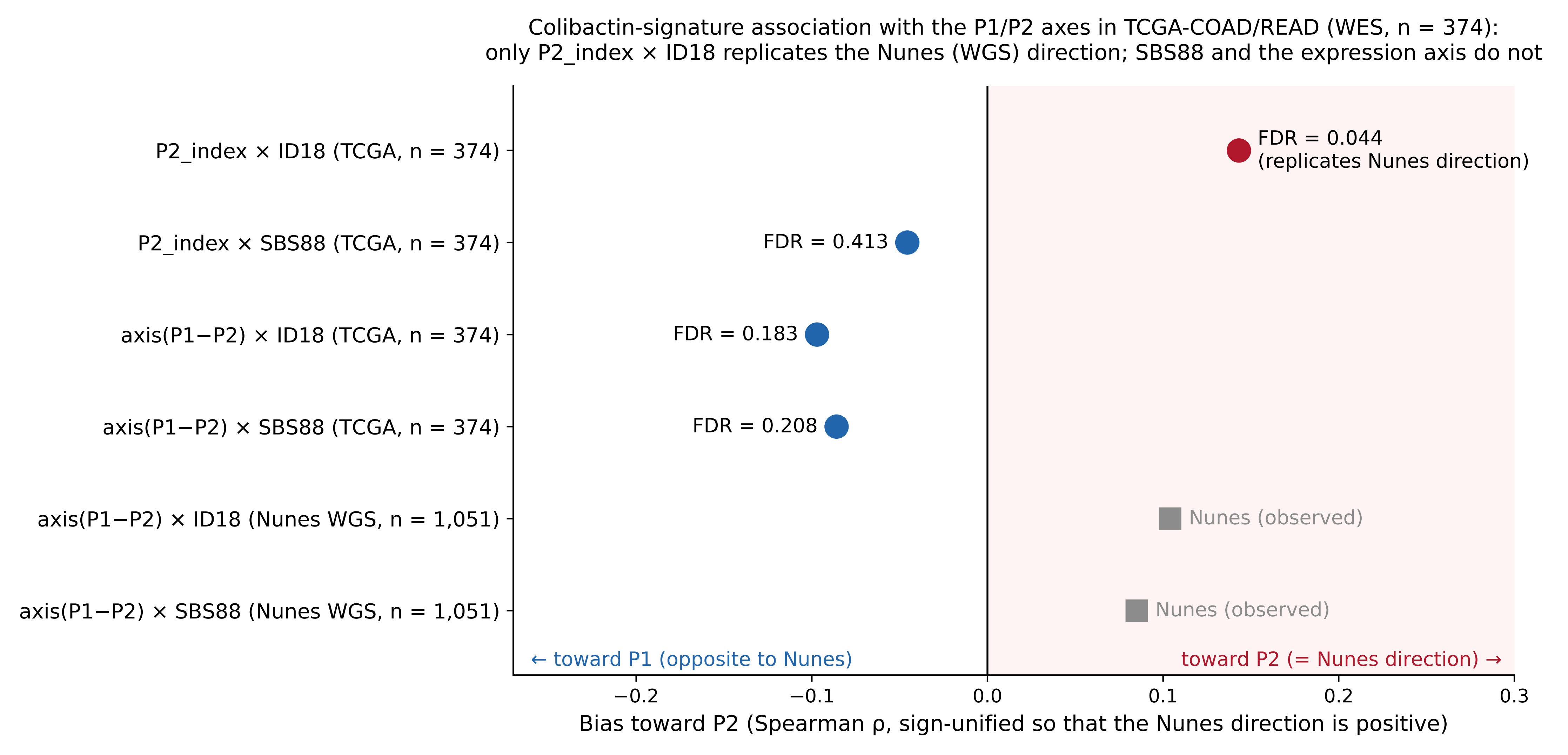
